# Functional diversification across yeast lineages of the nucleus-vacuole junction forming protein Nvj1

**DOI:** 10.64898/2026.09.05.749613

**Authors:** Florian Kramer, Jonathan Millen, Audrey M. Goldfarb, Jean-Claude Farre, John F. Wolters, Antonis Rokas, Chris Todd Hittinger, Suresh Subramani, Justin C. Fay, Michael Thumm, David S. Goldfarb

## Abstract

Nucleus-vacuole junctions (NVJs) in *Saccharomyces cerevisiae* serve as an inter-organellar hub for multiple cell processes, including lipid transport and biosynthesis, an intra-nuclear quality control mechanism, and piecemeal microautophagy of the nucleus. NVJs are formed by complexes between Vac8 in the vacuole membrane and Nvj1 in the nuclear envelope. Nvj1 also links the inner and outer nuclear membranes across the perinuclear lumen. Orthologs of Nvj1 had previously only been found in yeasts of the order *Saccharomycetales*, raising the possibility that NVJs are restricted to this clade. Using homology and synteny, we discovered scores of novel Nvj1 orthologs across 200 million years of evolution within the subphylum *Saccharomycotina.* Not all orthologs mediate NVJ formation when expressed in *S. cerevisiae,* and some lack a sequence motif necessary for this function. Furthermore, the sequence motif required for binding the oxysterol binding protein Osh1 and an associated novel sequence motif, both found only in the order *Saccharomycetales,* were independently lost in three different lineages. This collection suggests specific opportunities to explore evolutionary and functional adaptations of Nvj1s in ecologically and physiologically diverse yeasts.

## Introduction

Membrane contact sites (MCSs) are patches of protein-protein-mediated links between organellar membranes that coordinate cell signaling and homeostasis in response to growth, environmental, and nutritional cues (Voeltz et al., 2024; Cali et al., 2025). A broad theme among MCSs are their roles in nonvesicular lipid transport, metabolism, and membrane remodeling (Reinisch et al., 2021).

Nucleus-vacuole junctions (NVJs), described in *Saccharomyces cerevisiae,* are Velcro-like MCSs formed by complexes of Nvj1 in the nuclear envelope and Vac8 in the vacuolar membrane (Pan et al. 2000; Kvam and Goldfarb, 2006a; Kohler and Buttner, 2021; Hugenroth et al. 2025). Nvj1 is an integral endoplasmic reticulum (ER) membrane protein that, in addition to bridging to Vac8 in the cytoplasm, physically links the outer nuclear membrane (ONM or perinuclear ER) across the perinuclear lumen to the inner nuclear membrane (INM) via an N-terminal hydrophobic anchor sequence (Kvam and Goldfarb, 2006b; Millen et al., 2008). In addition to roles in lipid metabolism, NVJs also mediate piecemeal microautophagy of the nucleus (PMN) (Roberts et al. 2003; Kvam and Goldfarb, 2007; Tasnin et al. 2021; Li and Nakatogawa, 2022), which targets nonessential nuclear constituents to the vacuole, and mediate a nuclear protein quality control mechanism (Sontag et al. 2023).

Nvj1 expression is induced and NVJs expand during nutrient depletion and oxidative stress (Kvam and Goldfarb 2006a; Kohler and Buttner, 2021). In this regard, Snd3 is an especially interesting NVJ-associated protein (Tosal-Castano et al., 2021). Snd3, a Sec61 translocon-associated membrane insertase (Yang et al. 2025), redistributes from the bulk ER into NVJs during glucose exhaustion, and then back again when starved cells are fed glucose. Tosal-Castano et al. (2021) show that Snd3 is required for the formation and expansion of NVJs. Moreover, Snd3 depletion causes the rapid turnover of Nvj1 and disassembly of NVJs. Thus, Snd3 appears to play a key role in NVJ assembly, growth and disassembly in response to nutritional cues.

In addition to Vac8, at least 3 proteins with roles in lipid metabolism are recruited to NVJs by direct binding to Nvj1. These include Tsc13, a multi-pass ER-associated enzyme that functions in the biosynthesis of very long chain fatty acids (Kohlwein et al. 2001; Kvam et al. 2005), and Osh1, one of a seven-member family of yeast oxysterol binding proteins (Levine et al. 2001; Kvam et al. 2004). During log phase growth, Osh1 is distributed between Golgi membranes and NVJs, but beginning in late log phase, as nutrients become limiting and *NVJ1* expression is induced, it becomes increasingly sequestered within expanding NVJs (Kvam et al. 2004). Osh1 likely mediates the non-vesicular exchange between nuclear and vacuole membranes of ergosterol and phosphatidylinositol 4-phosphate (Manik et al. 2017; Nakatsu et al. 2021). A sequence within Nvj1 recruits HMG-CoA reductase into NVJs (Rogers et al. 2021), where it stimulates flux through the mevalonate biosynthetic pathway and promotes the biosynthesis of sterol esters.

The proteomes of MCSs shift in service of lipid homeostasis and involve the physiologically controlled shifting localization of various lipid-binding proteins among MCSs. This is especially true of NVJs. For example, the ergosterol binding protein Lam6 localizes to NVJs via an interaction with Vac8 during NVJ expansion, but can also be found at mitochondrial-ER and mitochondrial-vacuole MCSs (Elbaz-Alon et al., 2015). Yet3 localizes to NVJs and other MCSs, where it recruits the ERGosome, a multi-subunit complex containing ergosterol biosynthetic enzymes (Zung et al. 2024). The phosphatidic acid binding complex, Pex29/30, localizes to NVJs, peroxisomal and lipid droplet MCSs, where it regulates lipid homeostasis at multiple MCSs (Ferreira et al. 2021). Finally, the bulk lipid transporter Vps13 localizes to NVJs and ER-mitochondrial MCSs (Park et al. 2016; Leonzino et al. 2021).

Lipid droplets (LDs) are metabolically dynamic organelles that store triacylglycerides (TAGs) and sterol esters (SEs) (Zadoorian et al. 2023). Because they lack membrane bilayers, which precludes their fusion with membranes associated with the vesicular trafficking network, much of LD metabolism is mediated via interactions with MCSs (Renne and Hariri, 2021). LDs have a complicated and incompletely understood relationship with NVJs which, in addition to likely mediating the transfer of lipids into and out of lipid droplets, includes a role for NVJs in their biogenesis and macrolipophagy (Hariri et al. 2018; Diep et al. 2024). In response to nutrient starvation, Mdm1 links Tld-tagged LDs to the periphery of NVJs where they associate with fatty acyl-CoA synthetases for LD production and are simultaneously protected from lipolysis (Hariri et al. 2019; Speer et al. 2024). The microlipophagy of LDs requires an NVJ-adjacent MCS called vCLIP that links Ldo16/45-associated LDs to Vac8 (Alvarez-Guerra et al., 2024; Diep et al., 2024).

The nucleus is serviced by selective microautophagic (PMN) (Roberts et al. 2003; Kvam and Goldfarb, 2007; Otto and Thumm, 2021; Sakai and Oku, 2024) and Atg-39-mediated macroautophagic (Mochida et al. 2015; Mochida et al. 2022; Chandra et al. 2021) processes, both of which require core autophagy genes (Krick et al. 2008). During PMN small portions of NVJ-associated NE and underlying nucleoplasm are extruded into invaginations of the vacuole membrane (Roberts et al. 2003). PMN blebs containing NVJs pinch off and are released into the vacuole lumen by an unknown scission process, where they are digested by vacuolar hydrolases. Portions of the rough nucleolus, composed largely of pre-ribosomes, are selectively packaged into PMN blebs, while essential nuclear DNA, nuclear pore complexes and spindle pole bodies are excluded (Roberts et al., 2003). VAMP-associated proteins (VAPs), Scs2 and Scs22, localize throughout the ER and to NVJs, where they are required both for NVJ integrity and PMN (Manik et al. 2024).

The subphylum *Saccharomycotina* includes ecologically and metabolically diverse species that display a variety of cell shapes and morphologies, not all of which reproduce by budding (Groenewald et al. 2023; Chavez et al. 2024). The evolutionary and comparative functional biology of NVJs within this subphylum have both been understudied, in large part because Nvj1 orthologs have heretofore only been identified among species of the order *Saccharomycetales*, which includes *S. cerevisiae*. The reason for the restricted range was either that Nvj1s are indeed restricted to the *Saccharomycetales*, or, alternatively, Nvj1s are more widely conserved within the *Saccharomycotina*, but have escaped discovery due to a failure of homology detection (Weisman et al. 2020).

In this study we employed amino acid sequence-based similarity algorithms and gene order (synteny) to search for Nvj1 orthologs among more than 900 *Saccharomycotina* yeast genomes (Shen et al. 2018; Opulente et al. 2024). We conclude that the previous lack of identifiable orthologs across *Saccharomycotina* orders is due to the poor sequence conservation of Nvj1s. We compare and contrast the sequence properties and predicted structures of 137 candidate Nvj1 orthologs and demonstrate that many, but not all, mediate the *in vivo* formation of NVJs and PMN-structures. This collection of Nvj1s derives from physiologically and ecologically diverse species and provides investigators with a rich resource to investigate the comparative cell biology and functional diversity of a prototypic MCS-forming protein.

## Results

### Novel candidate Nvj1 orthologs

We found scores of novel Nvj1 orthologs encoded in 137 diverse yeast genomes (Table S1). The *Saccharomycetales*, of which *S. cerevisiae* is a member, comprise 17 genera in the family *Saccharomycetaceae* (Liu et al. 2024). We initially interrogated available *Saccharomycetaceae* genomes (Shen et al. 2018; Opulente et al. 2024) using BLAST searches with the *S. cerevisiae* sequence. This led to the identification of a number of novel likely orthologs encoded in pre- and post-whole genome duplication (WGD) clades (Fig. 1, Tables S1 & S2). BLAST searches failed to identify candidates in more divergent *Saccharomycetaceae* genomes. HMMER (hmmer.org) and gene order searches were then successfully used to extend the phylogenetic range of discoverable candidate orthologs outside the *Saccharomycetaceae*.

**Figure 1.**
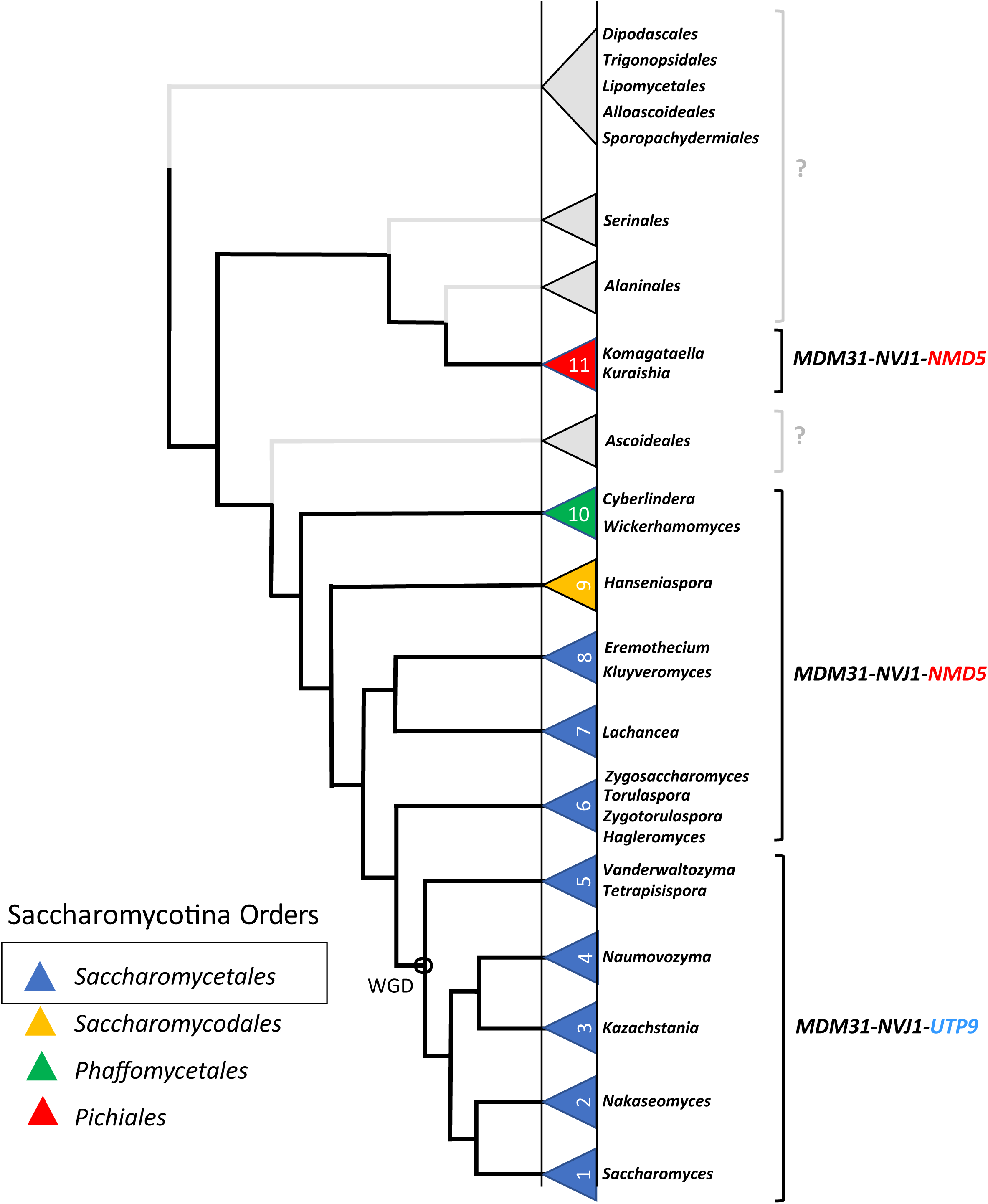
Phylogenetic distribution of Nvj1 orthologs. Clades where candidate orthologs were discovered are indicated by triangles colored according to their yeast orders. Clades from which we were unable to discover candidates are indicated by gray triangles labeled by their families. Gene orders on either side of the candidate genes are indicated by brackets. Note that the gene order changed in lineages post-WGD.

For simplicity, we divided relevant yeast clades into groups 1-11 (Fig. 1). Candidate orthologs from 12 genera of the *Saccharomycetaceae* family comprise groups 1-8. This family is further divided into non-WGD (groups 1-5) and post-WGD (groups 6-8) clades.

Candidates from the *Saccharomycodaceae* (*Hanseniaspora* species) comprise group 9, the *Phaffomycetaceae* (*Cyberlindnera* and *Wickerhamomyces* species) group 10, and the *Pichiaceae* (*Komagataella* and *Kuraishia* species) comprise group 11. Groups lacking candidate Nvj1 orthologs are left unnumbered in Fig. 1.

Outside the *Saccharomycetaceae* (groups 1-8), where BLAST and HMMER homology searches failed, gene order searches identified putative candidate Nvj1s within groups 9 and 10 (Table S2). Candidates from all discoverable non-WGD clades within and outside the *Saccharomycetaceae* are sandwiched between *MDM31* and *NMD5* genes (groups 6-11), whereas post-WGD orthologs map between *MDM31* and *UTP9* (groups 1-5). The loss of microsynteny in the genomes of family *Serinales* species, which include the human pathogen *Candida albicans*, precluded the discovery of candidate Nvj1s by gene order. Inspection of available gene order maps for the *Serinales* clade (http://cgob.ucd.ie) confirmed the lack of *NVJ1* candidate genes adjacent to *MDM31* and either *NMD5* or *UTP9* in these genomes. Surprisingly, candidate *NVJ1* genes were identified between *MDM31* and *NMD5* genes in several group 11 *Komagataella* (family *Pichiaceae*) genomes and, by BLAST to the *Ko. phaffii* (previously *Pichia pastoris*) protein sequence, and a candidate ortholog in a sister genus *Kuraishia floccula*. For convenience, *Hagleromyces aurorensis* and *Kuraishia floccosa candidates,* which might be grouped in their own clades, are instead included with their sister clades groups 6 and 11, respectively. *MDM31-NMD5* microsynteny is not conserved in *Sporopachydermia*, *Alloascoidaceae*, *Dipodascaceae*/*Trichomonascaceae*, *Trigonopsidaceae* and *Lipomycetaceae* families, precluding efforts to identify orthologs within these genomes.

Identity matrices comparing the sequences of orthologs from examples from each of the 11 groups are displayed in Fig. S1. These data reveal significant divergence of these protein sequences even within most genera.

Gene order is an exceptional tool for identifying orthologs when homology searches fail, even when gene order is imperfectly conserved. Group 10 candidate orthologs are sandwiched between *MDM31* and *NMD5.* However, in six of these genomes a paralog of *AMD2,* which encodes an amidase of unknown function, has been inserted immediately downstream of *NVJ1*. Moreover, in these genomes the orientation of the *NVJ1-AMD2* pair is flipped relative to the reading frame of *MDM31*. These *AMD2* paralogs are found adjacent to *NVJ1* candidates in the genomes of group 10 species *Starmera quercuum* and *Candida stellimalicola*. Thus, the insertion of *AMD2* paralogs adjacent to *NVJ1* candidate genes and their flipped orientation relative to *MDM31* and *NMD5* is ancestral within the group 10 lineage.

Another variation in gene order occurs in group 11 *Komagataella* genomes, where the gene order is *MDM31*-*ORF*-*NVJ1*-*NMD5*. This ORF encodes a short, reasonably well-conserved, polypeptide of uncharacterized function. The *MDM31-NMD5* gene order is not conserved in most other known group 11 genomes, although it does occur in the *Citeromyces hawaiiensis* genome, but without an intervening *NVJ1*-like gene or any other ORF.

In conclusion, we have compiled a list of 137 candidate Nvj1 protein sequences discovered by BLAST, HMMER and/or gene order (Table S1).

### Sequence characteristics of candidate Nvj1 proteins

We sought to identify shared characteristics based on the assumption that bona fide orthologs will abide to the principle that form follows function. In doing so, among such a diverse set of polypeptides, we hope to gain insight into the evolutionary dynamics—the gain and loss of functions—of Nvj1-mediated cellular processes. *S. cerevisiae* Nvj1 is divided into N- and C-terminal domains separated by a single transmembrane domain (TMD). The N-terminal domain localizes to the perinuclear lumen, while the C-terminal domain is exposed to the cytosol. All candidate Nvj1s harbor single hydrophobic sequences predicted to function as TMDs (Fig. S2, 2 and Table S2), while most, but not all, also contain N-terminus-adjacent hydrophobic sequences consistent with their being INM anchors (Fig. S2 & Table S2). The 137 candidates vary in length between 206-513 amino acids with commensurate masses (Fig. S2 & S3A).

To a first approximation, most of the length variability among candidates is due to variability of the lengths of their C-terminal domains. With exceptions, candidates from groups 1-7 & 11 have N-terminal domains of ∼90 aa. This makes sense, since N-terminal domains of NVJ-forming Nvj1s need to be long enough to span the width of the perinuclear lumen, which is clamped at ∼8.6 nm within NVJs (Millen et al. 2008).

Candidates belonging to groups 8 & 9 show atypical variability in the lengths of their N-terminal domains (Fig. S2). Group 9, the *Hanseniaspora*, are atypical in that they contain only one hydrophobic motif consistent with a membrane spanning function very near their N-termini. If these were not the only putative TMD-like sequences, for example, if a second TMD-like motif within the body of the protein was also present, these might have appropriately been designated INM anchors. One reason NVJs are not formed by group 9 orthologs is the apparent presence of only single INM anchor/TMD-like sequences in this clade (see below).

Isoelectric points (pI) are known to influence protein 3D structures, cellular localizations, and interactions with other biomolecules (Tokmakov et al. 2021). The pI of the *S. cerevisiae* proteome reflects a biphasic distribution with an average pI = 6.7 (Kozlowski, 2022). *S. cerevisiae* Nvj1 has a relatively strong acidic pI = 4.8, as do candidates from most of the groups (Fig. S3B), largely due to an abundance of aspartates and glutamates in their C-terminal domains (Fig. S2). The pIs of candidate Nvj1s within groups 7 and 8, which together comprise a clade within the *Saccharomycetaceae*, are more diverse, and include a number with basic pI’s (Fig. S3B). Note that the *K. africana* candidate (group 3) is unique among its cohort in having a basic pI = 9.5, owing in large part to a unique stretch of mostly basic residues in the C-terminal domain (KRKTKKNKRKEKKNSNKKTKR). There is no correlation between the lengths of Nvj1 candidates and their pIs.

In conclusion, with exceptions, the domain organization of our candidates—a single TMD separating N- and C-domains—and an overall acidic amino acid composition is conserved in yeast lineages spanning over 200 million years of evolution. Consistent with clade-specific variations, it is noteworthy that variations in pI and domain organization do not correlate directly with phylogenetic relationships. For example, the locations of the TMDs and the preponderance of acidic residues are more similar when comparing candidates belonging to groups 1 and 11 than groups 1 and 8.

### Conservation of consensus sequence motifs

*S. cerevisiae* Nvj1 harbors two types of known functional sequences: topological signals (INM anchor and TMD) and discrete partner binding sequences (e.g. Osh1 and Vac8). Sequences that are sufficient and necessary for binding to Vac8, Osh1, and Tsc13 have been mapped (Kvam and Goldfarb, 2006), while a sequence within the N-terminal domain is necessary for binding Hmg1 (Rogers et al. 2021). The mapping and distribution of sequence motifs among our candidates was explored using agnostic MEME motif search algorithms (https://meme-suite.org/meme/) (Fig. 2, Table S2). Whereas Osh1-binding motifs are restricted to groups 1, 4, 6-8 (Fig. 2 & 3), a consensus Vac8-binding motif is more broadly conserved among all but groups 9 and 11. The absence of consensus factor binding motifs does not mean that the factors do not bind. A good example is that of the *Ko. phaffii* Nvj1 ortholog, which lacks a consensus Vac8-binding motif but still forms *VAC8*-dependent NVJs (below). Either the Vac8 binding motif has diverged in these proteins, or, perhaps, another protein links Nvj1 to vacuoles in these cells.

**Figure 2.**
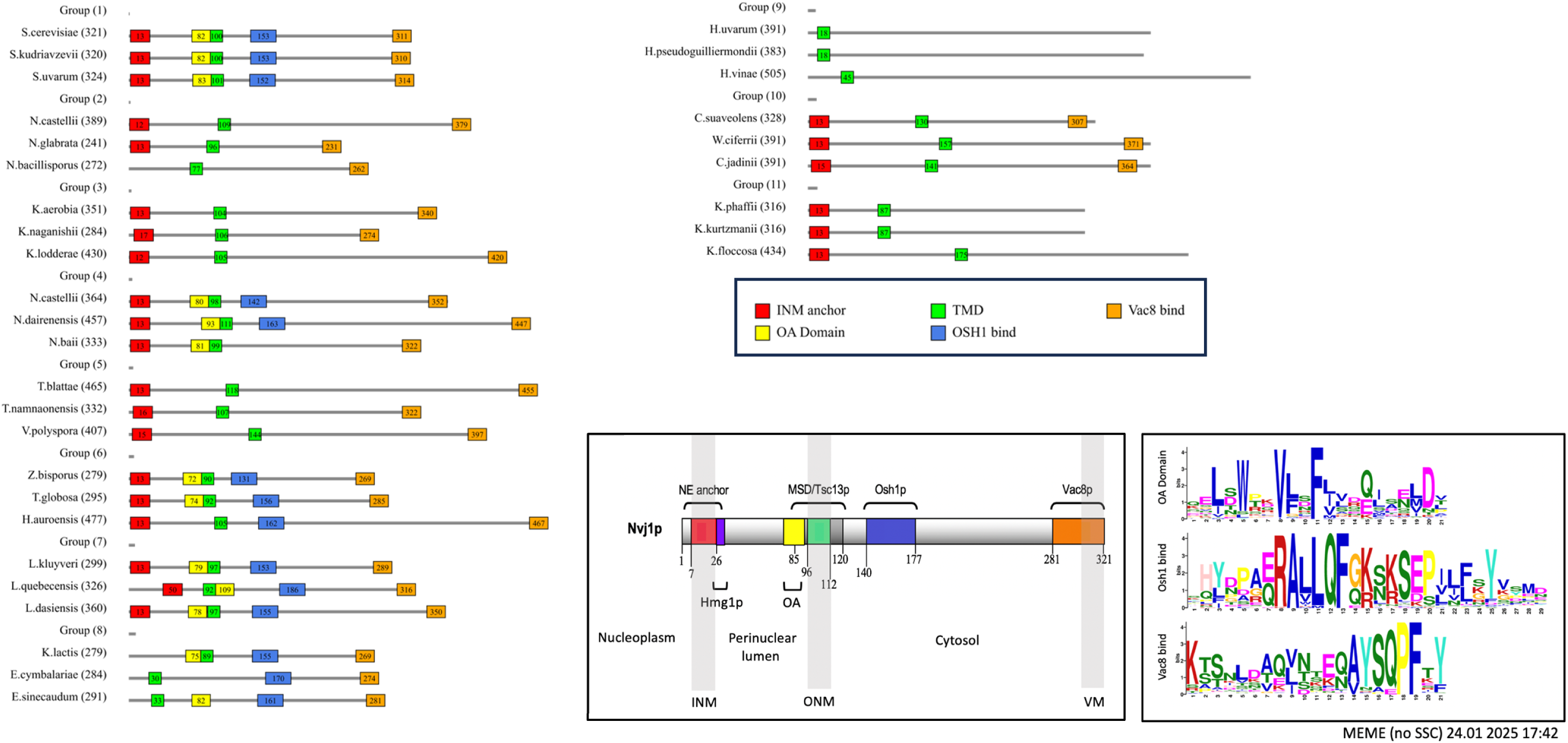
Domain organization and motifs within Nvj1 proteins from each of 11 groups. (**A**) Three candidates from each group display intragroup variations. Assignment of topogenic and sequence motifs are described in the text. (**B**) Box diagram showing the domain organization of *S. cerevisiae* Nvj1. (**C**) Consensus motifs for Osh1-binding, OA, and Vac8-binding motifs.

Meme searches identified a novel “Osh1-associated” (OA) consensus sequence that we had previously overlooked. OA motifs are associated almost exclusively with candidates from group 1,4, 6-8 that also harbor Osh1 binding motifs (Figs. 2 & 3). OA motifs are located within N-terminal perinuclear domains adjacent TMDs, which are on the other side of the ER membrane from Osh1-binding motifs (Fig. 2). The most parsimonious explanation for this punctuated distribution is that the OA and Osh1 binding motifs were coordinately lost in three separate events during the diversification of the *Saccharomycetaceae* family lineage (Fig. 3).

**Figure 3.**
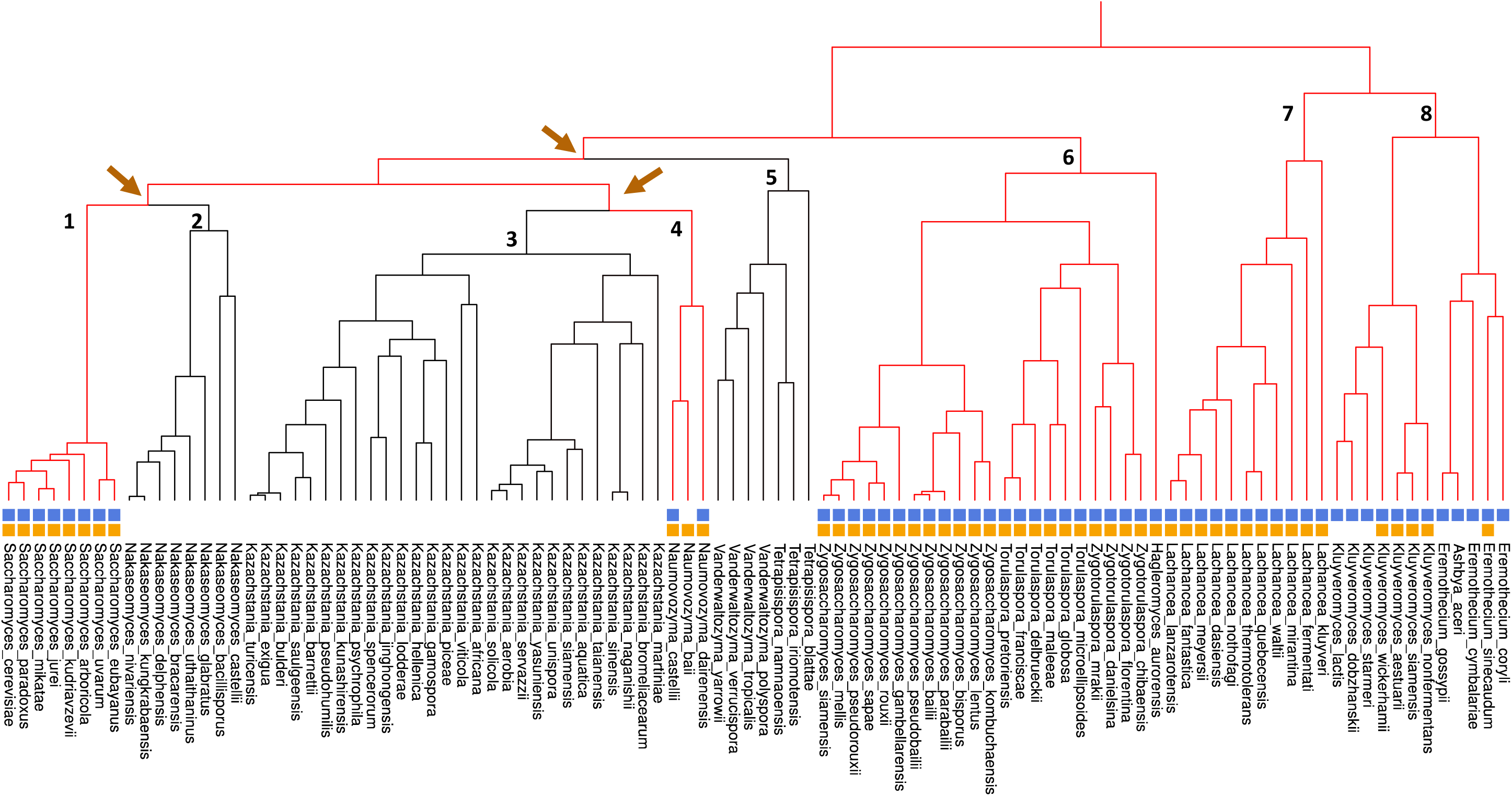
Osh1-binding and OA-motifs were independently lost three times during the diversification of *Saccharomycetaceae* lineages. Clades are identified at their branch points by group numbers (1-8). Presence of Osh1-binding (solid blue squares) and OA-associated motifs (solid orange squares) are indicated for individual species. Branch points where OA motifs were lost are indicated by a switch from red to black lineage lines.

Meme searches also predicted putative INM anchor and TMD sequence motifs in most candidates. Because INM anchors and TMDs are characterized more by their overall hydrophobic properties than by any particular sequence motif, we included likely TMDs which MEME failed to identify but which were predicted by a TMD predictor algorithm (https://services.healthtech.dtu.dk/services/TMHMM-2.0/). The *S. cerevisiae* INM anchor extends across the perinuclear lumen and physically associates with the INM (Millen et al., 2008). As a result, INM anchor sequences likely occur at or very near the N-termini of candidates. The characterization of INM anchors as having properties associated with TMDs is consistent with a mechanism of anchoring involving the direct insertion of these sequences into the INM bilayer.

Group 9 candidates (*Hanseniaspora*) are the most atypical in that they exhibit few of the characteristics that are shared by bona fide orthologs. They lack all the consensus motifs. They do share their syntenic location, contain putative TMDs, albeit very near their N-termini, and have acidic pI’s. Group 11 candidates are the most evolutionarily divergent. These contain putative INM anchors, TMDs and acidic pIs, and have N-terminal domains of lengths similar to *S. cerevisiae* Nvj1, but they lack consensus Vac8-binding motifs (see NVJ-forming capacity below).

### Predicted secondary and tertiary structures

Short of biophysical structural determinations, tentative insight into the structural properties of candidates can be obtained using prediction algorithms. AIPred (Manavalan et al., 2018) predicts that large tracts of most Nvj1 candidates are disordered, especially within their C-terminal domains (Fig. S4). Intrinsically disordered protein sequences typically contain an abundance of polar and charged residues that preclude the formation of globular domains. The prevalence of acidic over basic clusters of amino acids within the putative disordered domains of Nvj1 candidates (above) may have functional implications (Bigman et al., 2022). The presence of large segments of disordered sequence in most candidates is supported by AlphaFold 3D structural predictions (Fig. S5), albeit see (Ruff and Pappu, 2021). Disordered domains can serve as scaffolds that promote protein partner binding (Holehouse and Kragelund, 2024), a function that aligns well with the fact that the largely disordered C-terminal domain of *S. cerevisiae* Nvj1 associates with known binding partners. It is noteworthy that Osh1, OA, and TMD sequences, where they occur in these structures, tend to lie within segments that are predicted by AlphaFold to be alpha-helical (Fig. S5).

In conclusion, most candidate Nvj1s, except all of group 9 and some group 8 members, are predicted to contain large stretches of intrinsically disordered sequence. The presence of disordered sequences, which are characteristically poorly constrained, explaining in large part why swaths of candidate Nvj1 sequences from within and between genera are dissimilar (Fig. S1).

### Nvj1 candidates direct the formation of NVJs in *S. cerevisiae*

We next wished to test whether *in vivo* functions of these orthologs is conserved. We tested the capacity of seven candidate Nvj1s from groups 1, 2, 3, 9, 10 & 11 to mediate the formation of *VAC8*-dependent NVJs in *S. cerevisiae*, a hallmark of Nvj1 function. These seven genes were expressed in *S. cerevisiae* as C-terminal GFP-tagged reporters in *VAC8^+^ nvj1Δ* or *vac8Δ nvj1Δ*. All seven sequences contain predicted TMDs, and all but the *Han. uvarum* (group 9) and *Ko. phaffii* (group 11) sequences contain consensus Vac8-binding motifs. None but the *S. cerevisiae* protein contains Osh1-binding and OA motifs.

As shown in Fig. 4, all but the *Han. uvarum* reporter formed discrete *VAC8*-dependent NVJs. The *Han. uvarum* Nvj1 appears to localize throughout the peripheral and perinuclear ER without any proclivity to interact with Vac8-associated vacuoles. The inability of the *Han. uvarum* candidate to direct NVJ formation in *S. cerevisiae* can be explained three different ways. First, the gene might be unrelated to *NVJ1*, despite its positioning between *MDM31* and *NMD5*, its ER membrane protein localization and an acidic pI. Second, the gene could be orthologous to *NVJ1* but might have diverged either by drift or selection and lost the elements needed to form NVJs. Third, the protein might mediate the formation of NVJs in *Han. uvarum* cells but not in *S. cerevisiae*.

**Figure 4.**
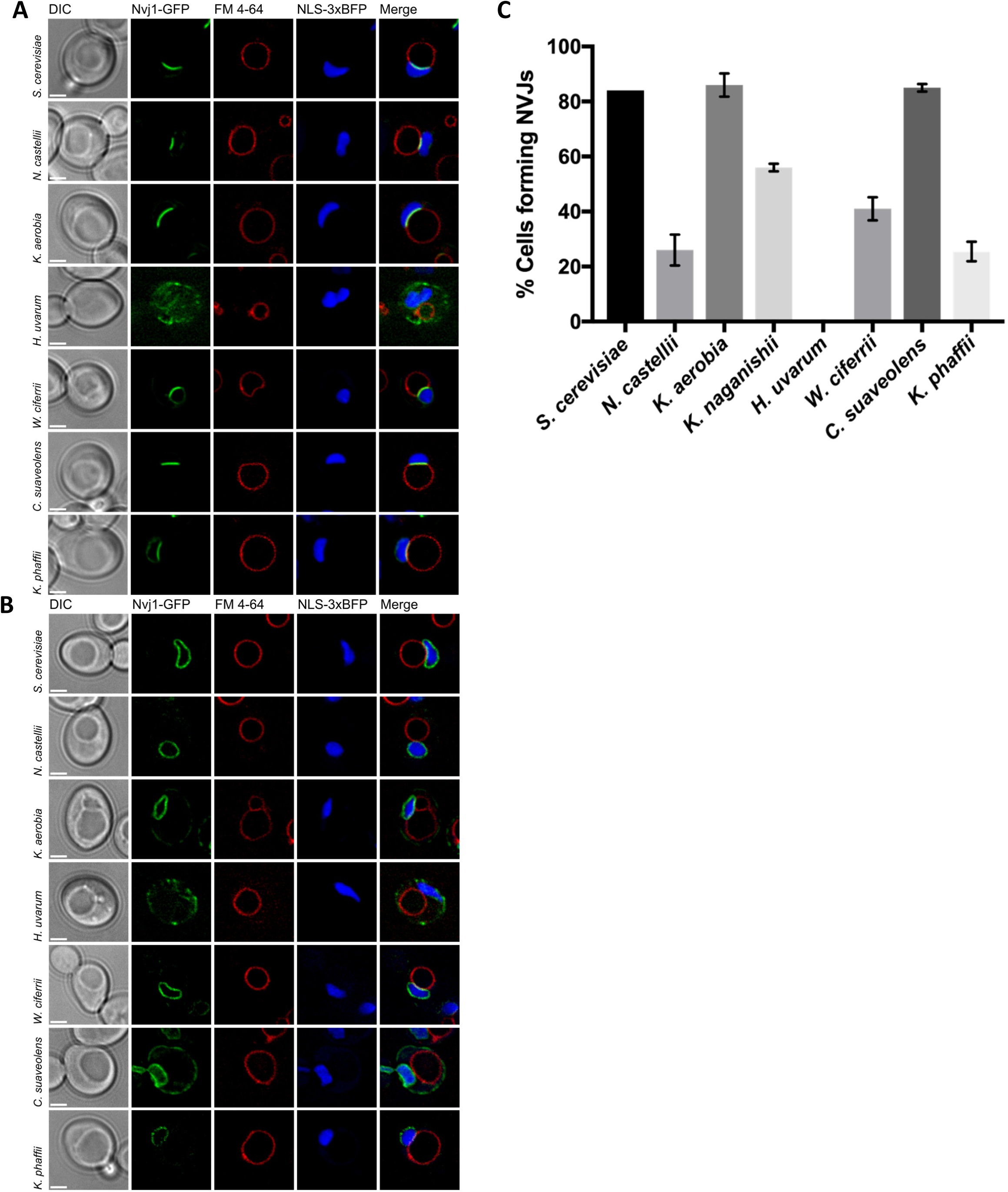
Formation of *VAC8*-dependent NVJs in *S. cerevisiae*. (**A**) Candidate orthologs expressed in *nvj1Δ VAC8*, and (**B**) *nvj1Δ vac8Δ S. cerevisiae* cells (strain WCG). (**C**) Frequency of NVJ formation from two independent biological trials of 200 cells each. Results from both *K. aerobia* and *K. naganishii* are shown. Cells were grown overnight to stationary phase in CM medium. C-terminal GFP fusions of candidate Nvj1s were expressed from the *Met17* promoter induced with 0.3 mM methionine. Strains were transformed with Nab2-NLS-3xBFP to mark the nucleoplasm. Vacuoles were stained with FM 4-64. (Scale bar: 2μm)

Two group 10 candidates (*W. ciferrii* and *Cy. suaveolens*) directed the formation of both discrete NVJs (Fig. 4) and ectopic ER-vacuole junctions (Fig. S6). Ectopic ER-vacuole junctions were previously observed when *S. cerevisiae* Nvj1 reporters with mutated INM anchor sequences were expressed in *S. cerevisiae* cells (Millen et al. 2008). It is instructive that both of the group 10 candidates we looked at lack predicted INM anchor sequences, but still formed NVJs in *S. cerevisiae*. Thus, the motif prediction programs we used are not robust predictors of whether or not a particular functional sequence occurs in a candidate ortholog.

Along these lines, the group 11 *Ko. phaffii* candidate lacks a consensus Vac8-binding motif but still mediates the *VAC8*-dependent formation of NVJs in both *S. cerevisiae* and *Ko. phaffii* cells (Figs. 4 & 5). Based on the formation of *VAC8*-dependent NVJs, the most likely conclusion is that *Ko. phaffii* Nvj1 contains a functional but divergent Vac8-binding sequence.

**Figure 5.**
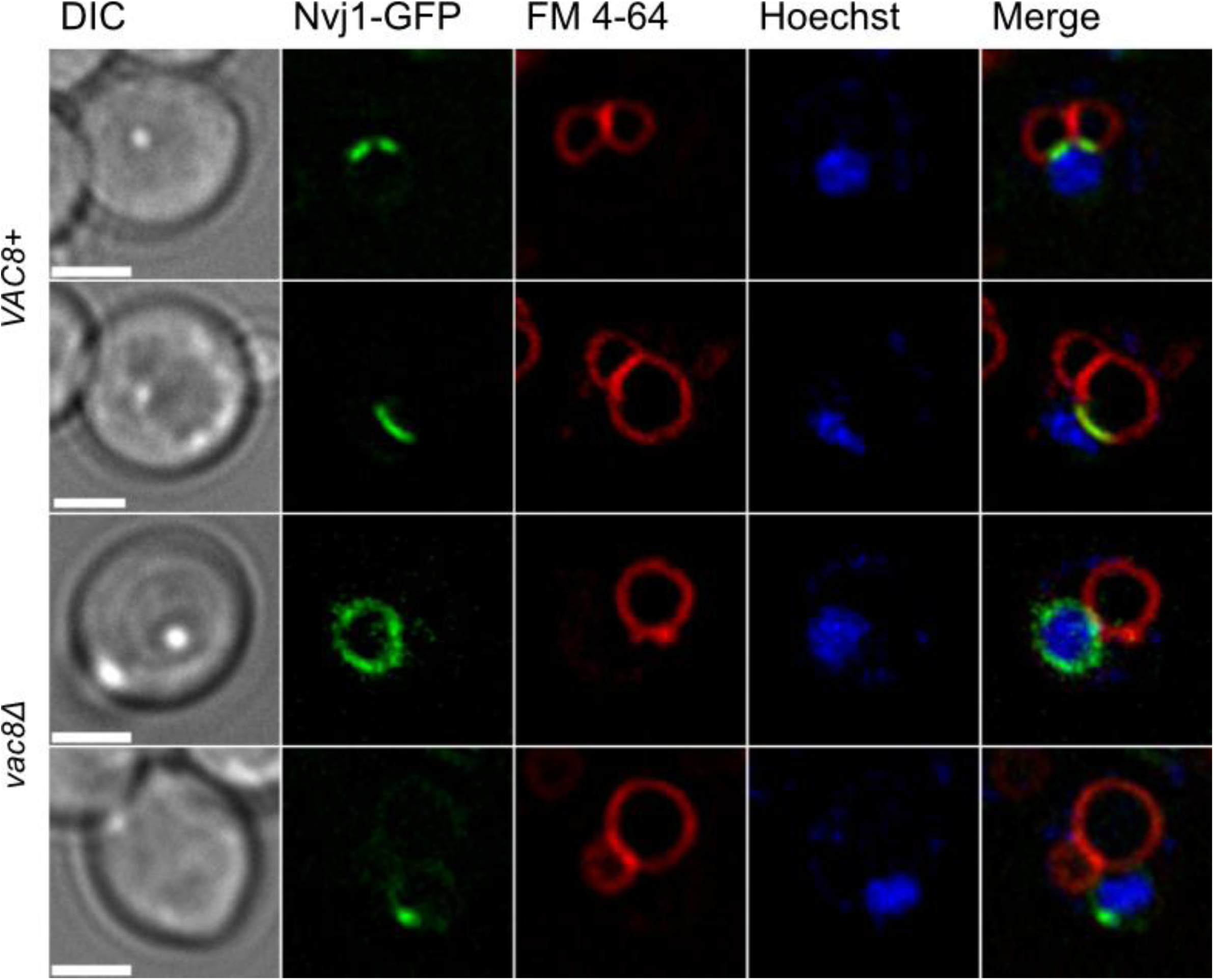
The *Ko. phaffii* Nvj1 candidate forms *VAC8*-dependent NVJs in *Ko. phaffii* cells. Integrated GFP-tagged *Ko. phaffii* Nvj1 candidate was expressed in *nvj1Δ VAC8* and *nvj1Δ vac8Δ Ko. phaffii* PPY12h cells. Cells were grown in CM media to stationary phase. Vacuoles were stained with FM 4-64 and nuclei with Hoechst-33342. The shown phenotype was present in 90% of observed cells. (Scale bar: 2μm)

### Divergent orthologs mediate the formation of PMN-like structures

Another striking phenomenon mediated by Nvj1 is PMN. Bona fide PMN structures characteristically contain nucleoplasm, often including pre-ribosomes (rough nucleolus). No other cellular process in any (non-*Saccharomycotina*) organism is known to produce these striking structures. We asked whether our candidates could mediate PMN when expressed in nitrogen-starved *S. cerevisiae* cells. As shown in Fig. 6, PMN-like structures appeared in nitrogen-starved *S. cerevisiae* cells expressing *S. cerevisiae*, *Nak. castellii*, *K. aerobia*, *W. ciferrii*, and *Cy. suaveolens* GFP-tagged Nvj1 reporters. These intravacuolar structures are indistinguishable from bona fide PMN blebs and vesicles in that they stained with Nvj1-GFP (nuclear envelope), FM4-64 (vacuole membrane), and NLS-3XBFP (nucleoplasm). We were unable to find PMN-like structures when expressing *Ko. phaffii* Nvj1-GFP in either S*. cerevisiae* or *Ko. phaffii* cells. We cannot rule out the possibility that *Ko. phaffii* Nvj1 is capable of mediating PMN, but that they occur at undetectably low frequencies or they are too small to visualize by fluorescence microscopy. PMN in *Ko. phaffii* could be induced by different nutritional or environmental cues than we used.

**Figure 6.**
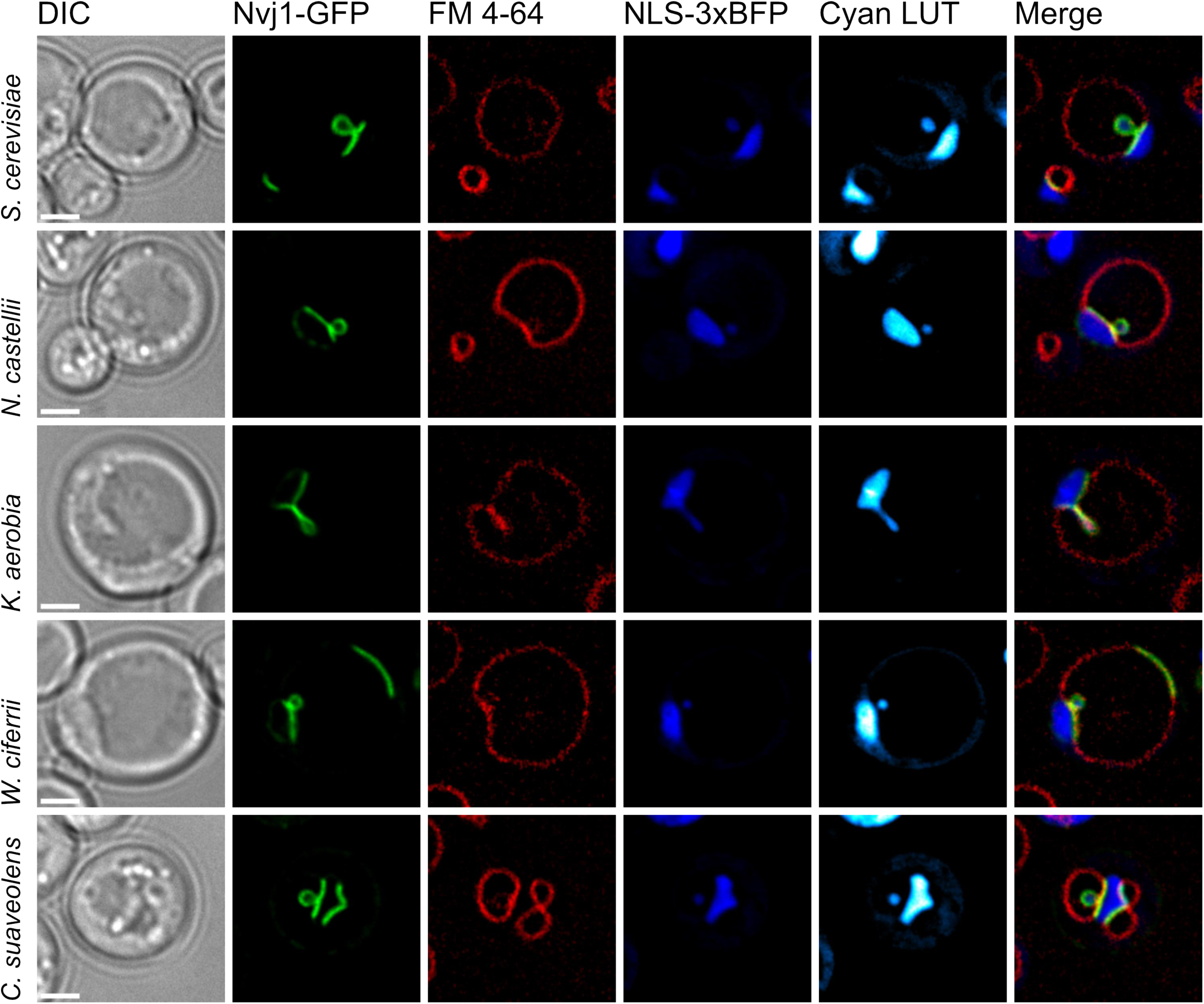
Production of PMN-like structures by Nvj1 candidate orthologs. GFP-tagged Nvj1 candidates were expressed in *S. cerevisiae* strain WCG from the *Met17* promoter induced with 0.3 mM methionine in CM media, grown overnight to stationary phase and suspended in SD-N medium for 2.5 hrs. Vacuole membranes are dyed with FM 4-64 and nucleoplasm with NLS-3xBFP. An enhanced Cyan LUT representation of the signal was used to highlight smaller NLS-3xBFP-stained PMN-like blebs and vesicles. (Scale bar: 2μm)

### Conservation of partner binding

In addition to mediating the formation of NVJs and PMN structures, *S. cerevisiae* Nvj1 binds directly to Vac8, Tsc13, Hmg1 and Osh1. Thus, we tested candidate orthologs for their ability to recruit these factors into NVJs in *S. cerevisiae*. As shown in Fig. 7, examples from groups 1, 2, 3, and 10 recruited Hmg1-mCherry into NVJs when expressed in *S. cerevisiae*. When expressed in *S. cerevisiae* cells, only candidates from group 1, 2 and 3 recruited mCherry-Tsc13 into NVJs (Fig. 8). Candidate orthologs from groups 10 (*W. ciferrii* and *Cy. suaveolens*) and 11 (*Ko. phaffii*) Nvj1-GFPs did not recruit Tsc13-mCherry into NVJs.

**Figure 7.**
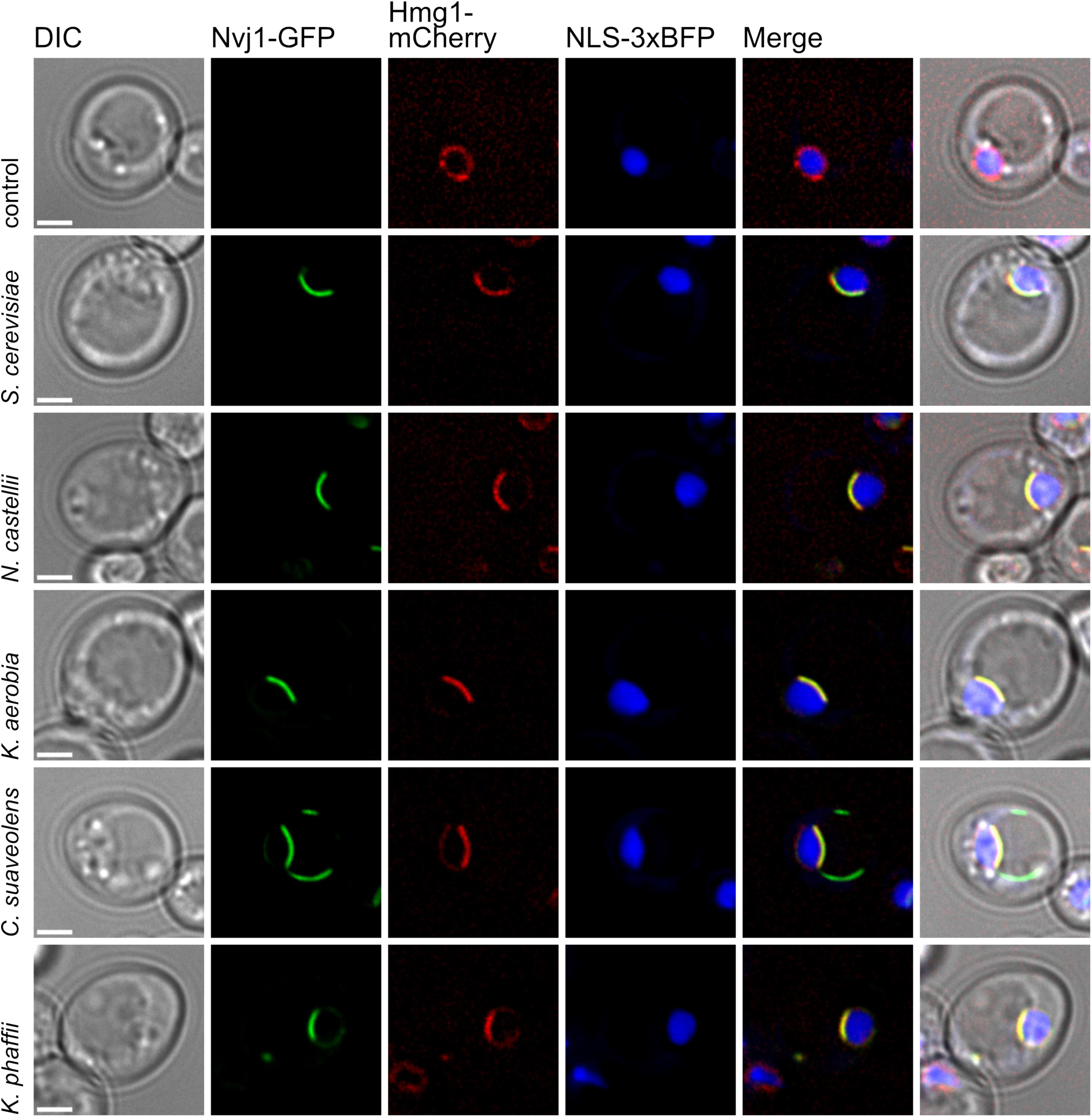
Candidate Nvj1 orthologs recruit Hmg1 to NVJs in *S. cerevisiae*. Strain WCG *nvj1Δ HMG1-mCherry* was transformed with plasmids producing Nab2-NLS-3xBFP for nucleus visualization and one Nvj1-GFP homolog respectively, indicated on the left side. The strains were grown in selective media with 0.3 mM methionine and subsequently starved for 2.5 hrs in SD-N media. A *nvj1Δ S. cerevisiae* strain served as negative control. When recruitment is depicted, it occurred in at least one third of captured cells. (Scale bar: 2μm)

**Figure 8.**
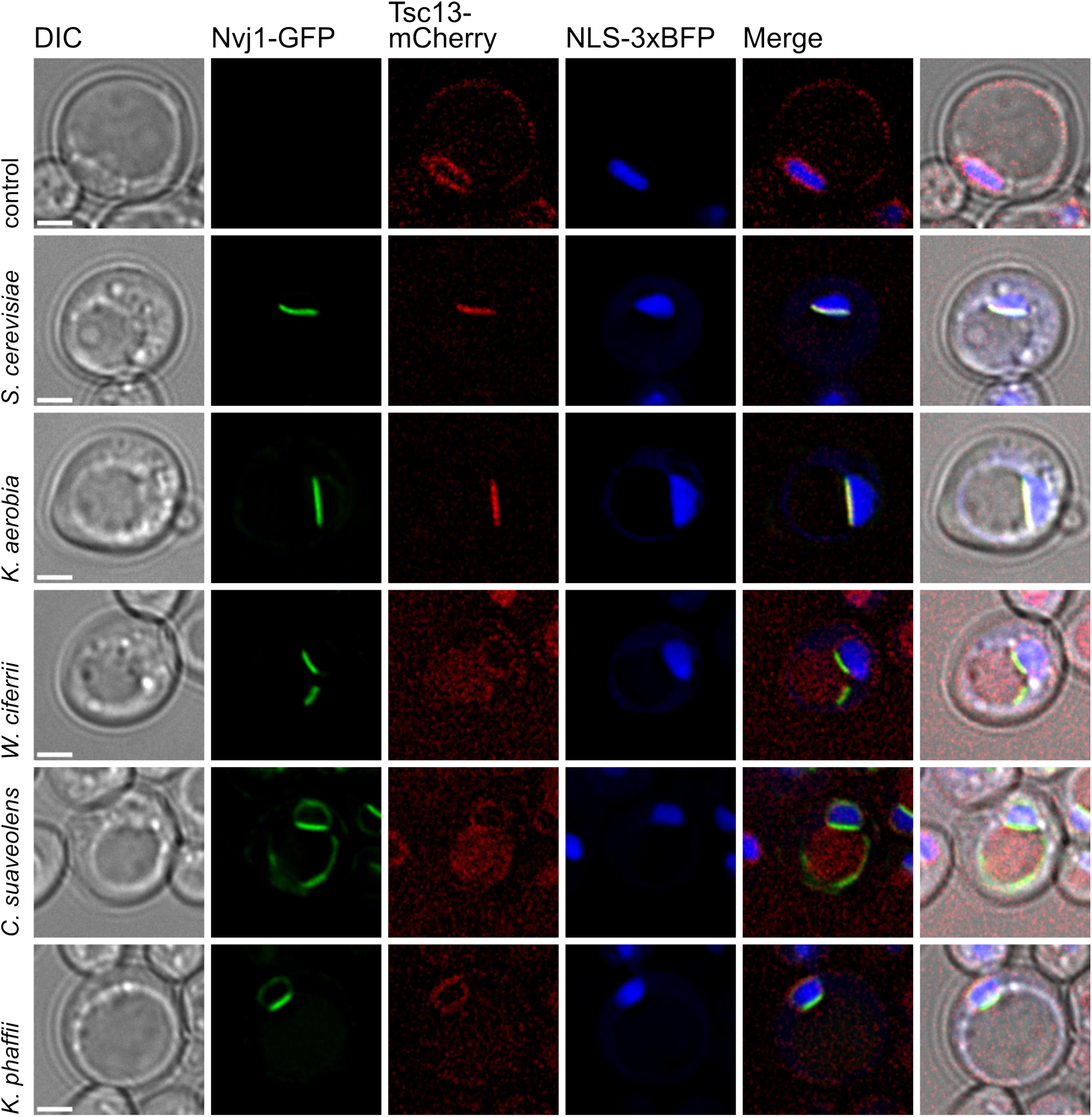
Select candidate Nvj1s recruit Tsc13 to NVJs *S. cerevisiae*. Strain WCG *nvj1Δ TSC13-mCherry* was transformed with plasmids expressing different candidate Nvj1s and induced in CM media with 0.3 mM methionine and subsequently starved in SD-N media for 2.5 hours. A *nvj1Δ S. cerevisiae* strain served as negative control. The shown phenotypes are representative of approximately 80% of analyzed cells. (Scale bar: 2μm)

As described above, both the Osh1 binding motif and the OA motif appear to have been lost three times during the diversification of the *Saccharomycetaceae* lineage. The question remains as to whether the lack of canonical Osh1-binding and OA motifs correlates with a lack of Osh1 binding. Fig. 9 shows that Osh1-mCherry is not recruited into NVJs in cells expressing candidates that lack consensus Osh1-binding and OA motifs.

**Figure 9.**
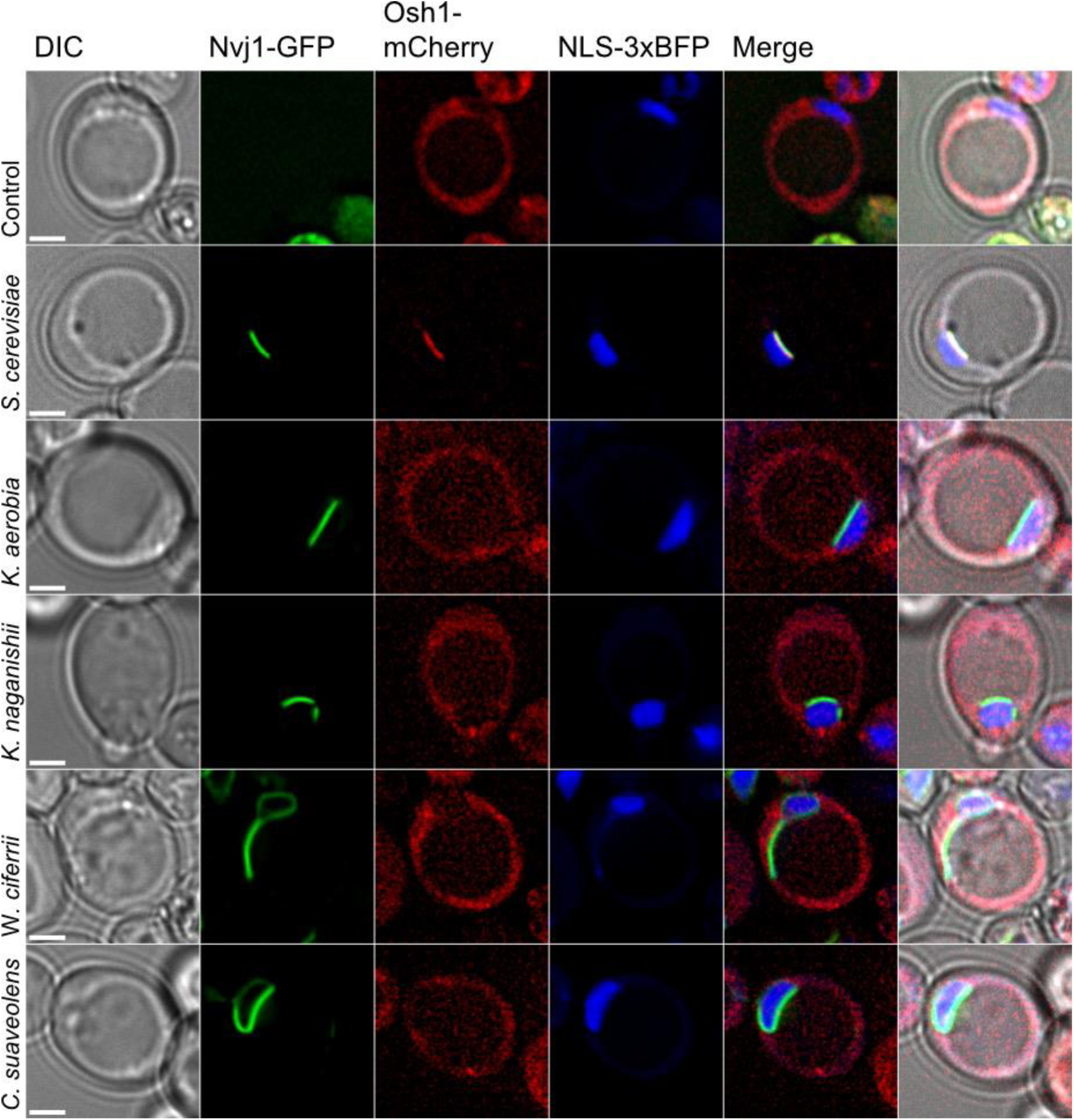
Nvj1 candidates lacking Osh1-binding and OA-motifs do not recruit Osh1 to NVJs. *S. cerevisiae* WCG *nvj1Δ* was transformed with plasmids producing Nab2-NLS-3xBFP for nucleus visualization, mCherry-Osh1 and one Nvj1-GFP homolog respectively, indicated on the left side. The strains were grown in selective media with 0.3 mM methionine and subsequently starved in SD-N media for 2.5 hours. A strain lacking any *NVJ1* was used as a negative control. The extent of the vacuole can be seen in the DIC merge image in the far-right column. All tested homologs form a concentrated junction containing Nvj1-GFP. Only Nvj1-GFP originating from *S. cerevisiae* was able to localize mCherry-Osh1 to the junction in the majority of captured images. Other homologs resemble the negative control. (Scale bar: 2μm)

If, in the course of yeast diversification, these motifs are readily dispensable, what then is the role of Osh1 at NVJs? To investigate this question, we posited that the inability to recruit Osh1 to NVJs might correlate with the gain or loss of other nonessential genes, for example, those with roles in lipid transport or storage (lipid droplets) or lipid metabolism. We performed a gene presence association analysis to search for gene gains and losses that correlate with the loss of Osh1-binding motifs based on gene orthology assignments (Opulente et al., 2024). Only one ortholog, *CTR3*, encoding a plasma membrane-associated copper transporter (Pena et al., 2000) was nearly perfectly correlated with the lack of Osh1 binding motifs in post-WGD lineages.

Unfortunately, there is no obvious functional relationship between Ctr3 and Osh1 or any NVJ-associated protein. The failure of this approach may be due to associated genes being modified (e.g. mutated or over- or under-expressed) rather than gained or lost, or the limited power of the analysis due to the small number of loss events analyzed.

### The OA motif is not required for factor binding

The discovery of a novel OA motif situated in the perinuclear lumen and found only in orthologs containing Osh1-binding sites suggests that the two sites might collaborate, perhaps to facilitate binding to Osh1 or to other Nvj1 binding partners. To investigate possible functions of the OA motif in this regard we created a set of mutant *S. cerevisiae* Nvj1 OA motifs, containing 2, 4 or 6 alanine substitutions. The M6 mutant replaces the six most highly conserved residues of the OA consensus motif. Nvj1 protein levels in M2, M4, and M6 mutant cells were reduced to 70%, 8%, and 4% respectively (Fig. S7A). Although the expression of M6 mutant Nvj1 was significantly reduced (Fig. S7A), enough was expressed to mediate the formation of NVJs (Fig. S7C). These results argue against a role of the OA motif in the formation of Vac8-dependent NVJs.

We further assessed the capacity of the M6 mutant OA motif Nvj1s to localize to NVJs Osh1, Tsc13, and Hmg1 mCherry-tagged reporters. As shown in Fig. S7C, all three of these reporters were recruited to NVJs formed by the M6 mutant Nvj1. We conclude that the OAM is either insensitive to these mutations, or, more likely, that it is not required for the in vivo association of these factors with Nvj1 in the context of NVJs.

Mutations within the OA motif affected both the frequency of NVJs and Nvj1 levels. These effects could be due to a defect in Snd3 binding. Snd3, which associates with both Nvj1 and Vac8, is required for normal levels of Nvj1 and efficient NVJ formation under glucose starvation and may function during the initial insertion of Nvj1 into the ER membrane (Tosal-Castano et al., 2021). We tested whether the reduced levels of Nvj1 in these mutants might be due to a defect in associated with a loss of appropriate engagement with Snd3, possibly during the targeting of Nvj1 to the ER membrane. Snd3-mCherry localized to NVJs in starved cells expressing the M2 and M4 mutants (Fig. S8A), but under these conditions the GFP-M6-Nvj1 fluorescence in the M6 mutant was too low to score. However, the M6 mutant bound in vitro to a Snd3-6xHA reporter (Fig. S8B), supporting the conclusion that the OA motif is not necessary for the interaction of Snd3 with Nvj1.

## Discussion

Model cell systems such as S. cerevisiae have contributed greatly to our understanding of molecular mechanisms of many cell processes. Extending the study of cell processes to evolutionarily and physiologically diverse systems broadens and deepens our understanding of these adaptations in unexpected and meaningful ways. In this study we describe scores of novel Nvj1 orthologs spanning 200 million years of evolution within the subphylum *Saccharomycotina* (see Opulente et al., 2024). Many of these orthologs display extreme clade-specific variability in amino acid sequence and in vivo function.

The *Saccharomycotina* yeasts are a veritable playground of evolutionary and physiological diversity. The clade includes both generalist and specialist species which are adapted to almost every imaginable biome (Chavez et al., 2024). These yeasts vary widely in rates of evolution and reflect adaptations to landmark events such as the WGD event and instances of massive reductions of genome size and gene losses, which in some cases compromised otherwise well-conserved metabolic and genome maintenance pathways.

We employed both sequence and functional criteria to assess whether putative candidates were true genetic orthologs. In cases where NVJ formation was observed following the expression of candidate *NVJ1* genes in *S. cerevisiae,* we are confident that these proteins are true genetic and functional orthologs. The demonstration that the *Ko. phaffii* ortholog formed NVJs in both *S. cerevisiae* and *Ko. phaffii* cells shows that even the most divergent ortholog (relative to *S. cerevisiae*) that we identified among the *Saccharomycotina* is able to mediate the formation of NVJs when heterologously expressed in *S. cerevisiae*.

Perhaps the most striking comparative feature among a number of orthologs is the lack of canonical N-terminal-adjacent hydrophobic sequences (INM anchors). We showed that this motif is required for NVJ formation in *S. cerevisiae* (Kvam and Goldfarb, 2006a; Kvam and Goldfarb, 2006b; Millen et al., 2008), suggesting most likely that the N-termini of Nvj1s insert directly into the lipid bilayer of the INM. However, some orthologs that lack an N-terminal INM anchor still mediate NVJ formation (Fig. 2 and described below). How these proteins link the two nuclear membranes without a canonical INM anchor warrants a re-examination of our proposed mechanism. It is possible that the current model is incorrect, or that it applies selectively to orthologs that display INM anchors, but not to those that do not. The latter group may link to the INM by other mechanisms.

Based on NVJ-forming capacity, candidates from groups 1, 2, 3, 4, 7, 10 and 11 are confirmed to be bona fide genetic and functional orthologs. Caveats persist for candidate orthologs that lack sequence elements that are required for NVJ formation (e.g. INM anchor sequences) and did not produce NVJs when expressed in *S. cerevisiae*. A test case for complications that arise in the functional analysis of candidate orthologs by expression in *S. cerevisiae* is the *Nak. castellii* Nvj1 (group 2). This protein formed mostly extra-nuclear, ER-vacuole junctions in *S. cerevisiae* (Millen et al., 2008). In contrast, the protein formed NVJs when expressed in native *Nak. castellii* cells. The heterologous localization “defect” was rescued by increasing the hydrophobicity of the relatively weak INM anchor sequence. Perhaps the INM anchor is better adapted to some membrane property or biogenetic process characteristic of *Nak. castellii* cells. In the present study, which involved scoring hundreds of cells, the *Nak. castellii* ortholog induced NVJs and very few ectopic junctions in *S. cerevisiae* cells. Moreover, the *Nak. castellii* reporter induced the formation of PMN-like structures in starved *S. cerevisiae* cells. The contrasting results from the previous and current studies are likely due to different strain and culture conditions, both of which are common confounding factors in these types of assays.

A second case involves the *Kl. lactis* (group 8) candidate which we had previously shown to localize to NVJs when expressed in either *S. cerevisiae* or *Kl. lactis* (Millen et al., 2008). This protein exhibits features that are characteristic of bona fide Nvj1s, including a single predicted TMD, acidic pI, Vac8- and Osh1-binding motifs, and an OA motif. However, it lacks anything resembling an N-terminal INM anchor, which, based on our genetic analysis of the linking function in *S. cerevisiae*, was predicted to be a prerequisite to forming discrete NVJs.

A third, striking case is the highly divergent *Han. uvarum* (group 9) candidate, which showed no proclivity to form NVJs in *S. cerevisiae*. Instead, it localized throughout the perinuclear and peripheral ER. Consistent with the protein lacking a consensus Vac8-binding motif, it does not form extra-nuclear, ER-vacuole junctions when expressed in S. cerevisiae. Vac8 is well-conserved among the *Saccharomycotina*. The *Han. uvarum* candidate harbors a single TMD-like hydrophobic sequence close to its N-terminus (aa 13-27), leaving only 12 aa to be potentially exposed to the perinuclear lumen. Most other Nvj1 candidates have at least 80 aa upstream of their TMDs, suggesting that more than 12 aa, even fully extended, is required to span the perinuclear lumen and anchor in the INM. The chromosomal location of the *Han. uvarum* ortholog, between *MDM31* and *NMD5* orthologs, and sequence characteristics (single TMD and ER localization, acidic pI and predominance of unfolded sequence) support the conclusion that this protein is a genetic ortholog. It remains to be determined whether or not this protein mediates NVJs in *Han. uvarum*.

That NVJs are generated in *Han. uvarum* by some means is suggested by a remarkable electron micrograph from a decades-old study on the bipolar budding of some yeast species, including *Han. uvarum* (Kreger-van Rij and Veenhuis, 1971). This image shows what appears to be a striking PMN bleb emerging into the vacuole lumen from an extensive NVJ. Though striking, this single image is insufficient evidence to support the conclusion that NVJs and PMN occur in *Han. uvarum*.

The *Saccharomycetaceae* (groups 1-8) is itself an evolutionarily diverse family, in part because groups 1-5 exhibit the consequences of the WGD event. Duplicated genomes underwent significant changes, including the deletion of hundreds of paralogs and extensive gene order rearrangements. All known *NVJ1* genes in non-WGD genomes, including those of earlier branching families, are sandwiched between *MDM31* and *NMD5* orthologs. Those encoded in post-WGD genomes are all located between *MDM31* and *UTP5* orthologs, indicating the change in gene order occurred soon after the WGD and before diversification of these lineages.

One notable variation among non- and post-WGD *Saccharomycetaceae* Nvj1s is the presence and absence of Osh1 binding and OA motifs among species. Non-WGD *Saccharomycetaceae* species (groups 6-8) all bear Vac8, Osh1, and OA motifs. In contrast, both Osh1 and OA motifs appear to have been lost three times independently during the diversification of post-WGD lineages. The loss of these motifs has functional consequences. We showed that the lack of Osh1 binding motifs in two *Kazachstania* Nvj1s (group 3) precludes their ability in *S. cerevisiae* to recruit Osh1 into NVJs. We found no examples of Osh1 binding motifs or binding in Nvj1 orthologs from lineages that are ancestral to the *Saccharomycetaceae*.

*Lachancea* (group 7), *Kluyveromyces* and *Eremothecium* genera (group 8), the latter of which comprise a separate clade (see Fig. 5), show a wider distribution of isoelectric points than any other group. It is unclear whether shifts in pI are the result of genetic drift or positive selection. As discussed above in the case of the *Kl. lactis* ortholog, group 8 orthologs deserve further attention because they lack both INM anchors and TMD sequences that are presumed to be prerequisite for NVJ formation. Some group 10 candidates also mediate the formation of NVJs without displaying canonical INM anchors (see below).

The situation with domain structures, isoelectric points, and sequence motifs is complicated, with some features showing significant clade-specific variations. *Eremothecium* candidate sequences (group 8) are especially odd. These are recognizable as candidate Nvj1 orthologs because they map between *MDM31* and *NMD5*, are predicted to contain single TMDs and, most telling, some have canonical Osh1/OA and Vac8 motifs. Their putative TMDs occur very near their N-termini. These also lack obvious INM anchors and, in the case of *E. sinecaudum*, for example, have predicted OA and Osh1-binding motifs that are not separated by a TMD. Our assignment of INM and TMD domains for these group 8 proteins and those of group 9, which are also atypical (below), are best guesses (Table S2).

Meme searches identified a motif that appears in numerous candidates from most groups. This motif is composed largely of clusters of acidic residues, i.e. aspartates and glutamates, though without apparent phylogenetic rhyme or reason (e.g. EEEDPEEEDPEE in *Kazachstania saulgeensis* (group 3), EGEEEEEDDDDD in *Cyberlindnera jadinii* (group 10), and EEEEEEEEDDEE in *Kuraishia floccula* (group 11)). Runs of acidic residues in intrinsically disordered domains can mediate selective electrostatic binding, usually to membrane lipids or proteins (Kleiger et al. 2009; Bigman et al. 2022). However, the lack of a consistent phylogenetic pattern suggests that these acidic motifs more likely serve to prevent the formation of stable, compact structures (see below). E (glutamate) is next only to P (proline) in its propensity to promote structural disorder (Uversky, 2013).

The extreme divergence of *Hanseniaspora* candidate orthologs, discussed above, is consistent with the overall rapid genome evolution exhibited by this genus. The *Hanseniaspora* are divided into two clades, a fast-evolving lineage (FEL) and a slow-evolving lineage (SEL), both of which have evolved faster than any other of the yeast lineages (Steenwyk et al., 2019).

The five *Hanseniaspora* orthologs in this group contain representatives from both FEL and SEL clades. All share similar sequence and domain characteristics. Thus, their divergence is not limited to FEL species, and their unique characteristics are ancestral to both clades. Compared to the *S. cerevisiae* genome, FEL and SEL genomes lost on average 1,409 and 771 genes, respectively. Genomes from both FEL and SEL groups lost dozens of genes associated with cell cycle, genome integrity, and DNA repair. Both groups also lost a number of key genes required for biosynthesis of the essential biomolecule thiamine, a fact that is consistent with their adaptation to growth in environments where exogenous thiamine is available, such as fruits. Both clades also lack several genes required for methionine salvage, gluconeogenesis, and many that are required for (efficient) growth on various carbohydrates.

Group 10 is comprised of *Cyberlindnera* and *Wickerhamomyces* species of the family *Phaffomycetaceae*. Based on their domain and sequence motif characteristics, it is remarkable that *Cy. suaveolens* and *W. ciferrii* orthologs form NVJs and PMN-like structures in *S. cerevisiae*. Neither protein harbors apparent INM anchors, though other members of the group are predicted to contain both INM and TMD sequences (*W. chambardii* and *C. jadinii*). In addition to forming NVJs, these proteins also mediate the formation of extra-nuclear, ER-vacuole junctions (Fig. S6). The latter proclivity is consistent with weak INM anchors, at least in the context of *S. cerevisiae* cells.

Based on mutagenesis analyses, the molecular mechanism responsible for linking inner and outer nuclear membranes is presumed to involve the insertion of hydrophobic INM anchor sequences into the lipid bilayer of the INM (Millen et al., 2008). However, alternative mechanisms such as binding to an INM protein have not been ruled out. The formation of NVJs in some cells by Nvj1s like those from *Cy. suaveolens* and *W. ciferrii*, which lack anything resembling typical hydrophobic INM anchor sequences, is a reminder that the mechanism of INM anchor function is not understood.

Methylotrophic *Komagataella* and *Kuraishia* species of the family *Pichiaceae* are the most evolutionarily distant species from *S. cerevisiae* that encode discoverable Nvj1 orthologs. The *Ko. phaffii* protein mediates the formation of NVJs in both *S. cerevisiae* and *Ko. phaffii* cells. The discovery of Nvj1 orthologs in this clade was enabled by the conservation of the *MDM31-NVJ1-NMD5* gene order, which is not conserved in the *Serinales* or *Alaninales* or in earlier branching orders (Fig. 1). Because the *Ko. phaffii* ortholog generates *VAC8*-dependent NVJs, the lack of consensus Vac8-binding motifs among the four group 11 orthologs is likely due to divergence from the consensus sequence, which was calculated mostly from group 1-8 and 10.

We still know little about the molecular mechanism of PMN, including initiation, bleb formation, expansion and scission. Our inability to induce observable PMN-like activity by expressing the *Ko. phaffii* ortholog in either *S. cerevisiae* or *Ko. phaffii* may provide opportunities to elucidate the sequences, factors and conditions that are required for PMN. It is curious that PMN-like structures were induced in *S. cerevisiae* by expressing *Cy. suaveolens* and *W. ciferrii* orthologs, even though these proteins are by some criteria less similar to *S. cerevisiae* Nvj1 than the *Ko. phaffii* ortholog.

In summary, we conclude that most of the candidate Nvj1s described in this report are orthologous to *S. cerevisiae* Nvj1, at least based on their synteny and protein characteristics. We showed that orthologs from groups 1, 2, 3, 10 and 11 mediate the formation of NVJs. Based on their strong similarity to functional orthologs, it is reasonable to predict that orthologs from groups, 4, 5, 6 and 7 would also mediate NVJ formation. Group 8 includes diverse orthologs, at least one of which displays all the elements characteristic of NVJ-forming orthologs, including Osh1-binding and OA motifs. The argument for the *Hanseniaspora* candidates (group 9), which do not mediate the formation of NVJs, being genetic orthologs is weaker than for the others.

Why have *NVJ1* genes, NVJs and PMN only been found in these yeasts? There is currently no evidence for NVJ-like structures or PMN-like nucleophagic processes outside *Saccharomycotina* yeasts (however, see Murayama et al., 2026). It could be due to a lack of looking or a lack of the tools to look. Due to the loss of synteny, we have been unable to determine whether or not they occur in a number of *Saccharomycotina* orders. Where these phenomena do occur, we hypothesize that NVJs and PMN may have evolved to service the closed mitoses and characteristic organization of the ER network of these yeasts, the latter of which is less extensive than in the cells of many other eukaryotic lineages. Closed mitoses, where the nuclear envelope neither breaks down nor ruptures, occur in other eukaryotic lineages, dinoflagellates for example, but are best understood in *S. cerevisiae* (Boettcher and Barral, 2013).

NVJs are positioned to mediate NE membrane expansion. Campbell et al. (2006) proposed that chromatin-associated subdomains of the *S. cerevisiae* NE are resistant to membrane expansion. Instead, the expansion of the NE occurs at chromatin-free regions that are adjacent to the nucleolus. NVJs are organized opposite chromatin-free subdomains of the NE that are instead actively associated with nucleolar pre-ribosomes (Tasnin et al., 2021; Boyle and Wilfling, 2023).

A second possible role for NVJs and PMN relates to the presumed need to employ autophagic mechanisms to degrade nuclear constituents in cells with closed mitoses. The production of PMN blebs and vesicles persists throughout mitosis in *S. cerevisiae* (Roberts et al., 2003). Thus, during both interphase and mitosis, PMN provides a means to transfer nuclear constituents to the vacuole without having to disassemble the NE. It deserves mentioning that Atg39, which serves as a linker protein for macronucleophagy in *S. cerevisiae*, also links the inner and outer nuclear membranes, but using a different mechanism than how Nvj1 is likely to accomplish the feat (Mochida et al., 2022). Moreover, like Nvj1, homology searches for Atg39 orthologs among *Saccharomycotina* genomes yielded limited results, also likely due to homology detection failure (results not shown).

In conclusion, we hope that this collection of Nvj1s from ecologically and physiologically diverse yeast species will encourage investigators to delve into the functional diversity of this fascinating protein and the processes it mediates.

## Materials and Methods

### Identification of Nvj1 candidate orthologs

BLASTp (v2.9.0+) was used to search predicted proteins in 332 yeast genome using NVJ1 as a query (Shen et al., 2018). Protein alignments were performed using PASTA (v1.8.7) and removed 2 hits that didn’t align well, 2 others that were merged with adjacent genes, and one identical duplicate. This alignment was used to generate an HMMER model and search for more distantly related homologs using HMMER (v3.3).

### Gene order searches

The identification of candidate *NVJ1* orthologs by gene order, we initially searched for orthologs of adjacent genes: *MDM31* (upstream) and *UTP9* (downstream). The search was supplemented for orthologs of *NMD5*, the adjacent downstream gene found in non-WGD clades (WGD browser, Ken Wolfe).

Syntenic regions were defined as those with *MDM31* hits and either *UTP9* or *NMD5* hits that were within 5 kb of one another. The absence of hits could be the result of assembly gaps, rearrangements, or insertions increasing the distance between the *NVJ1* adjacent genes. We also required that the orthologs were both hits on the plus and minus strand, as they are in *S. cerevisiae* and prior to the WGD.

### Meme analysis

Orthologs were searched for protein motifs using MEME (v5.5.1). Starting with the 65 HMMER orthologs, we manually trimmed adjacent genes and eliminated homologs that were missing sequences at the beginning or end of the multi-species alignment. Two *Eremothecium* species (*gossypii, sinecaudum*) were added. The resulting 56 homologs were run through MEME using the ZOOPS model with 8 motifs with widths of 5-50.

### Yeast strains and plasmids

Yeast strains used in this study are derived from the *S. cerevisiae* WT strain WCG4 *MATα his2-11,15 leu2-3, 112 ura3* (Thumm et al., 1994) and the *Ko. phaffii* WT strain PPY12 *arg4Δ, his4Δ* (Chang et al., 1995). Deletions and chromosomal integrations in *S. cerevisiae* were generated using the methods described by Janke et al. (2004) and verified via PCR. Auxotrophic selection was ensured by growing strains in a complete minimal medium (CM) containing 0.67% (w/v) yeast nitrogen base without amino acids (Becton Dickinson, 291920), 2% (w/v) glucose (Roth, 6780.2) set to pH 5.6 and supplemented with the appropriate amino acids. In order to induce autophagy, cells were harvested, washed and incubated in SD-N medium containing 0.17% (w/v) yeast nitrogen base without amino acids and ammonium sulphate (Becton Dickinson, 233520) and 2% (w/v) glucose.

*Komagataella phaffii (Pichia pastoris)* cells were transformed by electroporation as described in Cregg et al. (1985). The plasmid expressing Nvj1 was linearized at the *HIS4* gene and integrated into the *HIS4* locus of strain PPY12h and WDY53 by homologous recombination.

The following plasmids were synthesized by GenScript Biotech (Piscataway, USA): pRS316 *S. cerevisiae*-Nvj1-EGFP, pRS316 *Nak. castellii*-Nvj1-EGFP, pRS316 *K. aerobia*-Nvj1-EGFP, pRS316 *Ko. phaffii*-Nvj1-EGFP, pRS316 *W. ciferrii*-Nvj1-EGFP, pRS316 *Cy. suaveolens*-Nvj1-EGFP, pRS316 *Han. uvarum*-Nvj1-EGFP, pRS316 *S. cerevisiae*-Nvj1*^V79A^ ^F82A^-*EGFP (M2), pRS316 *S. cerevisiae*-Nvj1*^V79A^ ^L74A^ ^W76A^ ^F82A^-*EGFP (M4), pRS316 *S. cerevisiae*-Nvj1*^V79A^ ^L74A^ ^W76A^ ^F80A^ ^F82A^ ^I83A^-*EGFP (M6). The previously described plasmid pRS423 (Christianson et al., 1992) and Nab2NLS-2mCherry (Krick et al., 2008) were used in restriction cloning with BamHI and SalI to construct the plasmid pYX242-Nab2NLS-3xBFP. The plasmid pRS423-Nab2NLS-3xBFP was generated by replacing the backbone of pYX242-Nab2NLS-3xBFP with pRS423 using PvuII (Christianson et al., 1992). For the plasmid pUG36-cherry-Osh1 the gene Osh1 was amplified from chromosomal DNA and cloned into the vector pUG36-Cherry (Juris et al., 2015) using SpeI and XhoI. The plasmid pUG23 (Niedenthal et al., 1996) and pUG36-cherry-Osh1 were used to construct pUG23-mCherry-Osh1 through restriction cloning with MfeI and EagI. Yeast strains and plasmids employed in this study are listed in Table 1.

**Table 1.**
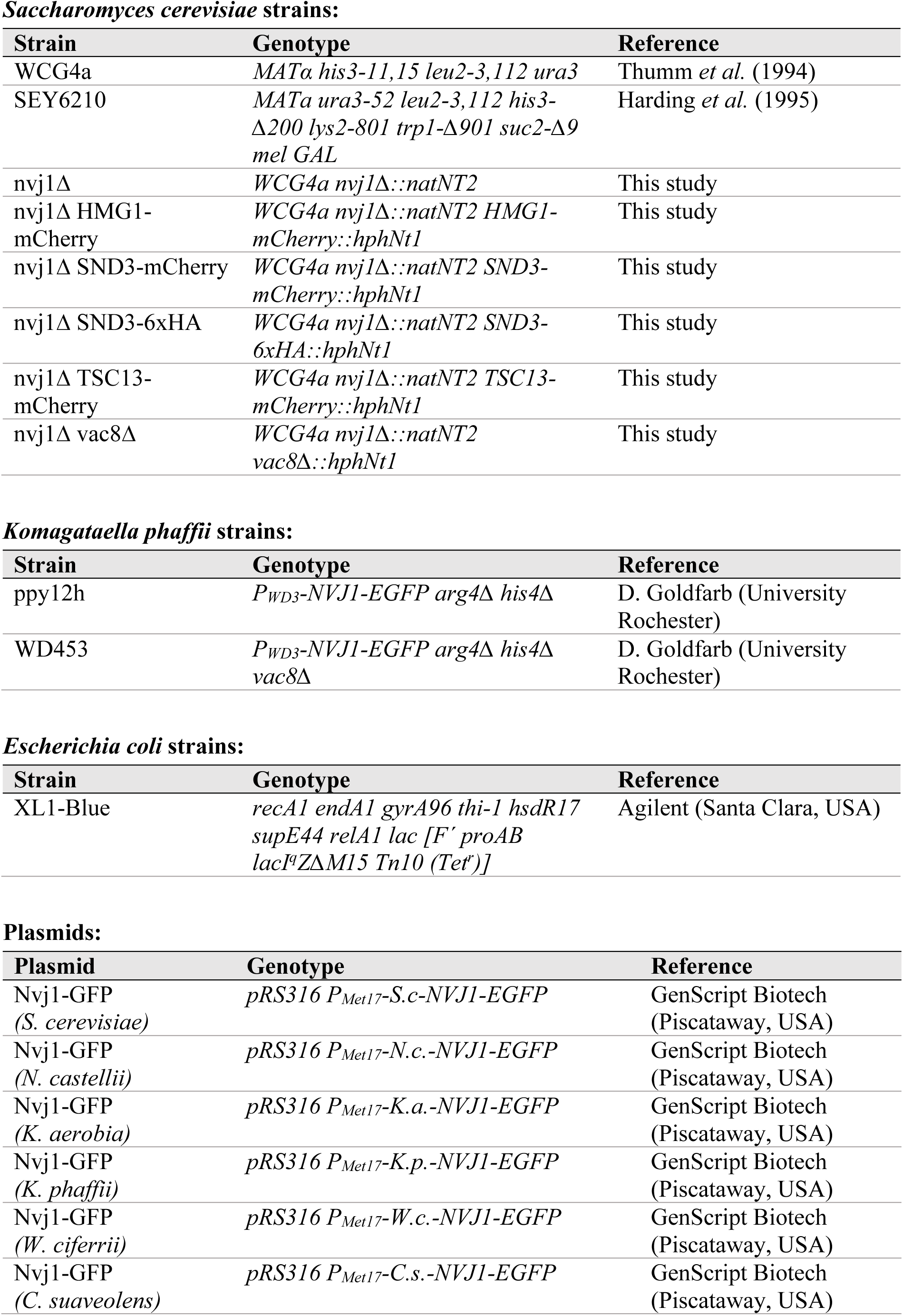
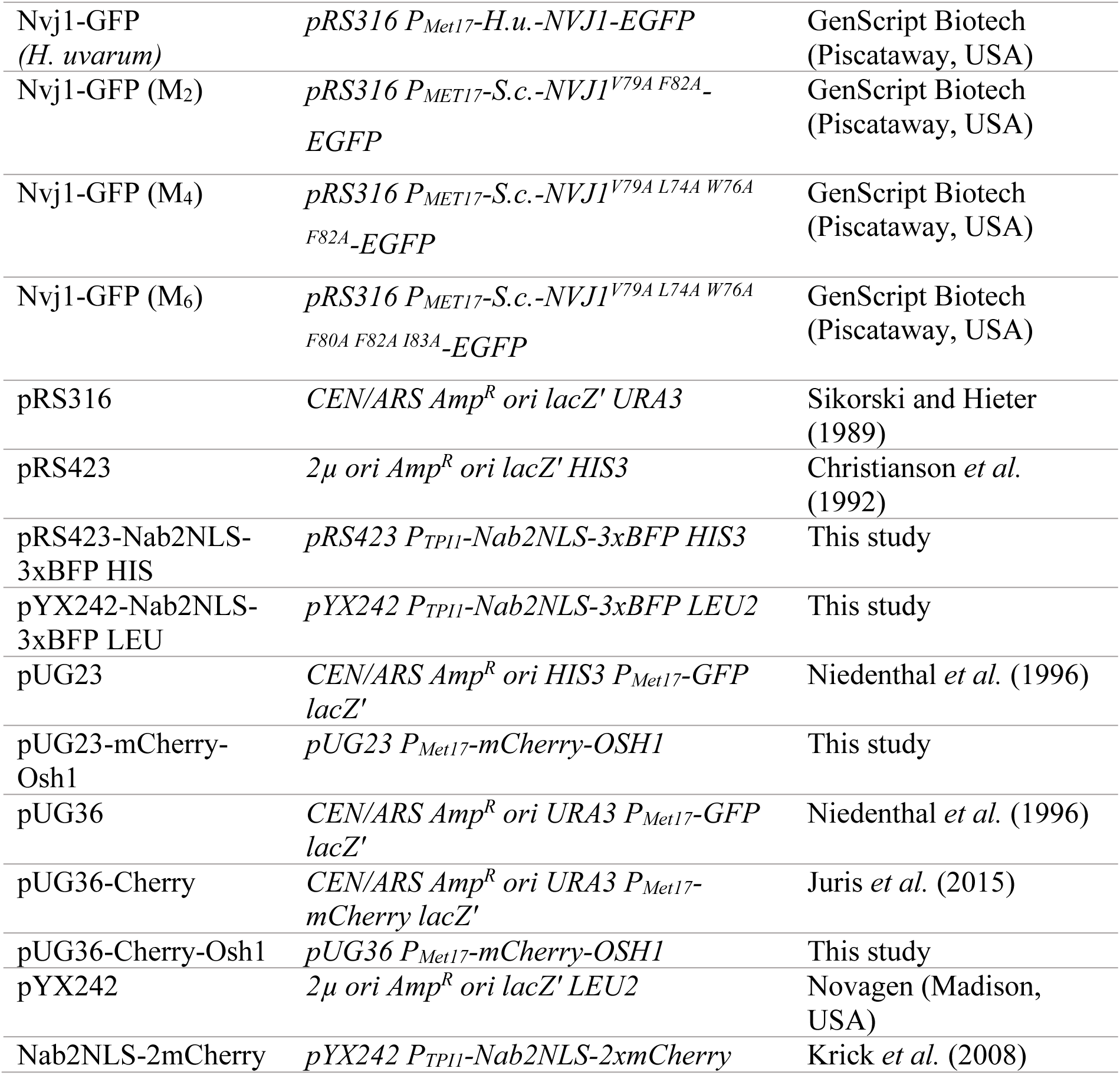
Strains and plasmids used in this study. (Supplied as a separate file, Table_1.pdf.)

### Antibodies

The primary antibodies used in this study were anti-GFP (from mouse IgG1κ; Roche, 11814460001) and the HA-probe antibody (F-7, mouse monoclonal IgG2a; Santa Cruz Biotechnology, sc-7392). Horseradish peroxidase-conjugated goat anti-mouse IgG (Dianova, 115-035-166) served as the secondary antibody.

### Alkaline Lysis

Cells equivalent to 2 OD_600_ were harvested from a logarithmic yeast culture in CM medium containing 0.3 mM methionine and resuspended in 1 ml alkaline lysis buffer (0.28 M sodium hydroxide, 1.125% (v/v) β-mercaptoethanol). After a 10 min incubation on ice, 150 μl of a 50% (v/v) trichloroacetic acid solution was added and incubated for another 10 min.

Afterwards, proteins were pelleted (16,000 g, 10 min, 4°C), washed twice with -20°C acetone and dried at 37°C. The pellet was resuspended in Laemmli buffer (117 mM Tris, pH 8, 3.4% (w/v) SDS, 12% (w/v) glycerol, 0.004% (w/v) bromophenol blue, 0.016% (w/v) β-mercaptoethanol) by shaking for 30 min at 30°C. The sample was analyzed by western blot or stored at -20°C.

### Co-Immunoprecipitation

Co-IPs were performed by binding GFP-tagged proteins to µMACS (magnetic activated cell sorting) columns according to the manufacturer’s protocol (µMACS GFP Isolation Kit, Miltenyi Biotec, 130-091-125). For this 200 OD_600_ of cells, grown overnight to an OD_600_ of 2-3, were harvested, washed and resuspended in lysis buffer (50 mM Tris-HCl pH 7.5, 1mM EDTA (Roth, 8043.1), 0.5% (w/v) Tween-20 (Sigma-Aldrich, P7949), 1 mM PMSF (Roth, 6367.1), 1x complete (EDTA-free Protease Inhibitor Cocktail, Roche, 05056489001), 1x of a 1000x Protease Inhibitor Mix containing 0.1% (w/v) of each Aprotinin, Chymostatin, Leupeptin, Pepstatin).

Lysis was performed with glass beads at and strong shaking at 4°C. Cell debris were removed by centrifugation (10,000 g, 4°C, 10 min) and the supernatant was incubated with 50 µl of µMACs anti-GFP microbeads (Miltenyi Biotec, 130-091-125). After 30 min incubation on ice, the samples were applied to µMAC columns (Miltenyi Biotec, 130-042-701) and washed in accordance with the manufacturers protocol. Proteins were treated with 20 µl 98°C Laemmli buffer (117 mM Tris, pH 8, 3.4% (w/v) SDS, 12% (w/v) glycerol, 0.004% (w/v) bromophenol blue, 0.016% (w/v) β-mercaptoethanol) for 5 min and were afterward eluted from the columns with 50 µl Laemmli buffer. The samples were later analyzed by Western blotting or stored at -20°C.

### Fluorescence microscopy

Fluorescence microscopy was performed using the DeltaVision^®^ microscope (Olympus IX71, Applied Precision) equipped with the UPlanSApo x100, 1.4 numerical aperture, oil immersion objective and a CoolSNAP_HQ2_^®^ couple-charged device (CCD) camera. Imaging occurred with a 100x objective and a 1x1 binning. At least 10 focal planes along the z-axis with a distance of <0.2 µm were captured. The resulting images were deconvolved using softWoRX^®^ (Applied Precision). Vacuole staining was achieved using 10 µg/ml FM 4-64 from Invitrogen (T3166). Nuclei were stained with 15 µg/ml Hoechst-33342 from Sigma (B2261). Both were applied to the growth media, incubated at 30°C for 30 minutes and removed before starvation and subsequent microscopy.

### Statistics

All graphs and statistics were done using GraphPad Prism version 7. All error bars are standard error of the mean (SEM). Statistical confidence was determined either with the one sample t-test or the two-tailed t-test, according to the experimental setup. Asterisks indicate P-values: ns, not significant; P>0.05; * for P<0.05; ** for P<0.01; *** for P<0.001; **** for P<0.0001.

## Acknowledgments

We thank Harmit Malik for suggesting MEME searches and Howard Ochman and Michael Henne for constructive conversations.

## Data availability

All the research data produced by this study are available in the figures and supplementary tables.

## Abbreviations

aa: amino acid
BFP: blue fluorescent protein
C-terminus: carboxy-terminus
ER: endoplasmic reticulum
GFP: green fluorescent protein
EGFP: enhanced green fluorescent protein
INM: inner nuclear membrane
IP: immunoprecipitation
NVJ: nucleus-vacuole junction
N-terminus: amino terminus
NE: nuclear envelope
OA: Osh1-associated
ONM: outer nuclear membrane
PMN: piecemeal micronucleophagy of the nucleus
TMD: transmembrane domain.

## Supplementary material

The following supplementary material is provided as a separate file (Supplementary_Material.pdf), except Table S2, which is supplied as a spreadsheet (Table_S2.xlsx).

**Figure S1.** Identity Matrix plots. (**A**) Identity matrix plot comparing examples of Nvj1 candidates from each of the 11 groups. (**B**). Intraclade identity matrix plots.

**Figure S2.** Heat map distribution of basic (blue) and acidic (red) amino acids in the N- and C-terminal domains of Nvj1 candidate proteins. Candidates are listed by group number (1-11) as defined in Figure 1. Net charges of 20 aa buckets along Nvj1 candidate sequences are interrupted by predicted TMDs, which are split down the middle. N-terminal domains are justified left and C-terminal domains are justified right.

**Figure S3.** Distributions of candidate ortholog (**A**) molecular masses and (**B**) isoelectric points by genus. Data generated by EMBOSS Pepstats (https://www.ebi.ac.uk/jdispatcher/seqstats/emboss_pepstats).

**Figure S4.** Prediction of intrinsically unstructured regions of Nvj1 candidates from each of 11 groups. Data generated by AIUPred (https://iupred.elte.hu). X-axes indicate amino acid number and Y-axes indicate “disorder tendency of each residue in the given protein, where higher values correspond to a higher probability of disorder.”

**Figure S5.** AlphaFold structure predictions of sample candidate Nvj1s from each of the 11 groups.

**Figure S6.** Ectopic localization of *W. ciferrii* and *Cy. suaveolens* Nvj1 candidates in *S. cerevisiae*. WCG *nvj1Δ* cells expressing *NVJ1* homologs from *W. ciferrii* and *Cy. suaveolens* show a unique mislocalization of Nvj1-GFP. These strains form clear elongated accumulations of Nvj1 that either stretch out from the NVJ or are located separately on the vacuolar membrane. Mislocalizations in other homologs are limited to the lack of NVJ formation and diffused peripheral GFP signal, reflecting their *vac8Δ* phenotype. These images were obtained from cultures in selective media containing 0.3 mM methionine. Vacuoles were stained with the dye FM 4-64 for 30 minutes prior to microscopy. The nucleus was visualized with Nab2-NLS-3xBFP, expressed from a plasmid. (Scale bar: 2μm)

**Figure S7.** Recruitment of Hmg1, Osh1 and Tsc13 to M6 mutant OA motif. (**A**) Immunoblot of M2, M4, and M6 OA motif mutants under conditions described below. (**B**) Quantification of OA motif mutant expression. (**C)** Immunofluorescent localization of Hmg1, Osh1 and Tsc13 mCherry reporters in *S. cerevisiae* cells expressing M6 OA mutant Nvj1. *S. cerevisiae* WCG *nvj1Δ,* WCG *nvj1Δ HMG1-mCherry,* WCG *nvj1Δ TSC13-mCherry* were transformed with plasmids producing Nab2-NLS-3xBFP for nucleus visualization and a plasmid producing the mutated protein Nvj1^L74A^ ^W76A^ ^V79A^ ^F80A^ ^F82A^ ^I83A^-GFP. Cells were grown in CM with 0.3 mM methionine and starved in SD-N media (for 2.5 hours). (Scale bar: 2μm)

**Figure S8.** OA motif mutant Nvj1s recruit Snd3 to NVJs. (**A**) Snd3*-mCherry* was expressed in *S. cerevisiae* cells by induction of the *MET17* promoter with 0.3 mM methionine. WCG *nvj1Δ SND3-mCherry* cells transformed with plasmids containing the wildtype *NVJ1- GFP* (WT), *NVJ1V79A F82A-EGFP* (M2) and *NVJ1V79A L74A W76A F82A-EGFP* (M4) OA mutants, respectively. Nuclei were stained with Nab2-NLS-3xBFP, and vacuoles with FM4-64. (Scale bar: 2μm). Microscopy was performed with cells in log-phase (log) as well as overnight cultures. The first two rows show cells expressing wildtype *NVJ1* form NVJs in both log and stationary phase. Snd3-mCherry shows a diffuse distribution on the nuclear membrane and cell periphery in log phase and then co-localizes with Nvj1-GFP in late stationary cells. This co-localization is also seen in the two tested mutant strains M2 and M4. (**B**) Snd3-6XHA copurifies with M2, M4, and M6 OA mutant Nvj1-GFPs (M6 mutant: *V79A L74A W76A F80A F82A I83A-EGFP*). Control cells expressed free *GFP*. The top row shows that all mutants were able to bind Snd3-6xHA, whereas the GFP control showed only a weak signal. The third row indicates bound free GFP. (**C**) Quantification experiment shown in panel B. scale bar: 2μm)

**Table S1.** Protein sequences of Nvj1 orthologs

**Table S2.** Properties and characterizations of *NVJ1* genes and Nvj1 proteins

