## Supplemental Figure S1 for "Functional diversification across yeast lineages of the nucleus-vacuole junction forming protein Nvj1"

### Kramer et al.

## A

| Group 6 | Z. gambellarensis | Z. pseudorouxii | Z. mellis | Z. siamensis | Z. kombuchaensis | Z. lentus | Z. bisporus | Z. bailii | Z. florentina | Z. chibaensis | Z. mrakii | Z. danielsina | H. aurorensis |
|---|---|---|---|---|---|---|---|---|---|---|---|---|---|
| Z. gambellarensis | - | | | | | | | | | | | | |
| Z. pseudorouxii | 72.1 | - | | | | | | | | | | | |
| Z. mellis | 65.7 | 74.5 | - | | | | | | | | | | |
| Z. siamensis | 64.3 | 75.2 | 93.9 | - | | | | | | | | | |
| Z. kombuchaensis | 57.4 | 59.3 | 55.8 | 57.7 | - | | | | | | | | |
| Z. lentus | 56.8 | 57.7 | 54.1 | 56.0 | 83.6 | - | | | | | | | |
| Z. bisporus | 59.7 | 57.8 | 53.9 | 53.9 | 70.6 | 68.6 | - | | | | | | |
| Z. bailii | 57.9 | 54.6 | 54.8 | 53.7 | 71.3 | 66.8 | 72.8 | - | | | | | |
| Z. florentina | 31.6 | 32.1 | 30.5 | 30.8 | 30.4 | 30.5 | 29.3 | 29.9 | - | | | | |
| Z. chibaensis | 31.2 | 31.1 | 28.3 | 28.3 | 28.0 | 28.9 | 30.4 | 30.7 | 73.4 | - | | | |
| Z. mrakii | 29.0 | 27.8 | 27.5 | 26.5 | 28.8 | 30.1 | 29.3 | 30.4 | 36.9 | 36.6 | - | | |
| Z. danielsina | 29.9 | 29.1 | 29.9 | 28.8 | 31.9 | 31.6 | 33.6 | 32.1 | 40.3 | 40.9 | 47.4 | - | |
| H. aurorensis | 24.7 | 22.9 | 24.7 | 24.7 | 27.7 | 27.1 | 25.8 | 27.4 | 20.7 | 22.0 | 21.7 | 21.6 | - |

| Group 7 | L. mirantina | L. kluyveri | L. meyersii | L. fantastica | L. lanzarotensis | L. nothofagi | L. dasiensis | L. fermentati | L. waltii | L. quebecensis | L. thermotolerans |
|---|---|---|---|---|---|---|---|---|---|---|---|
| L. mirantina | - | | | | | | | | | | |
| L. kluyveri | 27.4 | - | | | | | | | | | |
| L. meyersii | 25.5 | 33.1 | - | | | | | | | | |
| L. fantastica | 29.4 | 31.4 | 47.0 | - | | | | | | | |
| L. lanzarotensis | 25.8 | 29.1 | 47.5 | 66.8 | - | | | | | | |
| L. nothofagi | 28.2 | 33.0 | 41.4 | 37.6 | 38.2 | - | | | | | |
| L. dasiensis | 25.4 | 28.2 | 31.8 | 32.0 | 32.4 | 38.7 | - | | | | |
| L. fermentative | 32.9 | 32.9 | 29.7 | 27.2 | 27.3 | 29.2 | 26.9 | - | | | |
| L. wellie | 27.5 | 30.9 | 32.3 | 32.6 | 33.1 | 33.0 | 30.7 | 30.9 | - | | |
| L. quickness | 27.1 | 33.5 | 35.8 | 35.4 | 37.4 | 38.6 | 35.1 | 36.7 | 41.9 | - | |
| L. thermotolerance | 30.5 | 34.5 | 36.8 | 34.9 | 36.9 | 37.7 | 36.8 | 34.8 | 42.9 | 76.5 | - |

| Group 9 | H. vinae | H. valbyensis | H. pseudoguilliermondi | H. uvarum | H. clermontiae |
|---|---|---|---|---|---|
| H. vinae | - | | | | |
| H. valbyensis | 17.4 | - | | | |
| H. pseudoguilliermondi | 23.7 | 27.5 | - | | |
| H. uvarum | 24.3 | 27.2 | 60.5 | - | |
| H. clermontiae | 21.8 | 27.8 | 59.4 | 67.5 | - |

| Group 8 | E. gossypii | A. aceri | K. nonfermentans | K. aestuarii | K. siamensis | K. wickerhamii | K. lactis | K. dobzhanskii | K. starmeri | E. coryli | E. cymbalariae | E. sinecaudum |
|---|---|---|---|---|---|---|---|---|---|---|---|---|
| E. gossypii | - | | | | | | | | | | | |
| A. aceri | 48.0 | - | | | | | | | | | | |
| K. nonfermentans | 13.7 | 16.0 | - | | | | | | | | | |
| K. aestuarii | 17.0 | 19.3 | 45.6 | - | | | | | | | | |
| K. siamensis | 15.7 | 17.4 | 46.2 | 77.6 | - | | | | | | | |
| K. wickerhamii | 13.1 | 14.1 | 28.5 | 28.6 | 27.3 | - | | | | | | |
| K. lactis | 15.8 | 17.7 | 25.3 | 25.5 | 26.2 | 29.6 | - | | | | | |
| K. dobzhanskii | 16.4 | 17.7 | 29.4 | 30.0 | 27.9 | 31.0 | 53.4 | - | | | | |
| K. starmeri | 15.2 | 18.9 | 22.4 | 28.5 | 27.5 | 27.8 | 34.7 | 35.1 | - | | | |
| E. coryli | 15.7 | 20.9 | 17.4 | 19.4 | 18.2 | 18.6 | 19.6 | 22.5 | 20.0 | - | | |
| E. cymbalariae | 18.1 | 18.7 | 24.4 | 26.3 | 24.8 | 20.9 | 20.9 | 20.2 | 21.2 | 24.6 | - | |
| E. sinecaudum | 13.7 | 15.6 | 20.8 | 21.6 | 22.0 | 19.9 | 19.4 | 22.0 | 22.4 | 25.7 | 28.4 | - |

| Group 10 | W. chambardii | W. piperi | W. mucosus | W. hampshirensis | C. fabianii | C. jadinii | C. suaveolens | C. saturnus | C. misumaiensis | C. maclurae | W. ciferrii | W. alni | W. canadensis |
|---|---|---|---|---|---|---|---|---|---|---|---|---|---|
| W. chambardii | - | | | | | | | | | | | | |
| W. piperi | 15.7 | - | | | | | | | | | | | |
| W. mucosus | 18.2 | 24.4 | - | | | | | | | | | | |
| W. hampshirensis | 18.4 | 20.5 | 26.0 | - | | | | | | | | | |
| C. fabianii | 21.1 | 24.3 | 26.5 | 36.1 | - | | | | | | | | |
| C. jadinii | 19.8 | 18.5 | 19.2 | 32.2 | 31.3 | - | | | | | | | |
| C. suaveolens | 17.9 | 22.8 | 26.8 | 34.9 | 35.1 | 38.2 | - | | | | | | |
| C. saturnus | 18.9 | 22.4 | 26.8 | 35.9 | 32.8 | 38.5 | 86.3 | - | | | | | |
| C. misumaiensis | 18.9 | 18.4 | 21.7 | 36.4 | 38.9 | 33.7 | 40.8 | 38.6 | - | | | | |
| C. maclurae | 20.8 | 23.2 | 28.1 | 42.1 | 43.3 | 38.8 | 43.7 | 42.7 | 73.8 | - | | | |
| W. ciferrii | 17.1 | 17.6 | 21.2 | 26.5 | 26.8 | 24.6 | 31.4 | 30.7 | 23.9 | 30.8 | - | | |
| W. alni | 19.3 | 21.5 | 27.7 | 32.4 | 31.9 | 28.0 | 33.7 | 33.7 | 30.7 | 33.5 | 29.7 | - | |
| W. canadensis | 22.2 | 24.6 | 31.4 | 37.0 | 34.1 | 34.5 | 39.5 | 38.1 | 34.7 | 37.2 | 35.2 | 47.7 | - |

| Group 11 | K. floccosa | K. populi | Komatagataella | K. pastoris | K. phaffi | K. kurtzmanii |
|---|---|---|---|---|---|---|
| K. floccoa | - | | | | | |
| K. populi | 25.0 | - | | | | |
| Komatagataella | 23.9 | 73.4 | - | | | |
| K. pastoris | 23.0 | 72.7 | 72.7 | - | | |
| K. phaffi | 23.0 | 71.8 | 71.8 | 90.2 | - | |
| K. kurtzmanii | 23.0 | 71.8 | 71.8 | 90.2 | 98.7 | - |

## B

| Interclade | K. pastoris | H. pseudoguilliermondi | C. suaveolens | K. wickerhamii | V. tropicalis | L. quebecensis | N. baii | T. maleeae | S. cerevisiae | N. bracarensis | K. yasuniensis |
|---|---|---|---|---|---|---|---|---|---|---|---|
| K. pastoris | - | | | | | | | | | | |
| H. pseudoguilliermondi | 23.2 | - | | | | | | | | | |
| C. suaveolens | 13.1 | 22.6 | - | | | | | | | | |
| K. wickerhamii | 13.7 | 18.7 | 14.0 | - | | | | | | | |
| V. tropicalis | 17.0 | 11.9 | 14.5 | 13.2 | - | | | | | | |
| L. quebecensis | 13.4 | 17.4 | 13.1 | 20.4 | 16.0 | - | | | | | |
| N. baii | 18.5 | 15.1 | 17.8 | 14.2 | 22.6 | 18.4 | - | | | | |
| T. maleeae | 13.3 | 12.6 | 12.1 | 19.2 | 20.3 | 24.1 | 20.5 | - | | | |
| S. cerevisiae | 11.6 | 16.7 | 15.4 | 22.0 | 21.6 | 21.4 | 25.6 | 28.9 | - | | |
| N. bracarensis | 12.2 | 19.1 | 17.0 | 15.5 | 17.1 | 18.2 | 20.8 | 22.8 | 24.6 | - | |
| K. yasuniensis | 15.8 | 18.0 | 17.0 | 16.0 | 21.0 | 18.0 | 21.1 | 17.8 | 22.2 | 32.1 | - |

| Group 1 | S. uvarum | S. eubayanus | S. arboricola | S. kudriavzevii | S. jurei | S. mikatae | S. cerevisiae | S. paradoxus |
|---|---|---|---|---|---|---|---|---|
| S. uvarum | - | | | | | | | |
| S. eubayanus | 89.4 | - | | | | | | |
| S. arboricola | 67.6 | 67.5 | - | | | | | |
| S. kudriavzevii | 68.6 | 68.8 | 71.1 | - | | | | |
| S. jurei | 68.1 | 67.7 | 67.6 | 67.5 | - | | | |
| S. mikatae | 69.7 | 70.2 | 69.7 | 68.8 | 90.8 | - | | |
| S. cerevisiae | 69.9 | 69.8 | 72.5 | 67.7 | 75.7 | 78.5 | - | |
| S. paradoxus | 73.4 | 72.4 | 74.1 | 72.5 | 81.0 | 83.2 | 86.9 | - |

| Group 2 | N. bacillisporus | N. castellii | N. glabrata | N. uthaithanina | N. delphensis | N. bracarensis | N. kungkrabaensis | N. nivariensis |
|---|---|---|---|---|---|---|---|---|
| N. bacillisporus | - | | | | | | | |
| N. castellii | 26.9 | - | | | | | | |
| N. glabrata | 18.2 | 27.5 | - | | | | | |
| N. uthaithanina | 27.6 | 23.9 | 44.7 | - | | | | |
| N. delphensis | 22.2 | 28.9 | 45.6 | 55.9 | - | | | |
| N. bracarensis | 23.5 | 27.1 | 47.9 | 58.4 | 60.7 | - | | |
| N. kungkrabaensis | 23.2 | 28.0 | 46.4 | 55.7 | 64.7 | 69.7 | - | |
| N. nivariensis | 23.2 | 28.0 | 45.9 | 57.0 | 64.7 | 70.6 | 97.4 | - |

| Group 3 | K. viticola | K. bromeliacearum | K. sinensis | K. naganishii | K. taianensis | K. unispora | K. yasuniensis | K. servazzi | K. solicola | K. aerobia | K. aquatica | K. siamensis | K. martiniae | K. africana | K. jinghongensis | K. spencerorum | K. piceae | K. lodderae | K. gamospora | K. hellenica | K. psychrophila | K. kunashirensis | K. exigua | K. bulderi | K. turicensis | K. saulgeensis | K. barnettii |
|---|---|---|---|---|---|---|---|---|---|---|---|---|---|---|---|---|---|---|---|---|---|---|---|---|---|---|---|
| K. viticola | - | | | | | | | | | | | | | | | | | | | | | | | | | | |
| K. bromeliacearum | 23.9 | - | | | | | | | | | | | | | | | | | | | | | | | | | |
| K. sinensis | 23.0 | 31.9 | - | | | | | | | | | | | | | | | | | | | | | | | | |
| K. naganishii | 21.6 | 29.7 | 86.3 | - | | | | | | | | | | | | | | | | | | | | | | | |
| K. taianensis | 21.5 | 25.3 | 25.6 | 25.1 | - | | | | | | | | | | | | | | | | | | | | | | |
| K. unispora | 22.3 | 28.3 | 27.9 | 27.8 | 27.9 | - | | | | | | | | | | | | | | | | | | | | | |
| K. yasuniensis | 17.9 | 30.0 | 28.0 | 28.3 | 28.9 | 58.5 | - | | | | | | | | | | | | | | | | | | | | |
| K. servazzi | 19.5 | 29.0 | 26.5 | 26.9 | 32.2 | 55.0 | 56.8 | - | | | | | | | | | | | | | | | | | | | |
| K. solicola | 17.7 | 26.7 | 25.9 | 27.7 | 30.4 | 52.1 | 54.9 | 68.2 | - | | | | | | | | | | | | | | | | | | |
| K. aerobia | 20.7 | 27.8 | 27.0 | 28.4 | 30.0 | 53.7 | 53.7 | 66.4 | 75.9 | - | | | | | | | | | | | | | | | | | |
| K. aquatica | 23.4 | 32.4 | 28.8 | 28.6 | 28.4 | 51.4 | 53.2 | 51.5 | 49.7 | 48.6 | - | | | | | | | | | | | | | | | | |
| K. siamensis | 23.9 | 25.9 | 28.7 | 27.6 | 30.3 | 44.5 | 45.6 | 45.0 | 43.5 | 42.5 | 56.3 | - | | | | | | | | | | | | | | | |
| K. martiniae | 23.5 | 20.2 | 20.9 | 20.6 | 19.9 | 20.2 | 24.7 | 23.0 | 20.6 | 21.4 | 22.5 | 20.6 | - | | | | | | | | | | | | | | |
| K. africana | 23.6 | 23.3 | 24.2 | 23.5 | 18.7 | 18.5 | 21.7 | 20.5 | 19.0 | 21.8 | 22.7 | 22.9 | 30.2 | - | | | | | | | | | | | | | |
| K. jinghongensis | 22.7 | 19.4 | 16.9 | 18.2 | 17.2 | 17.1 | 17.8 | 18.0 | 17.8 | 18.9 | 18.2 | 15.6 | 22.7 | 22.7 | - | | | | | | | | | | | | |
| K. spencerorum | 20.6 | 22.9 | 18.1 | 19.0 | 16.5 | 16.9 | 16.4 | 17.7 | 18.7 | 19.8 | 19.9 | 17.3 | 25.0 | 22.8 | 49.8 | - | | | | | | | | | | | |
| K. piceae | 20.2 | 22.0 | 23.8 | 25.2 | 19.5 | 19.8 | 19.3 | 23.6 | 22.1 | 21.6 | 24.2 | 23.7 | 26.4 | 24.8 | 22.6 | 23.6 | - | | | | | | | | | | |
| K. lodderae | 26.7 | 20.6 | 22.7 | 22.2 | 19.9 | 18.8 | 15.9 | 18.9 | 18.4 | 18.8 | 19.4 | 19.2 | 26.8 | 26.6 | 29.9 | 27.4 | 26.8 | - | | | | | | | | | |
| K. gamospora | 23.4 | 18.3 | 18.1 | 17.1 | 15.6 | 17.6 | 15.3 | 17.2 | 15.2 | 17.7 | 18.9 | 19.7 | 23.3 | 25.5 | 26.8 | 25.1 | 25.1 | 31.6 | - | | | | | | | | |
| K. hellenica | 22.6 | 19.5 | 22.1 | 21.6 | 16.7 | 19.3 | 17.3 | 18.6 | 16.2 | 18.8 | 20.9 | 19.0 | 22.7 | 25.6 | 25.5 | 23.6 | 26.4 | 30.1 | 52.9 | - | | | | | | | |
| K. psychrophila | 22.2 | 22.2 | 25.1 | 25.6 | 21.1 | 21.7 | 20.5 | 19.3 | 21.5 | 22.3 | 22.5 | 23.2 | 25.8 | 21.5 | 25.1 | 27.3 | 25.9 | 26.1 | 29.8 | 27.4 | - | | | | | | |
| K. kunashirensis | 21.0 | 20.8 | 21.9 | 20.3 | 20.0 | 21.9 | 20.4 | 18.4 | 19.7 | 19.3 | 22.3 | 22.2 | 23.8 | 23.0 | 26.9 | 25.3 | 24.9 | 27.8 | 26.6 | 29.7 | 40.3 | - | | | | | |
| K. exigua | 19.3 | 20.8 | 22.8 | 24.7 | 19.5 | 19.4 | 19.2 | 17.3 | 19.5 | 20.2 | 20.5 | 21.7 | 21.7 | 21.8 | 24.6 | 24.4 | 25.5 | 26.6 | 26.0 | 27.4 | 37.6 | 40.4 | - | | | | |
| K. bulderi | 19.7 | 22.5 | 23.5 | 24.9 | 21.0 | 20.5 | 20.0 | 20.8 | 21.4 | 21.7 | 22.3 | 23.6 | 23.2 | 22.7 | 24.8 | 22.7 | 26.1 | 26.1 | 26.2 | 27.5 | 36.9 | 40.4 | 79.7 | - | | | |
| K. turicensis | 19.1 | 22.0 | 24.3 | 25.3 | 20.0 | 20.1 | 19.7 | 20.5 | 21.0 | 21.7 | 21.6 | 22.9 | 22.9 | 24.0 | 24.6 | 24.1 | 25.6 | 26.8 | 25.1 | 27.4 | 35.2 | 40.1 | 78.5 | 91.2 | - | | |
| K. saulgeensis | 19.5 | 20.3 | 23.0 | 21.4 | 22.5 | 19.4 | 20.8 | 19.8 | 19.1 | 21.2 | 20.5 | 23.1 | 23.3 | 22.0 | 25.2 | 23.7 | 26.1 | 28.4 | 28.0 | 27.8 | 37.9 | 42.1 | 56.3 | 56.3 | 54.7 | - | |
| K. barnettii | 20.1 | 21.2 | 23.8 | 22.8 | 22.0 | 20.0 | 21.5 | 21.6 | 21.9 | 22.5 | 21.7 | 25.5 | 22.5 | 23.6 | 25.5 | 22.7 | 23.5 | 29.1 | 28.1 | 28.8 | 35.5 | 40.7 | 57.2 | 55.9 | 54.0 | 71.7 | - |

| Group 4 | N. dairenensis | N. castellii | N. baii |
|---|---|---|---|
| N. dairenensis | - | | |
| N. castellii | 36.5 | - | |
| N. baii | 33.7 | 46.9 | - |

| Group 5 | T. iriomotensis | T. namnaonensis | T. blattae | V. polyspora | V. tropicalis | V. yarrowii | V. verrucispora |
|---|---|---|---|---|---|---|---|
| T. iriomotensis | - | | | | | | |
| T. namnaonensis | 34.8 | - | | | | | |
| T. blattae | 19.2 | 18.6 | - | | | | |
| V. polyspora | 23.2 | 20.9 | 18.6 | - | | | |
| V. tropicalis | 22.9 | 18.7 | 18.9 | 25.5 | - | | |
| V. yarrowii | 20.2 | 22.4 | 22.1 | 26.0 | 30.6 | - | |
| V. verrucispora | 24.2 | 18.3 | 19.2 | 23.8 | 26.5 | 35.9 | - |

### Figure. S1
