## Supplementary figures and images for "Functional diversification across yeast lineages of the nucleus-vacuole junction forming protein Nvj1"

### Supplemental Figure S2

Kramer et al.

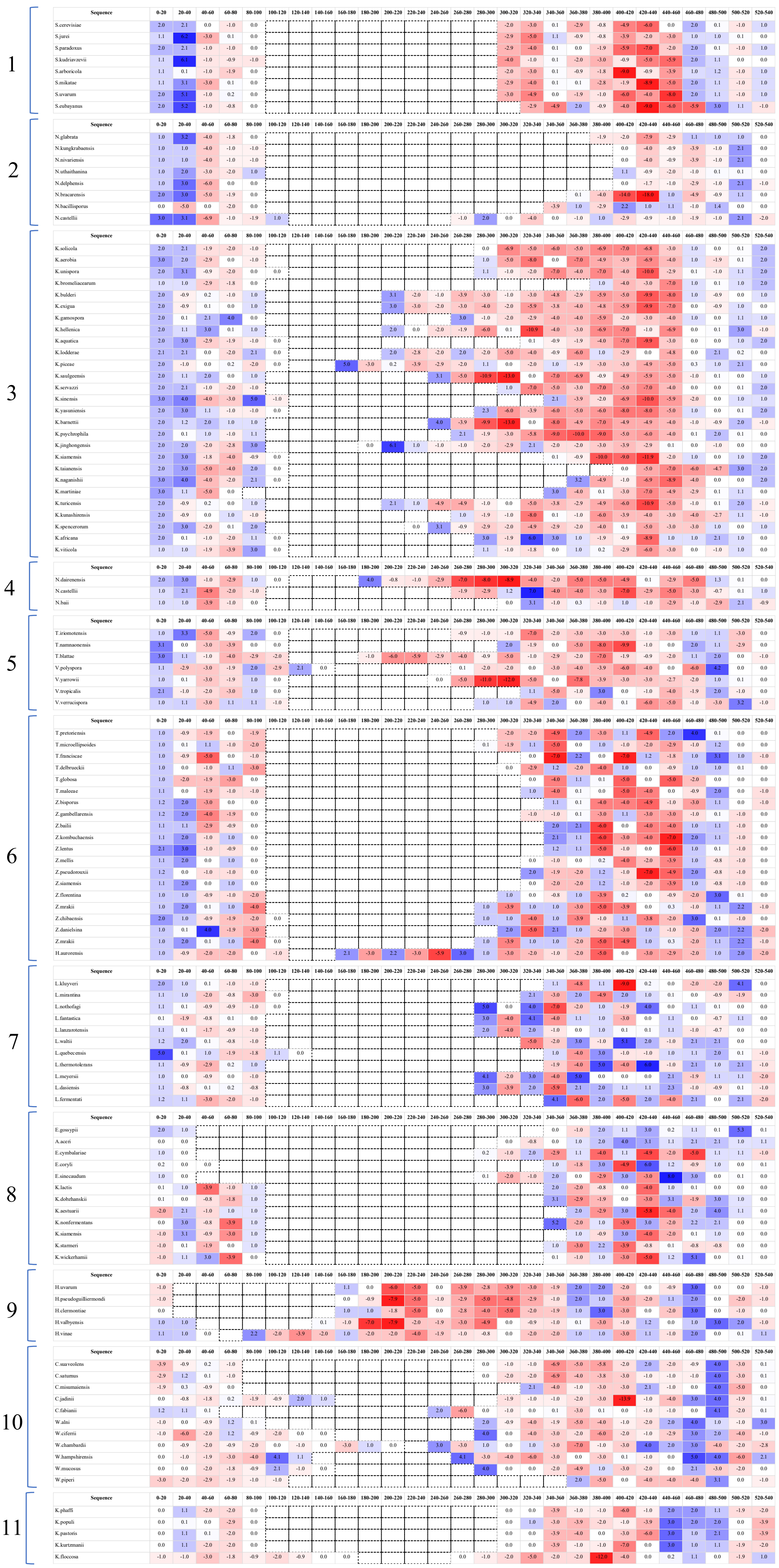

Figure. S2

### Supplemental Figure S3

Kramer et al.

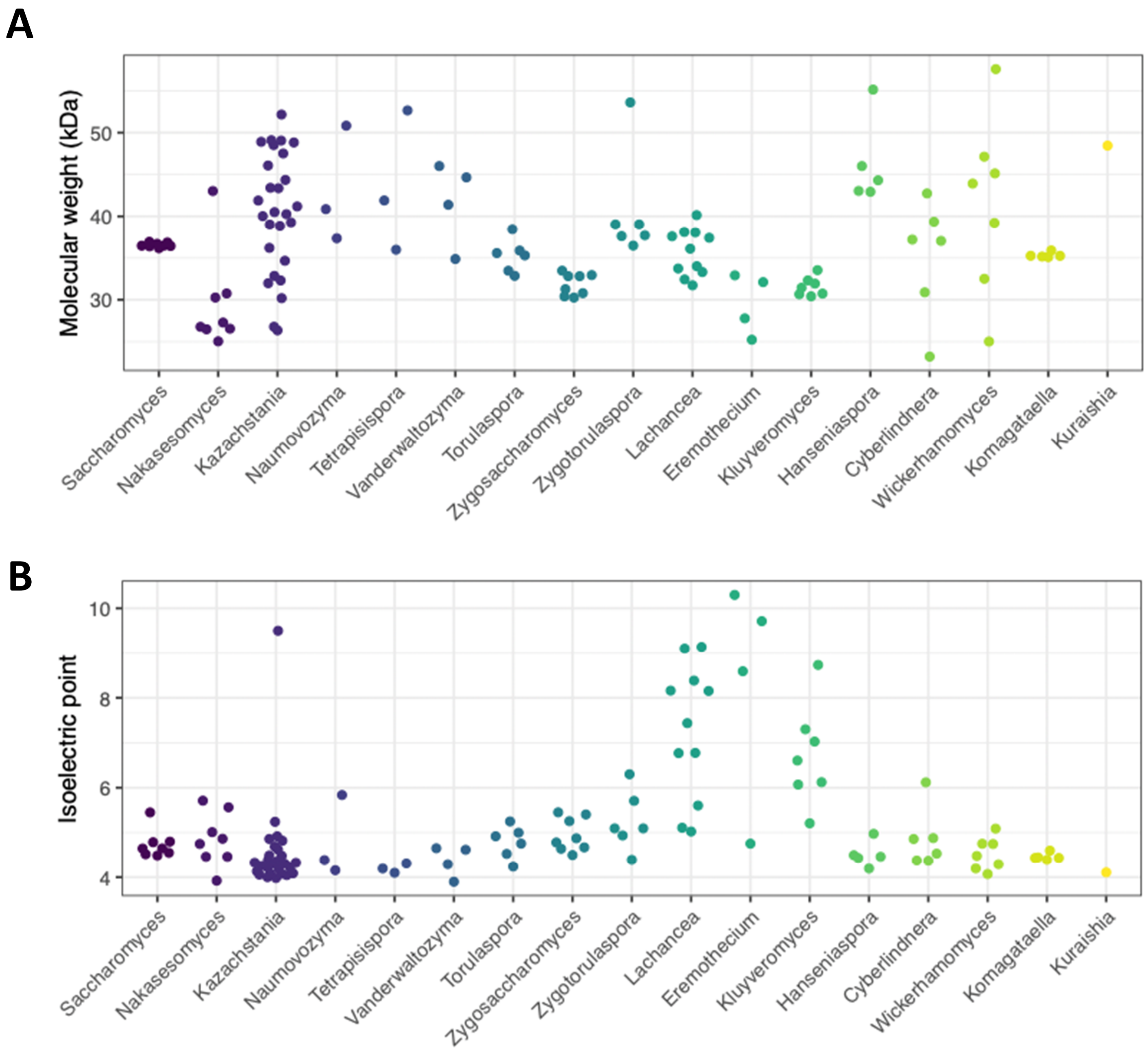


**Figure.** S3

### Supplemental Figure S4

#### Kramer et al.

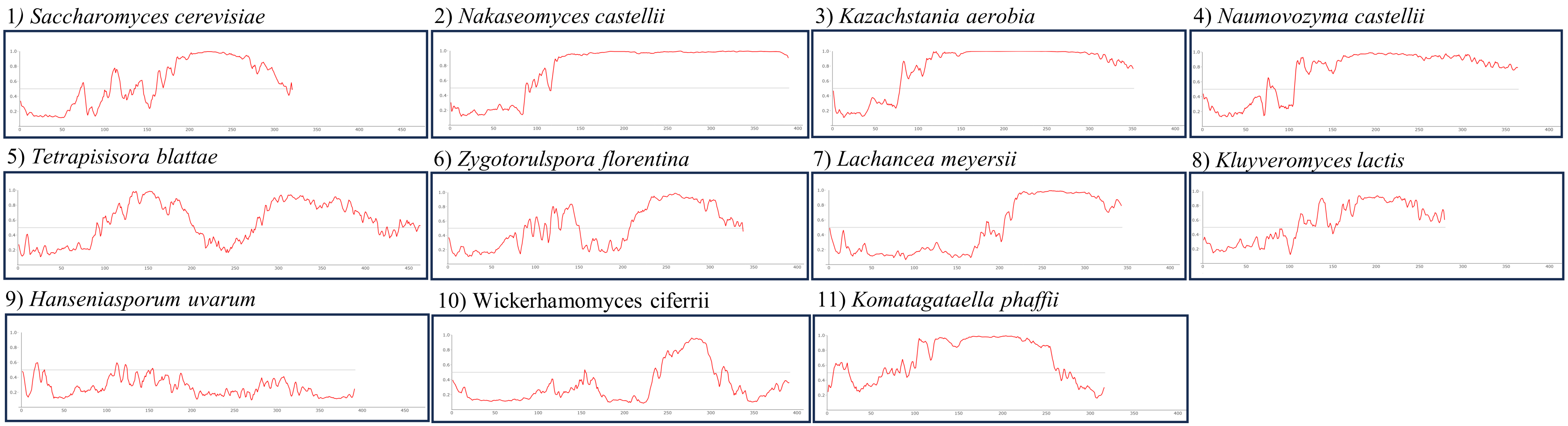


#### Figure. S4

### Supplemental Figure S5

#### Kramer et al.

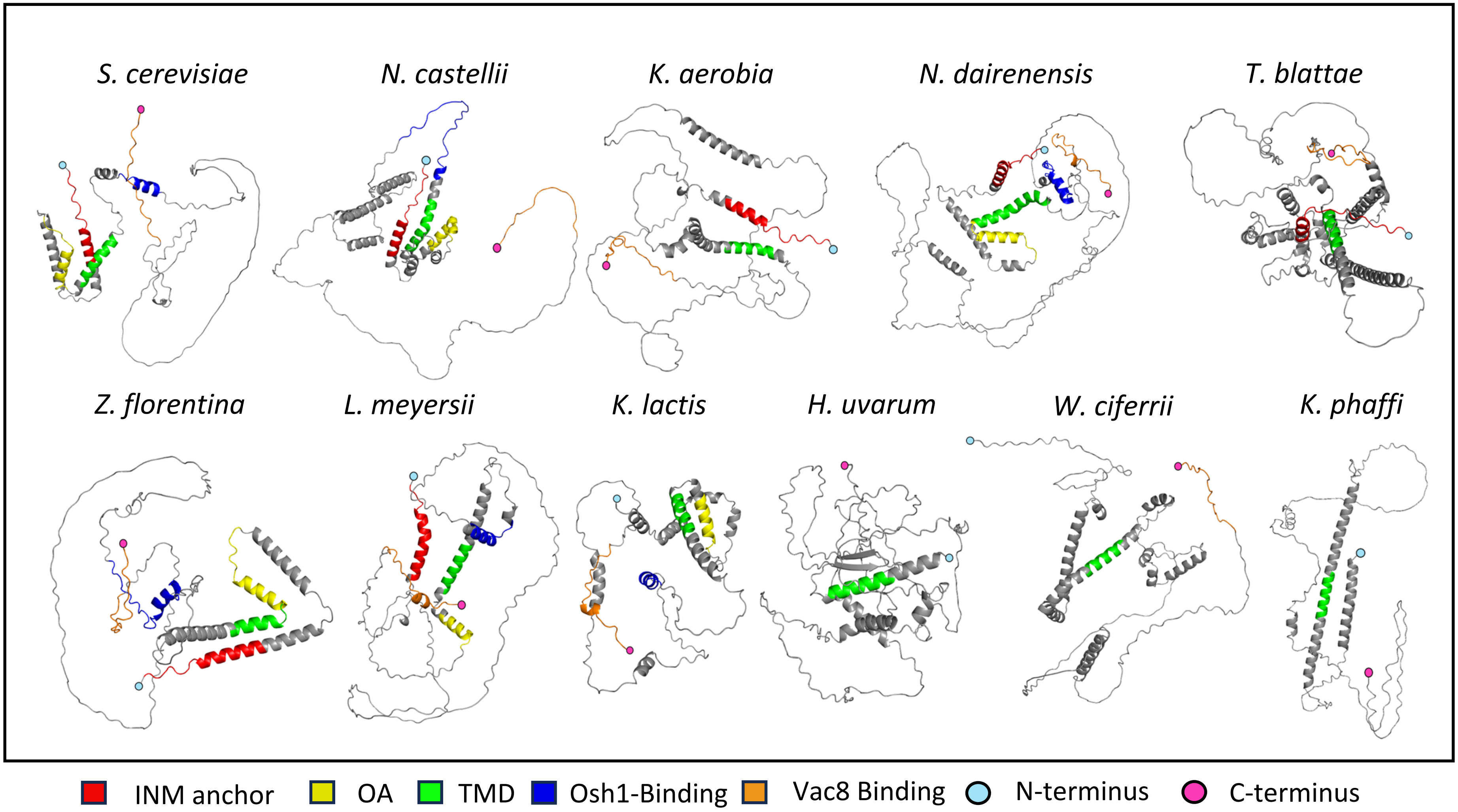


**Figure.** S5

### Supplemental Figure S6

Kramer et al.

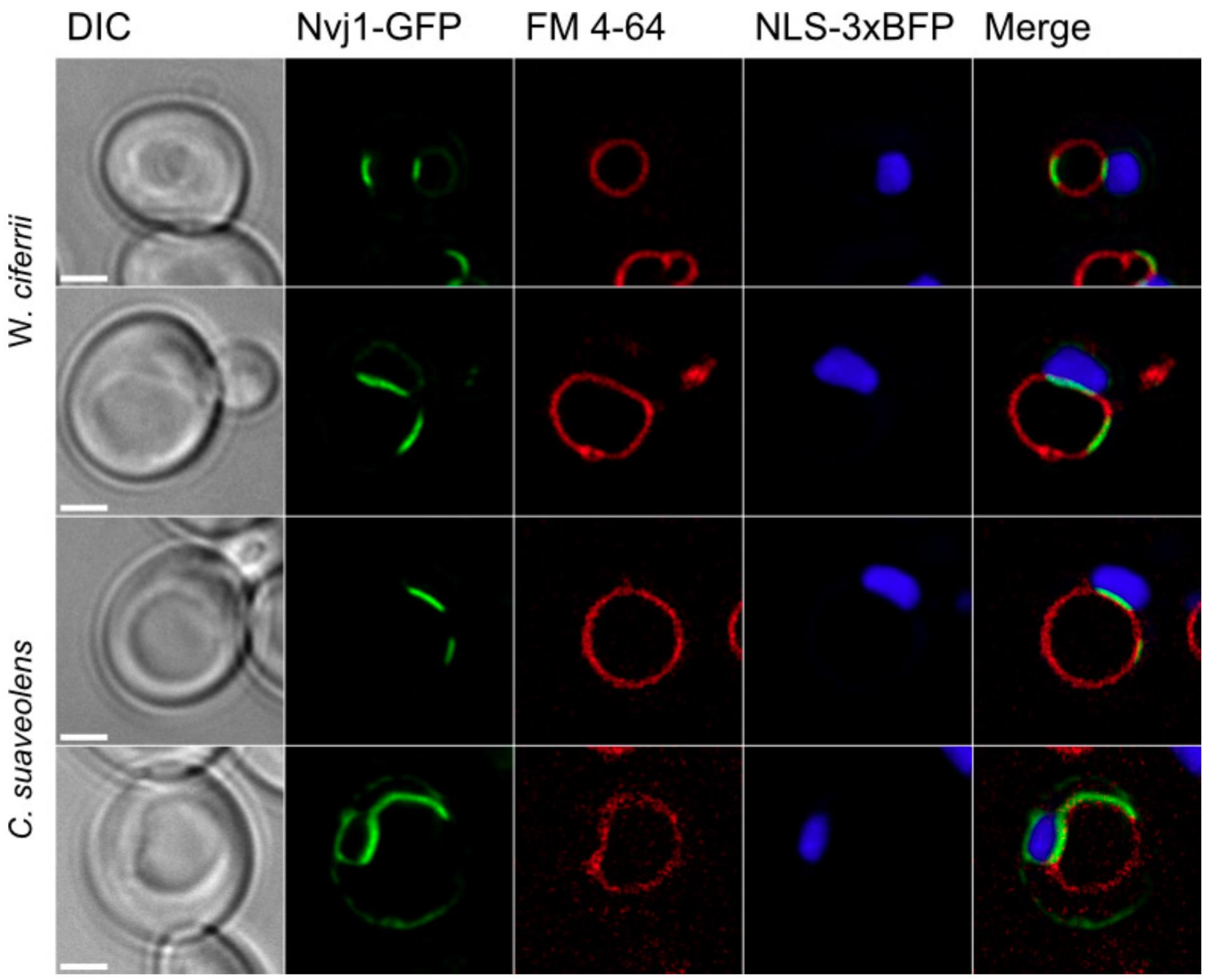


**Figure.** S6

### Supplemental Figure S7

#### Kramer et al.

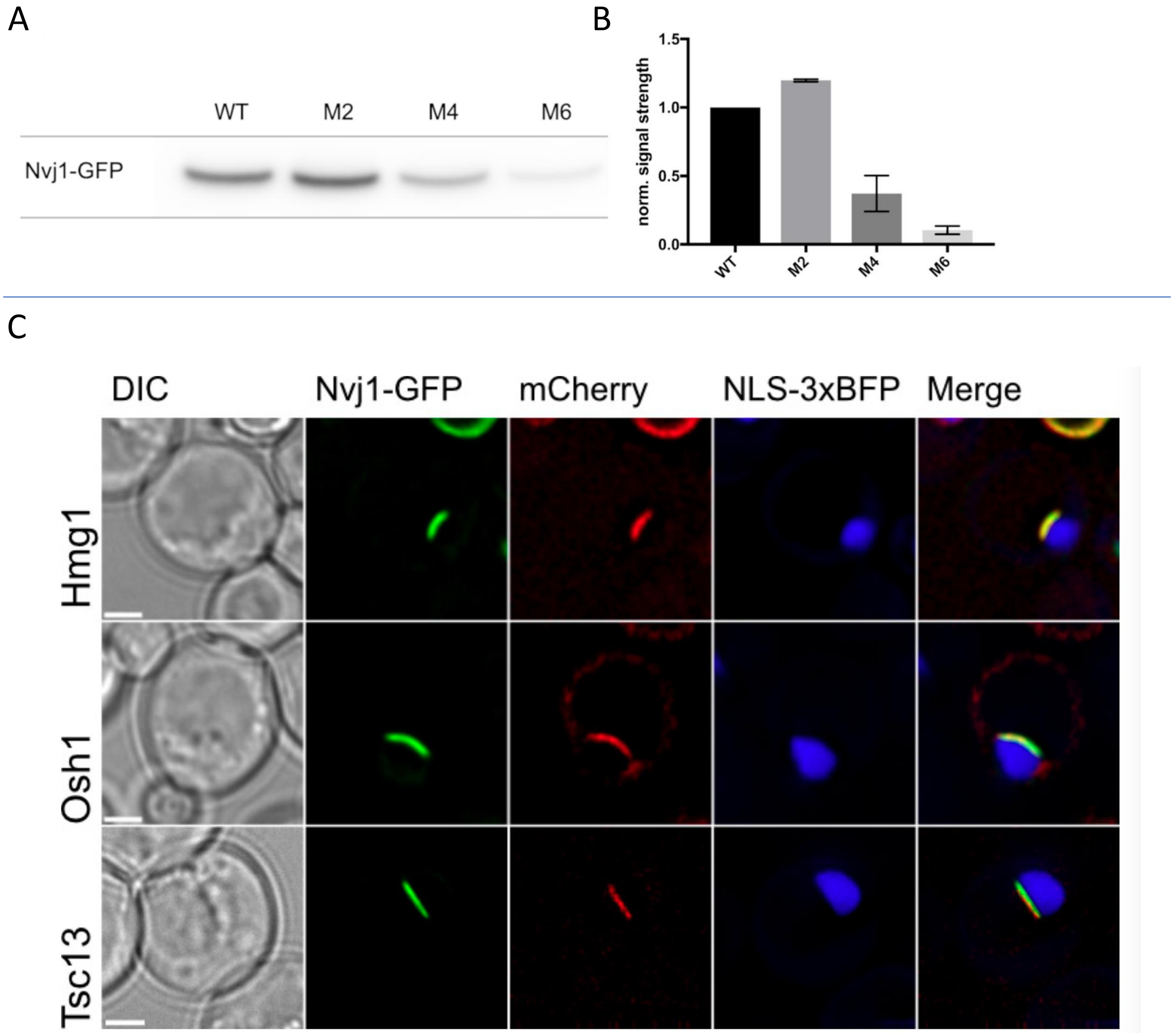


**Figure.** S7

### Supplemental Figure S8

#### Kramer et al.

**A**

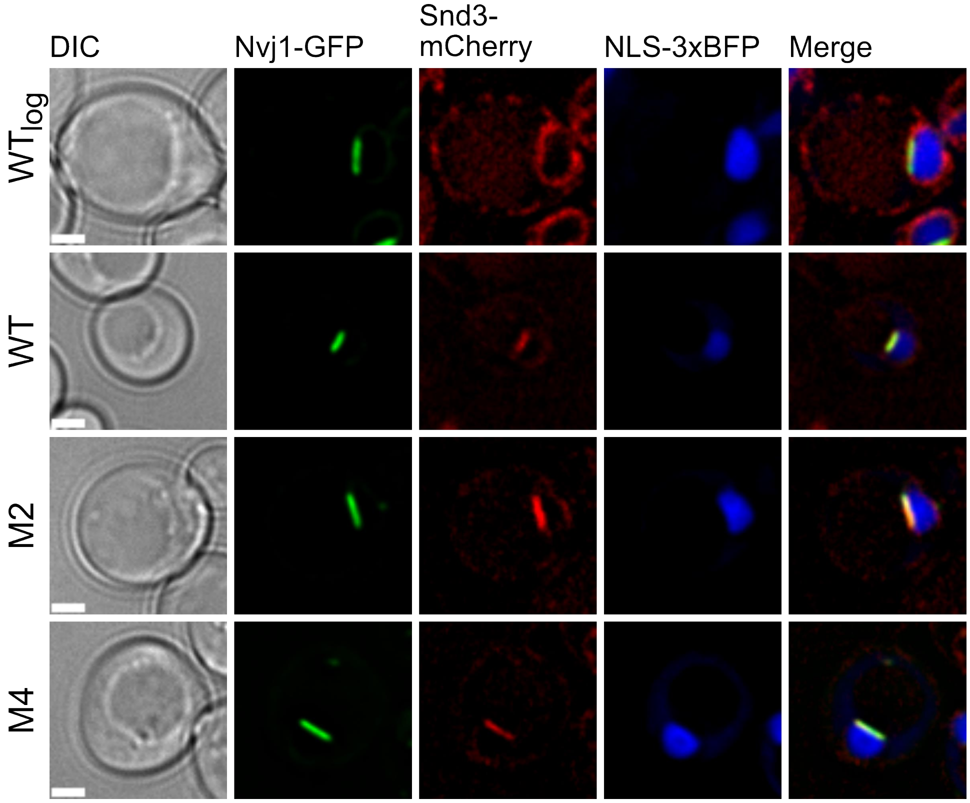


**B**

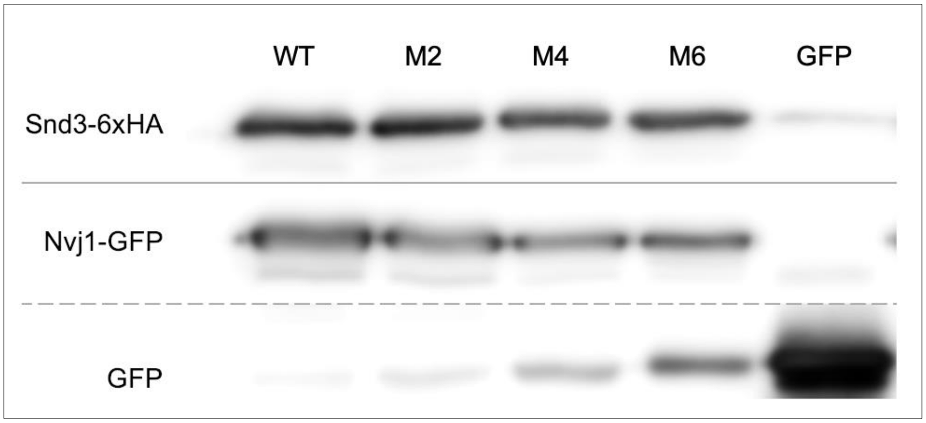


**C**

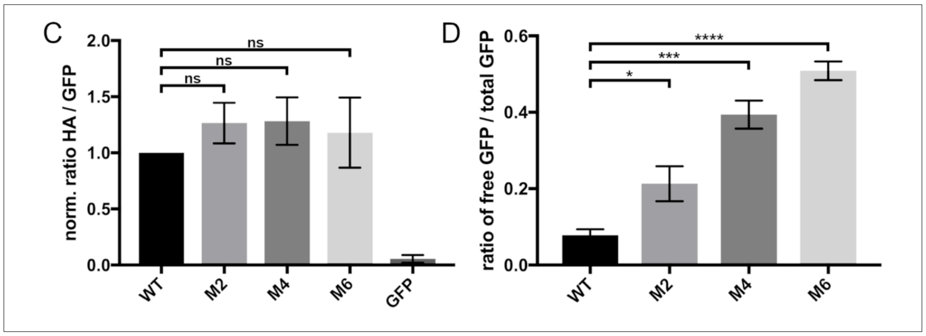


**Figure.** S8
