## Supplemental Table S2 for "Functional diversification across yeast lineages of the nucleus-vacuole junction forming protein Nvj1"

| Species Name | seqID | HMMProfileID | Y1000_TreeID | Y1000_DatasetID | Y1000_protID | ncbiID | group | inTableS1 | inY1000Hmmer | Copy Number | syntenyCheck | syntenyComments |
|---|---|---|---|---|---|---|---|---|---|---|---|---|
| saccharomycescerevisiae | saccharomyces_cerevisiae | YHR195W | Saccharomyces_cerevisiae | saccharomyces_cerevisiae | g005344.m1 | NP_012065.3 | Saccharomyces | 1 | 1 | 1 | MDM31-NVJ1-UTP9 | |
| saccharomycesjurei | saccharomyces_jurei | #N/A | Saccharomyces_jurei | saccharomyces_jurei | g004292.m1 | #N/A | Saccharomyces | 1 | 1 | 1 | MDM31-NVJ1-UTP9 | |
| saccharomycesparadoxus | saccharomyces_paradoxus | genemark-Spar_8-processed-gene-4.108 | Saccharomyces_paradoxus | saccharomyces_paradoxus | g005237.m1 | #N/A | Saccharomyces | 1 | 1 | 1 | MDM31-NVJ1-UTP9 | |
| saccharomyceskudriavzevii | saccharomyces_kudriavzevii | augustus_masked-Skud_8-processed-gene-3.158 | Saccharomyces_kudriavzevii | saccharomyces_kudriavzevii | g005184.m1 | EJT43435.1 | Saccharomyces | 1 | 1 | 1 | MDM31-NVJ1-UTP9 | |
| saccharomycesarboricola | saccharomyces_arboricola | augustus_masked-NC_026178.1-processed-gene-4.111 | Saccharomyces_arboricola | saccharomyces_arboricola | g005224.m1 | EJS43367.1 | Saccharomyces | 1 | 1 | 1 | MDM31-NVJ1-UTP9 | |
| saccharomycesmikatae | saccharomyces_mikatae | snap_masked-Smik_8-processed-gene-4.133 | Saccharomyces_mikatae | saccharomyces_mikatae | g005189.m1 | #N/A | Saccharomyces | 1 | 1 | 1 | MDM31-NVJ1-UTP9 | |
| saccharomycesuvarum | saccharomyces_uvarum | augustus_masked-Sbay_15-processed-gene-6.183 | Saccharomyces_uvarum | saccharomyces_uvarum | g002217.m1 | #N/A | Saccharomyces | 1 | 1 | 1 | MDM31-NVJ1-UTP9 | |
| saccharomyceseubayanus | saccharomyces_eubayanus | augustus_masked-chrXV-processed-gene-6.90 | Saccharomyces_eubayanus | saccharomyces_eubayanus | g005249.m1 | XP_018221971.1 | Saccharomyces | 1 | 1 | 1 | MDM31-NVJ1-UTP9 | |
| Nakasesomycesglabrata | Nakasesomyces_glabrata | #N/A | Nakaseomyces_glabratus | yHMPu5000034723_candida_glabrata_180604 | g000308.m1 | XP_449661.1 | Nakasesomyces | 1 | 1 | 1 | MDM31-NVJ1-UTP9 | |
| Nakaseomyceskungkrabaensis | Nakaseomyces_kungkrabaensis | #N/A | Nakaseomyces_kungkrabaensis | yHMPu5000034722_candida_kungkrabaensis_180604 | g004198.m1 | #N/A | Nakasesomyces | 1 | 1 | 1 | MDM31-NVJ1-UTP9 | |
| Nakaseomycesnivariensis | Nakaseomyces_nivariensis | snap_masked-CANI0S17-processed-gene-4.132 | Nakaseomyces_nivariensis | yHMPu5000034720_candida_nivariensis_180604 | g001471.m1 | #N/A | Nakasesomyces | 1 | 1 | 1 | MDM31-NVJ1-UTP9 | |
| Nakaseomycesuthaithanina | Nakaseomyces_uthaithanina | #N/A | Nakaseomyces_uthaithaninus | yHMPu5000034718_candida_uthaithanina_180604 | g005380.m1 | #N/A | Nakasesomyces | 1 | 1 | 1 | UTP9?-NVJ1-RTT105? | |
| nakaseomycesdelphensis | nakaseomyces_delphensis | augustus_masked-NADE0S27-processed-gene-3.39 | Nakaseomyces_delphensis | yHMPu5000026127_nakaseomyces_delphensis_160519 | g004209.m1 | CAO98811.1 | Nakasesomyces | 1 | 1 | 1 | MDM31-NVJ1-UTP9 | |
| nakaseomycesbracarensis | nakaseomyces_bracarensis | snap_masked-CABR0S29-processed-gene-6.0 | Nakaseomyces_bracarensis | yHMPu5000034725_candida_bracarensis_160928 | g001614.m1 | #N/A | Nakasesomyces | 1 | 1 | 1 | MDM31-NVJ1-UTP9 | |
| nakaseomycesbacillisporus | nakaseomyces_bacillisporus | augustus_masked-NABA0S29-processed-gene-0.108 | Nakaseomyces_bacillisporus | yHMPu5000035666_nakaseomyces_bacillisporus_160613 | g004314.m1 | #N/A | Nakasesomyces | 1 | 1 | 1 | MDM31-NVJ1-UTP9 | |
| nakaseomycescastellii_ | nakaseomyces_castellii_ | snap_masked-CACA0S33-processed-gene-9.107 | Nakaseomyces_castellii | yHMPu5000034724_candida_castellii_180604 | g001576.m1 | #N/A | Nakasesomyces | 1 | 1 | 1 | MDM31-NVJ1-UTP9 | YGOB disagrees |
| Nakaseomyces_sp._yHDO568 | Nakaseomyces_sp._yHDO568 | N/A | Nakaseomyces_sp._yHDO568 | yHDO568_nakaseomyces_sp_190924 | g004296.m1 | N/A | Nakasesomyces | 0 | 1 | 1 | MDM31-NVJ1-UTP9 | |
| kazachstaniasolicola | kazachstania_solicola | augustus_masked-scf7180000046332-processed-gene-1.206 | Kazachstania_solicola | yHAB159_kazachstania_solicola_160519.haplomerger2 | g005068.m1 | #N/A | Kazachstania | 1 | 0 | 1 | MDM31-NVJ1-RIX1? | UTP9 pseudpgenized? |
| kazachstaniaaerobia | kazachstania_aerobia | augustus_masked-scf7180000020169-processed-gene-0.188 | Kazachstania_aerobia | yHAB164_kazachstania_aerobia_160519 | g002718.m1 | #N/A | Kazachstania | 1 | 0 | 1 | MDM31-NVJ1-RIX1? | needs validation |
| kazachstaniaunispora | kazachstania_unispora | snap_masked-NODE_47_length_78712_cov_50.1529_ID_93-processed-gene-0.49 | Kazachstania_unispora | yHAB133_kazachstania_unispora_160519 | g003649.m1 | #N/A | Kazachstania | 1 | 0 | 1 | MDM31-NVJ1-RIX1? | needs validation |
| kazachstania_bromeliacearum | kazachstania_bromeliacearum | snap_masked-NODE_10_length_278180_cov_22.5073_ID_19-processed-gene-1.127 | Kazachstania_bromeliacearum | yHAB136_kazachstania_bromeliacearum_160519 | g000451.m1 | #N/A | Kazachstania | 1 | 1 | 1 | MDM31-NVJ1-RIX1? | needs validation |
| kazachstania_bulderi | kazachstania_bulderi | #N/A | Kazachstania_bulderi | yHAB157_kazachstania_bulderi_160519.haplomerger2 | g005958.m1 | #N/A | Kazachstania | 1 | 1 | 1 | MDM31-NVJ1-RIX1? | needs validation |
| kazachstania_exigua | kazachstania_exigua | #N/A | Kazachstania_exigua | yHAB150_kazachstania_exigua_160519 | g002519.m1 | #N/A | Kazachstania | 1 | 1 | 2 | MDM31-NVJ1-RIX1? | needs validation |
| kazachstania_exigua | kazachstania_exigua | #N/A | Kazachstania_exigua | yHAB150_kazachstania_exigua_160519 | g000048.m1 | #N/A | Kazachstania | 0 | 1 | 2 | MDM31-NVJ1-MRX19 | needs validation |
| kazachstania_gamospora | kazachstania_gamospora | #N/A | Kazachstania_gamospora | yHAB142_kazachstania_gamospora_170307.haplomerger2 | g004167.m1 | #N/A | Kazachstania | 1 | 1 | | MCM31-NVJ1-? | contig breakpoint in y1000 main assembly |
| kazachstania_hellenica | kazachstania_hellenica | #N/A | Kazachstania_hellenica | yHAB145_kazachstania_hellenica_160519.haplomerger2 | g003117.m1 | #N/A | Kazachstania | 1 | 1 | | MDM31-NVJ1-RIX1? | |
| kazachstania_aquatica | Missing! | #N/A | Kazachstania_aquatica | yHAB165_kazachstania_aquatica_160519 | g003032.m1 | #N/A | Kazachstania | 1 | 1 | | MDM31-NVJ1-RIX1? | |
| kazachstania_kunashirensis | kazachstania_kunashirensis | snap_masked-scf7180000008985-processed-gene-1.61 | Kazachstania_kunashirensis | yHAB160_kazachstania_kunashirensis_160519 | g005050.m1 | #N/A | Kazachstania | 1 | 1 | | MDM31-NVJ1-RIX1? | |
| kazachstania_lodderae | kazachstania_lodderae | #N/A | Kazachstania_lodderae | yHAB138_kazachstania_lodderae_160519 | g003605.m1 | #N/A | Kazachstania | 1 | 1 | | MDM31-NVJ1-RIX1? | |
| kazachstania_piceae | kazachstania_piceae | #N/A | Kazachstania_piceae | yHAB156_kazachstania_piceae_160519 | g005225.m1 | #N/A | Kazachstania | 1 | 1 | | MDM31-NVJ1-RIX1? | |
| kazachstania_saulgeensis | kazachstania_saulgeensis | #N/A | Kazachstania_saulgeensis | yHMPu5000037225_kazachstania_saulgeensis_210210 | g000550.m1 | SMN19139.1 | Kazachstania | 1 | 1 | | MDM31-NVJ1-RIX1? | |
| kazachstania_servazzi | kazachstania_servazzi | #N/A | Kazachstania_servazzii | yHAB151_kazachstania_servazzi_160519 | g000773.m1 | #N/A | Kazachstania | 1 | 1 | | MDM31-NVJ1-RIX1? | |
| kazachstania_sinensis | kazachstania_sinensis | #N/A | Kazachstania_sinensis | yHAB161_kazachstania_sinensis_160519 | g005599.m1 | #N/A | Kazachstania | 1 | 1 | | MDM31-NVJ1-RIX1? | |
| kazachstania_yasuniensis | kazachstania_yasuniensis | #N/A | Kazachstania_yasuniensis | yHMPu5000034708_kazachstania_yasuniensis_180604 | g001209.m1 | #N/A | Kazachstania | 1 | 1 | | MDM31-NVJ1-RIX1? | |
| Kazachstania_barnettii | Kazachstania_barnettii | #N/A | Kazachstania_barnettii | yHAB162_Kazachstania_barnettii_SPADES | g001263.m1 | #N/A | Kazachstania | 1 | 1 | | MDM31-NVJ1-RIX1? | |
| Kazachstania_psychrophila | Kazachstania_psychrophila | #N/A | Kazachstania_psychrophila | yHMPu5000034706_Kazachstania_psychrophila_SPADES | g002942.m1 | #N/A | Kazachstania | 1 | 1 | | MDM31-NVJ1-RIX1? | |
| Kazachstania_jinghongensis | Kazachstania_jinghongensis | #N/A | Kazachstania_jinghongensis | yHMPu5000037210_Kazachstania_jinghongensis_SPADES | g001801.m1 | #N/A | Kazachstania | 1 | 1 | | MDM31-NVJ1-RIX1? | |
| kazachstaniasiamensis | kazachstania_siamensis | augustus_masked-NODE_61_length_63804_cov_14.6622_ID_121-processed-gene-0.80 | Kazachstania_siamensis | yHAB143_kazachstania_siamensis_160519 | g004344.m1 | #N/A | Kazachstania | 1 | 1 | | MDM31-NVJ1-RIX1? | |
| kazachstaniataianensis | kazachstania_taianensis | augustus_masked-NODE_5_length_452406_cov_49.1731_ID_9-processed-gene-3.120 | Kazachstania_taianensis | yHAB147_kazachstania_taianensis_160519 | g004007.m1 | #N/A | Kazachstania | 1 | 1 | | MDM31-NVJ1-RIX1? | |
| kazachstanianaganishii | kazachstania_naganishii | snap_masked-HE978326-processed-gene-0.118 | Kazachstania_naganishii | kazachstania_naganishii | g001440.m1 | XP_022467125.1 | Kazachstania | 1 | 1 | | MDM31-NVJ1-RIX1? | |
| kazachstaniamartiniae | kazachstania_martiniae | snap_masked-scf7180000030514-processed-gene-0.103 | Kazachstania_martiniae | yHAB132_kazachstania_martiniae_160519 | g004439.m1 | #N/A | Kazachstania | 1 | 1 | | MDM31-NVJ1-RIX1? | |
| kazachstaniaturicensis | kazachstania_turicensis | snap_masked-scf7180000049581-processed-gene-4.63 | Kazachstania_turicensis | yHMPu5000040961_kazachstania_turicensis_201018 | g004713.m1 | #N/A | Kazachstania | 1 | 1 | | MDM31-NVJ1-RIX1? | |
| kazachstaniakunashirensis | kazachstania_kunashirensis | snap_masked-scf7180000008985-processed-gene-1.61 | Kazachstania_kunashirensis | yHAB160_kazachstania_kunashirensis_160519 | g005050.m1 | #N/A | Kazachstania | 1 | 1 | | MDM31-NVJ1-RIX1? | |
| kazachstaniaspencerorum | kazachstania_spencerorum | snap_masked-scf7180000055600-processed-gene-0.48 | Kazachstania_spencerorum | yHAB155_kazachstania_spencerorum_160519 | g002874.m1 | #N/A | Kazachstania | 1 | 0 | | MDM31-NVJ1-RIX1? | |
| kazachstaniaafricana | kazachstania_africana | augustus_masked-scaffold_2-processed-gene-13.205 | Kazachstania_africana | yHAB137_kazachstania_africana_160519 | g003755.m1 | XP_003956120.1 | Kazachstania | 1 | 1 | | MDM31-NVJ1-? | suspicious gap downstream |
| kazachstaniaviticola | kazachstania_viticola | snap_masked-scf7180000050361-processed-gene-0.67 | Kazachstania_viticola | yHAB158_kazachstania_viticola_160519.haplomerger2 | g001183.m1 | #N/A | Kazachstania | 1 | 1 | | MDM31-NVJ1-RIX1? | |
| Kazachtania_humilis | Kazachstania_humilis | | Kazachstania_humilis | yHAB152_candida_humilis_160519.haplomerger2 | g000693.m1 | | | 0 | 1 | | MDM31-NVJ1-RIX1? | |
| Kazachtania_pseudohumilis | Kazachtania_pseudohumilis | | Kazachtania_pseudohumilis | yHMPu5000034719_candida_pseudohumilis_180604.haplomerger2 | g002365.m1 | | | 0 | 1 | | MDM31-NVJ1-RIX1? | |
| Kazachstania_rosinii | Kazachstania_rosinii | | Kazachstania_rosinii | yHAB153_kazachstania_rosinii_160519 | g001213.m1 | | | 0 | 1 | | MDM31-NVJ1-? | downstream gene is small |
| Kazachstania_sp._UFMG-CM-Y273 | Kazachstania_sp._UFMG-CM-Y273 | | Kazachstania_sp._UFMG-CM-Y273 | yHDO576_kazachstania_sp_180604 | g000466.m1 | | | 0 | 1 | | MDM31-NVJ1-RIX1? | |
| naumovozymadairenensis | naumovozyma_dairenensis | augustus_masked-NC_016482.1-processed-gene-7.22 | Naumovozyma_dairenensis | yHMPu5000034872_naumovozyma_dairenensis_180604 | g002752.m1 | XP_003669889.1 | Naumovozyma | 1 | 1 | | MDM31-NVJ1-RIX1? | |
| naumovozymacastellii | naumovozyma_castellii | snap_masked-HE576752-processed-gene-12.71 | Naumovozyma_castellii | yHMPu5000034871_naumovozyma_castellii_180604 | g001098.m1 | #N/A | Naumovozyma | 1 | 1 | | MDM31-NVJ1-RIX1? | |
| naumovozyma_baii | naumovozyma_baii | #N/A | Naumovozyma_baii | yHMPu5000034894_naumovozyma_baii_190924 | g000794.m1 | #N/A | Naumovozyma | 1 | 1 | | MDM31-NVJ1-RIX1? | |
| tetrapisisporairiomotensis | tetrapisispora_iriomotensis | augustus_masked-flattened_line_57-processed-gene-0.130 | Tetrapisispora_iriomotensis | yHMPu5000034876_Tetrapisispora_iriomotensis_SPADES | g003416.m1 | #N/A | Tetrapisispora | 1 | 1 | | MDM31-NVJ1-NMD5-UTP9 | |
| tetrapisisporanamnaonensis | tetrapisispora_namnaonensis | #N/A | Tetrapisispora_namnaoensis | yHMPu5000034877_tetrapisispora_namnaonensis_160519 | g001993.m1 | #N/A | Tetrapisispora | 1 | 0 | | MDM31-NVJ1-NMD5-UTP9 | synteny graph hits wrong MDM31 |
| tetrapisisporablattae | tetrapisispora_blattae | snap_masked-NC_020185-processed-gene-25.62 | Tetrapisispora_blattae | yHMPu5000034874_tetrapisispora_blattae_190924 | g003018.m1 | XP_004178357.1 | Tetrapisispora | 1 | 0 | | PEX14-NVJ1-UTP9 | No MDM31 in vicinity |
| vanderwaltozymapolyspora | vanderwaltozyma_polyspora | snap_masked-NW_001834637.1-processed-gene-0.82 | Vanderwaltozyma_polyspora | yHMPu5000034869_vanderwaltozyma_polyspora_180604 | g005365.m1 | XP_001643905.1 | Vanderwaltozyma | 1 | 1 | | MDM31-NVJ1-NMD5-UTP9 | |
| Vanderwaltozyma_yarrowii | Vanderwaltozyma_yarrowii | #N/A | Vanderwaltozyma_yarrowii | yHMPu5000034868_Vanderwaltozyma_yarrowii_S33_SPADES | g001449.m1 | #N/A | Vanderwaltozyma | 1 | 1 | | MDM31-NVJ1-NMD5-UTP9 | |
| vanderwaltozyma_tropicalis | vanderwaltozyma_tropicalis | #N/A | Vanderwaltozyma_tropicalis | yHMPu5000026257_vanderwaltozyma_tropicalis_190924 | g003717.m1 | #N/A | Vanderwaltozyma | 1 | 1 | | MDM31-NVJ1-NMD5-UTP9 | |
| Vanderwaltozyma_verrucispora | Vanderwaltozyma_verrucispora | #N/A | Vanderwaltozyma_verrucispora | yHMPu5000037837_Vanderwaltozyma_verrucispora_S176_SPADES | g005143.m1 | #N/A | Vanderwaltozyma | 1 | 1 | | MDM31-NVJ1-NMD5-UTP9 | |
| torulasporapretoriensis | torulaspora_pretoriensis | augustus_masked-flattened_line_3-processed-gene-4.12 | Torulaspora_pretoriensis | yHMPu5000034881_Torulaspora_pretoriensis_SPADES | g004938.m1 | #N/A | Torulaspora | 1 | 1 | | MDM31-NVJ1-NMD5-UTP9 | |
| torulasporamicroellipsoides | torulaspora_microellipsoides | augustus_masked-flattened_line_52-processed-gene-0.132 | Torulaspora_microellipsoides | yHMPu5000035651_Torulaspora_microellipsoides_SPADES | g002639.m1 | #N/A | Torulaspora | 1 | 1 | | MDM31-NVJ1-FEN2? | weird gap after MDM31 |
| torulasporafranciscae | torulaspora_franciscae | snap_masked-flattened_line_10-processed-gene-1.40 | Torulaspora_franciscae | yHMPu5000026152_Torulaspora_franciscae_SPADES | g004752.m1 | #N/A | Torulaspora | 1 | 1 | | MDM31-NVJ1-NMD5-UTP9 | |
| torulasporadelbrueckii | torulaspora_delbrueckii | snap_masked-scaffold_4-processed-gene-1.34 | Torulaspora_delbrueckii | yHMPu5000035653_torulaspora_delbrueckii_160613 | g001776.m1 | XP_003680850.1 | Torulaspora | 1 | 1 | | MDM31-NVJ1-NMD5-UTP9 | |
| torulaspora_globosa | torulaspora_globosa | #N/A | Torulaspora_globosa | yHMPu5000034880_torulaspora_globosa_160519 | g001826.m1 | #N/A | Torulaspora | 1 | 1 | | MDM31-NVJ1-NMD5-UTP9 | |
| torulaspora_maleeae | torulaspora_maleeae | snap_masked-NODE_5_length_764704_cov_59.617_ID_1253-processed-gene-6.109 | Torulaspora_maleeae | yHMPu5000035652_torulaspora_maleeae_160613 | g003867.m1 | #N/A | Torulaspora | 1 | 1 | | MDM31-NVJ1-NMD5-UTP9 | |
| Torulaspora_sp._yHMJ407 | Torulaspora_sp._yHMJ407 | | Torulaspora_sp._yHMJ407 | yHMJ407_Torulaspora_sp_nov_plate34_SPADES | g002906.m1 | | | 0 | 1 | | MDM31-NVJ1-NMD5-UTP9 | |
| zygosaccharomyces_bisporus | zygosaccharomyces_bisporus | snap_masked-flattened_line_16-processed-gene-0.27 | Zygosaccharomyces_bisporus | yHMPu5000034866_zygosaccharomyces_bisporus_160519 | g005101.m1 | #N/A | Zygosaccharomyces | 1 | 1 | | MDM31-NVJ1-NMD5-UTP9 | |
| zygosaccharomyces_gambellarensis | zygosaccharomyces_gambellarensis | #N/A | Zygosaccharomyces_gambellarensis | yHDO565_zygosaccharomyces_gambellarensis_180604 | g005022.m1 | #N/A | Zygosaccharomyces | 1 | 1 | | MDM31-NVJ1-NMD5-UTP9 | |
| zygosaccharomycesbailii | Zygosaccharomyces_bailii | augustus_masked-ZYBA0S10-processed-gene-3.53 | Zygosaccharomyces_bailii | zygosaccharomyces_bailii | g001048.m1 | CDF91308.1 | Zygosaccharomyces | 1 | 1 | | MDM31-NVJ1-NMD5-UTP9 | |

| Species Name | seqID | HMMProfileID | Y1000_TreeID | Y1000_DatasetID | Y1000_protID | ncbiID | group | inTableS1 | inY1000Hmmer | CopyNumber | syntenyCheck | syntenyComments |
|---|---|---|---|---|---|---|---|---|---|---|---|---|
| zygosaccharomyceskombuchaensis | *Zygosaccharomyces_kombuchaensis* | *snap_masked-NODE_1_length_271579_cov_10.1807_ID_1-processed-gene-1.23* | *Zygosaccharomyces_kombuchaensis* | *yHMPu5000034865_zygosaccharomyces_kombuchaensis_160519* | *g000063.m1* | *#N/A* | *Zygosaccharomyces* | 1 | 1 | | MDM31-NVJ1-NMD5-UTP9 | |
| zygosaccharomyces_lentus | zygosaccharomyces_lentus | *#N/A* | *Zygosaccharomyces_lentus* | *yHMPu5000034864_zygosaccharomyces_lentus_170307* | *g004388.m1* | *#N/A* | *Zygosaccharomyces* | 1 | 1 | | MDM31-NVJ1-NMD5-UTP9 | |
| zygosaccharomyces_mellis | zygosaccharomyces_mellis | *#N/A* | *Zygosaccharomyces_mellis* | *yHDO572_zygosaccharomyces_mellis_180604* | *g002135.m1* | *#N/A* | *Zygosaccharomyces* | 1 | 1 | | MDM31-NVJ1-NMD5-UTP9 | |
| zygosaccharomyces_pseudorouxii | zygosaccharomyces_pseudorouxii | *#N/A* | *Zygosaccharomyces_pseudorouxii* | *yHMPu5000037836_zygosaccharomyces_pseudorouxii_201018.haplomerger2* | *g004410.m1* | *#N/A* | *Zygosaccharomyces* | 1 | 1 | | MDM31-NVJ1-NMD5-UTP9 | |
| zygosaccharomyces_siamensis | zygosaccharomyces_siamensis | *#N/A* | *Zygosaccharomyces_siamensis* | *yHMPu5000035626_zygosaccharomyces_siamensis_190924* | *g000372.m1* | *#N/A* | *Zygosaccharomyces* | 1 | 1 | | MDM31-NVJ1-NMD5-UTP9 | |
| Zygosaccharomyces_parabailii | Zygosaccharomyces_parabailii | | Zygosaccharomyces_parabailii | yHMPu5000037834_zygosaccharomyces_parabailii_190924 | g003412.m1 | | | 0 | 1 | | MDM31-NVJ1-NMD5-UTP9 | |
| Zygosaccharomyces_pseudobailii | Zygosaccharomyces_pseudobailii | | Zygosaccharomyces_pseudobailii | yHMPu5000035627_zygosaccharomyces_pseudobailii_201018 | g002625.m1 | | | 0 | 1 | | MDM31-NVJ1-? | small contig |
| Zygosaccharomyces_pseudobailii | Zygosaccharomyces_pseudobailii | | Zygosaccharomyces_pseudobailii | yHMPu5000035627_zygosaccharomyces_pseudobailii_201018 | g003112.m1 | | | 0 | 1 | | ?-NVJ1-NMD5 | |
| Zygosaccharomyces_rouxii | Zygosaccharomyces_rouxii | | Zygosaccharomyces_rouxii | *yHMPu5000034863_zygosaccharomyces_rouxii_180604* | g001572.m1 | | | 0 | 1 | | MDM31-NVJ1-NMD5-UTP9 | |
| Zygosaccharomyces_sapae | Zygosaccharomyces_sapae | | Zygosaccharomyces_sapae | *yHDO603_zygosaccharomyces_sapae_190924.haplomerger2* | g000017.m1 | | | 0 | 1 | | MDM31-NVJ1-NMD5-UTP9 | |
| zygotorulasporaflorentina | *zygotorulaspora_florentina* | *augustus_masked-NODE_6_length_562868_cov_24.3835_ID_11-processed-gene-2.47* | *Zygotorulaspora_florentina* | *yHMPu5000034862_zygotorulaspora_florentina_160519* | *g004327.m1* | *#N/A* | *Zygotorulaspora* | 1 | 1 | | MDM31-NVJ1-NMD5-UTP9 | |
| zygotorulasporamrakii | *zygotorulaspora_mrakii* | *snap_masked-NODE_2_length_708340_cov_18.0358_ID_2767-processed-gene-4.105* | *Zygotorulaspora_mrakii* | *yHMPu5000026256_zygotorulaspora_mrakii_170307* | *g002003.m1* | *#N/A* | *Zygotorulaspora* | 1 | 1 | | MDM31-NVJ1-NMD5-UTP9 | |
| zygotorulaspora_chibaensis | zygotorulaspora_chibaensis | *#N/A* | *Zygotorulaspora_chibaensis* | *yHMPu5000037204_zygotorulaspora_chibaensis_210210* | *g002973.m1* | *#N/A* | *Zygotorulaspora* | 1 | 1 | | MDM31-NVJ1-NMD5-UTP9 | |
| zygotorulaspora_danielsina | zygotorulaspora_danielsina | *#N/A* | *Zygotorulaspora_danielsina* | *yHMPu5000037205_zygotorulaspora_danielsina_210210* | *g002121.m1* | *#N/A* | *Zygotorulaspora* | 1 | 1 | | MDM31-NVJ1-NMD5-UTP9 | |
| zygotorulaspora_mrakii | zygotorulaspora_mrakii | *snap_masked-NODE_2_length_708340_cov_18.0358_ID_2767-processed-gene-4.105* | *Zygotorulaspora_mrakii* | *yHMPu5000026256_zygotorulaspora_mrakii_170307* | *g002003.m1* | *#N/A* | *Zygotorulaspora* | 1 | 1 | | MDM31-NVJ1-NMD5-UTP9 | |
| Zygotorulaspora_sp._yHDO592 | Zygotorulaspora_sp._yHDO592 | | Zygotorulaspora_sp._yHDO592 | *yHDO592_zygotorulaspora_sp_190924* | g004529.m1 | | | 0 | 1 | | MDM31-NVJ1-NMD5-UTP9 | |
| hagleromyces_aurorensis | hagleromyces_aurorensis | *#N/A* | *Hagleromyces_aurorensis* | *yHDO579_hagleromyces_aurorensis_180604* | *g002965.m1* | *#N/A* | *Zygotorulaspora* | 1 | 1 | | MDM31-NVJ1-NMD5-UTP9 | |
| lachanceakluyveri | *lachancea_kluyveri* | *snap_masked-SAKL0G-processed-gene-15.78* | *Lachancea_kluyveri* | *yHMPu5000034694_lachancea_kluyveri_180604* | *g001117.m1* | *#N/A* | *Lachancea* | 1 | 1 | | MDM31-NVJ1-NMD5-UTP9 | |
| lachancea_mirantina | lachancea_mirantina | *augustus_masked-LAMI0C-processed-gene-0.22* | *Lachancea_mirantina* | *yHMPu5000034691_lachancea_mirantina_180604* | *g001548.m1* | *SCU82809.1* | *Lachancea* | 1 | 1 | | MDM31-NVJ1-NMD5-? | |
| Lachancea_nothofagi | Lachancea_nothofagi | *snap_masked-LANO0B-processed-gene-1.74* | *Lachancea_nothofagi* | *yHMPu5000034690_lachancea_nothofagi_180604* | *g001406.m1* | *SCU80770.1* | *Lachancea* | 1 | 1 | | MDM31-NVJ1-NMD5-UTP9 | |
| lachanceafantastica | *lachancea_fantastica* | *#N/A* | Lachancea_fantastica_nom._nud. | *lachancea_fantastica* | *g003435.m1* | *SCU92434.1* | *Lachancea* | 1 | 1 | | MDM31-NVJ1-NMD5-UTP9 | |
| lachancealanzarotensis | *lachancea_lanzarotensis* | *augustus_masked-LN736361.1-processed-gene-3.12* | *Lachancea_lanzarotensis* | *yHMPu5000034693_Lachancea_lanzarotensis_SPADES* | *g004947.m1* | *XP_022627210.1* | *Lachancea* | 1 | 1 | | MDM31-NVJ1-NMD5-UTP9 | |
| lachanceawaltii | *lachancea_waltii* | *snap_masked-AADM01000119.1-processed-gene-0.11* | *Lachancea_waltii* | *yHMPu5000026219_lachancea_waltii_180604* | *g003199.m1* | *#N/A* | *Lachancea* | 1 | 1 | | MDM31-NVJ1-NMD5-UTP9 | |
| lachanceaquebecensis | *lachancea_quebecensis* | *snap_masked-LAQU0S22-processed-gene-0.116* | *Lachancea_quebecensis* | *yHMPu5000040963_lachancea_quebecuensis_200128* | *g002616.m1* | *#N/A* | *Lachancea* | 1 | 1 | | MDM31-NVJ1-NMD5-UTP9 | |
| lachanceathermotolerans | *lachancea_thermotolerans* | *snap_masked-CU928168-processed-gene-14.65* | *Lachancea_thermotolerans* | *yHMPu5000034678_lachancea_thermotolerans_180604* | *g003213.m1* | *XP_002553478.1* | *Lachancea* | 1 | 1 | | MDM31-NVJ1-NMD5-UTP9 | |
| lachanceameyersii | *lachancea_meyersii* | *snap_masked-LAME0G-processed-gene-17.24* | *Lachancea_meyersii* | *yHMPu5000034692_lachancea_meyersii_180604* | *g001214.m1* | *SCV01664.1* | *Lachancea* | 1 | 1 | | MDM31-NVJ1-NMD5-UTP9 | |
| lachanceadasiensis | *lachancea_dasiensis* | *augustus_masked-LADA0A-processed-gene-7.6* | *Lachancea_dasiensis* | *yHMPu5000034697_lachancea_dasiensis_190924* | *g003259.m1* | *SCU78828.1* | *Lachancea* | 1 | 1 | | MDM31-NVJ1-NMD5-UTP9 | |
| lachanceafermentati | *lachancea_fermentati* | *augustus_masked-LAFE0A-processed-gene-1.131* | *Lachancea_fermentati* | *yHMPu5000034695_lachancea_fermentati_201018* | *g004518.m1* | *SCV99344.1* | *Lachancea* | 1 | 1 | | MDM31-NVJ1-NMD5-UTP9 | |
| Lachancea_cidri_NRRL_Y-12635 | Lachancea_cidri_NRRL_Y-12635 | | Lachancea_cidri_NRRL_Y-12635 | *lookup* | | | | 0 | 1 | | MDM31-NVJ1-NMD5-UTP9 | |
| eremotheciugossypii | *eremotheciu_gossypii* | *#N/A* | Eremothecium_gossypii | *eremothecium_gossypii* | *NA* | *NP_982880.2* | *Eremothecium* | 1 | 0 | | MDM31-NMD5-UTP9 | NVJ1 not annotated in y1000 |
| eremotheciumcymbalariae | *eremothecium_cymbalariae* | *snap_masked-NC_016453.1-processed-gene-3.121* | *Eremothecium_cymbalariae* | *eremothecium_cymbalariae* | *g002832.m1* | *XP_003646754.1* | *Eremothecium* | 1 | 1 | | MDM31-NVJ1-NMD5-UTP9 | |
| eremotheciumcoryli | *eremothecium_coryli* | *maker-AZAH01000009.1-snap-gene-0.44* | *Eremothecium_coryli* | *eremothecium_coryli* | *g004362.m1* | *#N/A* | *Eremothecium* | 1 | 0 | | MDM31-NVJ1-NMD5-UTP9 | |
| eremotheciumsinecaudum | *eremothecium_sinecaudum* | *#N/A* | *Eremothecium_sinecaudum* | *eremothecium_sinecaudum* | *g003584.m1* | *XP_017989102.1* | *Eremothecium* | 1 | 1 | | MDM31-NVJ1-NMD5-? | |
| kluyveromyceslactis | *kluyveromyces_lactis* | *#N/A* | Kluyveromyces_lactis_CBS_2359 | *kluyveromyces_lactis* | *g000677.m1* | *XP_451721.1* | *Kluyveromyces* | 1 | 1 | | MDM31-NVJ1-NMD5-UTP9 | |
| Kluyveromyces_sp._yHMH660 | Kluyveromyces_sp._yHMH660 | | Kluyveromyces_sp._yHMH660 | *yHMH660_Kluyveromyces_sp_nov_plate33_SPADES* | g005129.m1 | | | 0 | 1 | | MDM31-NVJ1-NMD5-UTP9 | |
| kluyveromycesdobzhanskii | *kluyveromyces_dobzhanskii* | *snap_masked-NODE_13_length_251415_cov_20.6991_ID_25-processed-gene-1.144* | *Kluyveromyces_dobzhanskii* | *yHMPu5000034710_kluyveromyces_dobzhanskii_160519* | *g001256.m1* | *CDO92201.1* | *Kluyveromyces* | 1 | 1 | | MDM31-NVJ1-NMD5-UTP9 | |
| Kluyveromyces_aestuarii_NRRL_YB-4510 | Kluyveromyces_aestuarii_NRRL_YB-4510 | | Kluyveromyces_aestuarii_NRRL_YB-4510 | *yHMPu5000034709_kluyveromyces_aestuarii_160519* | g003160.m1 | | | 0 | 1 | | MDM31-NVJ1-NMD5-UTP9 | |
| kluyveromyces_nonfermentans | kluyveromyces_nonfermentans | *#N/A* | *Kluyveromyces_nonfermentans* | *yHMPu5000034701_kluyveromyces_nonfermentans_170307* | *g001343.m1* | *#N/A* | *Kluyveromyces* | 1 | 1 | | MDM31-NVJ1-NMD5-UTP9 | |
| kluyveromyces_siamensis | kluyveromyces_siamensis | *#N/A* | *Kluyveromyces_siamensis* | *yHMPu5000034700_kluyveromyces_siamensis_180604* | *g003460.m1* | *#N/A* | *Kluyveromyces* | 1 | 1 | | MDM31-NVJ1-NMD5-UTP9 | |
| kluyveromyces_starmeri | kluyveromyces_starmeri | *#N/A* | *Kluyveromyces_starmeri* | *yHDO570_kluyveromyces_starmeri_180604* | *g003161.m1* | *#N/A* | *Kluyveromyces* | 1 | 1 | | MDM31-NVJ1-NMD5-UTP9 | |
| kluyveromyces_wickerhamii | kluyveromyces_wickerhamii | *#N/A* | *Kluyveromyces_wickerhamii* | *yHMPu5000034699_kluyveromyces_wickerhamii_170307* | *g000532.m1* | *#N/A* | *Kluyveromyces* | 1 | 1 | | MDM31-NVJ1-NMD5-UTP9 | |
| hanseniasporauvarum | *hanseniaspora_uvarum* | *#N/A* | Hanseniaspora_uvarum_DSMZ_2768 | *hanseniaspora_uvarum* | *g000873.m1* | *KKA03541.1* | *Hanseniaspora* | 1 | 0 | | <-NVJ1-MDM31, UTP9 | inversion |
| hanseniasporapseudoguilliermondi | *hanseniaspora_pseudoguilliermondi* | *#N/A* | Hanseniaspora_pseudoguilliermondii | *yHMPu5000035695_hanseniaspora_pseudoguilliermondii_160519.haplomerger2* | *g000765.m1* | *#N/A* | *Hanseniaspora* | 1 | 0 | | <-NVJ1-MDM31, UTP9 | |
| hanseniasporaclermontiae | *hanseniaspora_clermontiae* | *#N/A* | *Hanseniaspora_clermontiae* | *yHMPu5000034963_hanseniaspora_clermontiae_160519.haplomerger2* | *g003387.m1* | *#N/A* | *Hanseniaspora* | 1 | 0 | | <-NVJ1-MDM31, UTP9 | |
| hanseniasporavalbyensis | *hanseniaspora_valbyensis* | *#N/A* | *Hanseniaspora_valbyensis* | *yHMPu5000034955_hanseniaspora_valbyensis_201018.haplomerger2* | g000788.m1 | *OBA26588.1* | *Hanseniaspora* | 1 | 0 | | <-NVJ1-MDM31, UTP9 | |
| hanseniasporavinae | *hanseniaspora_vinae* | *#N/A* | Hanseniaspora_vineae | *yHMPu5000026147_hanseniaspora_vineae_190924.haplomerger2* | *g000163.m1* | *#N/A* | *Hanseniaspora* | 1 | 0 | | ? | small contig |
| cyberlindnerasuaveolens | *cyberlindnera_suaveolens* | *#N/A* | *Cyberlindnera_suaveolens* | *yHMPu5000035687_cyberlindnera_suaveolens_160613* | *g005674.m1* | *#N/A* | *Cyberlindnera* | 1 | 0 | | MDM31-NVJ1-?-NMD5-UTP9 | |
| cyberlindnerasaturnus | *cyberlindnera_saturnus* | *#N/A* | *Cyberlindnera_saturnus* | *yHMPu5000035686_Cyberlindnera_saturnus_SPADES* | *g001009.m1* | *#N/A* | *Cyberlindnera* | 1 | 0 | | MDM31-NVJ1-?-NMD5-UTP9 | |
| cyberlindneramisumaiensis | *cyberlindnera_misumaiensis* | *#N/A* | *Cyberlindnera_misumaiensis* | *yHMPu5000034979_cyberlindnera_misumaiensis_160519* | *g002417.m1* | *#N/A* | *Cyberlindnera* | 1 | 0 | | MDM31-NVJ1-?-NMD5-? | |
| Cyberlindnerajadinii | *Cyberlindnera_jadinii* | *#N/A* | *Cyberlindnera_jadinii* | *yHMPu5000035336_cyberlindnera_jadinii_180604.haplomerger2* | *g002673.m1* | *XP_020072378.1* | *Cyberlindnera* | 1 | 0 | | ? | contig break |
| Cyberlindnerafabianii | *Cyberlindnera_fabianii* | *#N/A* | *Cyberlindnera_fabianii* | *yHMPu5000035701_cyberlindnera_fabianii_160519* | *g005778.m1* | *ONH68477.1* | *Cyberlindnera* | 1 | 0 | | ? | MDM31 nearby |
| cyberlindneramaclurae | *cyberlindnera_maclurae* | *#N/A* | *Cyberlindnera_maclurae* | *yHMPu5000035699_cyberlindnera_maclurae_160613* | *g003717.m1* | *#N/A* | *Cyberlindnera* | 1 | 1 | | MDM31-NVJ1-?-NMD5-UTP9 | |
| wickerhamomycesalni | *wickerhamomyce_salni* | *#N/A* | Wickerhamomyces_alni | *yHMPu5000035274_wickerhamomyces_alni_170307* | *g001244.m1* | *#N/A* | *Wickerhamomyces* | 1 | 0 | | MDM31-NVJ1-?-NMD5-? | |
| wickerhamomycescanadensis | *wickerhamomyces_canadensis* | *#N/A* | *Wickerhamomyces_canadensis* | *yHMPu5000035639_wickerhamomyces_canadensis_160613* | *g004559.m1* | *#N/A* | *Wickerhamomyces* | 1 | 0 | | MDM31-NVJ1-?-NMD5-UTP9 | |
| wickerhamomycesciferrii | *wickerhamomyces_ciferrii* | *#N/A* | *Wickerhamomyces_ciferrii* | *yHMPu5000035269_wickerhamomyces_ciferrii_170307* | g005482.m1,g003054.m1 | *XP_011277532.1* | *Wickerhamomyces* | 1 | 0 | | ? | |
| Wickerhamomyceschambardii | *Wickerhamomyces_chambardii* | *#N/A* | *Wickerhamomyces_chambardii* | *yHMPu5000035270_wickerhamomyces_chambardii_180604.haplomerger2* | *g001027.m1* | *#N/A* | *Wickerhamomyces* | 1 | 0 | | <-UTP9, MDM31,<-MVJ1-NMD5 | |
| Wickerhamomyceshampshirensis | *Wickerhamomyces_hampshirensis* | *#N/A* | *Wickerhamomyces_hampshirensis* | *yHMPu5000035268_wickerhamomyces_hampshirensis_160928* | *g002138.m1* | *#N/A* | *Wickerhamomyces* | 1 | 0 | | MDM31-NVJ1-?-NMD5-? | |
| Wickerhamomycesmucosus | *Wickerhamomyces_mucosus* | *#N/A* | *Wickerhamomyces_mucosus* | *yHMPu5000038048_wickerhamomyces_mucosus_170912.haplomerger2* | *g002404.m1* | *#N/A* | *Wickerhamomyces* | 1 | 0 | | ? | |
| Wickerhamomycespiperi | *Wickerhamomyces_piperi* | *#N/A* | Wickerhamomyces_pijperi | *yHMPu5000035254_wickerhamomyces_pijperi_160928.haplomerger2* | g002584.m1 | *#N/A* | *Wickerhamomyces* | 1 | 0 | | <-NVJ1, NMD5-MDM31,?-?-?-UTP9 | |
| komagataellaphaffi | *komagataella_phaffi* | *#N/A* | Komagataella_phaffii_GS115 | *komagataella_phaffi* | *g002262.m1* | *XP_002491603.1* | *Komagataella* | 1 | 0 | | MDM31-?,<-NVJ1, NMD5 | |
| komagataellapopuli | *komagataella_populi* | *#N/A* | *Komagataella_populi* | *yHMPu5000026274_komagataella_populi_160519* | *g004657.m1* | *#N/A* | *Komagataella* | 1 | 0 | | MDM31-?,<-NVJ1, NMD5 | |
| komagataellapastoris | *komagataella_pastoris* | *#N/A* | Komagataella_pastoris_ATCC_28485 | *komagataella_pastoris* | *g002679.m1* | *ANZ75330.1* | *Komagataella* | 1 | 0 | | MDM31-?,<-NVJ1, NMD5 | |

| Species Name | seqID | HMMProfileID | Y1000_TreeID | Y1000_DatasetID | Y1000_protID | ncbiID | group | inTableS1 | inY1000Hmmer | CopyNumber | syntenyCheck | syntenyComments |
|---|---|---|---|---|---|---|---|---|---|---|---|---|
| Komagataellakurtzmanii | *Komagataella_kurtzmanii* | *#N/A* | *Komagataella_kurtzmanii* | *yHMPu5000035676_komagataella_kurtzmanii_160613* | *g004231.m1* | *#N/A* | *Komagataella* | 1 | 0 | | MDM31-?,<-NVJ1, NMD5 | |
| Komatagataella | Komatagaella_mondaviorum | *#N/A* | Komagataella_mondaviorum | *yHMPu5000037220_komagataella_mondaviorum_210210* | *g004487.m1* | *#N/A* | *Komagataella* | 1 | 0 | | MDM31-?,<-NVJ1, NMD5 | |
| Kuraishia | Kuraishia_floccosa | *#N/A* | *Kuraishia_floccosa* | *yHMPu5000034935_kuraishia_floccosa_210210* | *g004700.m1* | *#N/A* | Kuraishia | 1 | 0 | | MDM31-?,<-NVJ1, NMD5 | |
| NA | Asbhya_aceri | *#N/A* | Ashbya_aceri | ashbya_aceri | NA | AGO10388.1 | Eremothecium | 1 | 0 | | MDM31-NMD5-UTP9 | No NVJ1 annotation in Y1000 data |
| Wickerhamomyces_anomalus | Wickerhamomyces_anomalus | *#N/A* | Wickerhamomyces_anomalus | yHMPu5000035273_wickerhamomyces_anomalus_180604.haplomerger2 | g001657.m1 | TBD | Wickerhamomyces | 1 | 0 | | MDM31-?,<-NVJ1-?, NMD5 | |

| Species Name | seqID | HMMProfileID | Y1000_TreeID | Y1000_DatasetID | Y1000_protID | ncbiID | group | , | Molecular Weight (KDa) | , | Isoelectric Point | | | Molecular Weight (Da) |
|---|---|---|---|---|---|---|---|---|---|---|---|---|---|---|
| saccharomycescerevisiae | saccharomyces_cerevisiae | YHR195W | Saccharomyces_cerevisiae | saccharomyces_cerevisiae | g005344.m1 | NP_012065.3 | Saccharomyces | , | 36.42145 | , | 4.7988 | | | 36421.45 |
| saccharomycesjurei | saccharomyces_jurei | #N/A | Saccharomyces_jurei | saccharomyces_jurei | g004292.m1 | #N/A | Saccharomyces | , | 36.85304 | , | 5.4497 | | | 36853.04 |
| saccharomycesparadoxus | saccharomyces_paradoxus | genemark-Spar_8-processed-gene-4.108 | Saccharomyces_paradoxus | saccharomyces_paradoxus | g005237.m1 | #N/A | Saccharomyces | , | 36.47436 | , | 4.5513 | | | 36474.36 |
| saccharomyceskudriavzevii | saccharomyces_kudriavzevii | augustus_masked-Skud_8-processed-gene-3.158 | Saccharomyces_kudriavzevii | saccharomyces_kudriavzevii | g005184.m1 | EJT43435.1 | Saccharomyces | , | 36.15633 | , | 4.7852 | | | 36156.33 |
| saccharomycesarboricola | saccharomyces_arboricola | augustus_masked-NC_026178.1-processed-gene-4.111 | Saccharomyces_arboricola | saccharomyces_arboricola | g005224.m1 | EJS43367.1 | Saccharomyces | , | 36.45834 | , | 4.6479 | | | 36458.34 |
| saccharomycesmikatae | saccharomyces_mikatae | snap_masked-Smik_8-processed-gene-4.133 | Saccharomyces_mikatae | saccharomyces_mikatae | g005189.m1 | #N/A | Saccharomyces | , | 36.92298 | , | 4.6452 | | | 36922.98 |
| saccharomycesuvarum | saccharomyces_uvarum | augustus_masked-Sbay_15-processed-gene-6.183 | Saccharomyces_uvarum | saccharomyces_uvarum | g002217.m1 | #N/A | Saccharomyces | , | 36.68756 | , | 4.5178 | | | 36687.56 |
| saccharomyceseubayanus | saccharomyces_eubayanus | augustus_masked-chrXV-processed-gene-6.90 | Saccharomyces_eubayanus | saccharomyces_eubayanus | g005249.m1 | XP_018221971.1 | Saccharomyces | , | 36.37819 | , | 4.4885 | | | 36378.19 |
| Nakasesomycesglabrata | Nakasesomyces_glabrata | #N/A | Nakaseomyces_glabratus | yHMPu5000034723_candida_glabrata_180604 | g000308.m1 | XP_449661.1 | Nakasesomyces | , | 27.2777 | , | 4.748 | | | 27277.7 |
| Nakaseomyceskungkrabaensis | Nakaseomyces_kungkrabaensis | #N/A | Nakaseomyces_kungkrabaensis | yHMPu5000034722_candida_kungkrabaensis_180604 | g004198.m1 | #N/A | Nakasesomyces | , | 26.47484 | , | 4.4612 | | | 26474.84 |
| Nakaseomycesnivariensis | Nakaseomyces_nivariensis | snap_masked-CANI0S17-processed-gene-4.132 | Nakaseomyces_nivariensis | yHMPu5000034720_candida_nivariensis_180604 | g001471.m1 | #N/A | Nakasesomyces | , | 26.54481 | , | 4.4612 | | | 26544.81 |
| Nakaseomycesuthaithanina | Nakaseomyces_uthaithanina | #N/A | Nakaseomyces_uthaithaninus | yHMPu5000034718_candida_uthaithanina_180604 | g005380.m1 | #N/A | Nakasesomyces | , | 25.03534 | , | 5.7115 | | | 25035.34 |
| nakaseomycesdelphensis | nakaseomyces_delphensis | augustus_masked-NADE0S27-processed-gene-3.39 | Nakaseomyces_delphensis | yHMPu5000026127_nakaseomyces_delphensis_160519 | g004209.m1 | CAO98811.1 | Nakasesomyces | , | 26.76638 | , | 5.0106 | | | 26766.38 |
| nakaseomycesbracarensis | nakaseomyces_bracarensis | snap_masked-CABR0S29-processed-gene-6.0 | Nakaseomyces_bracarensis | yHMPu5000034725_candida_bracarensis_160928 | g001614.m1 | #N/A | Nakasesomyces | , | 30.75206 | , | 3.9239 | | | 30752.06 |
| nakaseomycesbacillisporus | nakaseomyces_bacillisporus | augustus_masked-NABA0S29-processed-gene-0.108 | Nakaseomyces_bacillisporus | yHMPu5000035666_nakaseomyces_bacillisporus_160613 | g004314.m1 | #N/A | Nakasesomyces | , | 30.26465 | , | 5.5641 | | | 30264.65 |
| nakaseomycescastellii_ | nakaseomyces_castellii_ | snap_masked-CACA0S33-processed-gene-9.107 | Nakaseomyces_castellii | yHMPu5000034724_candida_castellii_180604 | g001576.m1 | #N/A | Nakasesomyces | , | 43.0299 | , | 4.8634 | | | 43029.9 |
| kazachstaniasolicola | kazachstania_solicola | augustus_masked-scf7180000046332-processed-gene-1.206 | Kazachstania_solicola | yHAB159_kazachstania_solicola_160519.haplomerger2 | g005068.m1 | #N/A | Kazachstania | , | 39.01031 | , | 3.9829 | | | 39010.31 |
| kazachstaniaaerobia | kazachstania_aerobia | augustus_masked-scf7180000020169-processed-gene-0.188 | Kazachstania_aerobia | yHAB164_kazachstania_aerobia_160519 | g002718.m1 | #N/A | Kazachstania | , | 40.26682 | , | 4.1348 | | | 40266.82 |
| kazachstaniaunispora | kazachstania_unispora | snap_masked-NODE_47_length_78712_cov_50.1529_ID_93-processed-gene-0.49 | Kazachstania_unispora | yHAB133_kazachstania_unispora_160519 | g003649.m1 | #N/A | Kazachstania | , | 38.83565 | , | 4.1879 | | | 38835.65 |
| kazachstania_bromeliacearum | kazachstania_bromeliacearum | snap_masked-NODE_10_length_278180_cov_22.5073_ID_19-processed-gene-1.127 | Kazachstania_bromeliacearum | yHAB136_kazachstania_bromeliacearum_160519 | g000451.m1 | #N/A | Kazachstania | , | 26.35025 | , | 4.4822 | | | 26350.25 |
| kazachstania_bulderi | kazachstania_bulderi | #N/A | Kazachstania_bulderi | yHAB157_kazachstania_bulderi_160519.haplomerger2 | g005958.m1 | #N/A | Kazachstania | , | 49.04927 | , | 4.3063 | | | 49049.27 |
| kazachstania_exigua | kazachstania_exigua | #N/A | Kazachstania_exigua | yHAB150_kazachstania_exigua_160519 | g002519.m1 | #N/A | Kazachstania | , | 49.09724 | , | 4.2164 | | | 49097.24 |
| kazachstania_gamospora | kazachstania_gamospora | #N/A | Kazachstania_gamospora | yHAB142_kazachstania_gamospora_170307.haplomerger2 | g004167.m1 | #N/A | Kazachstania | , | 41.89352 | , | 4.6891 | | | 41893.52 |
| kazachstania_hellenica | kazachstania_hellenica | #N/A | Kazachstania_hellenica | yHAB145_kazachstania_hellenica_160519.haplomerger2 | g003117.m1 | #N/A | Kazachstania | , | 47.51685 | , | 4.232 | | | 47516.85 |
| kazachstania_aquatica | Missing! | #N/A | Kazachstania_aquatica | yHAB165_kazachstania_aquatica_160519 | g003032.m1 | #N/A | | | | | | | | |
| kazachstania_kunashirensis | kazachstania_kunashirensis | snap_masked-scf7180000008985-processed-gene-1.61 | Kazachstania_kunashirensis | yHAB160_kazachstania_kunashirensis_160519 | g005050.m1 | #N/A | Kazachstania | , | 34.65823 | , | 4.3628 | | | 34658.23 |
| kazachstania_lodderae | kazachstania_lodderae | #N/A | Kazachstania_lodderae | yHAB138_kazachstania_lodderae_160519 | g003605.m1 | #N/A | Kazachstania | , | 48.90606 | , | 4.9175 | | | 48906.06 |
| kazachstania_piceae | kazachstania_piceae | #N/A | Kazachstania_piceae | yHAB156_kazachstania_piceae_160519 | g005225.m1 | #N/A | Kazachstania | , | 52.16235 | , | 4.8196 | | | 52162.35 |
| kazachstania_saulgeensis | kazachstania_saulgeensis | #N/A | Kazachstania_saulgeensis | yHMPu5000037225_kazachstania_saulgeensis_210210 | g000550.m1 | SMN19139.1 | Kazachstania | , | 44.35671 | , | 4.0589 | | | 44356.71 |
| kazachstania_servazzi | kazachstania_servazzi | #N/A | Kazachstania_servazzii | yHAB151_kazachstania_servazzi_160519 | g000773.m1 | #N/A | Kazachstania | , | 36.19702 | , | 4.0021 | | | 36197.02 |
| kazachstania_sinensis | kazachstania_sinensis | #N/A | Kazachstania_sinensis | yHAB161_kazachstania_sinensis_160519 | g005599.m1 | #N/A | Kazachstania | , | 32.84921 | , | 4.2566 | | | 32849.21 |
| kazachstania_yasuniensis | kazachstania_yasuniensis | #N/A | Kazachstania_yasuniensis | yHMPu5000034708_kazachstania_yasuniensis_180604 | g001209.m1 | #N/A | Kazachstania | , | 39.26319 | , | 4.0804 | | | 39263.19 |
| Kazachstania_barnettii | Kazachstania_barnettii | #N/A | Kazachstania_barnettii | yHAB162_Kazachstania_barnettii_SPADES | g001263.m1 | #N/A | Kazachstania | , | 43.36564 | , | 4.0478 | | | 43365.64 |
| Kazachstania_psychrophila | Kazachstania_psychrophila | #N/A | Kazachstania_psychrophila | yHMPu5000034706_Kazachstania_psychrophila_SPADES | g002942.m1 | #N/A | Kazachstania | , | 41.18863 | , | 4.0893 | | | 41188.63 |
| Kazachstania_jinghongensis | Kazachstania_jinghongensis | #N/A | Kazachstania_jinghongensis | yHMPu5000037210_Kazachstania_jinghongensis_SPADES | g001801.m1 | #N/A | Kazachstania | , | 48.51643 | , | 5.2415 | | | 48516.43 |
| kazachstaniasiamensis | kazachstania_siamensis | augustus_masked-NODE_61_length_63804_cov_14.6622_ID_121-processed-gene-0.80 | Kazachstania_siamensis | yHAB143_kazachstania_siamensis_160519 | g004344.m1 | #N/A | Kazachstania | , | 32.30443 | , | 4.0753 | | | 32304.43 |
| kazachstaniataianensis | kazachstania_taianensis | augustus_masked-NODE_5_length_452406_cov_49.1731_ID_9-processed-gene-3.120 | Kazachstania_taianensis | yHAB147_kazachstania_taianensis_160519 | g004007.m1 | #N/A | Kazachstania | , | 26.77711 | , | 4.3032 | | | 26777.11 |
| kazachstanianaganishii | kazachstania_naganishii | snap_masked-HE978326-processed-gene-0.118 | Kazachstania_naganishii | kazachstania_naganishii | g001440.m1 | XP_022467125.1 | Kazachstania | , | 31.97654 | , | 4.4904 | | | 31976.54 |
| kazachstaniamartiniae | kazachstania_martiniae | snap_masked-scf7180000030514-processed-gene-0.103 | Kazachstania_martiniae | yHAB132_kazachstania_martiniae_160519 | g004439.m1 | #N/A | Kazachstania | , | 30.1896 | , | 4.3839 | | | 30189.6 |
| kazachstaniaturicensis | kazachstania_turicensis | snap_masked-scf7180000049581-processed-gene-4.63 | Kazachstania_turicensis | yHMPu5000040961_kazachstania_turicensis_201018 | g004713.m1 | #N/A | Kazachstania | , | 48.82722 | , | 4.3272 | | | 48827.22 |
| kazachstaniakunashirensis | kazachstania_kunashirensis | snap_masked-scf7180000008985-processed-gene-1.61 | Kazachstania_kunashirensis | yHAB160_kazachstania_kunashirensis_160519 | g005050.m1 | #N/A | Kazachstania | , | 43.41379 | , | 4.328 | | | 43413.79 |
| kazachstaniaspencerorum | kazachstania_spencerorum | snap_masked-scf7180000055600-processed-gene-0.48 | Kazachstania_spencerorum | yHAB155_kazachstania_spencerorum_160519 | g002874.m1 | #N/A | Kazachstania | , | 46.07466 | , | 4.6212 | | | 46074.66 |
| kazachstaniaafricana | kazachstania_africana | augustus_masked-scaffold_2-processed-gene-13.205 | Kazachstania_africana | yHAB137_kazachstania_africana_160519 | g003755.m1 | XP_003956120.1 | Kazachstania | , | 40.02251 | , | 9.5003 | | | 40022.51 |
| kazachstaniaviticola | kazachstania_viticola | snap_masked-scf7180000050361-processed-gene-0.67 | Kazachstania_viticola | yHAB158_kazachstania_viticola_160519.haplomerger2 | g001183.m1 | #N/A | Kazachstania | , | 40.5042 | , | 4.8594 | | | 40504.2 |
| naumovozymadairenensis | naumovozyma_dairenensis | augustus_masked-NC_016482.1-processed-gene-7.22 | Naumovozyma_dairenensis | yHMPu5000034872_naumovozyma_dairenensis_180604 | g002752.m1 | XP_003669889.1 | Naumovozyma | , | 50.83124 | , | 4.1573 | | | 50831.24 |
| naumovozymacastellii | naumovozyma_castellii | snap_masked-HE576752-processed-gene-12.71 | Naumovozyma_castellii | yHMPu5000034871_naumovozyma_castellii_180604 | g001098.m1 | #N/A | Naumovozyma | , | 40.86545 | , | 4.3861 | | | 40865.45 |
| naumovozyma_baii | naumovozyma_baii | #N/A | Naumovozyma_baii | yHMPu5000034894_naumovozyma_baii_190924 | g000794.m1 | #N/A | Naumovozyma | , | 37.34486 | , | 5.8373 | | | 37344.86 |
| tetrapisisporairiomotensis | tetrapisispora_iriomotensis | augustus_masked-flattened_line_57-processed-gene-0.130 | Tetrapisispora_iriomotensis | yHMPu5000034876_Tetrapisispora_iriomotensis_SPADES | g003416.m1 | #N/A | Tetrapisispora | , | 41.90352 | , | 4.3115 | | | 41903.52 |
| tetrapisisporanamnaonensis | tetrapisispora_namnaonensis | #N/A | Tetrapisispora_namnaoensis | yHMPu5000034877_tetrapisispora_namnaonensis_160519 | g001993.m1 | #N/A | Tetrapisispora | , | 35.97886 | , | 4.1963 | | | 35978.86 |
| tetrapisisporablattae | tetrapisispora_blattae | snap_masked-NC_020185-processed-gene-25.62 | Tetrapisispora_blattae | yHMPu5000034874_tetrapisispora_blattae_190924 | g003018.m1 | XP_004178357.1 | Tetrapisispora | , | 52.65595 | , | 4.1007 | | | 52655.95 |
| vanderwaltozymapolyspora | vanderwaltozyma_polyspora | snap_masked-NW_001834637.1-processed-gene-0.82 | Vanderwaltozyma_polyspora | yHMPu5000034869_vanderwaltozyma_polyspora_180604 | g005365.m1 | XP_001643905.1 | Vanderwaltozyma | , | 46.01007 | , | 4.2926 | | | 46010.07 |
| Vanderwaltozyma_yarrowii | Vanderwaltozyma_yarrowii | #N/A | Vanderwaltozyma_yarrowii | yHMPu5000034868_Vanderwaltozyma_yarrowii_S33_SPADES | g001449.m1 | #N/A | Vanderwaltozyma | , | 44.66004 | , | 3.9007 | | | 44660.04 |
| vanderwaltozyma_tropicalis | vanderwaltozyma_tropicalis | #N/A | Vanderwaltozyma_tropicalis | yHMPu5000026257_vanderwaltozyma_tropicalis_190924 | g003717.m1 | #N/A | Vanderwaltozyma | , | 34.86475 | , | 4.6183 | | | 34864.75 |
| Vanderwaltozyma_verrucispora | Vanderwaltozyma_verrucispora | #N/A | Vanderwaltozyma_verrucispora | yHMPu5000037837_Vanderwaltozyma_verrucispora_S176_SPADES | g005143.m1 | #N/A | Vanderwaltozyma | , | 41.38653 | , | 4.6518 | | | 41386.53 |
| torulasporapretoriensis | torulaspora_pretoriensis | augustus_masked-flattened_line_3-processed-gene-4.12 | Torulaspora_pretoriensis | yHMPu5000034881_Torulaspora_pretoriensis_SPADES | g004938.m1 | #N/A | Torulaspora | , | 35.57586 | , | 4.9184 | | | 35575.86 |
| torulasporamicroellipsoides | torulaspora_microellipsoides | augustus_masked-flattened_line_52-processed-gene-0.132 | Torulaspora_microellipsoides | yHMPu5000035651_Torulaspora_microellipsoides_SPADES | g002639.m1 | #N/A | Torulaspora | , | 38.43885 | , | 4.9983 | | | 38438.85 |
| torulasporafranciscae | torulaspora_franciscae | snap_masked-flattened_line_10-processed-gene-1.40 | Torulaspora_franciscae | yHMPu5000026152_Torulaspora_franciscae_SPADES | g004752.m1 | #N/A | Torulaspora | , | 35.29763 | , | 4.7552 | | | 35297.63 |
| torulasporadelbrueckii | torulaspora_delbrueckii | snap_masked-scaffold_4-processed-gene-1.34 | Torulaspora_delbrueckii | yHMPu5000035653_torulaspora_delbrueckii_160613 | g001776.m1 | XP_003680850.1 | Torulaspora | , | 35.87571 | , | 5.2494 | | | 35875.71 |
| torulaspora_globosa | torulaspora_globosa | #N/A | Torulaspora_globosa | yHMPu5000034880_torulaspora_globosa_160519 | g001826.m1 | #N/A | Torulaspora | , | 32.85382 | , | 4.2401 | | | 32853.82 |
| torulaspora_maleeae | torulaspora_maleeae | snap_masked-NODE_5_length_764704_cov_59.617_ID_1253-processed-gene-6.109 | Torulaspora_maleeae | yHMPu5000035652_torulaspora_maleeae_160613 | g003867.m1 | #N/A | Torulaspora | , | 33.46367 | , | 4.5261 | | | 33463.67 |
| zygosaccharomyces_bisporus | zygosaccharomyces_bisporus | snap_masked-flattened_line_16-processed-gene-0.27 | Zygosaccharomyces_bisporus | yHMPu5000034866_zygosaccharomyces_bisporus_160519 | g005101.m1 | #N/A | Zygosaccharomyces | , | 30.78561 | , | 4.4988 | | | 30785.61 |
| zygosaccharomyces_gambellarensis | zygosaccharomyces_gambellarensis | #N/A | Zygosaccharomyces_gambellarensis | yHDO565_zygosaccharomyces_gambellarensis_180604 | g005022.m1 | #N/A | Zygosaccharomyces | , | 33.46945 | , | 4.6711 | | | 33469.45 |
| zygosaccharomycesbailii | Zygosaccharomyces_bailii | augustus_masked-ZYBA0S10-processed-gene-3.53 | Zygosaccharomyces_bailii | zygosaccharomyces_bailii | g001048.m1 | CDF91308.1 | Zygosaccharomyces | , | 31.26516 | , | 4.8738 | | | 31265.16 |
| zygosaccharomyceskombuchaensis | Zygosaccharomyces_kombuchaensis | snap_masked-NODE_1_length_271579_cov_10.1807_ID_1-processed-gene-1.23 | Zygosaccharomyces_kombuchaensis | yHMPu5000034865_zygosaccharomyces_kombuchaensis_160519 | g000063.m1 | #N/A | Zygosaccharomyces | , | 30.40245 | , | 4.6402 | | | 30402.45 |
| zygosaccharomyces_lentus | zygosaccharomyces_lentus | #N/A | Zygosaccharomyces_lentus | yHMPu5000034864_zygosaccharomyces_lentus_170307 | g004388.m1 | #N/A | Zygosaccharomyces | , | 30.25931 | , | 5.4041 | | | 30259.31 |
| zygosaccharomyces_mellis | zygosaccharomyces_mellis | #N/A | Zygosaccharomyces_mellis | yHDO572_zygosaccharomyces_mellis_180604 | g002135.m1 | #N/A | Zygosaccharomyces | , | 32.94388 | , | 5.2563 | | | 32943.88 |
| zygosaccharomyces_pseudorouxii | zygosaccharomyces_pseudorouxii | #N/A | Zygosaccharomyces_pseudorouxii | yHMPu5000037836_zygosaccharomyces_pseudorouxii_201018.haplomerger2 | g004410.m1 | #N/A | Zygosaccharomyces | , | 32.81375 | , | 4.7807 | | | 32813.75 |
| zygosaccharomyces_siamensis | zygosaccharomyces_siamensis | #N/A | Zygosaccharomyces_siamensis | yHMPu5000035626_zygosaccharomyces_siamensis_190924 | g000372.m1 | #N/A | Zygosaccharomyces | , | 32.81682 | , | 5.4515 | | | 32816.82 |

| Species Name | seqID | HMMProfileID | Y1000_TreeID | Y1000_DatasetID | Y1000_protID | ncbiID | group | , | Molecular Weight (KDa) | , | Isoelectric Point | | | Molecular Weight (Da) |
|---|---|---|---|---|---|---|---|---|---|---|---|---|---|---|
| zygotorulasporaflorentina | zygotorulaspora_florentina | augustus_masked-NODE_6_length_562868_cov_24.3835_ID_11-processed-gene-2.47 | Zygotorulaspora_florentina | yHMPu5000034862_zygotorulaspora_florentina_160519 | g004327.m1 | #N/A | Zygotorulaspora | , | 37.61049 | , | 6.2991 | | | 37610.49 |
| zygotorulasporamrakii | zygotorulaspora_mrakii | snap_masked-NODE_2_length_708340_cov_18.0358_ID_2767-processed-gene-4.105 | Zygotorulaspora_mrakii | yHMPu5000026256_zygotorulaspora_mrakii_170307 | g002003.m1 | #N/A | Zygotorulaspora | , | 39.02446 | , | 5.0952 | | | 39024.46 |
| zygotorulaspora_chibaensis | zygotorulaspora_chibaensis | #N/A | Zygotorulaspora_chibaensis | yHMPu5000037204_zygotorulaspora_chibaensis_210210 | g002973.m1 | #N/A | Zygotorulaspora | , | 37.70335 | , | 4.9361 | | | 37703.35 |
| zygotorulaspora_danielsina | zygotorulaspora_danielsina | #N/A | Zygotorulaspora_danielsina | yHMPu5000037205_zygotorulaspora_danielsina_210210 | g002121.m1 | #N/A | Zygotorulaspora | , | 36.47326 | , | 5.7074 | | | 36473.26 |
| zygotorulaspora_mrakii | zygotorulaspora_mrakii | snap_masked-NODE_2_length_708340_cov_18.0358_ID_2767-processed-gene-4.105 | Zygotorulaspora_mrakii | yHMPu5000026256_zygotorulaspora_mrakii_170307 | g002003.m1 | #N/A | Zygotorulaspora | , | 39.02446 | , | 5.0952 | | | 39024.46 |
| hagleromyces_aurorensis | hagleromyces_aurorensis | #N/A | Hagleromyces_aurorensis | yHDO579_hagleromyces_aurorensis_180604 | g002965.m1 | #N/A | Zygotorulaspora | , | 53.61132 | , | 4.3924 | | | 53611.32 |
| lachanceakluyveri | lachancea_kluyveri | snap_masked-SAKL0G-processed-gene-15.78 | Lachancea_kluyveri | yHMPu5000034694_lachancea_kluyveri_180604 | g001117.m1 | #N/A | Lachancea | , | 34.00936 | , | 5.1086 | | | 34009.36 |
| lachancea_mirantina | lachancea_mirantina | augustus_masked-LAMI0C-processed-gene-0.22 | Lachancea_mirantina | yHMPu5000034691_lachancea_mirantina_180604 | g001548.m1 | SCU82809.1 | Lachancea | , | 33.72982 | , | 5.6031 | | | 33729.82 |
| Lachancea_nothofagi | Lachancea_nothofagi | snap_masked-LANO0B-processed-gene-1.74 | Lachancea_nothofagi | yHMPu5000034690_lachancea_nothofagi_180604 | g001406.m1 | SCU80770.1 | Lachancea | , | 38.10047 | , | 8.1531 | | | 38100.47 |
| lachanceafantastica | lachancea_fantastica | #N/A | Lachancea_fantastica_nom._nud. | lachancea_fantastica | g003435.m1 | SCU92434.1 | Lachancea | , | 37.40871 | , | 6.7695 | | | 37408.71 |
| lachancealanzarotensis | lachancea_lanzarotensis | augustus_masked-LN736361.1-processed-gene-3.12 | Lachancea_lanzarotensis | yHMPu5000034693_Lachancea_lanzarotensis_SPADES | g004947.m1 | XP_022627210.1 | Lachancea | , | 37.58671 | , | 6.775 | | | 37586.71 |
| lachanceawaltii | lachancea_waltii | snap_masked-AADM01000119.1-processed-gene-0.11 | Lachancea_waltii | yHMPu5000026219_lachancea_waltii_180604 | g003199.m1 | #N/A | Lachancea | , | 33.30709 | , | 9.1367 | | | 33307.09 |
| lachanceaquebecensis | lachancea_quebecensis | snap_masked-LAQU0S22-processed-gene-0.116 | Lachancea_quebecensis | yHMPu5000040963_lachancea_quebecuensis_200128 | g002616.m1 | #N/A | Lachancea | , | 36.10041 | , | 8.3847 | | | 36100.41 |
| lachanceathermotolerans | lachancea_thermotolerans | snap_masked-CU928168-processed-gene-14.65 | Lachancea_thermotolerans | yHMPu5000034678_lachancea_thermotolerans_180604 | g003213.m1 | XP_002553478.1 | Lachancea | , | 32.42788 | , | 8.1613 | | | 32427.88 |
| lachanceameyersii | lachancea_meyersii | snap_masked-LAME0G-processed-gene-17.24 | Lachancea_meyersii | yHMPu5000034692_lachancea_meyersii_180604 | g001214.m1 | SCV01664.1 | Lachancea | , | 38.05227 | , | 9.1029 | | | 38052.27 |
| lachanceadasiensis | lachancea_dasiensis | augustus_masked-LADA0A-processed-gene-7.6 | Lachancea_dasiensis | yHMPu5000034697_lachancea_dasiensis_190924 | g003259.m1 | SCU78828.1 | Lachancea | , | 40.14557 | , | 7.4365 | | | 40145.57 |
| lachanceafermentati | lachancea_fermentati | augustus_masked-LAFE0A-processed-gene-1.131 | Lachancea_fermentati | yHMPu5000034695_lachancea_fermentati_201018 | g004518.m1 | SCV99344.1 | Lachancea | , | 31.72441 | , | 5.0251 | | | 31724.41 |
| eremotheciugossypii | eremotheciu_gossypii | #N/A | Eremothecium_gossypii | eremothecium_gossypii | NA | NP_982880.2 | Eremothecium | , | 25.22526 | , | 10.297 | | | 25225.26 |
| eremotheciumcymbalariae | eremothecium_cymbalariae | snap_masked-NC_016453.1-processed-gene-3.121 | Eremothecium_cymbalariae | eremothecium_cymbalariae | g002832.m1 | XP_003646754.1 | Eremothecium | , | 32.10557 | , | 4.7546 | | | 32105.57 |
| eremotheciumcoryli | eremothecium_coryli | maker-AZAH01000009.1-snap-gene-0.44 | Eremothecium_coryli | eremothecium_coryli | g004362.m1 | #N/A | Eremothecium | , | 27.77677 | , | 8.5921 | | | 27776.77 |
| eremotheciumsinecaudum | eremothecium_sinecaudum | #N/A | Eremothecium_sinecaudum | eremothecium_sinecaudum | g003584.m1 | XP_017989102.1 | Eremothecium | , | 32.91456 | , | 9.7132 | | | 32914.56 |
| kluyveromyceslactis | kluyveromyces_lactis | #N/A | Kluyveromyces_lactis_CBS_2359 | kluyveromyces_lactis | g000677.m1 | XP_451721.1 | Kluyveromyces | , | 31.42798 | , | 5.2075 | | | 31427.98 |
| kluyveromycesdobzhanskii | kluyveromyces_dobzhanskii | snap_masked-NODE_13_length_251415_cov_20.6991_ID_25-processed-gene-1.144 | Kluyveromyces_dobzhanskii | yHMPu5000034710_kluyveromyces_dobzhanskii_160519 | g001256.m1 | CDO92201.1 | Kluyveromyces | , | 32.31271 | , | 7.0271 | | | 32312.71 |
| kluyveromycesaestuarii | kluyveromyces_aestuarii | #N/A | Kluyveromyces_aestuarii_ATCC_18862 | kluyveromyces_aestuarii | g000823.m1 | #N/A | Kluyveromyces | , | 30.65803 | , | 7.3002 | | | 30658.03 |
| kluyveromyces_nonfermentans | kluyveromyces_nonfermentans | #N/A | Kluyveromyces_nonfermentans | yHMPu5000034701_kluyveromyces_nonfermentans_170307 | g001343.m1 | #N/A | Kluyveromyces | , | 31.92737 | , | 8.7328 | | | 31927.37 |
| kluyveromyces_siamensis | kluyveromyces_siamensis | #N/A | Kluyveromyces_siamensis | yHMPu5000034700_kluyveromyces_siamensis_180604 | g003460.m1 | #N/A | Kluyveromyces | , | 30.40989 | , | 6.1228 | | | 30409.89 |
| kluyveromyces_starmeri | kluyveromyces_starmeri | #N/A | Kluyveromyces_starmeri | yHDO570_kluyveromyces_starmeri_180604 | g003161.m1 | #N/A | Kluyveromyces | , | 30.73575 | , | 6.0696 | | | 30735.75 |
| kluyveromyces_wickerhamii | kluyveromyces_wickerhamii | #N/A | Kluyveromyces_wickerhamii | yHMPu5000034699_kluyveromyces_wickerhamii_170307 | g000532.m1 | #N/A | Kluyveromyces | , | 33.53587 | , | 6.6051 | | | 33535.87 |
| hanseniasporauvarum | hanseniaspora_uvarum | #N/A | Hanseniaspora_uvarum_DSMZ_2768 | hanseniaspora_uvarum | g000873.m1 | KKA03541.1 | Hanseniaspora | , | 44.31316 | , | 4.4615 | | | 44313.16 |
| hanseniasporapseudoguilliermondi | hanseniaspora_pseudoguilliermondi | #N/A | Hanseniaspora_pseudoguilliermondii | yHMPu5000035695_hanseniaspora_pseudoguilliermondii_160519.haplomerger2 | g000765.m1 | #N/A | Hanseniaspora | , | 42.93928 | , | 4.1968 | | | 42939.28 |
| hanseniasporaclermontiae | hanseniaspora_clermontiae | #N/A | Hanseniaspora_clermontiae | yHMPu5000034963_hanseniaspora_clermontiae_160519.haplomerger2 | g003387.m1 | #N/A | Hanseniaspora | , | 43.04594 | , | 4.4959 | | | 43045.94 |
| hanseniasporavalbyensis | hanseniaspora_valbyensis | #N/A | Hanseniaspora_valbyensis | yHMPu5000034955_hanseniaspora_valbyensis_201018.haplomerger2 | g000788.m1 | OBA26588.1 | Hanseniaspora | , | 46.0101 | , | 4.4316 | | | 46010.1 |
| hanseniasporavinae | hanseniaspora_vinae | #N/A | Hanseniaspora_vineae | yHMPu5000026147_hanseniaspora_vineae_190924.haplomerger2 | g000163.m1 | #N/A | Hanseniaspora | , | 55.14142 | , | 4.9736 | | | 55141.42 |
| cyberlindnerasuaveolens | cyberlindnera_suaveolens | #N/A | Cyberlindnera_suaveolens | yHMPu5000035687_cyberlindnera_suaveolens_160613 | g005674.m1 | #N/A | Cyberlindnera | , | 37.19332 | , | 4.377 | | | 37193.32 |
| cyberlindnerasaturnus | cyberlindnera_saturnus | #N/A | Cyberlindnera_saturnus | yHMPu5000035686_Cyberlindnera_saturnus_SPADES | g001009.m1 | #N/A | Cyberlindnera | , | 37.03333 | , | 4.3708 | | | 37033.33 |
| cyberlindneramisumaiensis | cyberlindnera_misumaiensis | #N/A | Cyberlindnera_misumaiensis | yHMPu5000034979_cyberlindnera_misumaiensis_160519 | g002417.m1 | #N/A | Cyberlindnera | , | 30.88568 | , | 4.8579 | | | 30885.68 |
| Cyberlindnerajadinii | Cyberlindnera_jadinii | #N/A | Cyberlindnera_jadinii | yHMPu5000035336_cyberlindnera_jadinii_180604.haplomerger2 | g002673.m1 | XP_020072378.1 | Cyberlindnera | , | 42.74321 | , | 4.5317 | | | 42743.21 |
| Cyberlindnerafabianii | Cyberlindnera_fabianii | #N/A | Cyberlindnera_fabianii | yHMPu5000035701_cyberlindnera_fabianii_160519 | g005778.m1 | ONH68477.1 | Cyberlindnera | , | 39.35018 | , | 6.1188 | | | 39350.18 |
| cyberlindneramaclurae | cyberlindnera_maclurae | #N/A | Cyberlindnera_maclurae | yHMPu5000035699_cyberlindnera_maclurae_160613 | g003717.m1 | #N/A | Cyberlindnera | , | 23.21122 | , | 4.8781 | | | 23211.22 |
| wickerhamomycesalni | wickerhamomyce_salni | #N/A | Wickerhamomyces_alni | yHMPu5000035274_wickerhamomyces_alni_170307 | g001244.m1 | #N/A | Wickerhamomyces | , | 39.20614 | , | 5.0895 | | | 39206.14 |
| wickerhamomycescanadensis | wickerhamomyces_canadensis | #N/A | Wickerhamomyces_canadensis | yHMPu5000035639_wickerhamomyces_canadensis_160613 | g004559.m1 | #N/A | Wickerhamomyces | , | 25.01526 | , | 4.2917 | | | 25015.26 |
| wickerhamomycesciferrii | wickerhamomyces_ciferrii | #N/A | Wickerhamomyces_ciferrii | yHMPu5000035269_wickerhamomyces_ciferrii_170307 | g005482.m1,g003054.m1 | XP_011277532.1 | Wickerhamomyces | , | 43.91976 | , | 4.1975 | | | 43919.76 |
| Wickerhamomyceschambardii | Wickerhamomyces_chambardii | #N/A | Wickerhamomyces_chambardii | yHMPu5000035270_wickerhamomyces_chambardii_180604.haplomerger2 | g001027.m1 | #N/A | Wickerhamomyces | , | 57.58908 | , | 4.48 | | | 57589.08 |
| Wickerhamomyceshampshirensis | Wickerhamomyces_hampshirensis | #N/A | Wickerhamomyces_hampshirensis | yHMPu5000035268_wickerhamomyces_hampshirensis_160928 | g002138.m1 | #N/A | Wickerhamomyces | , | 47.13197 | , | 4.7469 | | | 47131.97 |
| Wickerhamomycesmucosus | Wickerhamomyces_mucosus | #N/A | Wickerhamomyces_mucosus | yHMPu5000038048_wickerhamomyces_mucosus_170912.haplomerger2 | g002404.m1 | #N/A | Wickerhamomyces | , | 45.13403 | , | 4.7526 | | | 45134.03 |
| Wickerhamomycespiperi | Wickerhamomyces_piperi | #N/A | Wickerhamomyces_pijperi | yHMPu5000035254_wickerhamomyces_pijperi_160928.haplomerger2 | g002584.m1 | #N/A | Wickerhamomyces | , | 32.50732 | , | 4.0717 | | | 32507.32 |
| komagataellaphaffi | komagataella_phaffi | #N/A | Komagataella_phaffii_GS115 | komagataella_phaffi | g002262.m1 | XP_002491603.1 | Komagataella | , | 35.0953 | , | 4.4338 | | | 35095.3 |
| komagataellapopuli | komagataella_populi | #N/A | Komagataella_populi | yHMPu5000026274_komagataella_populi_160519 | g004657.m1 | #N/A | Komagataella | , | 35.2525 | , | 4.5982 | | | 35252.5 |
| komagataellapastoris | komagataella_pastoris | #N/A | Komagataella_pastoris_ATCC_28485 | komagataella_pastoris | g002679.m1 | ANZ75330.1 | Komagataella | , | 35.8892 | , | 4.4453 | | | 35889.2 |
| Komagataellakurtzmanii | Komagataella_kurtzmanii | #N/A | Komagataella_kurtzmanii | yHMPu5000035676_komagataella_kurtzmanii_160613 | g004231.m1 | #N/A | Komagataella | , | 35.15743 | , | 4.4338 | | | 35157.43 |
| Komatagataella | Komatagaella_mondaviorum | #N/A | Komagataella_mondaviorum | yHMPu5000037220_komagataella_mondaviorum_210210 | g004487.m1 | #N/A | Komagataella | , | 35.24556 | , | 4.3928 | | | 35245.56 |
| Kuraishia | Kuraishia_floccosa | #N/A | Kuraishia_floccosa | yHMPu5000034935_kuraishia_floccosa_210210 | g004700.m1 | #N/A | Kuraishia | , | 48.45106 | , | 4.1091 | | | 48451.06 |
| NA | Asbhya_aceri | #N/A | Ashbya_aceri | ashbya_aceri | NA | AGO10388.1 | Eremothecium | | | | | | | |
| Wickerhamomyces_anomalus | Wickerhamomyces_anomalus | #N/A | Wickerhamomyces_anomalus | yHMPu5000035273_wickerhamomyces_anomalus_180604.haplomerger2 | g001657.m1 | TBD | Wickerhamomyces | | | | | | | |

| | | | | | | | |
|---|---|---|---|---|---|---|---|
| saccharomyces_mikatae.4711/1-325 | saccharomyces_mikatae | 4711/1-325 | Saccharomyces_mikatae | 0 | 0 | 1 | *Saccharomyces_mikatae* |
| saccharomyces_paradoxus.4820/1-321 | saccharomyces_paradoxus | 4820/1-321 | Saccharomyces_paradoxus | 0 | 0 | 1 | |
| Saccharomyces_cerevisiae.2608/1-321 | Saccharomyces_cerevisiae | 2608/1-321 | Saccharomyces_cerevisiae | 0 | 0 | 1 | |
| saccharomyces_uvarum.2005/523-836 | saccharomyces_uvarum | 2005/523-836 | Saccharomyces_uvarum | 0 | 0 | 1 | 1 |
| saccharomyces_eubayanus.4603/581-888 | saccharomyces_eubayanus | 4603/581-888 | Saccharomyces_eubayanus | 0 | 0 | 1 | |
| saccharomyces_arboricola.2054/541-857 | saccharomyces_arboricola | 2054/541-857 | Saccharomyces_arboricola | 0 | 1 | 1 | |
| torulaspora_delbrueckii.2077/1-320 | torulaspora_delbrueckii | 2077/1-320 | Torulaspora_delbrueckii | 0 | 0 | 1 | |
| torulaspora_franciscae.1384/1-315 | torulaspora_franciscae | 1384/1-315 | Torulaspora_franciscae | 0 | 0 | 1 | |
| torulaspora_pretoriensis.3518/1-316 | torulaspora_pretoriensis | 3518/1-316 | Torulaspora_pretoriensis | 0 | 0 | 1 | |
| torulaspora_pretoriensis.3519/1-316 | torulaspora_pretoriensis | 3519/1-316 | Torulaspora_pretoriensis | 0 | 0 | 1 | |
| torulaspora_microellipsoides.4259/1-344 | torulaspora_microellipsoides | 4259/1-344 | Torulaspora_microellipsoides | 0 | 0 | 1 | |
| zygosaccharomyces_bailii.3872/1-282 | zygosaccharomyces_bailii | 3872/1-282 | Zygosaccharomyces_bailii | 0 | 0 | 1 | |
| zygotorulaspora_florentina.4182/586-923 | zygotorulaspora_florentina | 4182/586-923 | Zygotorulaspora_florentina | 0 | 0 | 1 | |
| zygosaccharomyces_kombuchaensis.1039/1-275 | zygosaccharomyces_kombuchaensis | 1039/1-275 | Zygosaccharomyces_kombuchaensis | 0 | 0 | 1 | |
| zygosaccharomyces_bisporus.1211/1-279 | zygosaccharomyces_bisporus | 1211/1-279 | Zygosaccharomyces_bisporus | 0 | 0 | 1 | |
| zygotorulaspora_mrakii.2527/1-350 | zygotorulaspora_mrakii | 2527/1-350 | Zygotorulaspora_mrakii | 0 | 0 | 2 | |
| zygosaccharomyces_rouxii.874/1-294 | zygosaccharomyces_rouxii | 874/1-294 | Zygosaccharomyces_rouxii | 0 | 0 | 0 | |
| torulaspora_maleeae.3383/1-304 | torulaspora_maleeae | 3383/1-304 | Torulaspora_maleeae | 0 | 0 | 1 | |
| saccharomyces_kudriavzevii.4688/573-833 | saccharomyces_kudriavzevii | 4688/573-833 | Saccharomyces_kudriavzevii | 0 | 0 | 1 | |
| lachancea_kluyveri.4105/1-299 | lachancea_kluyveri | 4105/1-299 | Lachancea_kluyveri | 0 | 0 | 1 | |
| naumovozyma_dairenensis.2516/1-258 | naumovozyma_dairenensis | 2516/1-258 | Naumovozyma_dairenensis | 0 | 0 | 1 | |
| naumovozyma_dairenensis.2516/421-457 | naumovozyma_dairenensis | 2516/421-457 | Naumovozyma_dairenensis | 0 | 0 | 1 | |
| kazachstania_viticola.4751/1-360 | kazachstania_viticola | 4751/1-360 | Kazachstania_viticola | 0 | 0 | 1 | |
| naumovozyma_castellii.578/448-789 | naumovozyma_castellii | 578/448-789 | Naumovozyma_castellii | 0 | 0 | 1 | |
| lachancea_fermentati.50/7-173 | lachancea_fermentati | 50/7-173 | Lachancea_fermentati | 0 | 0 | 1 | |
| lachancea_fermentati.50/195-281 | lachancea_fermentati | 50/195-281 | Lachancea_fermentati | 0 | 0 | 1 | |
| lachancea_cidri.2690/37-204 | lachancea_cidri | 2690/37-204 | Lachancea_cidri | 0 | 0 | 0 | |
| lachancea_cidri.2690/225-312 | lachancea_cidri | 2690/225-312 | Lachancea_cidri | 0 | 0 | 0 | |
| nakaseomyces_delphensis.4813/1-136 | nakaseomyces_delphensis | 4813/1-136 | Nakaseomyces_delphensis | 0 | 0 | 1 | |
| nakaseomyces_delphensis.4813/185-236 | nakaseomyces_delphensis | 4813/185-236 | Nakaseomyces_delphensis | 0 | 0 | 1 | |
| vanderwaltozyma_polyspora.1882/5-40 | vanderwaltozyma_polyspora | 1882/5-40 | Vanderwaltozyma_polyspora | 0 | 0 | 1 | |
| vanderwaltozyma_polyspora.1882/129-163 | vanderwaltozyma_polyspora | 1882/129-163 | Vanderwaltozyma_polyspora | 0 | 0 | 1 | |
| vanderwaltozyma_polyspora.1882/269-407 | vanderwaltozyma_polyspora | 1882/269-407 | Vanderwaltozyma_polyspora | 0 | 0 | 1 | |
| kluyveromyces_aestuarii.3100/2-165 | kluyveromyces_aestuarii | 3100/2-165 | Kluyveromyces_aestuarii | 0 | 0 | 0 | |
| kluyveromyces_aestuarii.3100/171-271 | kluyveromyces_aestuarii | 3100/171-271 | Kluyveromyces_aestuarii | 0 | 0 | 0 | |
| nakaseomyces_bacillisporus.3279/3-99 | nakaseomyces_bacillisporus | 3279/3-99 | Nakaseomyces_bacillisporus | 0 | 0 | 1 | |
| nakaseomyces_bacillisporus.3279/192-272 | nakaseomyces_bacillisporus | 3279/192-272 | Nakaseomyces_bacillisporus | 0 | 0 | 1 | |
| nakaseomyces_bracarensis.2954/1-119 | nakaseomyces_bracarensis | 2954/1-119 | Nakaseomyces_bracarensis | 0 | 0 | 1 | |
| nakaseomyces_bracarensis.2954/121-270 | nakaseomyces_bracarensis | 2954/121-270 | Nakaseomyces_bracarensis | 0 | 0 | 1 | |
| lachancea_mirantina.815/3-303 | lachancea_mirantina | 815/3-303 | Lachancea_mirantina | 0 | 0 | 1 | |
| nakaseomyces_nivariensis.2803/1-131 | nakaseomyces_nivariensis | 2803/1-131 | Nakaseomyces_nivariensis | 0 | 0 | 1 | |
| nakaseomyces_nivariensis.2803/113-233 | nakaseomyces_nivariensis | 2803/113-233 | Nakaseomyces_nivariensis | 0 | 0 | 1 | |
| lachancea_quebecensis.4341/20-160 | lachancea_quebecensis | 4341/20-160 | Lachancea_quebecensis | 0 | 0 | 1 | |
| lachancea_quebecensis.4341/175-262 | lachancea_quebecensis | 4341/175-262 | Lachancea_quebecensis | 0 | 0 | 1 | |
| lachancea_waltii.2068/10-274 | lachancea_waltii | 2068/10-274 | Lachancea_waltii | 0 | 0 | 1 | |
| candida_glabrata.4846/1-104 | candida_glabrata | 4846/1-104 | Candida_glabrata | 0 | 0 | 0 | |
| candida_glabrata.4846/113-241 | candida_glabrata | 4846/113-241 | Candida_glabrata | 0 | 0 | 0 | |
| kazachstania_bromeliacearum.151/1-128 | kazachstania_bromeliacearum | 151/1-128 | Kazachstania_bromeliacearum | 0 | 0 | 1 | |
| kazachstania_bromeliacearum.151/123-235 | kazachstania_bromeliacearum | 151/123-235 | Kazachstania_bromeliacearum | 0 | 0 | 1 | |
| nakaseomyces_castellii.4169/1-389 | nakaseomyces_castellii | 4169/1-389 | Nakaseomyces_castellii | 0 | 0 | 1 | |
| kazachstania_turicensis.4764/1-137 | kazachstania_turicensis | 4764/1-137 | Kazachstania_turicensis | 0 | 0 | 1 | |
| kazachstania_turicensis.4764/312-424 | kazachstania_turicensis | 4764/312-424 | Kazachstania_turicensis | 0 | 0 | 1 | |
| kazachstania_naganishii.4691/6-132 | kazachstania_naganishii | 4691/6-132 | Kazachstania_naganishii | 0 | 0 | 1 | |
| kazachstania_naganishii.4691/183-284 | kazachstania_naganishii | 4691/183-284 | Kazachstania_naganishii | 0 | 0 | 1 | |
| lachancea_lanzarotensis.976/4-340 | lachancea_lanzarotensis | 976/4-340 | Lachancea_lanzarotensis | 0 | 0 | 1 | |
| lachancea_dasiensis.337/5-360 | lachancea_dasiensis | 337/5-360 | Lachancea_dasiensis | 0 | 0 | 1 | |
| kazachstania_martiniae.4455/1-41 | kazachstania_martiniae | 4455/1-41 | Kazachstania_martiniae | 0 | 0 | 1 | |
| kazachstania_martiniae.4455/117-266 | kazachstania_martiniae | 4455/117-266 | Kazachstania_martiniae | 0 | 0 | 1 | |
| lachancea_nothofagi.389/11-289 | lachancea_nothofagi | 389/11-289 | Lachancea_nothofagi | 0 | 0 | 1 | |
| kazachstania_kunashirensis.4945/1-131 | kazachstania_kunashirensis | 4945/1-131 | Kazachstania_kunashirensis | 0 | 0 | 2 | |
| kazachstania_kunashirensis.4945/219-375 | kazachstania_kunashirensis | 4945/219-375 | Kazachstania_kunashirensis | 0 | 0 | 2 | |
| lachancea_fantastica.3409/4-159 | lachancea_fantastica | 3409/4-159 | Lachancea_fantastica | 0 | 0 | 0 | |
| lachancea_fantastica.3409/290-341 | lachancea_fantastica | 3409/290-341 | Lachancea_fantastica | 0 | 0 | 0 | |
| tetrapisispora_iriomotensis.4707/5-135 | tetrapisispora_iriomotensis | 4707/5-135 | Tetrapisispora_iriomotensis | 0 | 0 | 1 | |
| tetrapisispora_iriomotensis.4707/184-374 | tetrapisispora_iriomotensis | 4707/184-374 | Tetrapisispora_iriomotensis | 0 | 0 | 1 | |
| kazachstania_africana.2163/1-129 | kazachstania_africana | 2163/1-129 | Kazachstania_africana | 0 | 0 | 1 | |
| kazachstania_africana.2163/200-352 | kazachstania_africana | 2163/200-352 | Kazachstania_africana | 0 | 0 | 1 | |
| kluyveromyces_lactis.677/61-176 | kluyveromyces_lactis | 677/61-176 | Kluyveromyces_lactis | 0 | 0 | 0 | |
| kluyveromyces_lactis.677/248-278 | kluyveromyces_lactis | 677/248-278 | Kluyveromyces_lactis | 0 | 0 | 0 | |
| lachancea_meyersii.3908/11-299 | lachancea_meyersii | 3908/11-299 | Lachancea_meyersii | 0 | 0 | 1 | |
| kazachstania_siamensis.3992/1-129 | kazachstania_siamensis | 3992/1-129 | Kazachstania_siamensis | 0 | 0 | 1 | |
| kazachstania_siamensis.3992/152-287 | kazachstania_siamensis | 3992/152-287 | Kazachstania_siamensis | 0 | 0 | 1 | |
| kluyveromyces_dobzhanskii.483/44-194 | kluyveromyces_dobzhanskii | 483/44-194 | Kluyveromyces_dobzhanskii | 0 | 0 | 1 | |
| kluyveromyces_dobzhanskii.483/236-281 | kluyveromyces_dobzhanskii | 483/236-281 | Kluyveromyces_dobzhanskii | 0 | 0 | 1 | |
| eremothecium_cymbalariae.2716/144-220 | eremothecium_cymbalariae | 2716/144-220 | Eremothecium_cymbalariae | 0 | 0 | 1 | |

| | | | | | | |
|---|---|---|---|---|---|---|
| eremothecium_cymbalariae.2716/259-284 | eremothecium_cymbalariae | 2716/259-284 | Eremothecium_cymbalariae | 0 | 0 | 1 |
| lachancea_thermotolerans.1896/10-57 | lachancea_thermotolerans | 1896/10-57 | Lachancea_thermotolerans | 0 | 0 | 1 |
| lachancea_thermotolerans.1896/87-183 | lachancea_thermotolerans | 1896/87-183 | Lachancea_thermotolerans | 0 | 0 | 1 |
| kazachstania_taianensis.3897/1-44 | kazachstania_taianensis | 3897/1-44 | Kazachstania_taianensis | 0 | 0 | 1 |
| kazachstania_taianensis.3897/130-238 | kazachstania_taianensis | 3897/130-238 | Kazachstania_taianensis | 0 | 0 | 1 |
| kazachstania_aerobia.2672/1-349 | kazachstania_aerobia | 2672/1-349 | Kazachstania_aerobia | 0 | 0 | 1 |
| kazachstania_unispora.3458/1-342 | kazachstania_unispora | 3458/1-342 | Kazachstania_unispora | 0 | 0 | 1 |
| kazachstania_yakushimaensis.3740/307-438 | kazachstania_yakushimaensis | 3740/307-438 | Kazachstania_yakushimaensis | 0 | 0 | 0 |
| kazachstania_yakushimaensis.3740/716-872 | kazachstania_yakushimaensis | 3740/716-872 | Kazachstania_yakushimaensis | 0 | 0 | 0 |
| kazachstania_spencerorum.3580/3-278 | kazachstania_spencerorum | 3580/3-278 | Kazachstania_spencerorum | 0 | 0 | 1 |
| kazachstania_spencerorum.3580/351-408 | kazachstania_spencerorum | 3580/351-408 | Kazachstania_spencerorum | 0 | 0 | 1 |
| kazachstania_rosinii.1249/3-239 | kazachstania_rosinii | 1249/3-239 | Kazachstania_rosinii | 0 | 0 | 0 |
| kazachstania_solicola.5780/1-137 | kazachstania_solicola | 5780/1-137 | Kazachstania_solicola | 0 | 0 | 1 |
| kazachstania_solicola.5780/187-342 | kazachstania_solicola | 5780/187-342 | Kazachstania_solicola | 0 | 0 | 1 |
| tetrapisispora_blattae.1015/1-244 | tetrapisispora_blattae | 1015/1-244 | Tetrapisispora_blattae | 0 | 0 | 1 |
| tetrapisispora_blattae.1015/304-465 | tetrapisispora_blattae | 1015/304-465 | Tetrapisispora_blattae | 0 | 0 | 1 |
| tetrapisispora_namnaonensis.1469/6-127 | tetrapisispora_namnaonensis | 1469/6-127 | Tetrapisispora_namnaonensis | 0 | 0 | 0 |
| kluyveromyces_marxianus.2881/64-175 | kluyveromyces_marxianus | 2881/64-175 | Kluyveromyces_marxianus | 0 | 0 | 0 |
| eremothecium_coryli.3485/574-629 | eremothecium_coryli | 3485/574-629 | Eremothecium_coryli | 0 | 0 | 1 |
| lachancea_nothofagi.512/203-358 | lachancea_nothofagi | 512/203-358 | Lachancea_nothofagi | 0 | 0 | 1 |

| | | | | | | | | | | | | | | | | | | | | | | | | |
|---|---|---|---|---|---|---|---|---|---|---|---|---|---|---|---|---|---|---|---|---|---|---|---|---|
| #=GS | saccharomyces_mikatae.4711/1-325 | DE | [subseq | from] | snap_masked-Smik_8-processed-gene-4.133-mRNA-1_1 | gene=snap_masked-Smik_8-processed-gene-4.133 | CDS=1-975 | | | | | | | | | | | | | | | | | |
| #=GS | saccharomyces_paradoxus.4820/1-321 | DE | [subseq | from] | genemark-Spar_8-processed-gene-4.108-mRNA-1_1 | gene=genemark-Spar_8-processed-gene-4.108 | CDS=1-963 | | | | | | | | | | | | | | | | | |
| #=GS | Saccharomyces_cerevisiae.2608/1-321 | DE | [subseq | from] | YHR195W | NVJ1 | SGDID:S000001238, | Chr | VIII | from | 490747-491712 , | Genome | Release | 64-2-1, | Verified | ORF, | Nuclear envelope protein; anchored to the nuclear inner membrane, that interacts | during | piecemeal | microautophagy | of | the | nucleus | (PMN)" |
| #=GS | saccharomyces_uvarum.2005/523-836 | DE | [subseq | from] | augustus_masked-Sbay_15-processed-gene-6.183-mRNA-1_1 | gene=augustus_masked-Sbay_15-processed-gene-6.183 | CDS=1-4203 | | | | | | | | | | | | | | | | | |
| #=GS | saccharomyces_eubayanus.4603/581-888 | DE | [subseq | from] | augustus_masked-chrXV-processed-gene-6.90-mRNA-1_1 | gene=augustus_masked-chrXV-processed-gene-6.90 | CDS=1-4320 | | | | | | | | | | | | | | | | | |
| #=GS | saccharomyces_arboricola.2054/541-857 | DE | [subseq | from] | augustus_masked-NC_026178.1-processed-gene-4.111-mRNA-1_1 | gene=augustus_masked-NC_026178.1-processed-gene-4.111 | CDS=1-2571 | | | | | | | | | | | | | | | | | |
| #=GS | torulaspora_delbrueckii.2077/1-320 | DE | [subseq | from] | snap_masked-scaffold_4-processed-gene-1.34-mRNA-1_1 | gene=snap_masked-scaffold_4-processed-gene-1.34 | CDS=1-960 | | | | | | | | | | | | | | | | | |
| #=GS | torulaspora_franciscae.1384/1-315 | DE | [subseq | from] | snap_masked-flattened_line_10-processed-gene-1.40-mRNA-1_1 | gene=snap_masked-flattened_line_10-processed-gene-1.40 | CDS=1-945 | | | | | | | | | | | | | | | | | |
| #=GS | torulaspora_pretoriensis.3518/1-316 | DE | [subseq | from] | augustus_masked-flattened_line_3-processed-gene-4.12-mRNA-1_1 | gene=augustus_masked-flattened_line_3-processed-gene-4.12 | CDS=1-948 | | | | | | | | | | | | | | | | | |
| #=GS | torulaspora_pretoriensis.3519/1-316 | DE | [subseq | from] | augustus_masked-flattened_line_3-processed-gene-4.38-mRNA-1_1 | gene=augustus_masked-flattened_line_3-processed-gene-4.38 | CDS=1-948 | | | | | | | | | | | | | | | | | |
| #=GS | torulaspora_microellipsoides.4259/1-344 | DE | [subseq | from] | augustus_masked-flattened_line_52-processed-gene-0.132-mRNA-1_1 | gene=augustus_masked-flattened_line_52-processed-gene-0.132 | CDS=1-1032 | | | | | | | | | | | | | | | | | |
| #=GS | zygosaccharomyces_bailii.3872/1-282 | DE | [subseq | from] | augustus_masked-ZYBA0S10-processed-gene-3.53-mRNA-1_1 | gene=augustus_masked-ZYBA0S10-processed-gene-3.53 | CDS=1-846 | | | | | | | | | | | | | | | | | |
| #=GS | zygotorulaspora_florentina.4182/586-923 | DE | [subseq | from] | augustus_masked-NODE_6_length_562868_cov_24.3835_ID_11-processed-gene-2.47-mRNA-1_1 | gene=augustus_masked-NODE_6_length_562868_cov_24.3835_ID_11-processed-gene-2.47 | CDS=1-2769 | | | | | | | | | | | | | | | | | |
| #=GS | zygosaccharomyces_kombuchaensis.1039/1-275 | DE | [subseq | from] | snap_masked-NODE_1_length_271579_cov_10.1807_ID_1-processed-gene-1.23-mRNA-1_1 | gene=snap_masked-NODE_1_length_271579_cov_10.1807_ID_1-processed-gene-1.23 | CDS=1-825 | | | | | | | | | | | | | | | | | |
| #=GS | zygosaccharomyces_bisporus.1211/1-279 | DE | [subseq | from] | snap_masked-flattened_line_16-processed-gene-0.27-mRNA-1_1 | gene=snap_masked-flattened_line_16-processed-gene-0.27 | CDS=1-837 | | | | | | | | | | | | | | | | | |
| #=GS | zygotorulaspora_mrakii.2527/1-350 | DE | [subseq | from] | snap_masked-NODE_2_length_708340_cov_18.0358_ID_2767-processed-gene-4.105-mRNA-1_1 | gene=snap_masked-NODE_2_length_708340_cov_18.0358_ID_2767-processed-gene-4.105 | CDS=1-1050 | | | | | | | | | | | | | | | | | |
| #=GS | zygosaccharomyces_rouxii.874/1-294 | DE | [subseq | from] | snap_masked-Zyro0B-processed-gene-6.126-mRNA-1_1 | gene=snap_masked-Zyro0B-processed-gene-6.126 | CDS=1-882 | | | | | | | | | | | | | | | | | |
| #=GS | torulaspora_maleeae.3383/1-304 | DE | [subseq | from] | snap_masked-NODE_5_length_764704_cov_59.617_ID_1253-processed-gene-6.109-mRNA-1_1 | gene=snap_masked-NODE_5_length_764704_cov_59.617_ID_1253-processed-gene-6.109 | CDS=1-912 | | | | | | | | | | | | | | | | | |
| #=GS | saccharomyces_kudriavzevii.4688/573-833 | DE | [subseq | from] | augustus_masked-Skud_8-processed-gene-3.158-mRNA-1_1 | gene=augustus_masked-Skud_8-processed-gene-3.158 | CDS=1-6507 | | | | | | | | | | | | | | | | | |
| #=GS | lachancea_kluyveri.4105/1-299 | DE | [subseq | from] | snap_masked-SAKL0G-processed-gene-15.78-mRNA-1_1 | gene=snap_masked-SAKL0G-processed-gene-15.78 | CDS=1-897 | | | | | | | | | | | | | | | | | |
| #=GS | naumovozyma_dairenensis.2516/1-258 | DE | [subseq | from] | augustus_masked-NC_016482.1-processed-gene-7.22-mRNA-1_1 | gene=augustus_masked-NC_016482.1-processed-gene-7.22 | CDS=1-1371 | | | | | | | | | | | | | | | | | |
| #=GS | naumovozyma_dairenensis.2516/421-457 | DE | [subseq | from] | augustus_masked-NC_016482.1-processed-gene-7.22-mRNA-1_1 | gene=augustus_masked-NC_016482.1-processed-gene-7.22 | CDS=1-1371 | | | | | | | | | | | | | | | | | |
| #=GS | kazachstania_viticola.4751/1-360 | DE | [subseq | from] | snap_masked-scf7180000050361-processed-gene-0.67-mRNA-1_1 | gene=snap_masked-scf7180000050361-processed-gene-0.67 | CDS=1-1080 | | | | | | | | | | | | | | | | | |
| #=GS | naumovozyma_castellii.578/448-789 | DE | [subseq | from] | snap_masked-HE576752-processed-gene-12.71-mRNA-1_1 | gene=snap_masked-HE576752-processed-gene-12.71 | CDS=1-2379 | | | | | | | | | | | | | | | | | |
| #=GS | lachancea_fermentati.50/7-173 | DE | [subseq | from] | augustus_masked-LAFE0A-processed-gene-1.131-mRNA-1_1 | gene=augustus_masked-LAFE0A-processed-gene-1.131 | CDS=1-843 | | | | | | | | | | | | | | | | | |
| #=GS | lachancea_fermentati.50/195-281 | DE | [subseq | from] | augustus_masked-LAFE0A-processed-gene-1.131-mRNA-1_1 | gene=augustus_masked-LAFE0A-processed-gene-1.131 | CDS=1-843 | | | | | | | | | | | | | | | | | |
| #=GS | lachancea_cidri.2690/37-204 | DE | [subseq | from] | augustus_masked-LACI0G-processed-gene-1.33-mRNA-1_1 | gene=augustus_masked-LACI0G-processed-gene-1.33 | CDS=1-936 | | | | | | | | | | | | | | | | | |
| #=GS | lachancea_cidri.2690/225-312 | DE | [subseq | from] | augustus_masked-LACI0G-processed-gene-1.33-mRNA-1_1 | gene=augustus_masked-LACI0G-processed-gene-1.33 | CDS=1-936 | | | | | | | | | | | | | | | | | |
| #=GS | nakaseomyces_delphensis.4813/1-136 | DE | [subseq | from] | augustus_masked-NADE0S27-processed-gene-3.39-mRNA-1_1 | gene=augustus_masked-NADE0S27-processed-gene-3.39 | CDS=1-708 | | | | | | | | | | | | | | | | | |
| #=GS | nakaseomyces_delphensis.4813/185-236 | DE | [subseq | from] | augustus_masked-NADE0S27-processed-gene-3.39-mRNA-1_1 | gene=augustus_masked-NADE0S27-processed-gene-3.39 | CDS=1-708 | | | | | | | | | | | | | | | | | |
| #=GS | vanderwaltozyma_polyspora.1882/5-40 | DE | [subseq | from] | snap_masked-NW_001834637.1-processed-gene-0.82-mRNA-1_1 | gene=snap_masked-NW_001834637.1-processed-gene-0.82 | CDS=1-1221 | | | | | | | | | | | | | | | | | |
| #=GS | vanderwaltozyma_polyspora.1882/129-163 | DE | [subseq | from] | snap_masked-NW_001834637.1-processed-gene-0.82-mRNA-1_1 | gene=snap_masked-NW_001834637.1-processed-gene-0.82 | CDS=1-1221 | | | | | | | | | | | | | | | | | |
| #=GS | vanderwaltozyma_polyspora.1882/269-407 | DE | [subseq | from] | snap_masked-NW_001834637.1-processed-gene-0.82-mRNA-1_1 | gene=snap_masked-NW_001834637.1-processed-gene-0.82 | CDS=1-1221 | | | | | | | | | | | | | | | | | |
| #=GS | kluyveromyces_aestuarii.3100/2-165 | DE | [subseq | from] | snap_masked-NODE_4_length_706832_cov_27.5555_ID_7-processed-gene-0.39-mRNA-1_1 | gene=snap_masked-NODE_4_length_706832_cov_27.5555_ID_7-processed-gene-0.39 | CDS=1-816 | | | | | | | | | | | | | | | | | |
| #=GS | kluyveromyces_aestuarii.3100/171-271 | DE | [subseq | from] | snap_masked-NODE_4_length_706832_cov_27.5555_ID_7-processed-gene-0.39-mRNA-1_1 | gene=snap_masked-NODE_4_length_706832_cov_27.5555_ID_7-processed-gene-0.39 | CDS=1-816 | | | | | | | | | | | | | | | | | |
| #=GS | nakaseomyces_bacillisporus.3279/3-99 | DE | [subseq | from] | augustus_masked-NABA0S29-processed-gene-0.108-mRNA-1_1 | gene=augustus_masked-NABA0S29-processed-gene-0.108 | CDS=1-816 | | | | | | | | | | | | | | | | | |
| #=GS | nakaseomyces_bacillisporus.3279/192-272 | DE | [subseq | from] | augustus_masked-NABA0S29-processed-gene-0.108-mRNA-1_1 | gene=augustus_masked-NABA0S29-processed-gene-0.108 | CDS=1-816 | | | | | | | | | | | | | | | | | |
| #=GS | nakaseomyces_bracarensis.2954/1-119 | DE | [subseq | from] | snap_masked-CABR0S29-processed-gene-6.0-mRNA-1_1 | gene=snap_masked-CABR0S29-processed-gene-6.0 | CDS=1-810 | | | | | | | | | | | | | | | | | |
| #=GS | nakaseomyces_bracarensis.2954/121-270 | DE | [subseq | from] | snap_masked-CABR0S29-processed-gene-6.0-mRNA-1_1 | gene=snap_masked-CABR0S29-processed-gene-6.0 | CDS=1-810 | | | | | | | | | | | | | | | | | |
| #=GS | lachancea_mirantina.815/3-303 | DE | [subseq | from] | augustus_masked-LAMI0C-processed-gene-0.22-mRNA-1_1 | gene=augustus_masked-LAMI0C-processed-gene-0.22 | CDS=1-909 | | | | | | | | | | | | | | | | | |
| #=GS | nakaseomyces_nivariensis.2803/1-131 | DE | [subseq | from] | snap_masked-CANI0S17-processed-gene-4.132-mRNA-1_1 | gene=snap_masked-CANI0S17-processed-gene-4.132 | CDS=1-699 | | | | | | | | | | | | | | | | | |
| #=GS | nakaseomyces_nivariensis.2803/113-233 | DE | [subseq | from] | snap_masked-CANI0S17-processed-gene-4.132-mRNA-1_1 | gene=snap_masked-CANI0S17-processed-gene-4.132 | CDS=1-699 | | | | | | | | | | | | | | | | | |
| #=GS | lachancea_quebecensis.4341/20-160 | DE | [subseq | from] | snap_masked-LAQU0S22-processed-gene-0.116-mRNA-1_1 | gene=snap_masked-LAQU0S22-processed-gene-0.116 | CDS=1-786 | | | | | | | | | | | | | | | | | |
| #=GS | lachancea_quebecensis.4341/175-262 | DE | [subseq | from] | snap_masked-LAQU0S22-processed-gene-0.116-mRNA-1_1 | gene=snap_masked-LAQU0S22-processed-gene-0.116 | CDS=1-786 | | | | | | | | | | | | | | | | | |
| #=GS | lachancea_waltii.2068/10-274 | DE | [subseq | from] | snap_masked-AADM01000119.1-processed-gene-0.11-mRNA-1_1 | gene=snap_masked-AADM01000119.1-processed-gene-0.11 | CDS=1-822 | | | | | | | | | | | | | | | | | |
| #=GS | candida_glabrata.4846/1-104 | DE | [subseq | from] | snap_masked-CR380959-processed-gene-7.12-mRNA-1_1 | gene=snap_masked-CR380959-processed-gene-7.12 | CDS=1-723 | | | | | | | | | | | | | | | | | |
| #=GS | candida_glabrata.4846/113-241 | DE | [subseq | from] | snap_masked-CR380959-processed-gene-7.12-mRNA-1_1 | gene=snap_masked-CR380959-processed-gene-7.12 | CDS=1-723 | | | | | | | | | | | | | | | | | |
| #=GS | kazachstania_bromeliacearum.151/1-128 | DE | [subseq | from] | snap_masked-NODE_10_length_278180_cov_22.5073_ID_19-processed-gene-1.127-mRNA-1_1 | gene=snap_masked-NODE_10_length_278180_cov_22.5073_ID_19-processed-gene-1.127 | CDS=1-711 | | | | | | | | | | | | | | | | | |
| #=GS | kazachstania_bromeliacearum.151/123-235 | DE | [subseq | from] | snap_masked-NODE_10_length_278180_cov_22.5073_ID_19-processed-gene-1.127-mRNA-1_1 | gene=snap_masked-NODE_10_length_278180_cov_22.5073_ID_19-processed-gene-1.127 | CDS=1-711 | | | | | | | | | | | | | | | | | |

| | | | | | | | |
|---|---|---|---|---|---|---|---|
| #=GS | nakaseomyces_castellii.4169/1-389 | DE | [subseq | from] | snap_masked-CACA0S33-processed-gene-9.107-mRNA-1_1 | gene=snap_masked-CACA0S33-processed-gene-9.107 | CDS=1-1167 |
| #=GS | kazachstania_turicensis.4764/1-137 | DE | [subseq | from] | snap_masked-scf7180000049581-processed-gene-4.63-mRNA-1_1 | gene=snap_masked-scf7180000049581-processed-gene-4.63 | CDS=1-1272 |
| #=GS | kazachstania_turicensis.4764/312-424 | DE | [subseq | from] | snap_masked-scf7180000049581-processed-gene-4.63-mRNA-1_1 | gene=snap_masked-scf7180000049581-processed-gene-4.63 | CDS=1-1272 |
| #=GS | kazachstania_naganishii.4691/6-132 | DE | [subseq | from] | snap_masked-HE978326-processed-gene-0.118-mRNA-1_1 | gene=snap_masked-HE978326-processed-gene-0.118 | CDS=1-852 |
| #=GS | kazachstania_naganishii.4691/183-284 | DE | [subseq | from] | snap_masked-HE978326-processed-gene-0.118-mRNA-1_1 | gene=snap_masked-HE978326-processed-gene-0.118 | CDS=1-852 |
| #=GS | lachancea_lanzarotensis.976/4-340 | DE | [subseq | from] | augustus_masked-LN736361.1-processed-gene-3.12-mRNA-1_1 | gene=augustus_masked-LN736361.1-processed-gene-3.12 | CDS=1-1020 |
| #=GS | lachancea_dasiensis.337/5-360 | DE | [subseq | from] | augustus_masked-LADA0A-processed-gene-7.6-mRNA-1_1 | gene=augustus_masked-LADA0A-processed-gene-7.6 | CDS=1-1080 |
| #=GS | kazachstania_martiniae.4455/1-41 | DE | [subseq | from] | snap_masked-scf7180000030514-processed-gene-0.103-mRNA-1_1 | gene=snap_masked-scf7180000030514-processed-gene-0.103 | CDS=1-798 |
| #=GS | kazachstania_martiniae.4455/117-266 | DE | [subseq | from] | snap_masked-scf7180000030514-processed-gene-0.103-mRNA-1_1 | gene=snap_masked-scf7180000030514-processed-gene-0.103 | CDS=1-798 |
| #=GS | lachancea_nothofagi.389/11-289 | DE | [subseq | from] | snap_masked-LANO0B-processed-gene-1.74-mRNA-1_1 | gene=snap_masked-LANO0B-processed-gene-1.74 | CDS=1-867 |
| #=GS | kazachstania_kunashirensis.4945/1-131 | DE | [subseq | from] | snap_masked-scf7180000008985-processed-gene-1.61-mRNA-1_1 | gene=snap_masked-scf7180000008985-processed-gene-1.61 | CDS=1-1125 |
| #=GS | kazachstania_kunashirensis.4945/219-375 | DE | [subseq | from] | snap_masked-scf7180000008985-processed-gene-1.61-mRNA-1_1 | gene=snap_masked-scf7180000008985-processed-gene-1.61 | CDS=1-1125 |
| #=GS | lachancea_fantastica.3409/4-159 | DE | [subseq | from] | snap_masked-LAFA0F-processed-gene-10.9-mRNA-1_1 | gene=snap_masked-LAFA0F-processed-gene-10.9 | CDS=1-1023 |
| #=GS | lachancea_fantastica.3409/290-341 | DE | [subseq | from] | snap_masked-LAFA0F-processed-gene-10.9-mRNA-1_1 | gene=snap_masked-LAFA0F-processed-gene-10.9 | CDS=1-1023 |
| #=GS | tetrapisispora_iriomotensis.4707/5-135 | DE | [subseq | from] | augustus_masked-flattened_line_57-processed-gene-0.130-mRNA-1_1 | gene=augustus_masked-flattened_line_57-processed-gene-0.130 | CDS=1-1122 |
| #=GS | tetrapisispora_iriomotensis.4707/184-374 | DE | [subseq | from] | augustus_masked-flattened_line_57-processed-gene-0.130-mRNA-1_1 | gene=augustus_masked-flattened_line_57-processed-gene-0.130 | CDS=1-1122 |
| #=GS | kazachstania_africana.2163/1-129 | DE | [subseq | from] | augustus_masked-scaffold_2-processed-gene-13.205-mRNA-1_1 | gene=augustus_masked-scaffold_2-processed-gene-13.205 | CDS=1-1059 |
| #=GS | kazachstania_africana.2163/200-352 | DE | [subseq | from] | augustus_masked-scaffold_2-processed-gene-13.205-mRNA-1_1 | gene=augustus_masked-scaffold_2-processed-gene-13.205 | CDS=1-1059 |
| #=GS | kluyveromyces_lactis.677/61-176 | DE | [subseq | from] | snap_masked-CR382122-processed-gene-3.49-mRNA-1_1 | gene=snap_masked-CR382122-processed-gene-3.49 | CDS=1-837 |
| #=GS | kluyveromyces_lactis.677/248-278 | DE | [subseq | from] | snap_masked-CR382122-processed-gene-3.49-mRNA-1_1 | gene=snap_masked-CR382122-processed-gene-3.49 | CDS=1-837 |
| #=GS | lachancea_meyersii.3908/11-299 | DE | [subseq | from] | snap_masked-LAME0G-processed-gene-17.24-mRNA-1_1 | gene=snap_masked-LAME0G-processed-gene-17.24 | CDS=1-897 |
| #=GS | kazachstania_siamensis.3992/1-129 | DE | [subseq | from] | augustus_masked-NODE_61_length_63804_cov_14.6622_ID_121-processed-gene-0.80-mRNA-1_1 | gene=augustus_masked-NODE_61_length_63804_cov_14.6622_ID_121-processed-gene-0.80 | CDS=1-867 |
| #=GS | kazachstania_siamensis.3992/152-287 | DE | [subseq | from] | augustus_masked-NODE_61_length_63804_cov_14.6622_ID_121-processed-gene-0.80-mRNA-1_1 | gene=augustus_masked-NODE_61_length_63804_cov_14.6622_ID_121-processed-gene-0.80 | CDS=1-867 |
| #=GS | kluyveromyces_dobzhanskii.483/44-194 | DE | [subseq | from] | snap_masked-NODE_13_length_251415_cov_20.6991_ID_25-processed-gene-1.144-mRNA-1_1 | gene=snap_masked-NODE_13_length_251415_cov_20.6991_ID_25-processed-gene-1.144 | CDS=1-846 |
| #=GS | kluyveromyces_dobzhanskii.483/236-281 | DE | [subseq | from] | snap_masked-NODE_13_length_251415_cov_20.6991_ID_25-processed-gene-1.144-mRNA-1_1 | gene=snap_masked-NODE_13_length_251415_cov_20.6991_ID_25-processed-gene-1.144 | CDS=1-846 |
| #=GS | eremothecium_cymbalariae.2716/144-220 | DE | [subseq | from] | snap_masked-NC_016453.1-processed-gene-3.121-mRNA-1_1 | gene=snap_masked-NC_016453.1-processed-gene-3.121 | CDS=1-852 |
| #=GS | eremothecium_cymbalariae.2716/259-284 | DE | [subseq | from] | snap_masked-NC_016453.1-processed-gene-3.121-mRNA-1_1 | gene=snap_masked-NC_016453.1-processed-gene-3.121 | CDS=1-852 |
| #=GS | lachancea_thermotolerans.1896/10-57 | DE | [subseq | from] | snap_masked-CU928168-processed-gene-14.65-mRNA-1_1 | gene=snap_masked-CU928168-processed-gene-14.65 | CDS=1-549 |
| #=GS | lachancea_thermotolerans.1896/87-183 | DE | [subseq | from] | snap_masked-CU928168-processed-gene-14.65-mRNA-1_1 | gene=snap_masked-CU928168-processed-gene-14.65 | CDS=1-549 |
| #=GS | kazachstania_taianensis.3897/1-44 | DE | [subseq | from] | augustus_masked-NODE_5_length_452406_cov_49.1731_ID_9-processed-gene-3.120-mRNA-1_1 | gene=augustus_masked-NODE_5_length_452406_cov_49.1731_ID_9-processed-gene-3.120 | CDS=1-717 |
| #=GS | kazachstania_taianensis.3897/130-238 | DE | [subseq | from] | augustus_masked-NODE_5_length_452406_cov_49.1731_ID_9-processed-gene-3.120-mRNA-1_1 | gene=augustus_masked-NODE_5_length_452406_cov_49.1731_ID_9-processed-gene-3.120 | CDS=1-717 |
| #=GS | kazachstania_aerobia.2672/1-349 | DE | [subseq | from] | augustus_masked-scf7180000020169-processed-gene-0.188-mRNA-1_1 | gene=augustus_masked-scf7180000020169-processed-gene-0.188 | CDS=1-1053 |
| #=GS | kazachstania_unispora.3458/1-342 | DE | [subseq | from] | snap_masked-NODE_47_length_78712_cov_50.1529_ID_93-processed-gene-0.49-mRNA-1_1 | gene=snap_masked-NODE_47_length_78712_cov_50.1529_ID_93-processed-gene-0.49 | CDS=1-1032 |
| #=GS | kazachstania_yakushimaensis.3740/307-438 | DE | [subseq | from] | genemark-scf7180000044634-processed-gene-0.102-mRNA-1_1 | gene=genemark-scf7180000044634-processed-gene-0.102 | CDS=1-2616 |
| #=GS | kazachstania_yakushimaensis.3740/716-872 | DE | [subseq | from] | genemark-scf7180000044634-processed-gene-0.102-mRNA-1_1 | gene=genemark-scf7180000044634-processed-gene-0.102 | CDS=1-2616 |
| #=GS | kazachstania_spencerorum.3580/3-278 | DE | [subseq | from] | snap_masked-scf7180000055600-processed-gene-0.48-mRNA-1_1 | gene=snap_masked-scf7180000055600-processed-gene-0.48 | CDS=1-1224 |
| #=GS | kazachstania_spencerorum.3580/351-408 | DE | [subseq | from] | snap_masked-scf7180000055600-processed-gene-0.48-mRNA-1_1 | gene=snap_masked-scf7180000055600-processed-gene-0.48 | CDS=1-1224 |
| #=GS | kazachstania_rosinii.1249/3-239 | DE | [subseq | from] | snap_masked-scf7180000099615-processed-gene-0.31-mRNA-1_1 | gene=snap_masked-scf7180000099615-processed-gene-0.31 | CDS=1-1491 |
| #=GS | kazachstania_solicola.5780/1-137 | DE | [subseq | from] | augustus_masked-scf7180000046332-processed-gene-1.206-mRNA-1_1 | gene=augustus_masked-scf7180000046332-processed-gene-1.206 | CDS=1-1032 |
| #=GS | kazachstania_solicola.5780/187-342 | DE | [subseq | from] | augustus_masked-scf7180000046332-processed-gene-1.206-mRNA-1_1 | gene=augustus_masked-scf7180000046332-processed-gene-1.206 | CDS=1-1032 |
| #=GS | tetrapisispora_blattae.1015/1-244 | DE | [subseq | from] | snap_masked-NC_020185-processed-gene-25.62-mRNA-1_1 | gene=snap_masked-NC_020185-processed-gene-25.62 | CDS=1-1446 |
| #=GS | tetrapisispora_blattae.1015/304-465 | DE | [subseq | from] | snap_masked-NC_020185-processed-gene-25.62-mRNA-1_1 | gene=snap_masked-NC_020185-processed-gene-25.62 | CDS=1-1446 |
| #=GS | tetrapisispora_namnaonensis.1469/6-127 | DE | [subseq | from] | augustus_masked-NODE_17_length_366881_cov_78.9049_ID_33-processed-gene-2.114-mRNA-1_1 | gene=augustus_masked-NODE_17_length_366881_cov_78.9049_ID_33-processed-gene-2.114 | CDS=1-996 |
| #=GS | kluyveromyces_marxianus.2881/64-175 | DE | [subseq | from] | snap_masked-BBIL01000008.1-processed-gene-11.40-mRNA-1_1 | gene=snap_masked-BBIL01000008.1-processed-gene-11.40 | CDS=1-879 |
| #=GS | eremothecium_coryli.3485/574-629 | DE | [subseq | from] | maker-AZAH01000009.1-snap-gene-0.44-mRNA-1_1 | gene=maker-AZAH01000009.1-snap-gene-0.44 | CDS=1-2259 |
| #=GS | lachancea_nothofagi.512/203-358 | DE | [subseq | from] | snap_masked-LANO0B-processed-gene-3.66-mRNA-1_1 | gene=snap_masked-LANO0B-processed-gene-3.66 | CDS=1-1185 |

| Sequences | HMMProfileID |
|---|---|
| Saccharomyces_mikatae | snap_masked-Smik_8-processed-gene-4.133 |
| Saccharomyces_paradoxus | genemark-Spar_8-processed-gene-4.108 |
| Saccharomyces_cerevisiae | YHR195W |
| Saccharomyces_uvarum | augustus_masked-Sbay_15-processed-gene-6.183 |
| Saccharomyces_eubayanus | augustus_masked-chrXV-processed-gene-6.90 |
| Saccharomyces_arboricola | augustus_masked-NC_026178.1-processed-gene-4.111 |
| Torulaspora_delbrueckii | snap_masked-scaffold_4-processed-gene-1.34 |
| Torulaspora_franciscae | snap_masked-flattened_line_10-processed-gene-1.40 |
| Torulaspora_pretoriensis | augustus_masked-flattened_line_3-processed-gene-4.12 |
| Torulaspora_pretoriensis | augustus_masked-flattened_line_3-processed-gene-4.38 |
| Torulaspora_microellipsoides | augustus_masked-flattened_line_52-processed-gene-0.132 |
| Zygosaccharomyces_bailii | augustus_masked-ZYBA0S10-processed-gene-3.53 |
| Zygotorulaspora_florentina | augustus_masked-NODE_6_length_562868_cov_24.3835_ID_11-processed-gene-2.47 |
| Zygosaccharomyces_kombuchaensis | snap_masked-NODE_1_length_271579_cov_10.1807_ID_1-processed-gene-1.23 |
| Zygosaccharomyces_bisporus | snap_masked-flattened_line_16-processed-gene-0.27 |
| Zygotorulaspora_mrakii | snap_masked-NODE_2_length_708340_cov_18.0358_ID_2767-processed-gene-4.105 |
| Zygosaccharomyces_rouxii | snap_masked-Zyro0B-processed-gene-6.126 |
| Torulaspora_maleeae | snap_masked-NODE_5_length_764704_cov_59.617_ID_1253-processed-gene-6.109 |
| Saccharomyces_kudriavzevii | augustus_masked-Skud_8-processed-gene-3.158 |
| Lachancea_kluyveri | snap_masked-SAKL0G-processed-gene-15.78 |
| Naumovozyma_dairenensis | augustus_masked-NC_016482.1-processed-gene-7.22 |
| Naumovozyma_dairenensis | augustus_masked-NC_016482.1-processed-gene-7.22 |
| Kazachstania_viticola | snap_masked-scf7180000050361-processed-gene-0.67 |
| Naumovozyma_castellii | snap_masked-HE576752-processed-gene-12.71 |
| Lachancea_fermentati | augustus_masked-LAFE0A-processed-gene-1.131 |
| Lachancea_cidri | augustus_masked-LACI0G-processed-gene-1.33 |
| Nakaseomyces_delphensis | augustus_masked-NADE0S27-processed-gene-3.39 |
| Vanderwaltozyma_polyspora | snap_masked-NW_001834637.1-processed-gene-0.82 |
| Kluyveromyces_aestuarii | snap_masked-NODE_4_length_706832_cov_27.5555_ID_7-processed-gene-0.39 |
| Nakaseomyces_bacillisporus | augustus_masked-NABA0S29-processed-gene-0.108 |
| Nakaseomyces_bracarensis | snap_masked-CABR0S29-processed-gene-6.0 |
| Lachancea_mirantina | augustus_masked-LAMI0C-processed-gene-0.22 |
| Nakaseomyces_nivariensis | snap_masked-CANI0S17-processed-gene-4.132 |
| Lachancea_quebecensis | snap_masked-LAQU0S22-processed-gene-0.116 |
| Lachancea_waltii | snap_masked-AADM01000119.1-processed-gene-0.11 |
| Candida_glabrata | snap_masked-CR380959-processed-gene-7.12 |
| Kazachstania_bromeliacearum | snap_masked-NODE_10_length_278180_cov_22.5073_ID_19-processed-gene-1.127 |
| Nakaseomyces_castellii | snap_masked-CACA0S33-processed-gene-9.107 |
| Kazachstania_turicensis | snap_masked-scf7180000049581-processed-gene-4.63 |
| Kazachstania_naganishii | snap_masked-HE978326-processed-gene-0.118 |
| Lachancea_lanzarotensis | augustus_masked-LN736361.1-processed-gene-3.12 |
| Lachancea_dasiensis | augustus_masked-LADA0A-processed-gene-7.6 |
| Kazachstania_martiniae | snap_masked-scf7180000030514-processed-gene-0.103 |
| Lachancea_nothofagi | snap_masked-LANO0B-processed-gene-1.74 |
| Kazachstania_kunashirensis | snap_masked-scf7180000008985-processed-gene-1.61 |
| Lachancea_fantastica | snap_masked-LAFA0F-processed-gene-10.9 |
| Tetrapisispora_iriomotensis | augustus_masked-flattened_line_57-processed-gene-0.130 |
| Kazachstania_africana | augustus_masked-scaffold_2-processed-gene-13.205 |
| Kluyveromyces_lactis | snap_masked-CR382122-processed-gene-3.49 |
| Lachancea_meyersii | snap_masked-LAME0G-processed-gene-17.24 |
| Kazachstania_siamensis | augustus_masked-NODE_61_length_63804_cov_14.6622_ID_121-processed-gene-0.80 |
| Kluyveromyces_dobzhanskii | snap_masked-NODE_13_length_251415_cov_20.6991_ID_25-processed-gene-1.144 |
| Eremothecium_cymbalariae | snap_masked-NC_016453.1-processed-gene-3.121 |
| Lachancea_thermotolerans | snap_masked-CU928168-processed-gene-14.65 |
| Kazachstania_taianensis | augustus_masked-NODE_5_length_452406_cov_49.1731_ID_9-processed-gene-3.120 |
| Kazachstania_aerobia | augustus_masked-scf7180000020169-processed-gene-0.188 |
| Kazachstania_unispora | snap_masked-NODE_47_length_78712_cov_50.1529_ID_93-processed-gene-0.49 |
| Kazachstania_yakushimaensis | genemark-scf7180000044634-processed-gene-0.102 |
| Kazachstania_spencerorum | snap_masked-scf7180000055600-processed-gene-0.48 |
| Kazachstania_rosinii | snap_masked-scf7180000099615-processed-gene-0.31 |
| Kazachstania_solicola | augustus_masked-scf7180000046332-processed-gene-1.206 |
| Tetrapisispora_blattae | snap_masked-NC_020185-processed-gene-25.62 |
| Tetrapisispora_namnaonensis | augustus_masked-NODE_17_length_366881_cov_78.9049_ID_33-processed-gene-2.114 |
| Kluyveromyces_marxianus | snap_masked-BBIL01000008.1-processed-gene-11.40 |
| Eremothecium_coryli | maker-AZAH01000009.1-snap-gene-0.44 |
| Lachancea_nothofagi | snap_masked-LANO0B-processed-gene-3.66 |

| | | | | | | | | |
|---|---|---|---|---|---|---|---|---|
| >Nakaseomyces_bracarensis_g001614.m1 g001614.m1 | g001614.m1 | Nakaseomyces_bracarensis | 1 | | | | Y1000_protID | Y1000_TreeID |
| >Nakaseomyces_castellii_g001576.m1 g001576.m1 | g001576.m1 | Nakaseomyces_castellii | 1 | | | | g005344.m1 | Saccharomyces_cerevisiae |
| >Nakaseomyces_glabratus_g000308.m1 g000308.m1 | g000308.m1 | Nakaseomyces_glabratus | 1 | | | | g004292.m1 | Saccharomyces_jurei |
| >Nakaseomyces_kungkrabaensis_g004198.m1 g004198.m1 | g004198.m1 | Nakaseomyces_kungkrabaensis | 1 | | | | g005237.m1 | Saccharomyces_paradoxus |
| >Nakaseomyces_nivariensis_g001471.m1 g001471.m1 | g001471.m1 | Nakaseomyces_nivariensis | 1 | | | | g005184.m1 | Saccharomyces_kudriavzevii |
| >Nakaseomyces_uthaithaninus_g005380.m1 g005380.m1 | g005380.m1 | Nakaseomyces_uthaithaninus | 1 | | | | g005224.m1 | Saccharomyces_arboricola |
| >Eremothecium_cymbalariae_g002832.m1 g002832.m1 | g002832.m1 | Eremothecium_cymbalariae | 1 | | | | g005189.m1 | Saccharomyces_mikatae |
| >Eremothecium_sinecaudum_g003584.m1 g003584.m1 | g003584.m1 | Eremothecium_sinecaudum | 1 | | | | g002217.m1 | Saccharomyces_uvarum |
| >Hagleromyces_aurorensis_g002965.m1 g002965.m1 | g002965.m1 | Hagleromyces_aurorensis | 1 | | | | g005249.m1 | Saccharomyces_eubayanus |
| >Kazachstania_africana_g003755.m1 g003755.m1 | g003755.m1 | Kazachstania_africana | 1 | | | | g000308.m1 | Nakaseomyces_glabratus |
| >Kazachstania_aquatica_g003032.m1 g003032.m1 | g003032.m1 | #N/A | 0 | | | | g004198.m1 | Nakaseomyces_kungkrabaensis |
| >Kazachstania_bromeliacearum_g000451.m1 g000451.m1 | g000451.m1 | Kazachstania_bromeliacearum | 1 | | | | g001471.m1 | Nakaseomyces_nivariensis |
| >Kazachstania_bulderi_g005958.m1 g005958.m1 | g005958.m1 | Kazachstania_bulderi | 1 | | | | g005380.m1 | Nakaseomyces_uthaithaninus |
| >Kazachstania_exigua_g000048.m1 g000048.m1 | g000048.m1 | #N/A | 0 | | | | g004209.m1 | Nakaseomyces_delphensis |
| >Kazachstania_exigua_g002519.m1 g002519.m1 | g002519.m1 | Kazachstania_exigua | 1 | | 1 | g001614.m1 | g001614.m1 | Nakaseomyces_bracarensis |
| >Kazachstania_gamospora_g004167.m1 g004167.m1 | g004167.m1 | Kazachstania_gamospora | 1 | | | | g004314.m1 | Nakaseomyces_bacillisporus |
| >Kazachstania_hellenica_g003117.m1 g003117.m1 | g003117.m1 | Kazachstania_hellenica | 1 | | | | g001576.m1 | Nakaseomyces_castellii |
| >Kazachstania_humilis_NRRL_Y-7245_g000693.m1 g000693.m1 | g000693.m1 | #N/A | 0 | | | | g005068.m1 | Kazachstania_solicola |
| >Kazachstania_kunashirensis_g005050.m1 g005050.m1 | g005050.m1 | Kazachstania_kunashirensis | 2 | | | | g002718.m1 | Kazachstania_aerobia |
| >Kazachstania_lodderae_g003605.m1 g003605.m1 | g003605.m1 | Kazachstania_lodderae | 1 | | | | g003649.m1 | Kazachstania_unispora |
| >Kazachstania_martiniae_g004439.m1 g004439.m1 | g004439.m1 | Kazachstania_martiniae | 1 | | | | g000451.m1 | Kazachstania_bromeliacearum |
| >Kazachstania_naganishii_g001440.m1 g001440.m1 | g001440.m1 | Kazachstania_naganishii | 1 | | | | g005958.m1 | Kazachstania_bulderi |
| >Kazachstania_piceae_g005225.m1 g005225.m1 | g005225.m1 | Kazachstania_piceae | 1 | | | | g002519.m1 | Kazachstania_exigua |
| >Kazachstania_pseudohumilis_g002365.m1 g002365.m1 | g002365.m1 | #N/A | 0 | | | | g004167.m1 | Kazachstania_gamospora |
| >Kazachstania_rosinii_g001213.m1 g001213.m1 | g001213.m1 | #N/A | 0 | | | | g003117.m1 | Kazachstania_hellenica |
| >Kazachstania_saulgeensis_g000550.m1 g000550.m1 | g000550.m1 | Kazachstania_saulgeensis | 1 | | | | g005050.m1 | Kazachstania_kunashirensis |
| >Kazachstania_servazzii_g000773.m1 g000773.m1 | g000773.m1 | Kazachstania_servazzii | 1 | | | | g003605.m1 | Kazachstania_lodderae |
| >Kazachstania_siamensis_g004344.m1 g004344.m1 | g004344.m1 | Kazachstania_siamensis | 1 | | | | g005225.m1 | Kazachstania_piceae |
| >Kazachstania_sinensis_g005599.m1 g005599.m1 | g005599.m1 | Kazachstania_sinensis | 1 | | | | g000550.m1 | Kazachstania_saulgeensis |
| >Kazachstania_sp._UFMG-CM-Y273_g000466.m1 g000466.m1 | g000466.m1 | #N/A | 0 | | | | g000773.m1 | Kazachstania_servazzii |
| >Kazachstania_taianensis_g004007.m1 g004007.m1 | g004007.m1 | Kazachstania_taianensis | 1 | | | | g005599.m1 | Kazachstania_sinensis |
| >Kazachstania_turicensis_g004713.m1 g004713.m1 | g004713.m1 | Kazachstania_turicensis | 1 | | | | g001209.m1 | Kazachstania_yasuniensis |
| >Kazachstania_viticola_g001183.m1 g001183.m1 | g001183.m1 | Kazachstania_viticola | 1 | | | | g001263.m1 | Kazachstania_barnettii |
| >Kazachstania_yasuniensis_g001209.m1 g001209.m1 | g001209.m1 | Kazachstania_yasuniensis | 1 | | | | g002942.m1 | Kazachstania_psychrophila |
| >Kluyveromyces_aestuarii_ATCC_18862_g000823.m1 g000823.m1 | g000823.m1 | Kluyveromyces_aestuarii_ATCC_18862 | 1 | | | | g001801.m1 | Kazachstania_jinghongensis |
| >Kluyveromyces_aestuarii_NRRL_YB-4510_g003160.m1 g003160.m1 | g003160.m1 | #N/A | 0 | | | | g004344.m1 | Kazachstania_siamensis |
| >Kluyveromyces_dobzhanskii_g001256.m1 g001256.m1 | g001256.m1 | Kluyveromyces_dobzhanskii | 1 | | | | g004007.m1 | Kazachstania_taianensis |
| >Kluyveromyces_lactis_CBS_2359_g000677.m1 g000677.m1 | g000677.m1 | Kluyveromyces_lactis_CBS_2359 | 1 | | | | g001440.m1 | Kazachstania_naganishii |
| >Kluyveromyces_lactis_var._drosophilarum_NRRL_Y-8278_g001346.m1 g001346.m1 | g001346.m1 | #N/A | 0 | | | | g004439.m1 | Kazachstania_martiniae |
| >Kluyveromyces_lactis_var._lactis_NRRL_Y-8279_g002882.m1 g002882.m1 | g002882.m1 | #N/A | 0 | | | | g004713.m1 | Kazachstania_turicensis |
| >Kluyveromyces_nonfermentans_g001343.m1 g001343.m1 | g001343.m1 | Kluyveromyces_nonfermentans | 1 | | | | g005050.m1 | Kazachstania_kunashirensis |
| >Kluyveromyces_siamensis_g003460.m1 g003460.m1 | g003460.m1 | Kluyveromyces_siamensis | 1 | | | | g002874.m1 | Kazachstania_spencerorum |
| >Kluyveromyces_sp._yHMH660_g005129.m1 g005129.m1 | g005129.m1 | #N/A | 0 | | | | g003755.m1 | Kazachstania_africana |
| >Kluyveromyces_starmeri_g003161.m1 g003161.m1 | g003161.m1 | Kluyveromyces_starmeri | 1 | | | | g001183.m1 | Kazachstania_viticola |
| >Kluyveromyces_wickerhamii_g000532.m1 g000532.m1 | g000532.m1 | Kluyveromyces_wickerhamii | 1 | | | | g002752.m1 | Naumovozyma_dairenensis |
| >Lachancea_cidri_NRRL_Y-12635_g002916.m1 g002916.m1 | g002916.m1 | #N/A | 0 | | | | g001098.m1 | Naumovozyma_castellii |
| >Lachancea_cidri_NRRL_Y-12634_g003378.m1 g003378.m1 | g003378.m1 | #N/A | 0 | | | | g000794.m1 | Naumovozyma_baii |
| >Lachancea_dasiensis_g003259.m1 g003259.m1 | g003259.m1 | Lachancea_dasiensis | 1 | | | | g003416.m1 | Tetrapisispora_iriomotensis |
| >Lachancea_fantastica_nom._nud._g003435.m1 g003435.m1 | g003435.m1 | Lachancea_fantastica_nom._nud. | 1 | | | | g001993.m1 | Tetrapisispora_namnaoensis |
| >Lachancea_fermentati_g004518.m1 g004518.m1 | g004518.m1 | Lachancea_fermentati | 1 | | | | g003018.m1 | Tetrapisispora_blattae |
| >Lachancea_kluyveri_g001117.m1 g001117.m1 | g001117.m1 | Lachancea_kluyveri | 1 | | | | g005365.m1 | Vanderwaltozyma_polyspora |
| >Lachancea_meyersii_g001214.m1 g001214.m1 | g001214.m1 | Lachancea_meyersii | 1 | | | | g001449.m1 | Vanderwaltozyma_yarrowii |
| >Lachancea_mirantina_g001548.m1 g001548.m1 | g001548.m1 | Lachancea_mirantina | 1 | | | | g003717.m1 | Vanderwaltozyma_tropicalis |
| >Lachancea_nothofagi_g001406.m1 g001406.m1 | g001406.m1 | Lachancea_nothofagi | 1 | | | | g005143.m1 | Vanderwaltozyma_verrucispora |
| >Lachancea_quebecensis_g002616.m1 g002616.m1 | g002616.m1 | Lachancea_quebecensis | 1 | | | | g004938.m1 | Torulaspora_pretoriensis |
| >Lachancea_sp._yHQL494_g002832.m1 g002832.m1 | g002832.m1 | Eremothecium_cymbalariae | 1 | | | | g002639.m1 | Torulaspora_microellipsoides |
| >Lachancea_thermotolerans_g003213.m1 g003213.m1 | g003213.m1 | Lachancea_thermotolerans | 1 | | | | g004752.m1 | Torulaspora_franciscae |
| >Lachancea_waltii_g003199.m1 g003199.m1 | g003199.m1 | Lachancea_waltii | 1 | | | | g001776.m1 | Torulaspora_delbrueckii |
| >Kazachstania_barnettii_g001263.m1 g001263.m1 | g001263.m1 | Kazachstania_barnettii | 1 | | | | g001826.m1 | Torulaspora_globosa |
| >Torulaspora_franciscae_g004752.m1 g004752.m1 | g004752.m1 | Torulaspora_franciscae | 1 | | | | g003867.m1 | Torulaspora_maleeae |
| >Lachancea_lanzarotensis_g004947.m1 g004947.m1 | g004947.m1 | Lachancea_lanzarotensis | 1 | | | | g005101.m1 | Zygosaccharomyces_bisporus |
| >Kazachstania_psychrophila_g002942.m1 g002942.m1 | g002942.m1 | Kazachstania_psychrophila | 1 | | | | g005022.m1 | Zygosaccharomyces_gambellarensis |
| >Vanderwaltozyma_yarrowii_g001449.m1 g001449.m1 | g001449.m1 | Vanderwaltozyma_yarrowii | 1 | | | | g001048.m1 | Zygosaccharomyces_bailii |
| >Tetrapisispora_iriomotensis_g003416.m1 g003416.m1 | g003416.m1 | Tetrapisispora_iriomotensis | 1 | | | | g000063.m1 | Zygosaccharomyces_kombuchaensis |
| >Torulaspora_pretoriensis_g004938.m1 g004938.m1 | g004938.m1 | Torulaspora_pretoriensis | 1 | | | | g004388.m1 | Zygosaccharomyces_lentus |
| >Torulaspora_microellipsoides_g002639.m1 g002639.m1 | g002639.m1 | Torulaspora_microellipsoides | 1 | | | | g002135.m1 | Zygosaccharomyces_mellis |
| >Kazachstania_jinghongensis_g001801.m1 g001801.m1 | g001801.m1 | Kazachstania_jinghongensis | 1 | | | | g004410.m1 | Zygosaccharomyces_pseudorouxii |
| >Vanderwaltozyma_verrucispora_g005143.m1 g005143.m1 | g005143.m1 | Vanderwaltozyma_verrucispora | 1 | | | | g000372.m1 | Zygosaccharomyces_siamensis |
| >Nakaseomyces_bacillisporus_g004314.m1 g004314.m1 | g004314.m1 | Nakaseomyces_bacillisporus | 1 | | | | g004327.m1 | Zygotorulaspora_florentina |
| >Nakaseomyces_delphensis_g004209.m1 g004209.m1 | g004209.m1 | Nakaseomyces_delphensis | 1 | | | | g002003.m1 | Zygotorulaspora_mrakii |
| >Nakaseomyces_sp._yHDO568_g004296.m1 g004296.m1 | g004296.m1 | #N/A | 0 | | | | g002973.m1 | Zygotorulaspora_chibaensis |
| >Naumovozyma_baii_g000794.m1 g000794.m1 | g000794.m1 | Naumovozyma_baii | 1 | | | | g002121.m1 | Zygotorulaspora_danielsina |
| >Naumovozyma_castellii_g001098.m1 g001098.m1 | g001098.m1 | Naumovozyma_castellii | 1 | | | | g002003.m1 | Zygotorulaspora_mrakii |
| >Naumovozyma_dairenensis_g002752.m1 g002752.m1 | g002752.m1 | Naumovozyma_dairenensis | 1 | | | | g002965.m1 | Hagleromyces_aurorensis |

| | | | | | | | | |
|---|---|---|---|---|---|---|---|---|
| >Saccharomyces_arboricola_g005224.m1 g005224.m1 | g005224.m1 | Saccharomyces_arboricola | 1 | | | | *g001117.m1* | *Lachancea_kluyveri* |
| >Saccharomyces_cerevisiae_g005344.m1 g005344.m1 | g005344.m1 | Saccharomyces_cerevisiae | 1 | | | | *g001548.m1* | *Lachancea_mirantina* |
| >Saccharomyces_eubayanus_g005249.m1 g005249.m1 | g005249.m1 | Saccharomyces_eubayanus | 1 | | | | *g001406.m1* | *Lachancea_nothofagi* |
| >Saccharomyces_jurei_g004292.m1 g004292.m1 | g004292.m1 | Saccharomyces_jurei | 1 | | | | *g003435.m1* | Lachancea_fantastica_nom._nud. |
| >Saccharomyces_kudriavzevii_g005184.m1 g005184.m1 | g005184.m1 | Saccharomyces_kudriavzevii | 1 | | | | *g004947.m1* | *Lachancea_lanzarotensis* |
| >Saccharomyces_mikatae_g005189.m1 g005189.m1 | g005189.m1 | Saccharomyces_mikatae | 1 | | | | *g003199.m1* | *Lachancea_waltii* |
| >Saccharomyces_paradoxus_g005237.m1 g005237.m1 | g005237.m1 | Saccharomyces_paradoxus | 1 | | | | g002616.m1 | *Lachancea_quebecensis* |
| >Saccharomyces_uvarum_g002217.m1 g002217.m1 | g002217.m1 | Saccharomyces_uvarum | 1 | | | | *g003213.m1* | *Lachancea_thermotolerans* |
| >Torulaspora_delbrueckii_g001776.m1 g001776.m1 | g001776.m1 | Torulaspora_delbrueckii | 1 | | | | *g001214.m1* | *Lachancea_meyersii* |
| >Torulaspora_globosa_g001826.m1 g001826.m1 | g001826.m1 | Torulaspora_globosa | 1 | | | | *g003259.m1* | *Lachancea_dasiensis* |
| >Torulaspora_maleeae_g003867.m1 g003867.m1 | g003867.m1 | Torulaspora_maleeae | 1 | | | | g004518.m1 | *Lachancea_fermentati* |
| >Torulaspora_sp._yHMJ407_g002906.m1 g002906.m1 | g002906.m1 | #N/A | 0 | | | | *NA* | Eremothecium_gossypii |
| >Vanderwaltozyma_polyspora_g005365.m1 g005365.m1 | g005365.m1 | Vanderwaltozyma_polyspora | 1 | | | | *g002832.m1* | *Eremothecium_cymbalariae* |
| >Vanderwaltozyma_tropicalis_g003717.m1 g003717.m1 | g003717.m1 | Vanderwaltozyma_tropicalis | 2 | | | | *g004362.m1* | *Eremothecium_coryli* |
| >Zygosaccharomyces_bailii_g001048.m1 g001048.m1 | g001048.m1 | Zygosaccharomyces_bailii | 1 | | | | *g003584.m1* | *Eremothecium_sinecaudum* |
| >Zygosaccharomyces_bisporus_g005101.m1 g005101.m1 | g005101.m1 | Zygosaccharomyces_bisporus | 1 | | | | g000677.m1 | Kluyveromyces_lactis_CBS_2359 |
| >Zygosaccharomyces_gambellarensis_g005022.m1 g005022.m1 | g005022.m1 | Zygosaccharomyces_gambellarensis | 1 | | | | *g001256.m1* | *Kluyveromyces_dobzhanskii* |
| >Zygosaccharomyces_kombuchaensis_g000063.m1 g000063.m1 | g000063.m1 | Zygosaccharomyces_kombuchaensis | 1 | | | | g000823.m1 | Kluyveromyces_aestuarii_ATCC_18862 |
| >Zygosaccharomyces_lentus_g004388.m1 g004388.m1 | g004388.m1 | Zygosaccharomyces_lentus | 1 | | | | *g001343.m1* | *Kluyveromyces_nonfermentans* |
| >Zygosaccharomyces_mellis_g002135.m1 g002135.m1 | g002135.m1 | Zygosaccharomyces_mellis | 1 | | | | *g003460.m1* | *Kluyveromyces_siamensis* |
| >Zygosaccharomyces_parabailii_g003412.m1 g003412.m1 | g003412.m1 | #N/A | 0 | | | | *g003161.m1* | *Kluyveromyces_starmeri* |
| >Zygosaccharomyces_pseudobailii_g001287.m1 g001287.m1 | g001287.m1 | #N/A | 0 | | | | *g000532.m1* | *Kluyveromyces_wickerhamii* |
| >Zygosaccharomyces_pseudobailii_g002625.m1 g002625.m1 | g002625.m1 | #N/A | 0 | | | | *g000873.m1* | Hanseniaspora_uvarum_DSMZ_2768 |
| >Zygosaccharomyces_pseudobailii_g003112.m1 g003112.m1 | g003112.m1 | #N/A | 0 | | | | *g000765.m1* | Hanseniaspora_pseudoguilliermondii |
| >Zygosaccharomyces_pseudorouxii_g004410.m1 g004410.m1 | g004410.m1 | Zygosaccharomyces_pseudorouxii | 1 | | | | *g003387.m1* | *Hanseniaspora_clermontiae* |
| >Zygosaccharomyces_rouxii_g001572.m1 g001572.m1 | g001572.m1 | #N/A | 0 | | | | g000788.m1 | *Hanseniaspora_valbyensis* |
| >Zygosaccharomyces_sapae_g000017.m1 g000017.m1 | g000017.m1 | #N/A | 0 | | | | g000163.m1 | Hanseniaspora_vineae |
| >Zygosaccharomyces_sapae_g006118.m1 g006118.m1 | g006118.m1 | #N/A | 0 | | | | *g005674.m1* | *Cyberlindnera_suaveolens* |
| >Zygosaccharomyces_siamensis_g000372.m1 g000372.m1 | g000372.m1 | Zygosaccharomyces_siamensis | 1 | | | | *g001009.m1* | *Cyberlindnera_saturnus* |
| >Zygotorulaspora_chibaensis_g002973.m1 g002973.m1 | g002973.m1 | Zygotorulaspora_chibaensis | 1 | | | | *g002417.m1* | *Cyberlindnera_misumaiensis* |
| >Zygotorulaspora_danielsina_g002121.m1 g002121.m1 | g002121.m1 | Zygotorulaspora_danielsina | 1 | | | | *g002673.m1* | *Cyberlindnera_jadinii* |
| >Zygotorulaspora_florentina_g004327.m1 g004327.m1 | g004327.m1 | Zygotorulaspora_florentina | 1 | | | | *g005778.m1* | *Cyberlindnera_fabianii* |
| >Zygotorulaspora_mrakii_g002003.m1 g002003.m1 | g002003.m1 | Zygotorulaspora_mrakii | 2 | | | | *g003717.m1* | *Cyberlindnera_maclurae* |
| >Zygotorulaspora_sp._yHDO592_g004529.m1 g004529.m1 | g004529.m1 | #N/A | 0 | | | | *g001244.m1* | Wickerhamomyces_alni |
| | | | | | | | *g004559.m1* | *Wickerhamomyces_canadensis* |
| | | | | | | | g005482.m1,g003054.m1 | *Wickerhamomyces_ciferrii* |
| | | | | | | | *g001027.m1* | *Wickerhamomyces_chambardii* |
| | | | | | | | *g002138.m1* | *Wickerhamomyces_hampshirensis* |
| | | | | | | | *g002404.m1* | *Wickerhamomyces_mucosus* |
| | | | | | | | g002584.m1 | Wickerhamomyces_pijperi |
| | | | | | | | g002262.m1 | Komagataella_phaffii_GS115 |
| | | | | | | | *g004657.m1* | *Komagataella_populi* |
| | | | | | | | g002679.m1 | Komagataella_pastoris_ATCC_28485 |
| | | | | | | | *g004231.m1* | *Komagataella_kurtzmanii* |
| | | | | | | | *g004487.m1* | Komagataella_mondaviorum |
| | | | | | | | *g004700.m1* | *Kuraishia_floccosa* |
| | | | | | | | NA | Ashbya_aceri |
| | | | | | | | g001657.m1 | Wickerhamomyces_anomalus |

| Species | gID | evalue | score | fastaID | countInTableS1 | | Y1000_protID_TableS1 | count_in_hmm_results | Present_in_hmm_results |
|---|---|---|---|---|---|---|---|---|---|
| Nakaseomyces_bracarensis | g001614.m1 | 1.00E-23 | 82.7 | >Nakaseomyces_bracarensis_g001614.m1 g001614.m1 | 1 | | *g005344.m1* | 1 | 1 |
| Nakaseomyces_castellii | g001576.m1 | 8.20E-21 | 73.1 | >Nakaseomyces_castellii_g001576.m1 g001576.m1 | 1 | | *g004292.m1* | 1 | 1 |
| Nakaseomyces_glabratus | g000308.m1 | 2.00E-21 | 75.2 | >Nakaseomyces_glabratus_g000308.m1 g000308.m1 | 1 | | *g005237.m1* | 1 | 1 |
| Nakaseomyces_kungkrabaensis | g004198.m1 | 6.30E-23 | 80.1 | >Nakaseomyces_kungkrabaensis_g004198.m1 g004198.m1 | 1 | | *g005184.m1* | 1 | 1 |
| Nakaseomyces_nivariensis | g001471.m1 | 8.60E-23 | 79.6 | >Nakaseomyces_nivariensis_g001471.m1 g001471.m1 | 1 | | *g005224.m1* | 1 | 1 |
| Nakaseomyces_uthaithaninus | g005380.m1 | 1.30E-22 | 79.1 | >Nakaseomyces_uthaithaninus_g005380.m1 g005380.m1 | 1 | | *g005189.m1* | 1 | 1 |
| Eremothecium_cymbalariae | g002832.m1 | 5.50E-15 | 53.8 | >Eremothecium_cymbalariae_g002832.m1 g002832.m1 | 1 | | *g002217.m1* | 1 | 1 |
| Eremothecium_sinecaudum | g003584.m1 | 8.20E-17 | 59.8 | >Eremothecium_sinecaudum_g003584.m1 g003584.m1 | 1 | | *g005249.m1* | 1 | 1 |
| Hagleromyces_aurorensis | g002965.m1 | 4.10E-38 | 129.9 | >Hagleromyces_aurorensis_g002965.m1 g002965.m1 | 1 | | *g000308.m1* | 1 | 1 |
| Kazachstania_africana | g003755.m1 | 1.40E-17 | 62.6 | >Kazachstania_africana_g003755.m1 g003755.m1 | 1 | | *g004198.m1* | 1 | 1 |
| Kazachstania_aquatica | g003032.m1 | 5.10E-20 | 70.7 | >Kazachstania_aquatica_g003032.m1 g003032.m1 | 1 | | *g001471.m1* | 1 | 1 |
| Kazachstania_bromeliacearum | g000451.m1 | 6.40E-21 | 73.6 | >Kazachstania_bromeliacearum_g000451.m1 g000451.m1 | 1 | | *g005380.m1* | 1 | 1 |
| Kazachstania_bulderi | g005958.m1 | 2.50E-16 | 58.6 | >Kazachstania_bulderi_g005958.m1 g005958.m1 | 1 | | *g004209.m1* | 1 | 1 |
| Kazachstania_exigua | g002519.m1 | 5.00E-17 | 61.7 | >Kazachstania_exigua_g002519.m1 g002519.m1 | 1 | | *g001614.m1* | 1 | 1 |
| Kazachstania_exigua | g000048.m1 | 1.50E-14 | 53.6 | >Kazachstania_exigua_g000048.m1 g000048.m1 | 0 | | *g004314.m1* | 1 | 1 |
| Kazachstania_gamospora | g004167.m1 | 3.00E-17 | 61.5 | >Kazachstania_gamospora_g004167.m1 g004167.m1 | 1 | | *g001576.m1* | 1 | 1 |
| Kazachstania_hellenica | g003117.m1 | 7.90E-20 | 69.9 | >Kazachstania_hellenica_g003117.m1 g003117.m1 | 1 | | *g005068.m1* | 0 | 0 |
| Kazachstania_humilis_NRRL_Y-7245 | g000693.m1 | 2.10E-17 | 62.3 | >Kazachstania_humilis_NRRL_Y-7245_g000693.m1 g000693.m1 | 0 | | *g002718.m1* | 0 | 0 |
| Kazachstania_kunashirensis | g005050.m1 | 9.30E-19 | 66.5 | >Kazachstania_kunashirensis_g005050.m1 g005050.m1 | 2 | | *g003649.m1* | 0 | 0 |
| Kazachstania_lodderae | g003605.m1 | 4.30E-19 | 67.6 | >Kazachstania_lodderae_g003605.m1 g003605.m1 | 1 | | *g000451.m1* | 1 | 1 |
| Kazachstania_martiniae | g004439.m1 | 5.20E-18 | 64.1 | >Kazachstania_martiniae_g004439.m1 g004439.m1 | 1 | | *g005958.m1* | 1 | 1 |
| Kazachstania_naganishii | g001440.m1 | 1.20E-19 | 69.3 | >Kazachstania_naganishii_g001440.m1 g001440.m1 | 1 | | *g002519.m1* | 1 | 1 |
| Kazachstania_piceae | g005225.m1 | 5.50E-18 | 64 | >Kazachstania_piceae_g005225.m1 g005225.m1 | 1 | | *g004167.m1* | 1 | 1 |
| Kazachstania_pseudohumilis | g002365.m1 | 1.00E-16 | 59.9 | >Kazachstania_pseudohumilis_g002365.m1 g002365.m1 | 0 | | *g003117.m1* | 1 | 1 |
| Kazachstania_rosinii | g001213.m1 | 3.70E-20 | 71.1 | >Kazachstania_rosinii_g001213.m1 g001213.m1 | 0 | | g003032.m1 | 1 | 1 |
| Kazachstania_saulgeensis | g000550.m1 | 2.80E-18 | 64.9 | >Kazachstania_saulgeensis_g000550.m1 g000550.m1 | 1 | | *g005050.m1* | 1 | 1 |
| Kazachstania_servazzii | g000773.m1 | 6.60E-17 | 60.4 | >Kazachstania_servazzii_g000773.m1 g000773.m1 | 1 | | *g003605.m1* | 1 | 1 |
| Kazachstania_siamensis | g004344.m1 | 4.50E-17 | 60.9 | >Kazachstania_siamensis_g004344.m1 g004344.m1 | 1 | | *g005225.m1* | 1 | 1 |
| Kazachstania_sinensis | g005599.m1 | 3.60E-21 | 74.4 | >Kazachstania_sinensis_g005599.m1 g005599.m1 | 1 | | *g000550.m1* | 1 | 1 |
| Kazachstania_sp._UFMG-CM-Y273 | g000466.m1 | 2.20E-18 | 65.2 | >Kazachstania_sp._UFMG-CM-Y273_g000466.m1 g000466.m1 | 0 | | *g000773.m1* | 1 | 1 |
| Kazachstania_taianensis | g004007.m1 | 3.30E-14 | 51.5 | >Kazachstania_taianensis_g004007.m1 g004007.m1 | 1 | | *g005599.m1* | 1 | 1 |
| Kazachstania_turicensis | g004713.m1 | 5.80E-20 | 70.4 | >Kazachstania_turicensis_g004713.m1 g004713.m1 | 1 | | *g001209.m1* | 1 | 1 |
| Kazachstania_viticola | g001183.m1 | 3.00E-34 | 117.3 | >Kazachstania_viticola_g001183.m1 g001183.m1 | 1 | | *g001263.m1* | 1 | 1 |
| Kazachstania_yasuniensis | g001209.m1 | 2.90E-15 | 55 | >Kazachstania_yasuniensis_g001209.m1 g001209.m1 | 1 | | *g002942.m1* | 1 | 1 |
| Kluyveromyces_aestuarii_ATCC_18862 | g000823.m1 | 1.00E-24 | 85.9 | >Kluyveromyces_aestuarii_ATCC_18862_g000823.m1 g000823.m1 | 1 | | *g001801.m1* | 1 | 1 |
| Kluyveromyces_aestuarii_NRRL_YB-4510 | g003160.m1 | 1.00E-24 | 85.9 | >Kluyveromyces_aestuarii_NRRL_YB-4510_g003160.m1 g003160.m1 | 0 | | *g004344.m1* | 1 | 1 |
| Kluyveromyces_dobzhanskii | g001256.m1 | 2.00E-15 | 55.4 | >Kluyveromyces_dobzhanskii_g001256.m1 g001256.m1 | 1 | | *g004007.m1* | 1 | 1 |
| Kluyveromyces_lactis_CBS_2359 | g000677.m1 | 2.70E-17 | 61.5 | >Kluyveromyces_lactis_CBS_2359_g000677.m1 g000677.m1 | 1 | | *g001440.m1* | 1 | 1 |
| Kluyveromyces_lactis_var._drosophilarum_NRRL_Y-8278 | g001346.m1 | 6.30E-18 | 63.6 | >Kluyveromyces_lactis_var._drosophilarum_NRRL_Y-8278_g001346.m1 g001346.m1 | 0 | | g004439.m1 | 1 | 1 |
| Kluyveromyces_lactis_var._lactis_NRRL_Y-8279 | g002882.m1 | 2.70E-17 | 61.5 | >Kluyveromyces_lactis_var._lactis_NRRL_Y-8279_g002882.m1 g002882.m1 | 0 | | *g004713.m1* | 1 | 1 |
| Kluyveromyces_nonfermentans | g001343.m1 | 1.10E-21 | 75.9 | >Kluyveromyces_nonfermentans_g001343.m1 g001343.m1 | 1 | | *g005050.m1* | 1 | 1 |
| Kluyveromyces_siamensis | g003460.m1 | 1.50E-25 | 88.7 | >Kluyveromyces_siamensis_g003460.m1 g003460.m1 | 1 | | *g002874.m1* | 0 | 0 |
| Kluyveromyces_sp._yHMH660 | g005129.m1 | 8.70E-14 | 50.1 | >Kluyveromyces_sp._yHMH660_g005129.m1 g005129.m1 | 0 | | *g003755.m1* | 1 | 1 |
| Kluyveromyces_starmeri | g003161.m1 | 5.50E-15 | 53.9 | >Kluyveromyces_starmeri_g003161.m1 g003161.m1 | 1 | | *g001183.m1* | 1 | 1 |
| Kluyveromyces_wickerhamii | g000532.m1 | 1.90E-18 | 65.3 | >Kluyveromyces_wickerhamii_g000532.m1 g000532.m1 | 1 | | g002752.m1 | 1 | 1 |
| Lachancea_cidri_NRRL_Y-12635 | g002916.m1 | 6.50E-29 | 99.7 | >Lachancea_cidri_NRRL_Y-12635_g002916.m1 g002916.m1 | 0 | | g001098.m1 | 1 | 1 |
| Lachancea_cidri_NRRL_Y-12634 | g003378.m1 | 6.60E-29 | 99.7 | >Lachancea_cidri_NRRL_Y-12634_g003378.m1 g003378.m1 | 0 | | *g000794.m1* | 1 | 1 |
| Lachancea_dasiensis | g003259.m1 | 7.30E-19 | 66.7 | >Lachancea_dasiensis_g003259.m1 g003259.m1 | 1 | | *g003416.m1* | 1 | 1 |
| Lachancea_fantastica_nom._nud. | g003435.m1 | 9.60E-19 | 66.3 | >Lachancea_fantastica_nom._nud._g003435.m1 g003435.m1 | 1 | | *g001993.m1* | 0 | 0 |
| Lachancea_fermentati | g004518.m1 | 2.30E-29 | 101.2 | >Lachancea_fermentati_g004518.m1 g004518.m1 | 1 | | *g003018.m1* | 0 | 0 |
| Lachancea_kluyveri | g001117.m1 | 5.90E-82 | 274.1 | >Lachancea_kluyveri_g001117.m1 g001117.m1 | 1 | | *g005365.m1* | 1 | 1 |
| Lachancea_meyersii | g001214.m1 | 1.20E-18 | 66 | >Lachancea_meyersii_g001214.m1 g001214.m1 | 1 | | *g001449.m1* | 1 | 1 |
| Lachancea_mirantina | g001548.m1 | 3.50E-23 | 80.9 | >Lachancea_mirantina_g001548.m1 g001548.m1 | 1 | | *g003717.m1* | 1 | 1 |
| Lachancea_nothofagi | g001406.m1 | 5.90E-24 | 83.4 | >Lachancea_nothofagi_g001406.m1 g001406.m1 | 1 | | *g005143.m1* | 1 | 1 |
| Lachancea_quebecensis | g002616.m1 | 4.00E-28 | 97.1 | >Lachancea_quebecensis_g002616.m1 g002616.m1 | 1 | | *g004938.m1* | 1 | 1 |
| Lachancea_sp._yHQL494 | g002832.m1 | 5.20E-29 | 100 | >Lachancea_sp._yHQL494_g002832.m1 g002832.m1 | 1 | | *g002639.m1* | 1 | 1 |
| Lachancea_thermotolerans | g003213.m1 | 3.60E-28 | 97.3 | >Lachancea_thermotolerans_g003213.m1 g003213.m1 | 1 | | *g004752.m1* | 1 | 1 |
| Lachancea_waltii | g003199.m1 | 5.20E-25 | 86.9 | >Lachancea_waltii_g003199.m1 g003199.m1 | 1 | | *g001776.m1* | 1 | 1 |
| Kazachstania_barnettii | g001263.m1 | 3.00E-18 | 64.8 | >Kazachstania_barnettii_g001263.m1 g001263.m1 | 1 | | *g001826.m1* | 1 | 1 |
| Torulaspora_franciscae | g004752.m1 | 1.30E-107 | 358.3 | >Torulaspora_franciscae_g004752.m1 g004752.m1 | 1 | | *g003867.m1* | 1 | 1 |
| Lachancea_lanzarotensis | g004947.m1 | 1.30E-19 | 69.2 | >Lachancea_lanzarotensis_g004947.m1 g004947.m1 | 1 | | *g005101.m1* | 1 | 1 |
| Kazachstania_psychrophila | g002942.m1 | 5.50E-19 | 67.2 | >Kazachstania_psychrophila_g002942.m1 g002942.m1 | 1 | | *g005022.m1* | 1 | 1 |
| Vanderwaltozyma_yarrowii | g001449.m1 | 1.90E-27 | 95 | >Vanderwaltozyma_yarrowii_g001449.m1 g001449.m1 | 1 | | g001048.m1 | 1 | 1 |
| Tetrapisispora_iriomotensis | g003416.m1 | 2.40E-18 | 65 | >Tetrapisispora_iriomotensis_g003416.m1 g003416.m1 | 1 | | *g000063.m1* | 1 | 1 |
| Torulaspora_pretoriensis | g004938.m1 | 6.40E-105 | 349.4 | >Torulaspora_pretoriensis_g004938.m1 g004938.m1 | 1 | | *g004388.m1* | 1 | 1 |
| Torulaspora_microellipsoides | g002639.m1 | 1.00E-101 | 339 | >Torulaspora_microellipsoides_g002639.m1 g002639.m1 | 1 | | *g002135.m1* | 1 | 1 |
| Kazachstania_jinghongensis | g001801.m1 | 1.20E-15 | 56.2 | >Kazachstania_jinghongensis_g001801.m1 g001801.m1 | 1 | | *g004410.m1* | 1 | 1 |
| Vanderwaltozyma_verrucispora | g005143.m1 | 2.30E-33 | 114.5 | >Vanderwaltozyma_verrucispora_g005143.m1 g005143.m1 | 1 | | *g000372.m1* | 1 | 1 |
| Nakaseomyces_bacillisporus | g004314.m1 | 1.50E-24 | 85.4 | >Nakaseomyces_bacillisporus_g004314.m1 g004314.m1 | 1 | | *g004327.m1* | 1 | 1 |
| Nakaseomyces_delphensis | g004209.m1 | 1.70E-27 | 95.1 | >Nakaseomyces_delphensis_g004209.m1 g004209.m1 | 1 | | g002003.m1 | 1 | 1 |

| Species | gID | evalue | score | fastaID | countInTableS1 | | Y1000_protID_TableS1 | count_in_hmm_results | Present_in_hmm_results |
|---|---|---|---|---|---|---|---|---|---|
| Nakaseomyces_sp._yHDO568 | g004296.m1 | 1.30E-24 | 85.7 | >Nakaseomyces_sp._yHDO568_g004296.m1 g004296.m1 | 0 | | *g002973.m1* | 1 | 1 |
| Naumovozyma_baii | g000794.m1 | 3.60E-30 | 104 | >Naumovozyma_baii_g000794.m1 g000794.m1 | 1 | | *g002121.m1* | 1 | 1 |
| Naumovozyma_castellii | g001098.m1 | 5.40E-47 | 159.3 | >Naumovozyma_castellii_g001098.m1 g001098.m1 | 1 | | g002003.m1 | 1 | 1 |
| Naumovozyma_dairenensis | g002752.m1 | 1.70E-51 | 174.1 | >Naumovozyma_dairenensis_g002752.m1 g002752.m1 | 1 | | *g002965.m1* | 1 | 1 |
| Saccharomyces_arboricola | g005224.m1 | 1.60E-116 | 387.7 | >Saccharomyces_arboricola_g005224.m1 g005224.m1 | 1 | | *g001117.m1* | 1 | 1 |
| Saccharomyces_cerevisiae | g005344.m1 | 2.70E-131 | 436.2 | >Saccharomyces_cerevisiae_g005344.m1 g005344.m1 | 1 | | *g001548.m1* | 1 | 1 |
| Saccharomyces_eubayanus | g005249.m1 | 8.10E-123 | 408.4 | >Saccharomyces_eubayanus_g005249.m1 g005249.m1 | 1 | | *g001406.m1* | 1 | 1 |
| Saccharomyces_jurei | g004292.m1 | 3.90E-128 | 425.9 | >Saccharomyces_jurei_g004292.m1 g004292.m1 | 1 | | *g003435.m1* | 1 | 1 |
| Saccharomyces_kudriavzevii | g005184.m1 | 1.60E-126 | 420.5 | >Saccharomyces_kudriavzevii_g005184.m1 g005184.m1 | 1 | | *g004947.m1* | 1 | 1 |
| Saccharomyces_mikatae | g005189.m1 | 4.50E-133 | 442.1 | >Saccharomyces_mikatae_g005189.m1 g005189.m1 | 1 | | *g003199.m1* | 1 | 1 |
| Saccharomyces_paradoxus | g005237.m1 | 6.30E-133 | 441.6 | >Saccharomyces_paradoxus_g005237.m1 g005237.m1 | 1 | | g002616.m1 | 1 | 1 |
| Saccharomyces_uvarum | g002217.m1 | 3.10E-128 | 426.2 | >Saccharomyces_uvarum_g002217.m1 g002217.m1 | 1 | | *g003213.m1* | 1 | 1 |
| Torulaspora_delbrueckii | g001776.m1 | 2.40E-108 | 360.7 | >Torulaspora_delbrueckii_g001776.m1 g001776.m1 | 1 | | *g001214.m1* | 1 | 1 |
| Torulaspora_globosa | g001826.m1 | 1.20E-71 | 240.1 | >Torulaspora_globosa_g001826.m1 g001826.m1 | 1 | | *g003259.m1* | 1 | 1 |
| Torulaspora_maleeae | g003867.m1 | 9.50E-89 | 296.3 | >Torulaspora_maleeae_g003867.m1 g003867.m1 | 1 | | g004518.m1 | 1 | 1 |
| Torulaspora_sp._yHMJ407 | g002906.m1 | 9.60E-81 | 270 | >Torulaspora_sp._yHMJ407_g002906.m1 g002906.m1 | 0 | | *NA* | 0 | 0 |
| Vanderwaltozyma_polyspora | g005365.m1 | 3.40E-25 | 87.6 | >Vanderwaltozyma_polyspora_g005365.m1 g005365.m1 | 1 | | *g002832.m1* | 2 | 1 |
| Vanderwaltozyma_tropicalis | g003717.m1 | 2.40E-26 | 91.4 | >Vanderwaltozyma_tropicalis_g003717.m1 g003717.m1 | 2 | | *g004362.m1* | 0 | 0 |
| Zygosaccharomyces_bailii | g001048.m1 | 1.20E-100 | 335.5 | >Zygosaccharomyces_bailii_g001048.m1 g001048.m1 | 1 | | *g003584.m1* | 1 | 1 |
| Zygosaccharomyces_bisporus | g005101.m1 | 4.00E-95 | 317.3 | >Zygosaccharomyces_bisporus_g005101.m1 g005101.m1 | 1 | | g000677.m1 | 1 | 1 |
| Zygosaccharomyces_gambellarensis | g005022.m1 | 3.20E-88 | 294.6 | >Zygosaccharomyces_gambellarensis_g005022.m1 g005022.m1 | 1 | | *g001256.m1* | 1 | 1 |
| Zygosaccharomyces_kombuchaensis | g000063.m1 | 1.00E-95 | 319.2 | >Zygosaccharomyces_kombuchaensis_g000063.m1 g000063.m1 | 1 | | g000823.m1 | 1 | 1 |
| Zygosaccharomyces_lentus | g004388.m1 | 8.20E-88 | 293.3 | >Zygosaccharomyces_lentus_g004388.m1 g004388.m1 | 1 | | *g001343.m1* | 1 | 1 |
| Zygosaccharomyces_mellis | g002135.m1 | 4.30E-86 | 287.6 | >Zygosaccharomyces_mellis_g002135.m1 g002135.m1 | 1 | | *g003460.m1* | 1 | 1 |
| Zygosaccharomyces_parabailii | g003412.m1 | 1.20E-100 | 335.5 | >Zygosaccharomyces_parabailii_g003412.m1 g003412.m1 | 0 | | *g003161.m1* | 1 | 1 |
| Zygosaccharomyces_pseudobailii | g001287.m1 | 2.00E-100 | 335.5 | >Zygosaccharomyces_pseudobailii_g001287.m1 g001287.m1 | 0 | | *g000532.m1* | 1 | 1 |
| Zygosaccharomyces_pseudobailii | g002625.m1 | 2.90E-67 | 226.5 | >Zygosaccharomyces_pseudobailii_g002625.m1 g002625.m1 | 0 | | *g000873.m1* | 0 | 0 |
| Zygosaccharomyces_pseudobailii | g003112.m1 | 7.60E-58 | 195.6 | >Zygosaccharomyces_pseudobailii_g003112.m1 g003112.m1 | 0 | | *g000765.m1* | 0 | 0 |
| Zygosaccharomyces_pseudorouxii | g004410.m1 | 6.80E-94 | 313.2 | >Zygosaccharomyces_pseudorouxii_g004410.m1 g004410.m1 | 1 | | *g003387.m1* | 0 | 0 |
| Zygosaccharomyces_rouxii | g001572.m1 | 9.90E-94 | 312.7 | >Zygosaccharomyces_rouxii_g001572.m1 g001572.m1 | 0 | | g000788.m1 | 0 | 0 |
| Zygosaccharomyces_sapae | g000017.m1 | 1.40E-93 | 312.7 | >Zygosaccharomyces_sapae_g000017.m1 g000017.m1 | 0 | | g000163.m1 | 0 | 0 |
| Zygosaccharomyces_sapae | g006118.m1 | 2.00E-89 | 299 | >Zygosaccharomyces_sapae_g006118.m1 g006118.m1 | 0 | | *g005674.m1* | 0 | 0 |
| Zygosaccharomyces_siamensis | g000372.m1 | 2.40E-83 | 278.5 | >Zygosaccharomyces_siamensis_g000372.m1 g000372.m1 | 1 | | *g001009.m1* | 0 | 0 |
| Zygotorulaspora_chibaensis | g002973.m1 | 5.30E-86 | 287.3 | >Zygotorulaspora_chibaensis_g002973.m1 g002973.m1 | 1 | | *g002417.m1* | 0 | 0 |
| Zygotorulaspora_danielsina | g002121.m1 | 3.90E-77 | 258.2 | >Zygotorulaspora_danielsina_g002121.m1 g002121.m1 | 1 | | *g002673.m1* | 0 | 0 |
| Zygotorulaspora_florentina | g004327.m1 | 2.20E-100 | 334.5 | >Zygotorulaspora_florentina_g004327.m1 g004327.m1 | 1 | | *g005778.m1* | 0 | 0 |
| Zygotorulaspora_mrakii | g002003.m1 | 4.20E-94 | 313.9 | >Zygotorulaspora_mrakii_g002003.m1 g002003.m1 | 2 | | *g003717.m1* | 1 | 1 |
| Zygotorulaspora_sp._yHDO592 | g004529.m1 | 3.50E-60 | 202.5 | >Zygotorulaspora_sp._yHDO592_g004529.m1 g004529.m1 | 0 | | *g001244.m1* | 0 | 0 |
| | | | | | | | *g004559.m1* | 0 | 0 |
| | | | | | | | g005482.m1,g003054.m1 | 0 | 0 |
| | | | | | | | *g001027.m1* | 0 | 0 |
| | | | | | | | *g002138.m1* | 0 | 0 |
| | | | | | | | *g002404.m1* | 0 | 0 |
| | | | | | | | g002584.m1 | 0 | 0 |
| | | | | | | | g002262.m1 | 0 | 0 |
| | | | | | | | *g004657.m1* | 0 | 0 |
| | | | | | | | g002679.m1 | 0 | 0 |
| | | | | | | | *g004231.m1* | 0 | 0 |
| | | | | | | | *g004487.m1* | 0 | 0 |
| | | | | | | | *g004700.m1* | 0 | 0 |
| | | | | | | | NA | 0 | 0 |
| | | | | | | | g001657.m1 | 0 | 0 |
