## Supplemental Table S1 for "Functional diversification across yeast lineages of the nucleus-vacuole junction forming protein Nvj1"

1 >saccharomyces_cerevisiae
MTRPPLVRGIFSLGLSVAVLKGVEKTVRKHLERQGWIEPQKVDYELIFTIDRLKNLVDNK
REALTAEQPDAGELSWRKVFNFISRQSSELDTRIYVLILLLSFLLPIAWTVLDGDRETTL
EDKDNDCNVDLIENERRLKHYNDGERAVLQFGKNRSEPIILSYKDMNVLEGEHEFTSKEE
HSNSHLTSKSENALNQVGSEDLLGCHLEKQLEEDKNEPNGEADGEDDNNREKDCSSSSEV
ESQSKCRKESTAEPDSLSRDTRTTSSLKSSTSFPISFKGSIDLKSLNQPSSLLHIQVSPT
KSSNLDAQVNTEQAYSQPFRY

2 >saccharomyces_jurei
MTHPPVVRGIFSLGLSVAVLKGVKKTVRKHLEKQGWIKPHEVDYELIFTIDRLKNLIDDK
HKPLTGVQQDTSELSWQKVLNFISRQFSELDTRIYVLILLLSVLVPILWSVLDGDHEGVL
EDNDSDINVDIIENERRQKHYNDGERAVLQFGKNRSEPIILSYKNMNVLEGEHEFSNKKD
HDTSHLTSKSENALDKAENKDVSDGHPEQQPKGDIECDKNDLSKEKNGEDRDNKEKDSSS
SSSELESQIESKVESTAEPDVLSRNTRTTSSLKSSASFPASFKGSMDLRSLNPPSSLLHL
QVSPTKSTNLDAQVNTEQAYSQPFRY

3 >saccharomyces_paradoxus
MTRPPVVRGIFSLGLSVAVLKGVEKTVRKHLEKQGWIEPQKVDYELIFTIDRLKNLVDDK
RESLTAEQLDTGELSWRKVFNFISRQSSELDTRIYVLILLLSFLVPIAWTVLDGDHEGTL
EDNDNDINMDLIENERRQKHYNDGERAVLQFGKNRSEPIILSYKDMNVSEGEREFTTKKE
HGNGRLISKSENALDEVGSEDVSGCHPEKQLEEDNNELSEEENGEDDDNKEKDRCSSSEV
ESQSESKKESTAEPDLLSRDTRTTSSLKSSTSFPISFKGSIDLKSLNQPSSLLHLQVSPT
KSTNLDAQVNTEQAYSQPFRY

4 >saccharomyces_kudriavzevii
MTRPPVVHGAFSLGLSVAILKGVKKTVRKHLEKRGWIEPRKVDYEFIFTIDRLKNLVDEK
DGTAAVWQHDTSELSWRKVLNFVSNQFSELDTRIYVLLLLLCVLLPIAWTVLDGDSEDLL
NDNDNDDNVGLIGSKIRLKHYNDGERAVLQFGKSRSEPIILSYKDMDMPEEDREFTTKKE
REDCQLVSKSENALDKVENVESPGGQAKQHLEVDTQCNRDESGEKINGEGDSSKEEDRCS
SSEVGSHDESKEEVVTDPEFLSRDTRTTSSLKSSISFPISFKGSINQPSSLLHLQVSPTK
STNLDAQVNTEQAYSQPFRY

5 >saccharomyces_arboricola
MTRPPVVHGAFALGLSMAVFKGVEKTVREHLEKQGWIEPQKVDYELIFTIEKLKNLVDEK
YSQAVEHCDDTSELSWRKVLNFISRQSSELDTRIYVLILLLSVLLPIAWTVLDGDREGTP
DDNSNDVDENLIASAKQLKHYNDGERAVLQFGKNRSEPIILSYRDMNVSDGDHRDFTTKR
ELEHSHLTSKSENALAEMNDEDTPGDQQEQPLEIDIECEKNEPGEETKNVNKEKDHSSSS
EVESHNEAKKETVADPEILSQDTRTTSSLKSSTSFPISFRGSIDLKSINHPSSLHLQVSP
TKSTNLDAQVNTEQAYSQPFRY

6 >saccharomyces_mikatae
MTHPPVVRGIFSLGLSVAVLKGVQKTVRKHLEKQGWIEPQEVDYELIFTIDRLKNLIDDK
HNPLTAVQQDISELSWRKVLNFISRQFSELDTRIYVLILLLSLLVPIVWSVLDGDHEGAL
EDNDNDINVDIIENERRLKHYNDGERAVLQFGKNRSEPIILSYKNMNVLEGEHEFTNKKD
HDASHLTSKSENALDKAESKDVFDDHPEQQPKVDIKCDKNDLSEEENGEDSDNKEEDHSS
SLELESQSESKVESTAEPDVLSRDTRTTSSLKSSTSFPVSFKGSIDLRSLNPPSSLLHLQ
VSPTKSTNLDAQVNTEQAYSQPFRY

7 >saccharomyces_uvarum
MTRPPVMRGAFSLGLSVAVLKGVKKTVRKHLERQGWLEPRKVDYELIFTIDRLKNLVDEK
HGSVPAVGQHDTAELSWRKVFNFISRQSSELDTRIYVLILLLCLVVPVAWTVLDGDNDSS
LDENDDYDDLNEDARRLKHYGDGERAVLQFGKNRSEPIILSYKDMDVPEGEREFTAKRDH
NSRSLTSRSENALDRMNDGDTLRDQPEEQVEAGIGCDESELGEETNSNDIDDMTKDRCSS
PEVESQDEFNEDNSTEPECLSRDTKTTSSLKSSTSFPLSFKGSMDLRSINQPSSLLHIQV
SPTKSTNLDAQVNTEQAYSQPFRY

8 >saccharomyces_eubayanus
MTRPPVMRGAFSVGLSVAVLKGVKKTVRKHLERRGWLEPHKVDYELVFTIDRLKNLVDEK
HDSVLAVGQHDAGELSWRKVFNFISRQSSELDTRIYIFILLLCLVVPVVWTVLDGDHDNS
LDDNDDYEDARRLKHYGDGERAVLQFGKNRSEPIILSYKDMDVPEGEREFTAKRDHNSRS
LTSKSENALDGLNNEDTLSDQPEEQVEVDIGCDENELGEETNTNDIDDQEKDCCSSPEVE
PHDEFNEDNPTEPEFLTRDTKTTSSLKSSTSFPLSFKGSIDLRSINQPSSLLHIQVSPTK
STNLDAQVNTEQAYSQPFRY

9 >nakasesomyces_glabrata
MTRPPVLQYGIPFGVTVGAWKSLRKYLGHRMPYSSLHEPSQFEQWADDFFVPETLQEKII
EHIPEPVKHNKLSELFEQVTQRFFDMDTRLSIIMLIVIVLAPLLSPSRKETHENCNDNEN
ELQQSTAEERLKSAETTAVDLDEQEKEIHMEEIEDSTVVAVKLPVATAEEVELMHETFGA
VPALKREPNPSHTASSSSLDSANSKLSRESPRVPLQVSPAKACNSNAQINNFQAYSQPFT
Y

10 >Nakaseomyces_kungkrabaensis
MTKPPVLQYGVPISVTIGMWKGLRKYLADKLPYSVFDEPSRYEQWADEFSIPEINEVAKE
IIPDEIKTSKIGVWAQDIYDKFWDMDPRLSIFILVFLVLSPLLTRSSGDKVQESFDKQEI
NVQNNLVQQEIEEVKEIQQESAHVLAVKEINISPNVLKDELANVPSLAHDDEYIKIEADT
RQTPSASTSSLDSQRSKINDSSLHLPLQVSPAKTSNSNAQINNFQAYSQPFTY

11 >nakaseomyces_nivariensis
MTKPPVLQYGVPISVTIGMWKGLRKYLADKLPYSVFDEPSRYEQWADEFSIPEINEVAKE
IIPDEIKTSRIGVWAQDIYDKFWDMDPRLSIFILVFLVLSPLLTRSSGDKVQESYDRQEI
NVQNNLVQQEIEEVKEVQQESAHVLAVKEINISPNVLKDELANVPSLAHDDEYIKIEADT
RQTPSASTSSLDSQRSRINDSSPHLPLQVSPAKTSNSNAQINNFQAYSQPFTY

12 >nakaseomyces_uthaithanina
MAKPQVLQYGVPIGVTVGMWKGLRSYLAGRLPYSVFEGPSRFEQWSDEYLVPVEVVEVVR
EMVPEEIKSSRLVGVAEGVYNKFWNMDPRVSVAVLLVVVFGPLLVRSEVHAGGKDRNEHT
QVRAEERTASAAQTERAAPLEVVAVKDFTIEPELLKNELANVPAPVLAPAGYDSRQTPSA
STSSLESHRSRVNESTPHLPLQVSPAKTSNSDAQINNFQAYSQPFTY

13 >nakaseomyces_delphensis
MARPPVLQYGIPISVTIGMWKGLRKYLAPRLPYNMLEQPSEYEQWSDDYLVPEIPDLVKE
IVPENIKNSSIGTWVQKVNDKFWEMDPRISIFILLLIILSPLLAKTSNEKVQEREGIDMN
HKIRQLSIEKEEHVDHELYAERRESAQVVAVKDFGLSPNIIKDELANVPSLGHDEYCGLE
SDVRKTPSASSSSLDSKGSKINESSPHLPLQVSPAKTSNSDAQINNFQAYSQPFTY

14 >nakaseomyces_bracarensis
MTRPPVLRYGVPIGVTVGMWKGLRKYLAKRLPYSVFEEPSQFEQWTDEYLVPEIGIPEVV
KEIIPEEVKTSHVGVWMQDIYNKFWDMDPRLSILILALLVLSPLLTRTSEEKIESHKETL
TEERLADPPKISALEQDEEVDVVVVEEEDVEEEEEEEEEEDEEEEEEEDEEEEEEVVEIK
PSVVAVKDINISPVVIKDEIANVPTLAHDDFKTENETRQTPSASTSSLESHGSGINDSSP
HLPLQVSPAKTSNSNAQINNFQAYSQPFTY

15 >nakaseomyces_bacillisporus
MFPDSWFWKREGAQTTTYTTYDNVQNEARDLLEYLVNEESMDKMLNTESYVGNNRLLSYI
WQEVQDMNIGVGFAIVLLCIFGPMILLLLENNGDDKGHGEKEDAPRESIDPFPKLTRKST
PAVSEQPGKITDPIGGHDDENIENKRPETPRKTATDTALNHGHMKEPAKSVQPYPVLIKN
VNLASSTPQTHISLTEQKSFMPIGKLNVGVPDNVQDNASSSSRSTTHSNSASRSTSTSHS
ASHLHLQVSPAKSSNLDAQLNKEQAYSQPFMY

16 >nakaseomyces_castellii
MARPIVRNVFSIGASWAVLRGVRRALRTRMPDDWLWVSHKLVDPHWGELDAEPEELDREV
ISSILGLELDPLSLQRMQNSTLYKWNEKLHYIWDATMDMDARISIVLLIVLVLGPSFYLL
LDKSGNETQRNTVSKKAPSVSVDANEGRTKNEQPMRQRVSQTDTSSLPEISDDEELGDEI

KNLMRSSSALNERNGFQAKSVSETTAMTIMKANDAASNAAERSTNDDQRAEPNPKVLSEV
KSMLSRSKDSSISVQELPTIADHISNNDSSDITAEISGSTPNHTQKAPSLPSGSEEQISV
LSDVRMNRTVAESHTEFVLEKTSKMSSSEDGPPPLAPRSIDLLDGRSERSSRETLSSGSR
QPLHVSPAKAPNSGTQVNEEPAYSQPFSY

17 >Nakaseomyces_sp._yHDO568
MTRPPVVQYGIPVSVTIGMWKGLRKFLADKLPYSVLDEPTRFEQWSDEFSVPEIQEVVKE
IIPEDIKSSSVGKWVQDIYDKFWEMDPRISLLILVVLVLSPLFTKSSSEKEQIYEEQHAT
VEKNISRQELENIREVQEVTNSTTPGETVVAVKDININPNVIKKELENVPSLAHDEDCKA
ESDIRQTPSASTSSLDSQRSKMNDSSPHLPLQVSPAKTSNSNAQINNFQAYSQPFTY

18 >kazachstania_solicola
MRRGPLIGNVFSITSSYGFVVGLSRFLSSRLPYNDKYPWHKDQPCPSPSQHECLLPDLSQ
NIITAESELLNNTRAEYILGWMDKGNNWVDKMDVLILYIVGIICILALPLYDSWVGRGHS
RINRGDEGRDAEEEEEEEDLNEEVTEEESNIVPRKENVTNPPKDDSEDSREQSAEDLELI
DESEKEQKQDSNLSETPEDNIEEVPEEATLEHTSFQQDDSEITPNPDISLEEEVLSNEQE
HLQEPDQEHLQESDQEPLQEQRNTYVPQISEESSPDNGVSSPVPIPKGSRTLNDSISLQS
KPDLLNPNSQSSSSSFLQFSPSKSSHFDAQVDTKNAYSQPFKFK

19 >kazachstania_aerobia
MRRGPVIRNVFSITSSYGFVVGLSRFLSSRLPYNDKYPWYNEQPCPSPSQHECLLPDLSQ
NIITAESELLNNTRAQYILRWLDKGNNWIDKMDVLILYSIGIICLLSLPLYDLWVGRRPG
RINRDNGGRDAEEEEENEDINEDINEEVAEEENNNDSRKEEIINLPKDNLKPIEDNRKQS
EEVLEANEESEKELKEDNEASETSKDHTKEVPEDAIVEHTTSQQDSNEITPNPDISLEEE
TQSNEQEYIQESDNEHLQEPDQELLQEQSNNNIPQIKEENFSNNGVSSPVPIPRITRPSD
DSISLHSKPDLLNPNSQSSSSSFLQFSPTKSSHFDAQVDTKNAYSQPFKFK

20 >kazachstania_unispora
MRRGPTIGNVFSITTSYGFVVGLSRFLSRRLPYIDKHPWYENNNNNPRPSPSQHECPLPN
LSQEVITVESELLNNTSAKYILGWLEKGNKWVDKMDILLLYIVGVLCLLSLPLHDLWVSK
RSKTTNKEVGGSTDLEVRDIATSKTSEEEVAKEEVRSLKETQEQLNANEGDEEPLDEAKY
DEQLEKVSQEQQDKPVEIVVENVEEDIPSTEDIPSTEERDITFSQDTNNITPNPDFTLDE
QDINNEEAEVEEDSEETQSEQDDKDSLHVKEENSTNSAVSSPVPIPRLSHPLDDTLNLAL
KPDLVNPNSQSSSSSFLQFSPSKSSHLDAKVDTKNAYSQPFKFK

21 >kazachstania_bromeliacearum
MTRPPVVTNALTMGASYALFKGIARGLRGRIPEDWLTGDQNDTKSEEELIADHLTQFLKS
TLPFEHVGESHPVLKEYWNWISQSSFLFVLQIVILIAVLLGPPILATIYSKSPQGSEPVV
NDQNKDATNSEPDNYDDEPQQSNNSIPEPQTRGSTTTPSEDIFVMEPLQEDEVDSKRNAT
DDDQASSRISLLRGQPMLKSSSSDSSSFIHISPSKGANLQTQVDTKNAYSQPFKYKY

22 >kazachstania_bulderi
MARPVLQTTGSLGVSYIVFRGMKQLLSKHVPERWLQVEEEHMSKVNELANKMTERLHDSL
PNIPEQTRNDNIILKTIYESFQKVKVYEIAIYLLTILVVIILPPLLSLHMDKKKKTAIEA
FERKQKKEKDEDEEKQKKKEEAKELGDRLEDMRQNEKMEEEEETRKDSNQREESTHEEEG
NLSNRSINNYTEEIRLSLTKEERQKLKEEEKEKEQNLIVPDVTEIDEVTKTGPQNESLVN
PALHHSDQEGTEEVDSKIEDKVAYEQHNFIDKIDPLRENDGIDLEDGFNEVDHIVSPLEP
EIIADSVQDISQDNIDTVEEIHEETVQDQEEDDDGQEEQEPDEEKETLGPEQIIVLEKTA
EQPRRIPSLTNPTGSSFDSQSSLHTQVLLPSATSSSYIQFSPTKTSDLKVEINKKNAYSQ
PFEY

23 >kazachstania_exigua_g002519.m1
MARPVMQTTGSLGVSYIVFRSMKQLLSKHVPEKWLQVEEEHMNKVNELANKMTERLQESL
PNIPETTRNNNIILRTIYESFQKVKVYEIAVYLLTILVVIILPPLLSVNMDKKKKAAIEA
FEKKQKKEKDEDEEDKKNQKEIEQLGDRLENMRQDEKKEEQARKDQDQDQESETVQEEEE
NLSNRSINKDTEEIRLSLTKEERQKLKEEEEREQSLLEPDVTEIDEVTKTEPHNEFLVSP
ALHHPDQDETELKDNEVENKVEYEQQNFIDKVDPMRENDGVDLEHGFNEIDHIVSPLEPE

IIEESVHDISADNIDPAEEVHEETVQEQDEVEDEQELENNEEKETLEPEQAIVLEKTAEQ
PRTIPSLTNPTAASFDSQSSLHTQVLFPSANSSSYIQFSPTKTSDLKVEINKKNAYSQPF
EY

24 >kazachstania_exigua_g000048.m1
MARPVLQTTGSLGVSYIVFRGMKQLLSNHVPERWLQVEEAHMSKVNELANKMTERLHDS
LPNIPEKTRNDNIILRTIYESFQKVKVYEIAIYLLTILVVIILPPLLSLHMDKKKKAAI
EAFEKKQKKEKDEDEERQKKQEEIEDLGHRLEEMRQNEKVEEEETRKDLNQTAESTHVE
ETEEGNLSNRSINKNTEEIRLSLTKEERQKLKEEKEKEQNLIVPDVTEIDEITETGPKN
ESLVSPALRHSGQEGTEEIDSAIEDKVAYEQHNFIDKIDPLRENDGIDLEDGFKEVDHL
VSPLEPEIIVESVQDISQDNIDTVQEVHEETVKDQEEDDDDQEEQEPDKEKETLGPEQI
IVLEKTTEQPRIPSLTNPTGSSFDSQSSLHTQVLLPPATSSSYIQFSPTKTSDLKNAYS
QPFEY

25 >kazachstania_gamospora
MARPVIQSGFSLGITYIMTKTIKNLLSKHIPQKWLDIENEQKIKVNKIIEKVTDNLQLHI
IEPKSNLKFNKFKINFFKFLNNYQFIIYFIIIFTILFISSIILILNDGKKIIKKKENQCL
NDNLSRIESPNPIDQRNIIETNQLINSFNNLNEDNTKLDHINEQDNESNITTKIEDFEKE
STNEISDNKSTNEILDNKSIEKIDIKDENDSLDLENNDKNSIEDKHEGSNEELDIQEGFK
VVDSINSKDNSDSINSKDNSDSNSTKDNSDNIKDSNDNNDSIKDSKDIENEFKNNNVKII
NTLDEPFSKSSTVSSNIDTKTNSLYSLQLSHSNTSPFIQFSPTKTSTINVKLNKNNVYSQ
PFEY

26 >kazachstania_hellenica
MARPVIQSGFSLGVTYIMAKSIKKLLSRHIPQSWIDTQNEQKVKVSKAVEKVTESLQLRL
PEIPEPKILTSNDHFRNFKNNLVKFLNKNEILIYIIIILMFLFVPSIISILNMQKKRNKN
EGKDNYIEKTSNVIEPRNDNITEKDEIINFNNEKPIESVPHKDELMKNLNVDISSLDEVN
EFEQNCQNNDIETGNNHESKVKEIQDNSLEAKKLDDDEDSANDEHEGSIEELDIQEGFKV
VDSISSESNDDTNSSKYSSGSTGNNLEDDNLIDDNTGDCVAEDDDVEYGNAGHDNASDNN
ASDDNLKDDDVENENLKDDKLITDSLNSTNNKVENELKEGELETHNENIEDTQLSNDRFD
ERFSKGSSASSNLDTKSNSLYSLQYSHSSSAPFIQFSPTKTSPINVKLNKGNVYSQPFEY

27 >kazachstania_aquatica
MRRGPVVGNIFSIASSYAFVAGFARYLNRRLPRNIENGPWWMNIPPPSPSEHDSPLPDLS
QDIITVESELLNHTSAKYVLDWVEKGNKWVDKMDIFILYLIGLICLLMLPLHDVWVERSR
RSPRSRDRHDETSLTQQEDTQEEKGNKEVHKEENTPKNEEVQREEIQPNILNQEREEITP
IEDITLDASEGSELEEKERDTEENEHPIQIEVEDEDEEDKTDQDRGEEETKPSSSATSSP
VAIPKPLKQESESLNLPFKPDLPNQSSQSSSSSFIQFSPSKSSQLNAQLDTKNAYSQPFK
FK

28 >kazachstania_lodderae
MARPVVQHSFSFGASYIILKSVNKLASRYVPEHWVRKKDDKMDKVNLLIQKVTESLETQL
PSISEAATTVRTPSEGEFAFVKTFVIKLREKFQQHELLTYLITVFIIVFVPVIWKLYKNS
GKETTPKTDHEDSHLEILSPEPVDKYQINETEKLLNNDFKQSNQVLPNNENKNSQKCTLK
KTPEDTFTSPDNMTSDSKVDGDQLNNKGGSITDDSDDRDQKVEYIENNITEKLLLDDESS
AKDDELISPSHNTVSKNIDIDNEIDIEAGFKAIDKIDENQTRKTITGKDNTISEPNNNEQ
ESDTHKTGFQDGRLDSLEQDPNTTRLQNIADSKLEKKLESELKEIEHVLERKEENHEAIR
YNSNPALQLIEENSSKRSSDSSNRHSSNSIQSTKSLHSNIPSVIHYSPTKSSDLSVEINT
GNAYSQPFKY

29 >kazachstania_piceae
MVRPIIQNTISFGLSVSALRGVERVLLRILPDKWITSQDESVDASKLIQKMSESVQTPIP
AGIEQFLQKSSSDAHGWKHIIFIIWNAILTFEIEIYITTVCLLIFTPAIFALVKIRSRKI
QKEKVKVDEPVDKSETYAINDSHLLSPELLLHKQNYCEAKNPIETFDELHYPICETVGKE
TNAVNQRIEDGDHEDTKYTTSDKNYLDTEEKIKEIEQPQKVVNENTLKLNILRKEAQINH
TNENKINQILNNDTREPKNNDGNNILIPSDSIDFRVQSGNLNKQILNFNKANEDGNKSNL
INYNPNNFIESQHEGSIKEIDIQEGFKPIDAISKDANEMINEPSEKESIVKLEEKLEAVH

EEQQCIQIANEDSNPTNDIDYNSENYEEIEEPKAEYPKNNSQTVSLKISVHPTHDTVHSP
SLLYSNTSSFIQFSPSKSGHLSAELNKKNAYSQPFKY

30 >kazachstania_saulgeensis
MARPVLQTTGSLGVSYIIFRGVKQLLSKHVPEKWLRVEEQQRNRVNGLAQRVGEKLQDSL
PNIPEDTRNKNILLKTVYDSFQKIKLYEVAIYLLTILVVVILPPLLSLHMEKKKKAALEE
FEKRTQQNNDNNNDDDNTNVSNIHEEDPEEEDPEEEDPEEGDPEEGEEEEEEDVSEAENK
LSNISMNDNAPLSLIKDELPLADDIPKEQEIADVDNVEEELSHMDENDNDFSQSNITLSS
YHIIPDNVNNGEYKGGKNLESEVAERTHDEIDLETGFDEIDQIGSPLEPEHNEVFKENIE
TVEEPQDVQEKEVLEPEQGIVLEKSVKQPRETPSLSNPKALSFDSQSSLHTQMLFPSAGT
SSYIQFSPTKASDLKVEINKKNAYSQPFEY

31 >kazachstania_servazzi
MRRGPVIGNVLSITSSYGFVVGLSRFLSSRLPYNDKYPWHKDPPCPSPSQRECLLPDLSQ
DIITAESELLNNTRAQYILGWVEKGNNWVDKMDVLILYIVGIFCLLSLPIYDLLVGRLRD
KANGDDGDRDMNEEEELSEEESYNAPIQEEAGDPPKDKLEPIEERKKEVDEYGEALKMEE
NNVEEEPAEATMEPAILQQDINEITPNPNISLEEEVFDEQENQQELVENPEQNDVKEQND
EDIPPIKEESSLNIGVSSPVSIPKVSQALDDSLSLRSKSDVLNPNSQSSSSSFLQFSPSK
ASHFDAQVDTKNAYSQPFKFK

32 >kazachstania_sinensis
MARSLTRPALGRSAFSFGVSYGIVKGISKILQRRLPRDWIRSSGLDESGSEANNVVIDDM
EFLQQQLAQFLKDTLPVDEKLVAQSRWARWFSRGKTYVQDASILIWLLLLVLLVFLPPFW
SLVSRKHKGNSELVEDNHINESVPTVEIVEPVPSRRSSASIVDTDSDTPVDNPRDDETHD
YDTDLEPSEENVEEQEQEQEQEQEQEIAEEKHEEVVNETPDVTIENVLDEQSPKSTATNS
TESPTRSLQAPTLEKSESFSSSFVQFSPTRATNLSTQVDTKNAYSQPFKY

33 >kazachstania_yasuniensis
MRRGSVIGNVYSITSSYGFVVGLSRFLSRRIPYIDKQPWYKQHPRPSPSQNECPLPNLSQ
DIITVESELLNNKGAKYLLGWVDIGNKWVDKIDVLVLYILGIVCLLLLPLHDLWTRRHHG
RVNIDRYDDNQEATGEEENVNNHDTLQDIPQDQAEPPKDVTPEVEEVIRQPEEVENILEN
RAIVESEEDNKPQEIQIEQEPEKEELEETQPADTTFQQDIADITPNPNITLDEEEPEEPE
EPEEPEKPEEPEKLEEEQSEQDNEELPQVKEEISSNSALSSPVPIPRSSQSLNNSLNLQL
KPDLVNPNSQSSSSSFLQFSPSKASHLDAQVDTKNAYSQPFKFK

34 >kazachstania_barnettii
MARPVLQTTGSLGVSYIIFRGVKHLLSKHVPEKWLRVEEQQRNRVNDLAQKVGGKIQDSL
PSIPEKTRNNSALLKTVYDSFQKVKLYEVAIYLLTILVVVILPPLLSLRMDKKKRIALEE
FEKRNKENNDKHDDNDNINLSKTHEEEEEPEEEPEEEPEEEPEEPEEEEYEEEVSETSNE
LSNGSINNNAQLSLTKYEQTTEDETPQEQELPELEGIEDDKDIDESNITLASYHGVASDV
NNNEIEIEETPESEVTNITHNGVDLEAGFEEIDQIGSPLQPVHIEVPDENIEPIEASNDV
QEKEVLEPEQAIILEKSAEQPREVPSLSNPIALSFDSQSSLHTQMLFPSAGSSSYIQFSP
TKTSDLKVEINKKNAYSQPFEY

35 >kazachstania_psychrophila
MARPVMQTSCSMGVSYVVFRGLKRLLSKHVPEEWINVEQQQKKKVESLVQRVADSVSEQF
QIPEINEATTSIFIKELRNVIKQLKTHELILYLVIIFLLIFIPPLTSLLGASKRKSHDKE
QPEANNLGPVPNLTHEPVVKQVYIIDEPKPVPENNEDDFTGQHKSETMDECIHEEESLEA
IKEELEEEYQEELQEEFLVDPEEEEEIEETRTEVEDIQESLENDRLLEEEEEEMVESYES
LRNEKDNQEGPEDQDTEGTADVTLEPNEKEYFEPEMIFTLEKNGDEFNVSELEPSKTFYK
HTEILSDSQSSIQSPKLFYSTTSSFIQFSPSKSSHLIVETNKNNAYSQPFEYKAQ

36 >kazachstania_jinghongensis
MTRPVIQNTFTLGGSYIIIKGVKKLVSKYVPEDWVSLQRSKTDEANELIKKVSEQVQEQI
VDQIHRTDNEHFNAVIELVLACWKWFKKNELKIYIILIIIIGLLPGILLVIFTNPPPPFR
KPRQRPAKLLSPKPMHMRKISDTRKMISEYEDGKQLAADQNVVQGSLESPALYGNRNASE
SLSTNTSETKANNVIVDFTQQDSNGASKESSSNNSNIIDEVSQSKHLINSDNDLQSNNPI

GNVKHPLATAKVINLEGNDNVSSLISNSIPVKDKEAIQIKPESATNDGTDRQESNIEGSE
KTLSMPIEQSQNDTISGHTGSDDEADIESNFKATEYLGNEESGRLTSMDSQLHQELKKEE
ETLNKHTKIEKLNISDGLESKVSLNSLERKSFSSVQSPTLLLAPTNAYEIQFSPTKAASL
SVDINMDNARPQPFTY

37 >kazachstania_siamensis
MRRGPVVGNVFSIASSYAFVVGLGRYLNRRLPSLNGPWWIEHDTSSSPSLSPSPSQHNVP
LPDLSQDIITVESELLNTTGAKHILKWLEMGNQWVDTMDVVILYLICGVCLISLPLHDIW
MARGSHKETVISTAEVVQSLKESSPIESVPQKEEEQEEQEEAEEEQEEKEKQEEEEEQEE
ESFSSGEDSQPEEHEEEEAQEVEDDNLQTESSINLTNEPASQSPSTTSKPITIPKPAKSS
SQDLDQFKPDLTNSPSTGSSFIQFSPSKPSQLNAQLDTKNAYSQPFKFK

38 >kazachstania_taianensis
MTRRPVLGGAVGLGASWLFAEGVYRLLDARLARTRWWWRGAGEEEGDFADQLTRFLADTL
QGVEEGVAEGGVLEESVWRVRGRGAVAWVRGRWDSVEVSVLLGVLVVCLLWVPLTGLAAE
RSNERERARSEREREEGEEGEPELGEPENESEREEDERELVYSQDTAGTQDDESSSLENS
SLEQNSQDSFDHVSHDSHDSQSDQASVARSFVQFSPRKARDGQAQVDAKNAYSQPFKYK

39 >kazachstania_naganishii
MARPLARPALGRSVFSFGVSYGVVKGISKILQRRLPRDWIRSSGVAEGGSEVGDISMDDM
EFLQLQLAQFLRETFPVDEKLGTQSRWTHWFSRGKAYVQDVSILIWLLLLVLLVFLPPFW
SLVSRKHKGHCELVEDNHIKESVPEIVEPVPGRRSSASIVETVSDTPVDNPRDDETHDYG
TDFEPSEESVEEQEQGQEQETTEEEHGEVVNETPDVTIENILDEQSPKSTATNSTESPTR
SLQAPTLEKSESFSSSFVQFSPTRATNLSTQVDKKNAYSQPFKY

40 >kazachstania_martiniae
MARPVVKSGLSIGLSVVIIKGVSRIVQQVAPEHWFFSKAQYMEEDLYRKIDEANDYLIGE
ISTSLLVGLLVFLVFIPIMRAAWRVFFSKKQPDDDEDNNKQYSVIQEPTPEHQRQTYELQ
KTFLDKDKNASPFPKFDGIDAIELNSGDNKVDVQEEEDGEEEAEILNPRDFENEIYDDSQ
SEIKPLVYHPQLENEDAPLSNSSASSHSSINHGNKSELLTTPQTSFMAIPSLKLTPQSHI
QFSPSKTSELDAQLNKNNAYSQPFTY

41 >kazachstania_turicensis
MARPVLQTTGSLGVSYIVFRGMKQLLSNHVPERWLQVEEAHMSKVNELANKMTERLHDSL
PNIPEKTRNDNIILRTIYESFQKVKVYEIAIYLLTILVVIILPPLLSLHMDKKKKAAIEA
FEKKQKKEKDEDEERQKKQEEIEDLGHRLEEMRQNEKVEEEETRKDLNQTAESTHVEETE
EGNLSNRSINKNTEEIRLSLTKEERQKLKEEKEKEQNLIVPDVTEIDEITETGPKNESLV
SPALHHSGQEGTEEIDSAIEDKVAYEQHNFIDKIDPLRENDGIDLEDGFKEVDHLVSPLE
PEIIVESVQDISQDNIDTVQEVHEETVKDQEEDDDDQEEQEPDEEKETLGPEQIIVLEKT
TEQPRIPSLTNPTGSSFDSQSSLHTQVLLPPATSSSYIQFSPTKTSDLKVEINKKNAYSQ
PFEY

42 >kazachstania_kunashirensis
MARPVLQTTGSIGVSYIIFRGIKQLLSKHVPEEWIEIEKQQKNKVDTLVQKLSNVVDEQL
RVTDQEKQGMNSLYRIFYSTVEQIRINEILLYIVTIIVIIILPPLLAIYMKNKRDEALKT
MENRNRDDDTNNGFSHESRSGNSTETKVLQASETENSQNDEDYSVEDQEQKQDEREQDTF
HTTKEIKNFETNKPIDFSLRSIKQTEPEANKEGRENLDSDTDEIQEETEDIKSNRLNDQK
QEQEREEIGMNHNGTVDEIDIEDGFQKLDKIISPLEADVYDVSKDQLETQIKESDVEKEN
LEPEQTIVLEKPSEPREIPDSTFHTDFYTDSHSSLHSPKLLHSGASSYIQFSPSKSSDLS
VEIDKRNAYSQPFEY

43 >kazachstania_spencerorum
MPRPVIQNSLSLCGSYFLFRGIQKLVSKYVPDKWITLQRTKTDEANELLKKIAEQVQENI
VEKVGKTNNRHFDALIEFLLPYWRWFKVNEMKIYIILIIIIGLLPGILLVIYTNPPPPFR
KSRQPTAATLLSPNPEHMRKLSDSKKMLQKFEEGNDEHSLQQQGEKVNNDHDTFSTSNET
EKATNVRKDDPKEENSNDLDDATNDLDDIITKITVDAHAKEIVNNELQKIEPIHNIEDSK
IYISKESDDSSSLVGSESFEKIPGDDNESAVKGSDLSPDRTKRIVDDKDSSFVISEHIGS

DDEADIESSFKVAESLGSAERDELNQLELELELKLKKDAKDDPLILQIDSESKDSVSCIE
RKSFSSIQSPTLLLPSTNSAAIQFSPARSANLSVDINKDNARPQPFTY

44 >kazachstania_africana
MTRAPVVRSGLSFAASYVIVRGVKRFVENHVPEEWLSIKDQKEIEATEALRKVAEVIQAQ
LPPIAELELDDRNSTGLKNVYHIFMRKMLALRQYEIEIYVFIIFALIMIPAIVSFWYGTR
SNRKHLDRDSDDKGSKSYNVQDMSEKELNSTPPSSKRKTKKNKRKEKKNSNKKTKRADDS
TNENSPSGNKNTVRSNPSDDNGKNRLFREPHNLLLNDDASNASDFLDHINAVSNSSGNKS
DNVLGDEDTDENTHEINDNELDNLAKSYIIDRVREKEGSIPREFPKLELSSPLNDGLSNR
PLRSSSQTSAGTNSLDRARHSRTPSFIQFSPTKQSTQSVQVDKNNAYSQPFSF

45 >kazachstania_viticola
MTRQPVVQGAFSFGSSVALIKGIQRLLQKRLPEEWLFFPDVSELEGGDNIDKAIRHLSLA
VETQLGNLHDFNTDGNDNTVKLLKKMLISISYYQIGIYLFIIFICVFSPILHYILRSFNT
TTSETNLQPMENIIKQATISSLPTPFDKTHIHDTEIAILGYNKSRSEPIFSSLSQGTFIS
DIFPKPSNSNTNKILRYGDASEPAKLDHNTVPNERRQVHDLNVTNLLGDHFSSEEIKLEY
RDGLGQVDEDEYISSEDDSKLSNEGGLEKISSSTSIPETGTEFKSNSDLKLLAEEEKNKG
TNGGPKENTEQTSDSYFFPTNRKSNSFLNQLEYLQVSPISSKFTNVEVNTKNAYSQPFTY

46 >kazachstania_humilis
MTTTPTAPQGVQMTDSEKQKIAELADLKYQFDVLSNELFHLKEYISLIDHNPKLNNETES
YGKFLMRENLVVTDGLNADPLTNSRRGVIRRSHRHQAQQPTSVINETSGEPLKGLDAVQA
IVQQRFSSQANEIAQGISEEKSHTRAEKQKPKVKTVKHSVPDTHVKKEEHTLPISSKSTK
AASSSNQPLKRKHEKIRETPSIKKDDVDDVPTLHSEHVPEFDSDYYFTTSEEELEDPSRK
SSRKRPRVKVSFSEPKQTITNPLHVVHPKFGSLTAYLDSFRSLDEDMTPDEYDNFIKELV
IAVKHVKDGLRDGKLTYDVGTSSIQPVVTKDIKPIQHQRPDPISYIYKEQHKHVHQDYLV
NQGVHMSKLFQSTRRARIARAKKVSQMIEQHFKHIAGAEERKIKEEEKHKKSMIRGIVQA
LKKRWNLAERAYRVLRKDEEEQLKRIQGKKHLSKILKHSTELLEAQLNQGQSDSDETGDM
SESNISDTELSSEDRDDILSSSSDEQDNDGSENDTEQLNGTSVGSDDALSVNELKRRYEN
LDAVSDEIFDESVVSRTPSESKISETDSQVSSIVKPSVSLSDLFTKNYDSDDASEEDDQD
MSGSYESSDNSGDMELSDSDESSREESEDSKKSSNQQESESQNNETRSPDGTAAYEPPSE
QLSVVDVPIPSLLRGTLRVYQKQGLNWLASLYNNNTNGILADEMGLGKTIQTISLLAYLA
CEKENWGPHLIIVPTSVLLNWEMEFKRFAPGLKVLTYYGTPQQRKEKRKGWNKQDAFHVC
IVSYQLVVQDQHSFKRKKWEYMILDEAHNIKNFRSTRWQALLNFNTKRRLLLTGTPLQNN
LAELWSLLYFLMPQTVVNGKKVSGFADLDAFQQWFGHPVDKIIETSGGVQDEETKKTVTK
LHQVLRPYLLRRLKADVEKQMPAKYEHIVYCRLSKRQRFLYDDFMARSKTRETLASGNFM
SIVNCLMQLRKVCNHPDLFEVRPILTSFEGGDSVMSDYSMINKTIINMISANKVTREVDL
GNLGLTFAGRDSELESHISKSINTLQCSDQFSGRILDLEKCLANNGDMNDNDTSYQDASK
FFLHYGEEKVQRKIDMLKFKKYINELRCERYPVYGKNLINLLTVVDDVKSDDIQNVPLIT
PVTSRLLTDKKVIDNFAVLTPKAVTLDNRMLTLGLDDDSVVPEITRNNMLEEFYNMNNPF
HHLQTKSTIAFPDKSLLQYDCGKLQKLAVLLQELKDNGHRALIFTQMTKVLDILEQFLNF
HGYLYMRLDGATKVEDRQILTERFNSDNRVTVFILSSRSGGLGINLTGADTVIFYDSDWN
PAMDKQCQDRCHRIGQTRDVHIYRFVSEHTIESNILKKANQKRQLDNVVIQEGDFTTDYF
SKLSITDVLGAELPPGKSDDQLLFENGTEVSKNPKTLEKMLAQAEDADDVKAANLAMKEV
EVDDEDFTEETSPNGKDAPDSDDEYEGTSHVEEYMIRLIANGYYH

47 >kazachtania_pseudohumilis
MARPVLQTGGSLGVSYIIFRGIKHLLADRVPEQWLKAEEEQLDKVSGVMKKVAGAVQDKL
PTISEDTRENNTLLRTLYNTIQQAKLNEIALYLLTIIVIVILPPLMSIRNDKKKKELMAR
WEQPDKDQSSKDASGMGNGTSDELNEEDHHQDNKVDHQLSEQDKGEEQHDDSEQLSRESG
NDQSPNLSLTKDNEIEEVVSETEEVDDTVLHESITDETLEEEEESGHDGSVVNNEPEEHS
KIQEELNRLAAEKEHQLLQEEQTASNEIDLETGFEEVDTVARPMEPEHIEVPEEQVSLNE
DTDEEDGNDLIEQHSHESEVDNSPNEKETLEAVQEIPLEKTSEQPRKIPSLIEQTGLSFD
SQSSLNSQFLHAAATPTYIQFSPTKTSHLKVELNKKNAYSQPFEY

48 >kazachstania_rosinii

MVRPVVHNTFSLGLSYSALRGLQRILSRYLPEEWIVQGKKQCSKVVKVVESVQKQQQEAS
RWQRVLALVWDTVVVFEIQIYMAVVCLLVFIPVIYALAGKGNGDRELIPSSQELIKRQTM
CNIEKPVASNSDTNPALNEHLEKSVPPADNSTSNLVQNTETEHQEDSGPGVTPDHVSEVS
VLHKETPTDIEAEPIGDSDADKIVDETNKTIDQSHKTTEQLHESSEQSHNMDSTACSQNS
FIESEHEGSIMEVDIQEGFKPIDSICRDNNPATGNPDKTTSPSQLEEQLEAELEAERPEQ
DSAHPLDVNVTDNSPEQSMKVEEHSPNLIKNPSLHIERQVLYSPGLAHSSTSSYIQLSPS
KSERLSTELNVKNAYSQPFKY

49 >kazachstania_sp._UFMG-CM-Y273
MARPVLQTTGSLGVSYIIFRGVKQLLSKHVPEKWLRVEEQQRNRVNGLAQRVGEKLQDSL
PNIPEDTRNNNILLKTVYDSFQKIKLYEVAIYLLTILVVIILPPLLSLHMEKKKKAALEE
FEKRTQQNNDNDNDNNNDDDDDDEGDNADVGNIHEADPEEEDPEEEDPEEEDPEEEEEVS
EAENELSNVSMNDNATLSLTKDEPPLIEDIPQEQEISNGDNVEDEELSHMNENDNDFSQS
NITLSSYHIIPDDVNNDELEDGKNLESEVAERTHDEIDLETGFDEIDQIGSPLEPKRNEF
SKENIETVEEPQDVQEKELLEPEQAIVLEKSVKQPRETPSLSNPKALSFDSQSSLHTQML
FPSAGTSSYIQFSPTKASDLKVEINKKNAYSQPFEY

50 >naumovozyma_dairenensis
MTRRPVIQSIFSLGGSLAMVKGLKRVLRKYDLPENWLTPTQGNSTQENINKTNELISILQ
EVSNAMHSRDDISFSKQLEEAGDGITWKKVGLFVLKESINMDTKIAAGIIFTILFLPLLS
ALLSNSIGKKLRKKKNDNVDFATQTEQSHHYGPAERALLQFGKSKSEPILLEDNDIHPIL
SNLAGEKIEEGNQSEEEEKGITGGKEDEGKDEEQQEEGDDYGQGESEENVEIISPEVDDD
LNEEGNHIQVEDDENTNSFENVTSFSNNIEDINSFLRPIEEEEGNVNDNKNKADNEQNEV
EADQVEGAQTSIKEDNNGEDRNETDTTYDMAIRTIDFGETPSPNSLTHDVENETSTTNNS
SYKLSHPNNADVALIHLEPDTTALERNTLTFPEEIPDLNEAEALVGGEEKSKDLRVSSNG
SLSSHSNSPHIHVSPTKSMHPDAQVNKEQAYSQPFTY

51 >naumovozyma_castellii
MTRPPVMQTAFSLTGSIAIVKSIERLCRAHLPENWISQKNFEDSELVEVLQDISNAFSFH
ENVNFDESIDNLTWKKVGLFILQEILNLDIKISLGILLLLLLLPLLGVLLHAEITDDRND
DMEPQMLRHYDPAERALLQFGKNHSEPVLVKDHINRGSVSSNRNTRKNTKKAVVKRQNID
KPQEEEETKGKEDLTKEGKIVSYIQEEDISDFKEESNLNSDESPSPVGFKPSKPEEEIVQ
TGESIQDVDNIDNAEEEAIKNNETSDSLLVEHTDVTGFESSDNRLENTMNFPEEVPDLVE
ATVIADASSQFPPDKSSDSLPSIHDSISSHSNSAQLHVHPSKTTNLEGKVTNEQVFSQPF
VVSK

52 >naumovozyma_baii
MTRPPVLQTAFSLTGSIAIVKTIEQLLKPYLPKDWISYQQSNDTELIQILQNVSDAFELH
ERVTFFDESIDNITWKKVGLFIFKEILNLDIKISTGILLFLLMLPLLGLLLNNNKEQAQI
CRIPKSSKLIRHYDSVNPRTPQAEQTVAESAYINASVLSHNERNKDTGHDQHFPMIRGSS
AETTLNNNIENDKKRQEPRARIESSLTSNSNEVARQFDTASNLKEEGKVITMPDKQSINE
NANIHIPVGADGTFDNTIKFPEKSPDISESQKLIEGNTSQYPTEDSTDSLPSTHNSASSR
SNSTSLHIPPTKTSNLEGKVTNEQVFSQPFVHT

53 >tetrapisispora_iriomotensis
MGRTSPLVGSIFSIGTSISIVKGIQHLLRGHLPSNWLHSRGKLDDLVELLENIETVYNEL
DKDASRYSGLMKNTHSLDEYWYGDITIIQLYKRLQRWLFTVDIKIWVGILVLVMVIHLLS
NVFEQEAKGSPVNNTSGYDKRVYTTLGNVSEDQVRMSETPESDNKQSSFTPNASRGAVDE
FDDDDEPEARLDIIDNKCMDKINATFEGVNESELSIAPRYQERLPDTLDTDLEGLDTAQR
LPDINSLQKTDEALVKEIDSKSSMDELKLDESTVEPNVDDNGSIKVNKIQGPFPANITNE
GTPDQESCESKQSSITLSSQYRAYGDKLSFSSDKRHIITTSILSTSPIQTTSNELADIQA
QVTTEQAYSQPFSY

54 >tetrapisispora_namnaonensis
MSRGHRRPVLGWVFSVGTTFAAVRGFEYLVKPLVPEEWLKRSSVEVVTGVLEGEVGAGEV
VEAVLAGVDAQVLDAYWHGELTLVAAWKRLLTWLDDLDSRVLVGVLVLALVISVIPSVQR
RPSTSAVCVETATPTGATALAGHTLSAEAAGRGVSVQPAMPVAPAVPDMAASGDSFTSVE

DPGTIEPGDEFDDDQEQKEQEQEEQKGQEEEEEEREEEREEELHEQQASNIELKQSSASA
SSTDTTQAAVIRSSPNQEGLPAPPTLFLSTSPMTLRSQYRTFGDKLSFSSDKRHIITTNI
LSQSPTQTLSHSAEQPEAKVTTEDAYSQPFQY

55 >tetrapisispora_blattae
MRRNRVLYPTFSFTSPIFLFSVIRTISKRYLPDDWIHSSISTENGTIINPASQWSIKQPE
KSLLEYISEQEIESDILNIADIQVLPQGLTDLKEWHGIDPAIEYFTELNTGVYVFVVVLC
LIASLVSGVYQKVEEEEQGLIETDSENGENNSNSMNEHIENNTDTEATVLQVEFESSEKI
HDEQEKEMNEIKMNCKEEEERSDCKTINDEDYLLKESIIDNQIKNLDEIMHYNKKCDISE
EELNVPIRELNIGENSNSSIFSPTETSTNIYTSINERNIENLVHEDKSQINLENMKDEYD
STNKEAKIVSNSELSNVENSQDLPISSDTTEIDEDESIKKNLETSNSIIDHTAETNAENG
IKNSHTQDLKAMEVNNQGIILENNQVSTVQDNMIVSDKANSPKSLSPSISMHSGSPSIAS
SLKINERRSSSYSINPIQLQVTPTVLSNPSKVINDDTVYSQPFTY

56 >vanderwaltozyma_polyspora
MVRQQHPTLQSMVSFGASFVFVKVFKELIEGYLPEEWIHEVRYLGIDCDDSADKFYTVEA
VGLKDRETVALLDRIDDIIHSRSNKEITTRLTYNNNNNINSNNNYNHISGIWSDLYEMDM
VSLIHNLKKFYLNIDIRVLVVILILLLFGPVVLSLYHDYKGNAADAAVLSPGTPVLRVVP
DDKVDLKLSNMNCAGSSSDEENSGSSLQKLDKFTGETFPKDTETINTSIMEDLKLQYISS
SETGIVIEDDESGDIKSQSGSPLIIYGSDSIVPTAREEEHLDQERAQDQEQEQEQEQEQK
DVPEQTNEIEINITNLPEPDSKIEIQTIKPVVSSPQKQQEKENENENEQQDNESELAASF
TSSKLKSSSRRISTINPTHHIQISPTKTIKLDAQVNTEQAYSQPFTY

57 >vanderwaltozyma_yarrowii
MTRPPVMQSVFSLGTSYALVRGISHTIQKYLPDRWLEVDVGSGTLNNTLSMIESEDFPLL
NKMEEILQRQTEIIHNENSNVIAAVNKIRLQLLDIDIKYLIVLVLLLLIVPIISTILQRV
PNEGKEDNEDNNTNIPVESYIEDKEEERCSIEEDEQTDEVTEDEDEQQDEITEDEEGVEE
NDELEERDTNSIEVNTPLDAINYEQVSISNLITPSKSIVSNKEEGDSIHHESIDIIDEDC
DESVIHTEQTMIGAVEGQNIFYDESKNIIPEENEEKSINILVNENSNTNGVGFYTEEQDL
PALTNNKKFENEKHENIEHDHDQTDSSISFQKDELLVTTPTEKPKLPYINNNSDSPLSRT
PTQLHLQISPTKTIKLDAQVNTELAYSQPFTY

58 >vanderwaltozyma_tropicalis
MKSIFSVGTSIAFVKGVQHFAEKYLPEDWINSKNGLQSNSKLGNMINSMVNEDSVFNELS
NDINIDNLTFNDIYSQDKFNWILFIQYIRAKMIQVDLSISFLVLLLLVFSPIIKSLLLKH
SIPNDKIVNDNNNNNNEYEKDDEIDEDENNNIPNNNLKFNSRPVNTIQSKKLSSKDNQIK
QNTAGVENKTNITTSGKETNVIEKINNQLDETPLTNNNINTRLPTEEVAPLKSVISNDDG
NDNKENIPIIDETYTKNHKQDIINDNASKFSNNSFQTFNIKISPTKTTKLDEQVNTENAY
SQPFTY

59 >vanderwaltozyma_verrucispora
MTRPPVIQSIFSFSTSYALVRSISNALKNHLPSQWLDIKDNENSLIAKELPILNNIEEIL
DRNLNKVIGNNNDKSERHGSKHLDSLNYLDRIQMKLSDIRIEVYIIILLVVLLYPILSTI
AKGRPQEKRSSNVQNEKEVSNPNINNNDNNDGDDDDCNTAAKAIAHNDYEKIKRNDTPKY
SSNFVARTPSKSTSNGVVKIKCDEEVNNSDKINDDDSFNLSITTSIDENNNKNNNSSNNH
IEAKKNEWNSLNIDDDEEPVLFDNRESLIQSEESNVIEDNKIDDQVQTKKQGLDSNNFEA
DTENKQLLQTPEEELKLKSYENLQEQNSKHSPTTLHLQISPNKTIKLDAQVNTEQAYSQP
FKY

60 >torulaspora_pretoriensis
MARPVAAQSLLSLGVSVALVKVIKRFLSESTYEDWWYEHSHVQSNAPTTELVRESVERLD
ANLMQGQRLTWPQVLNFLVEQVTELDVHFAYFVLFLSVAGPLLWLLLDKESSDVDSKTIP
CNEDERVSSSVSTQTEEEPEEKITVRYFHYDPAERALLQFGRSKSESVLDAFKYMPAQKE
YTEPEDDIDQMVKRCTLEAEKAHNLATARLPGPKEQTSQECAQEDSAELQHDKSQLPAKQ
QALLRSSSGIISSTSSRGVPSVKSSRSLLSFKAVPSKSDSSTHSPSFLQLQISPRRTTQV
EVQLSPEQAYSQPFTY

61 >torulaspora_microellipsoides
MTRPPVLQGIFSLGASLAMFKGVKNMLQVYVPDDWLALHSREPSSTYVIHRVTTAVAERI
DPYYSSPKDDLNWTRVLTFIAQEAAELDLRISAALLLVCVLGPALWLLLTRESSHAVGVQ
VSQHATQTEQPPQEPVTFQYYHYDPAERALLQFGKSRSEPILIGYNMPMNFEDEDYAPGN
VENTNKITAGSDLRKEGGPTSKRPSNDTIQAEKQLGQNSVRTQPQDPNADTMVVSPAFSE
TQEEQSKERGLTNLQPQEKAQCSEPCHRESSSWEDSIVFESLENRLESSGRSSSSLPSLK
CTPTDRETSRHSLAHIQLQISPRKTSNVDAQLNPEQAYSQPFTY

62 >torulaspora_franciscae
MARPVAAQGLLSLGVSVALVKVVKRFLGENTLEEWWYEHPQGDEPTTELVRESVETLEAN
LMQCQRLTWPQVLDFLVQQVTELDVRMAYLVLFSFLVGPLLWLLLDKGSGNVDSKTVPNS
EEECVSNNVSIQTEEEPEEKITVRYFHYHPAERALLQFGRSKSESVLDNFKYMPVQKEYI
EPEEDIDQMVTGCTLEAENYAHSSATTRLSGPKGQSSHECLGEDSTESQHGKSQHPAEQR
ALLRSSSGIVSFTPSREVPSVKSSGSLLSFKAAPLKSHSSTQSLSFLQLQISPRRTTQVE
VQLSPEQAYSQPFTY

63 >torulaspora_delbrueckii
MARPVAAQGLFSLGTSITLVKLVKKLMEGGTVEEWWNVSPQAQQSSTYIVDYPIITTEPR
NIHNQKLTWPQVLRFFIEQVEEMDLQLACFVLFLCLLGPLIWSLLDTKSSSAAGNSLLHN
VSTQTEQEPEEKVTERHFHYDPAERALLQFGRSKSESVLSVFKYSPIQFDYNEPEEEVDE
IVAGCTIKTEPIQKTPLEILEEKRRSGVVKPFAPIDQEVILRLDTALSSQKQFKDSSQLC
PKVEHQVLVQNSSGNISPGNREAPSVKSAGSLPSFRGAPLERASSAQSLSFLQIQISPRK
TSHVEVQVNPEQAYSQPFTY

64 >torulaspora_globosa
MARMTAPYSLFFAGTSIALLKLVQKLVERNGADEWLEPSESEYSTYVIEAPKIVHGDLSV
PDTQELTWGQVFEFLAREVERVDLRIWFAVLLFCTLGPLAWSLSTGRSPSEQPPLAPSTC
VSTQTEPEPEVERKDPAPMRCYHYEPGERALLQFGRSKSDSVLFNYVPLEFHEPELDEFE
EVVGRCTLRQERANEVTPKEEPTAVPSVRLAQLVSEALADNGTETDASMVEDDSANRTTL
GDTLGSLPNLRDSSLEIAGRAQSPTALQIQLSPRKTSIVDVQVNPEQAYSQPFSY

65 >torulaspora_maleeae
MARATAPHGLFLAGTSIALLKFVQKLVDRNGAEEWLEPRLGEVSSSTYVVETPKILHGEI
GTSVSQELTWGQVIEFVVKETEKVDVRLWCAVLLVCVLGPVIWSLMEKKPAREPCVVGSE
SASTQTDQEQETTDRKDPVPIRSYHYEPAERALLQFGRSKSDSVIFNYVPTDFNEEELDD
YEGIVAGCTLRGEESEQLTDKEQDQQVTPKKEPAKMPLAELPERLSQTAPSPATGDTSGE
STSVHQASLGKSLGSLSSLRDGSLRRAPSAQSLPFLPIQLSPCKTSIVEVQVNPEQVYSQ
PFNY

66 >torulaspora_sp._yHMJ407
MARAIAPYGLFLGGTSIALLKLVQKLVDRNGADEWLEPSLDEDSSSTYVVEVPKILHGAS
STPISQELTWRQVIDFLMKELERVDVRVWCVILLVCLLGPVARNLLGSKPPQKPYVADSK
SVSTQTDQEAEPADRKDPVAIRSHHYEPAERALLQFGRSKSESVLFNYVPTDFNEEELQD
FEGIVAGCTLRAEELEPVKEKEQDQEVTPRKEPEKRSSVEPPEQLKQTDSNAATGDSSGE
STSVHQTSLGKSLGLLSSLKDDSLRRATSAQSLPFLPIQLSPRKTSIVEVQVNPEQVYSQ
PFNY

67 >zygosaccharomyces_bisporus
MTRHPVVHSALSVSGSVGAVKGIQRVLQKYLPDEWLFSGKVQSSPTNIVPSALEPTFVED
IPLEKGLSYTRVLSFVVDQLANMDARIAITIVVLTVIGPVLPYLFGIQKHVPQTAAEHLD
PAQRALLQFGKSKSEPILLSYVSMDLEQEVEDFSEGEDIQDPLRRREVSLEASQELVQED
GVVFKEEHTEVMQVLKSEEPEPMGKGLSAQEESDDSTSGKQESSSRSVCTLPSAQTSPLG
RRDSYTSHANVYIQFSPMKTSNLDAQLSSEQAYSQPFTY

68 >zygosaccharomyces_gambellarensis
MTRHPIFHGILSVSGTLGAARGFQRILQRYLNDEWLFGNKKDEVSPTSVVPSILEPSFEE
LPEKSKELDWTHVLSFLVEQIGNMDVRLAVAIVVLAVVGPVLPYVFGYREDTSEGTTTTT

TTTATITTTKESEIVASHLDPAQRALLQFGKSKSEPLLLGYVSMKLDEIEENRNAEQENL
LRRQLQLERPHFGKELEDEVRFAERLVERQESRGREREEEIKEPLPQVDESDTSVGGKPD
SSTRSEYTLPSAQTSPLDRRDSYVSHSNVCIQLSPIKTSNLDAQLTSEQAYSQPFTY

69 >zygosaccharomyces_bailii
MTRHPVLQNALSLSGSVGAVKGIERLLQKHLPDEWLFSPKTHSSPTDIVPSALEPTLIDN
DRLTNSLSSTHVLSFIVDQIANMDARIAVTIVVLAVLGPVLPYIFGSRKQVPQTTAQHID
PAQRALLQFGKSKSEPILLSYVSLDLEREEEDVSEKEDSSATLRREKVSMEVKQEIVQED
EPAFKEVMKEVNIVEQVVEKEGPEPTDKEFCVQEESDGSASRKRGSSSRSVITLPSAQTS
PLGRRESYTSHSNVYIQFSPIKTSNVDAQLSSEQAYSQPFTY

70 >zygosaccharomyces_kombuchaensis
MTRHPVVQGALSLSGSIGAVKGMDKLLRKYLPDEWLFKAQSSPTDILPSVLEPTLVKNSP
LEKGLNLTRILSFMVDKIANMDVRIAVIIVVLTVAGPVLPYVFGFHKRVAQKSAEHLDPG
QRALLQFGKSKSEPVLLSYVSLDLEEEVDFSEREDTPAVLRREEVGLEVEKEGDPVIEE
VYSEVKVLQQVAEEPQESTEKDFIMQEESDGSASAKQGSSSRSVCTLPSAQTSPLGRRDS
YVSHSNVYIQFSPIKTSNLDAQLSSEQAYSQPFTY

71 >zygosaccharomyces_lentus
MTRHPVVRGALSLSGSFGAVKGMERALRKYIPNEWLFKAQSSPTDMVPSVLEPTLVKSSP
LQNGLNLTHVLSFMVDNIANMDVRIAVIIVLLTVVGPVMPYVFGFHKHVAQKAVEHLDPA
QRALLQFGKSKSEPVLLSYVSMDLEEGEDFSERENTPAALRREEPRLEVEKKVEPVIEKV
YSEVKPLQQLVEEPQEPAERDLTSQEESDGSGSAKQGSSGRSVCTLPSAQTSPLGRSDSY
VSHSNVYIQFSPIKTSNLDAQLSSEQAYSQPFTY

72 >zygosaccharomyces_mellis
MVRHPILQGTISIGGTFGAVKGLQRLLQRYLSNEWLFENKKSKISPTNVIPSVLEPSSQD
FSNKGKEVNWSRVLSFIVDQISNIDLRLAVVIVVLAVMGPVLPYVFGYGKSDTSITESNI
TGSNLDSSQRALLQFGKSKSEPVLLGYVSMELDELSDRRNSEQVPHLRRHSELEELQAGR
NLQDEVKIAEQLVQRQESRELEKREEQEKEMQQEKGEMDSHLSRAEESDTSAGGKPGSSV
RSEYTLPSAQTSPLDHRDSYISHSNVCIQFSPIKTSNLDAQLTAEQAYSQPFTY

73 >zygosaccharomyces_pseudorouxii
MARHPVVHGALSISGTIGAVKGFQKLLQRYLSNEWLFENDKSEVSPTSVIPSVLESSSQG
LSDKGKELDWSRVLSFMVDQIANMDVRLAVAIVVLAVMGPVLPYVFGYGKVKAASVESEI
IASHLDPAQRALLQFGKSKSEPLLLGYVSMELDEFREHGTSEQVPHLRRQLRLEKQQVGK
DFQDEVKIAEQLVQRQESRELEKEEEQEAEEEQVEEETGSRSPHAEEESVTSAGGKPRSS
TRSEYTLPSAQTSPLDHRDSCVSHSNVCIQLSPIKTSNLDAQLTSEQVYSQPFTY

74 >zygosaccharomyces_siamensis
MVRHPVLQGAISIGGTFGAVKGFQRLLQRYLSNEWLFENKKSKISPTNVIPSVLESSSQD
FSNKGKEVNLPRVLSFMVDQISNIDLRLAVVIVVLAVMGPVLPYVFGYGKCDASTTESDI
TGSNLDSSQRALLQFGKSKSEPVLLGCVSMELDELSDRRNSERVPHLRRHSELEEMQVGR
NLQDEVKIAQQLVQRQESRELEKREEQEKEVQQEKGELDSHLSRAEESDTSAGGKPGSSV
RSEYTLPSAQTSPLDHRDSYISHSNVCIQFSPIKTSNLDAQLTAEQAYSQPFTY

75 >zygosaccharomyces_parabailii
MTRHPVLQNALSLSGSVGAVKGIERLLQKHLPDEWLFSPKTHSSPTDIVPSALEPTLIDNDRLTNSLSST
HVLSFIVDQIANMDARIAVTIVVLAVLGPVLPYIFGSRKQVPQTTAQHIDPAQRALLQFGKSKSEPILLS
YVSLDLEREEEDVSEKEDSSATLRREKVSMEVKQEIVQEDEPAFKEVMKEVNIVEQVVEKEGPEPTDKEF
CVQEESDGSASRKRGSSSRSVITLPSAQTSPLGRRESYTSHSNVYIQFSPIKTSNVDAQLSSEQAYSQPF
TY

76 >zygosaccharomyces_pseudobailii_g001287.m1
MTRHPVLQNALSLSGSVGAVKGIERLLQKHLPDEWLFSPKTHSSPTDIVPSALEPTLIDN
DRLTNSLSSTHVLSFIVDQIANMDARIAVTIVVLAVLGPVLPYIFGSRKQVPQTTAQHID

PAQRALLQFGKSKSEPILLSYVSLDLEREEEDVSEKEDSSATLRREKVSMEVKQEIVQED
EPAFKEVMKEVNIVEQVVEKEGPEPTDKEFCVQEESDGSASRKRGSSSRSVITLPSAQTS
PLGRRESYTSHSNVYIQFSPIKTSNVDAQLSSEQAYSQPFTY

77 >zygosaccharomyces_pseudobailii_g002625.m1_ g003112.m1
MTRHPVIQNALSVSGSFGAVKGIERLLQKYLPDEWLFSQKSHSSPTDIVPSPLEPTFIEK
DCLDNSLSSTRVLSFIVDQIANMDVRIAVTIVVLAVIGPVLPYLFRLKKEIPQTAAQHID
PAQRALLQFGKSKSEPILLSYVSLDLEREGEDLSEREDSSATLRREKVSMEVKQEIVQED
EAAFKEMTKEVSIVQQVVEKEGPEHTDKDFCVQEESDGSASRKRGSSSRSVITLPSAQT
SPLARRESYTSHSNVYIQFSPIKTSNVDAQLSSEQAYSQPFTY

78 >zygosaccharomyces_rouxii
MARHPVVHGVLSISGTFGAAKGFQKLLQRYLSNEWLFENEKDKISPTSLVPSILESSSQD
LSNKSKELDWTGVLSFVVDQIANVDVRLAVAIVVLAVMGPVLPYAFGYGKVDAASTESEI
IASHLDPAQRALLQFGKSKSEPVLFGYVSMELDELEEHGSSEHVLHLRRQLQLEDQHFVK
ELRDEVNIAEQLVQRQESKELEKEEEQEVVVAHSCSRSHSGSPQVEESDTSADGKPSSST
RSEYTLPSAQTSPLDHRDSCVSHSNTCIQLSPIKTSNLDAQLTSEQAYSQPFTY

79 >zygosaccharomyces_sapae_g000017.m1
MARHPVVHGVLSISGTFGAAKGFQKLLQRYLSNEWLFENEKDKISPTSLVPSILESSSQD
LSNKSKELDWTGVLSFVVDQIANVDVRLAVAIVVLAVMGPVLPYAFGYGKVDAASTESEI
IASHLDPAQRALLQFGKSKSEPVLFGYVSMELDELEEHGSSEHVLHLRRQLQLEDQHFVK
ELRDEVNIAEQLVQRQESKELEKEEEQEVVVAHSCSRSHSGSPQVEESDTSADGKPSSST
RSEYTLPSAQTSPLDHRDSCVSHSNTCIQLSPIKTSNLDAQLTSEQAYSQPFTY

80 >zygotorulaspora_florentina
MTRPPVLQGVFALGAGFAMFEGVMKLYNLYGVQRVQANMSEGFTEPSLVHLVTRVTTTVT
ASAGSLQENLGGKPTLDWPQVFDFLLQQMDQLDLRIPIIITTLCVFGPVLFYLWEGKLKA
RKEIRCLTREGSTQTEEISQHENPNILKPQVPEASNHFCQINPAERALLQFGKSKSDSLL
FGYVPMNYDSTDFHLDAEPELIGSFINSHELQKTPHRTPRRVYVNNLSEPSIASQPLPEC
QHKNLDIETCRETSPGSERPGTSDKGDSSFAESVTTNTFDTKKPSSIKSLCSLPSSRGSP
FNTSILSSSDNHLRLQMTPSKTNTQLNSDIAYSQPFTY

81 >zygotorulaspora_mrakii
MTRVPVSQGLFGLAATVAIFKGLKEAMNFYSLGALRLNVSDGNMNPTMTAESHTVTKKIV
TTVTATINTGTKQKLDWPLVLDFVLEELDMIDPRIWIATIALCVLGPVAVYLFDGRRSVF
STNTHSTQTDEVEEVRIPDLPGPSSGKGPSSYSPYFQYNPAERAWMQFGKSKSEPFLFTY
VPISYSTKDSSSEIEQEVKEYENEPSLEFCDKQVAEPATDNFNREESATDNYNHDESAQS
AQSISDQLPSSGNKFRSIQEKKSLHVMNHQQQHLLEDREVSSLDESFTINPFDKRPISSN
SSHSIPSSRSSPMASSVLNSSNSHLRLQMSPTKTFTHLSSELAYSQPFSY

82 >zygotorulaspora_chibaensis
MTRPPVLRSALVLGAGVAMFRGVMQLYDLYGVPRIQANLPEGFTIPLPVHMATTVTATVT
ASARPNHENLYGEQTLDWPQVLDFLLQQMDQLDLRIPIIIIALCVLGPTTFYIWEGKKNA
KPTKRQSTREESTQTDNPSQDENLETPEIKVSEASTQIYQINPAERALLQFGKSKSDSLL
FGYVPMNYDSTDFHSDMEQELIGCYIKNADLQTTPDRTPKHVYVNGLSEPSISYRALPEY
QHENQDIEICHVTPPGSERPGTSDKGESSFAESVTTNTFDTKKISSIKSFSSLPSSRGSP
FNTSILSSSDNHLRLQMTPSKTDTQLNSDIAYPQPFTY

83 >zygotorulaspora_danielsina
MTRFPVFQSLFGLGAGIALFKGLKGIVSSYEDVADRLHLSLPRSVIKPTKVYTVTKKAIE
TVTVTANASPEHELNWPLVFDFILQQIDELDLRLLIAIVVFCVIGPVALYAVKGQKSVMT
EIQSTQTDAPTEEDSCNTLKSDKPQPWLRHPYYQYNPAERAWLQFGKSKSESILFGYVPM
SYQIKDFTPDGDQEVLRFSSTADAPQKGDEQDAAVGQASVEIIGTGSGEIPKKQGKARIA
QNDGLVQPLGCQLQDLSDRDVSSFEESITINPFDKRIPSSKSSSSMPSSRSSPIATSVIN
NSHSHLRIQMTPTKTDTQLSSDLAYSQPFSY

84 >zygotorulaspora_mrakii
MTRVPVSQGLFGLAATVAIFKGLKEAMNFYSLGALRLNVSDGNMNPTMTAESHTVTKKIV
TTVTATINTGTKQKLDWPLVLDFVLEELDMIDPRIWIATIALCVLGPVAVYLFDGRRSVF
STNTHSTQTDEVEEVRIPDLPGPSSGKGPSSYSPYFQYNPAERAWMQFGKSKSEPFLFTY
VPISYSTKDSSSEIEQEVKEYENEPSLEFCDKQVAEPATDNFNREESATDNYNHDESAQS
AQSISDQLPSSGNKFRSIQEKKSLHVMNHQQQHLLEDREVSSLDESFTINPFDKRPISSN
SSHSIPSSRSSPMASSVLNSSNSHLRLQMSPTKTFTHLSSELAYSQPFSY

85 >Zygotorulaspora_sp._yHDO592
MGPPLLRSIFGIGVAAVVLKGIKEFLEVYDYEPDQLHFTPTVLSPNHEYTLTKSIITTVT
VSVERPEKPELDWPTVIDFLFQQLDKLDPRIPAAILGFLIAGPVLIHIIQGRKSNSKIDE
SIQRSSTQTSSTQTSSTQAGFKTESTQTEEVLKNKSSYELKSWRTDFSDLPHQYNPAERA
WLQFGKSKSESVLLGLVPRSYESEDLPPDDEQETNSIINDLQEQKQDLNSESKQMPKIAP
VSLSNEISIGFDRQEKNKMQPSSYSVGELMQQGSRERETFTSEESITVNPFDKRVVSSKS
LSSLSSSRSSPIASSVLHNSHNHLIIQMMPTKTDTQISSDLAYSQPFSY

86 >hagleromyces_aurorensis
MTPQPRIGSIFSLGTSVVIFKLATDWLGDKVPHIWPDIAPSERSQDLTLIETKNDIIKCD
WEDLLDIGKPVVPGVTNGVSWRGIAKVFYDNITDLPVFVDIGIIIVLVLWPVMWGVLRRE
GRTHKEPEAISQKPEEKVSDIVIERTIYHYGPNDRAILQFGKSHSEPFLINSTFYPLVDM
NYDDNDSESSIDYDSHAPSDAGNTRRQCISQGMPNTLCERNKTDNYRKKPDLKMEALSAS
LSIFSEEIENIQIRPDTATTATGRETSTEDSETNSTQPITRIYEFDVRSDNRWETQKKVT
EEQENKDFIKDSSIRKPDMNEGTGADDNETSAIPTEVQDTNSSQKQVKEGCSINLSSEVL
PQQPSDQNSSCLGEADKADVIYSTEVSNHGEKEPKENAETETSKRQDYNFKPDLDLQDSS
DVPTSISNKSSPMSRSYERRIPSQSTTLNYLQVSPIKSANVGGQVTIEEAYSQPFLY

87 >lachancea_kluyveri
MSNRPILQNRIILGIPVALVAGILTKVAVPFPAWLSLGNTEQAKHSNPTRTFEPQMEKVV
YRCVCPQKQQIDWQDVFDFVGTELLRLDPQWVVMFVSISLLLLHKVFGALRNIKDTNEVV
ECHTQTVDIDYEETVHSIPHYEPGQRALLQFQRNKSEPIVFKYSYNEENEIEEEEEEEGE
DISEHTVEQIIRRHLRRGGSEPPGTPTYRSVDVTKPNQLSEAEDGTFVFKNESSVSIRTS
SASVSPTGTLEDRQDLSMTLARKPSKSSITLHFQPSPPKPTKLNIGVTQEQVYSQPFTY

88 >lachancea_mirantina
MITRSSVHTGLLFGIPTAVFVGVITSVPFLLPTWNESRKIEAADCRTAPTSVSVLRTIDE
KTIGPVCHCPLHETSWTDVLDFLSNELMKVDPEWWLLSGCLLVMLLQEFVKKKPQQKDHQ
EDPQIVEKASIRVDDSVVGSPLGLRERALLQFQRNRSEPVDFRYGHPSEQDGFDLDISES
TFNEFLQRFSAKSILSSRPSIQMSPDRTQQPERKAATFTENRTHGYQGKPLGETTGSGAA
SSSVPSSSTNPASVTLSISSPHDVLSRSSENSLNIYFTPQSHSLDDNKAIVTQEQVYSQP
FNY

89 >lachancea_nothofagi
MHNRSTLQNGLWFGAPVALFAGILSQFHIPMPAWLASMSPPSSTIIKTTNPVEMYDVVHI
CHCNERAIEWQDVFDFLRRKLTQPDLEWMSLAVNLLLTLNLLLIVLRKIASNRRAKVTAS
SAAGSSSREVQVLQYSPRERALLQFQKLKLMPLNFKCDREDVWDFEGDDLINWEREYVPG
EMSEQSTDTLRSAIPTVSPPTQVASVLNVTLTPQRVPATAPQPHIVPTEGPIGQSDTPSL
RTPKLKIRATTISSSPTQLKSSSPVLKSVPTTAELLENICQPPTQSSSRTSSHASSVESN
ARSTLRTPENFIPSYHQPSPAKSTIVRTEVTQEQVYSQPFLY

90 >lachancea_fantastica
MHSRPSLVDGVIFGAPLAVVAGLLTHFEIPFPQWLTGPAEVRSHMEPESIVYVCRCPEHG
MRWPDVFRFIASQLAELTHAGWVPLILNIALVLQTLLVLGKLLRKSYSREMTQLPNSALC
DQTDADGQSIDISPYGPRERALLQFEKLRHKKLNFKYCNEDSWILEEDFLLPAGVVKPLL
KQHGPGAMFSDLSPSTVVASVGVQTAISQRSLSLPQSDELPVDNSTAGLASSPSSQDYPQ
LFTPKRSLDAALSSPPTSKTPVSPGFEPRHAPAAVSLSKDPLVLEIQNSSRTSFSSSQNA
KDTLCTPEHVLHAFHHPSPAKSSILRTAVTQEQVYSQPFIY

91 >lachancea_lanzarotensis
MHSRPGIQNGVIFGAPLAVIAGLLAHFGLPYPQWLSDPVKAPSDIEPHSVVHICRCPEHS
IGWPDVFQFVASQLTELTHRDWVSLTFNMILVIQMLLTLSKLWKKSFSGETTQLASTASV
EQREADVVDADLLPYTSRERALLQFEKTRDKTLNFKFCNEDSWILKEDFLLPAKSVKPLF
KDSPETTLSNLSPSTVVTSVGVQTSISQRLPPLSRSNDLHVGKIMSDLPSSPLSHEYPQL
FTPKQSLSAALSSPPTSKTPTSPGFESRHAPAAVSLSGDPLVLETQNSSRTSFSSSPNVR
EVLHTPEYLSHAFHQPSPAKSTILKTEVTQEQVYSQPFIY

92 >lachancea_waltii
MDVHSRSFLHRSLLFGAPVALIAGLMSAPIPKWLWFQKQAVALSPRSPVPVELNNIVHSC
HCVEQVIGWHEVFLFLKSQAMTIDPQWALLALNLLLVLVQTLSSSKDESEEDDRRGFVSG
CEADVTEEEVSKNTKSLPQFNARERALLQFQRIKVEPLSPKYGYAASNDVQDLKENAAKI
AAMIRKYSKRHSSKCKELKSAVSAISQETLKAPPPAQIIINDVAALGMPSEQRAYPSPSH
STPRTSCSSRRELKQAFISSPATSEGRSIKHLLRPSPSRSSFFSTEVTQEQVYSQPFTY

93 >lachancea_quebecensis
MISRRLTQRRSCRLLYDVKVFTTTESVGTKQRESVTGMHNRSILQNSLLFGAPVALIAGI
LSQLMPSAEWLLGHGQVGELSLAAPTTVELENVVHTCHCVKQVLGWHEVFVFLRNELLRV
NPQVALLVFNLLLILLQMLSSRRLPTAPSGDTIATDLEGEDDETSKLMSFVPRYDSRERA
LLQFQKLKTKPLNFKYGCENSCSMRDSTADAASLEELLRRYSSKGESPDVNAAKLKKGLL
SEASVSTQTSAIDLPTVAIAPQLTNSIVRSHCTSRASSFSRRELKNCPVGTPDSQSMIFH
QPSPAKSSVLSTEVTQEQVYSQPFIY

94 >lachancea_thermotolerans
MHNRSILQNSLLFGAPVALIAGILSQFFMPSAEWLLAHGQAGEFSPAVPTTVELENVVHT
CHCARQVLSWHDVLVFLKNELLRVNPQVALLAFNLMLILLQMLSSRKMPRAHDEDNAEAD
LEGEDDETSKFMSFVPKYDSRERALLQFQKLKSKPLNFKYGSDDSYSVRDSSDDAVSLER
LLRKFSRRGRSSEVNSTKLKKSQPPETPSSAQTSTIDLSIAVMTPQLTNSVGRSHCTSRT
SSFSKRELKNCLVGTPESQSMSFHQPSPAKSSVLSTKVTQEQVYSQPFIY

95 >lachancea_meyersii
MQNRSTLQNGAIFGAPLALLVGVISRFSIRVPEWLSGPTETASNMTAYDVVHVCRCQEQV
IEWPDVFQFLGRQLRQLINPDWVPLALNIVLTLNVILATSKHLKARFKKEVTNSSVITKP
CEFDADQSSPYSSLFGPRERAVLQFEKAKLKQLNFKYSNEDSWILEEDCLLPKTVVKKTS
NGSERTLRSPRTLLSNLSPSTLVTSAVVQVSTPQKRSPSPQSENLQSEALLPAVPESFTD
SSLLTPKLKFGSTSRPPHSPRPPKSPNMDSNHAPPAVSISNDVFVLDPRDSSPTASSSRS
NVKSLLQTPEHFSPPFHQPSPIKSSILKTEVTQEEVYSQPFTY

96 >lachancea_dasiensis
MHDRSSLQSRLLFGIPVALVAGLLTQLQVPIPTWLHLTDHERPPVSSAISPKHMNSYDLV
HICHCPERSLEWRDVSRFLLHQLHFVTTDWASLAMNLLFALNVLMWIAQWLASWKKAKNL
LHEEPTATTLDSTFQDERCAALYTAKERALLQFQKLNSQPPHFVYGFEDPWDCDPDELLP
PLRSKGTESVETLHPSSRPKSLSSTQTPITGKTLGKAVPTGANTNQNPAMNVHSSSIRTV
VSRSSLTPDRHSEPPIKPSCHTSSKLKFHAVTVPSTPIHLERSSPEKKPLVSTVEQSTEL
SLPESLAASQDSSSTSSHAKNQTLTPENLPLSFHQPSPAKSTILKTELTQEQVYSQPFIY

97 >lachancea_fermentati
MHSRSSIHSSVVFGLSAAVIASVISNMTMSLPSWKHPPSNTSEVETTFTLMEPVCQCSCP
PLQVSWYDVFQFLGKEFSEVDPQWAIIAASLLVMLAHRLRRNFKPVMQNNSTSTETDLSD
SCLDESNSNSDFTQYELRQRALLQFQKNKSEPINFKYGYKDNDDLDDSDIDEKSIAEIIR
RYSLRNPSPSVRASSEDPTDRTLDTMDQDNTTLSHASKLASKSDRVSGTSIFSRDEDISC
KVVSKSSNSSLTIHFQPSYPKPKLMKAELNQEQVYSEPFIY

98 >lachancea_cidri_NRRL_Y-12635
MLSRSNLHNSVVFGLSAAIIAGVVSKMPINLHSWQSPHSELQNAHNTSTNRNPLINECNC
SQHEVNWHDVLQFLGMEVAKVDPQWAVLAASLLLMLIHRLRHDFAPRLQNKSIGLDSDVK
SELTNEGVDCPTSPPYELRERALLQFQKNKSEPINFKYGYKDDEDLDDSDVDENSIAEVI

RRYSLRNRTPSVKGASESPSTHIYEVADQENATLSHTSKATSKNDRISGASIISKDEEKS
YKVVSKSSNSSLTIHFQPTYPKPNSIQTELNEEQVYSEPFIY

99 >eremotheciu_gossypii
mssgsaligftlrrlsacatirtpllgvdivrlaallvtagvitmwlwfcgqcsasctlf
lllaayrlweegylpvwqpapalplppptttkartcegnsahasevrrallqfrrglrsl
swtsrrnalgvpdspppraqhlhiepalsayvtgvpilapfvckrtphleqqqlapctcs
lasassyyafpphsatrharphsrtsarvhsrlytdtrdiasakgflhyi

100 >ashbya_aceri
MSSGSALIGFTLGQLFSCATIGAPPPGADKLLLASMLVATAVATIWLWVCTQRSAFCALV
LVLAAHHLWEEGYLPLWQPAPALPPPPPLPLNARAREGDAARAREVRRALLQFRRESAVP
LVNIRATRPRGPRPSASSRPAPAHRPCDIRIHGQRAQPSTSGLQAKTAHRAAVARSVHVQ
PGFGIRLSVMPPSFSDSACKATLSNVRASAQPFIYRH

101 >eremothecium_cymbalariae
MERLSVSLGLRYVTVATLIFIAGIVSVILFLCLLGMLLVISIVMPSKAHIVLEHSSSDLA
SKELNTSVVEFQWRVELRSPFKEILARILRISTVLDAYCCTLGLHFSIFEILDSPGKYAN
SHVKENMNKTILESGIIYDPELDDSTEEKFTNIKTCHYTEAQRALLQFQRTKSEPIVFKY
GHPYTVDDFDDDFEGTFDLTILEDRCAKKQLDNDTEYDDFTVDSSGKSEELGTTDASTSN
FITHIMPKVNDRKQGYVYQLIPSKSSIISTALSHEQANAQPFLY

102 >eremothecium_coryli
MAPDHTPFMTLKHTVYSPFFLDTIWRKLEQTTPFSTLFWIFVILSASLFITAFLIFLIRI
VPEKTVKETTNGYDVEFVKESHQYAAHRYTEAQRAILQFQRKKQEPQLISYIHPYDEDGC
FDCCNPDLPSIVITPSVRCRLPSRRKYSNASKMHASSSIKKHESSKLDTNWASDNEVCHL
GLLKTSSTSKPDIDVRGLKLKKSFTGTESLEQPSILYPRTNSSGVKDTDISYEHAVALPF
RF

103 >eremothecium_sinecaudum
MISGLPNTSDIVRYTQIARLIILTLIIVVLFLLTLLIGIVLVHLSTLLSVLIQETAERWP
ELTSDLLCAKSLDINWTAVLRFLWAELKETVSGRVSSPTLILFGSSVYIYKAFKTRLSST
GSIATEGIKDFDEWEYHVNEQLGKQAGLYPEAQRARLQFQRNKFEPISFNYIEPYDEGDD
DFKNSSKTKSTSKKYKGRRGLNSFKKSSKTYTRKGASQNGGFDSSKKSSSSLSKDEEISR
KYKGSDTDPQKALVEYSSETKTLIYHAYPPKTAELKAQLTREQVTAQPFLY

104 >kluyveromyces_lactis
MVPESMAHRGWLEQARINFGELFQSIGNLVKTPNKPIMDKVADMDPHLIEDICSISCPNA
LTRPVKELEWITLYEVIEYKVKNKDPIILMIILITVPFLIYKIMKSAAYLLSAKNTKDFD
TTNIELNTEVTGDEVSTKPSHHFPAEQRALLQFQRNKSEPIVFRYGFNFEDNSSDMELPD
QSTEEVIRGSLNGTRTPSPMPSPPLPSNTFKDDDLRNMHSNILKLTETPQKFKDTSIVES
TASKLPGGMLIPLSISPSKPTINTTQVSSEQANAEPFKF

105 >Kluyveromyces_sp._yHMH660
MMLEAFKRYDWYRQLQVNFNELFNYLRSPTVSPNERYLDQIDQMEPQLIDFVCRTSCNRL
YDKALLYTEWAELFKVLEFKVKKHDPVLMAILAITLLIVIVSVTGLLRSKKTETIAAYKG
CVEEVDSIEIDTSMKVPHLPAEQRALLQFQRNKSEPIVFRYGFELDDSNLCSEINDQSSI
ESVRRSLDHNHSPSPITSPSPRPVTMSETAADTEQMHKRILKLTEPVTINSKITHLESPC
SQMTKTVPVPSHLSISRSKSSIRTTQISNEQVNAQPFSF

106 >kluyveromyces_dobzhanskii
MVRGDVLHSGWIEQAKINFSELFQSICDFARTPNTSSFKQTVDTNQVKPHLIESIHSTSC
HDLRKEYVEDTSWLTVYHILKYKAETKDPILLWIILTTALVVIYKVLKWHTSTKYITNLE
EIKHIATEPNAELVEEEFVTRPLVHFPAEQRALLQFQRNKSEPIVFRYGFNFDENKSDFG
LPDESTDELIRNSLSNTRTPSPLPSHSLSPRLFKETDVQDIHSNILKFTETPQKTKQLSI
LNSARSKISANLQLPLSISPAKPTVNTTKLSSEQASAEPFKF

107 >kluyveromyces_aestuarii
MSIAAVSTIENFYGQLQEQWIRLLAKLNKNSPIPNAFENHIGGRELNPDMVTYLCSTYCQ
KDEENLNWAVVMKFLLSKIRSSKFVIPVVALSLILLLLGILIKLDKKSTKYEPTSPSSEE
VIEEKEFGRERTHFPAAQRALLQFQKNRSDPIVFKYSYDLNEFDDIDTNGHICHEFKDNS
QDLNCQDVEFESLPSASPIQKRSNDVINITTPTKSPSGKTVGNKAVCTPRKSAQNLPILL
NPHLPLSVSPIKTSIQTTKLSSEQANAEPFKF

108 >Kluyveromyces_aestuarii_NRRL_YB-4510
MSIAAVSTIENFYGQLQEQWIRLLAKLNKNSPIPNAFENHIGGRELNPDMVTYLCSTYCQ
KDEENLNWAVVMKFLLSKIRSSKFVIPVVALSLILLLLGILIKLDKKSTKYEPTSPSSEE
VIEEKEFGRERTHFPAAQRALLQFQKNRSDPIVFKYSYDLNEFDDIDTNGHICHEFKDNS
QDLNCQDVEFESLPSASPIQKRSNDVINITTPTKSPSGKTVGNKAVCTPRKSAQNLPILL
NPHLPLSVSPIKTSIQTTKLSSEQANAEPFKF

109 >kluyveromyces_nonfermentans
MPISGFIVNDCWYGQLKFQWNKLLAKLVESRRTPNTLEKQIKIEDLSPDTITHFCSTYHK
ETTEDLDWPKVLQFVLHEINSLNFLSRLAALSIALVLLGMLIKQQHKKSVKHKAVTSFTE
EETEERIILKKSTQFTAAQKAMLQFQKNNSDPLVFRYSYGLCEFDDIETNEHVYQTFERK
RTKPSKFNYSQPDSLPRTTETMNYSDDVSYTTTPIKKHAKSGHLIDRVELCTPKYFRQNI
PHFTSTPLSLSVSPIKTSSQTTKLSSEQANAEPFTF

110 >kluyveromyces_siamensis
MSIAAVSTIGKFYEQLQEQWTRLLGKLNKNSPIPNALGNHIGGKELNPDMVTYLCSTYCH
KDEENLDWSMVLKFLLSEISSSKFVLPLVALSLILLLLGVLIKLDKKSTKDEAAPVSSEE
VIEDKELGKKRTHFPPAQRALLQFQKNRSDPIVFKYSYGLNEFDDIDTNAQICQQFKNQN
LQGSTCQDFEFETLPSASAIQTHSNDVINMTTPTKIQPDKAVGDKDLCTPKKSAQDLTIL
LSAPLPLCLSPIKTSIQTTKLSSEQANAEPFKF

111 >kluyveromyces_starmeri
MFDVITRNNWYPQLQANLQELLHAVRGSTASVNERYLEQIDQMEPQMIDFICRTSCNTLH
QRSQMYTEWSELLRILEYKAKNRDPILLGLVAIMLILGTVNVVLLSRSSVAEKRSQYVEE
TESPLQADPGHAVAHLPAEQRALLQFQRNKSEPIVFRYGFELDDSHLCTDLNDQSSVESV
RRSLDHNHSLSPVASPSPSPVAKSTHDTNLMHQKILKFTEELTEPSQHTLQQLLQDHAAV
HIPLLSISPSKASTETTQISIEQAKAQPFSF

112 >kluyveromyces_wickerhamii
MGFDNILSSQWFNTTPSTISDVFKRLFKFTKTPNTVHAEELLKLNPELINYICSKSCYRK
QYEPEDLDWRTVFDFLEHKMSSMNPLFVTLLISVILLLLHKYAFTVVNVSIDTFRDTNST
QSCTSSSNEIIEPKQLQYPAEQRALLQFQRNKSDPMIFRYSYDIGDTINETELTEDSDSE
LIGISLNDSGSLSPLPSPSPPSRIFSRDHSNHNYDRCQHAVKSNKQMKNAPRKLSQGTIL
KEQLPKVETPTLKEDMNHSKQTEPFAFTLSPAKSSTQTTKLSITQVNGEPFQF

113 >hanseniaspora_uvarum
MDYIKEQNNKNIDFFILVFIIASVYTYVNTKKDDRHSIKFTFKNQQIIMNLVDKKDEELY
VKPTLEDAEFEVQIDSEESFSSSDEDLYEPLINNETPVVKIISPSKSYKQSIQDMDLDHE
TEVINNEYYNESNVVNLLQNLQHEGDTLDESRHNSEKTEEIEDIFTLHPSSSKVLINDET
QFDNSTIKTIVSAGFDDSNTHNNNVDYALSTLDSREQQQPVKLLIKKFETISSASSVNSG
ENSFINKNILNKANTKSNETLKSAIVPILNYEDRIEVVENAIPKEPLNPELKTPVKRDSI
SDTISVHSKDSFNSNTSYKFFTPLIERNNKRGLSISKEGGSNFLKFKSGNENINFNDLTT
SSISKSVLTQEICATATIVLPEIVTSKAAVD

114 >hanseniaspora_pseudoguilliermondi
MDYLKEQNNKNIDFVILVVVIASLYTYINSKNDETKLLKFKFKNHDIVIDMVDKFEETDL
EQVDLTEPEFEAHITSDESVSSENEEEYNPIINMESPVVKIISPSISYKESIRDLELDNQ
NEIINNEYHSESNVMSLLQNLQNEDDTFDESATGSEKTDEVDFNAVHPLNSKLDAHDDIH
LDNSTIKTTDSICPEIESSIQQNSTDYALSTIESNKQQQPVKLLIRKFESISSESSVKSA
ENSFINKNILNSANTKSNETLRANVLPLEELKESIEIAEVPGEPLNTPSKEESKSDRLSV

NSKDSLSSHASNKFFTPLIERNNGRRLSTTQDGSNFLKFKSGNETINFSEITTTSISKSV
LTQEICATATMVLPEVGNASSAD

115 >hanseniaspora_clermontiae
MEYIKEQNNKNIGFVLLVLFVTSIYTYANSKKEDTRLLKFTFKDKQITINMVGDFNEKLH
VKPAVDRAAFHIQIESEEVSSSDDESLYEPQPNNEYPVVKIISPSKSYKQSIEDLGLEHG
TEIINSEYHTESSVVNLLQNLQQEEDPLDNSGASTEKTEDIEDFTAINPLNSKEIINDEF
QFDNSTIRTIGSADPAAQINTHSANLDYALSTVESREQQQPVKRLIQKFETISSESSVKS
VDNSFINKNVLNKGNTKSNETLRNTIVPLAIPEDCIEVVEDAIPKEPLNPELKTPVKKES
ESDVVSVNSAASNKFFTPLIERNKGRRLSTTKDVGSNFLKFNSGNENINFNNITTSSVSK
SVLTQEICATATIVLPEIATSTPVVN

116 >hanseniaspora_valbyensis
MGLIDKKFILFVLQQIITNIKMILFLLSVIAIAHILESLENNQLKKFLADKDIISFEFNG
KRIVIDLQDESSDSEDEKESEYEFKNIPHIDESMDMTSISEEDNDEEQESREISTPLIIE
NISPQRVFHSFDLNDMEKPEVIKNIYRSESSLLEALSIVKDEENTLINDTTTTTDHKDDN
NMEDNSSNTSQSTADCTNNSNATFIKNFKNTHVASLEPISSTENISNSCDSRSVSLSSDA
RESQLPVKDLILKFESIHSNSSSSSPENSFLNNNILNNGETKSDDTIIAKRTNNNGFDVN
TIEEVNSLHFYTPAKKELIHETNMIKNEIINSASSKASSNVFFTPYIQSSVIRPPGNSNG
ANYLVFKDDKKNIKAAAVARLIDSKSSCSVTKTVVTQKIVVNGTIIPPDASQ

117 >hanseniaspora_vinae
MTAFLNRWIQDQCLHFGTQRVDYLNERNGIKKLGTVDDKKLFVPVVFLITAFLLFHTRTT
KHSSPNTIYQTSNASFESTTSDIFFAKENEQHISNKPDEFADQEKDTTSLEETKVVTLDS
KTLSENSSQKIAEETDISPSSKKLDKKALQEKGTLIETCNTKPESLLDKCNSVLEIQDTV
TERVLNAEKVLEIEETSLVETAEITAASVPSSNGYTETPKDLPQHFDNKEIDQVSTPGKE
PLANTNITKQDTHDKISQIFRVHEEDIKSAITASEKDLQKTNKILESQELEPGANQKVMI
IDKDIAQPNNSSPLISEQEENKISLPVVNLLPKLRSKSSAGSLDSGNDSERTSKLSNLGS
IKALSSSESPKVKELIEIFENIKEPEIDIFSAHSTGSGGKGKATDSVAVPVLPSPSMFEG
NQPSSFSTSPRRNFNNNITLVSVSQASSQGSALSADLDKTPTKTGNSMGFSSVGLMTSTS
AHPSVTTAVNETPTKGAISTFTHKP

118 >cyberlindnera_suaveolens
MVETSKTIVDEDLLHQMESFVQENKLHMSDRFVQFYQEYVPGNLNEQLSHLLKLWKEHPS
YAFYKEYENSINTLIILIVFSTIGKLVSLFVDIYGAKQEELPQRFVLDKRYTEFETFPIC
TAKGDAEELLVLKSYEERHIELPDIDIEEVEYNISIVESPTPLNSIFIEDEEVLAAEPST
HIHVAVEETSDEILDMDILTSRSDDGGSGPTSSSSEENTKKINSKDGSRNARSSSSEDSF
EREEKLLDRTAVKMNEEKAYTPSKVSNHSFELSPSKKLVYKKSSSILTAFPSPSVISAET
TVGDITQETVYSQGFSSEDSTPSSLRRH

119 >cyberlindnera_saturnus
MVETSKKIVDEDLLHQMESFVQEHKLHLSDRLVQLYQKYVPENLNEQLSRLLKLWKEHPS
FAFYKDYENPINTLIILIGFSMIGKLVSLLVDICGAEPEELPQRVVLDKRYTDFETFPIC
TAKGDAEELLVLKSYEERHIELPDIDIEEAEYNISIVQSPAPLNFNFIEDEEAVVAEPSK
HIHVAVEETSDEILDIDIPISRSDEGSSAPTSSSSEEYTNKINGNDGSRSAISSSREDSF
EREEKLLDRSAVKMNEDKAYTPSKLSNNSFELSPSKKLVYKKSSSILTAFPSPSVISAET
TVGDITQETVYSQGFSSEDSTPSSLRRH

120 >cyberlindnera_misumaiensis
MEALPTTISQPSNHTVQLGDQISSFYHQHTSYHVTAKVLELYDAWTSHASYKVYKEYESF
ISGALILMGLVITYRVIHSFYKMFQQQEEPEVIVLDKRYTEFDTIPVCTGRGDMEELFML
RTYRETEIQLPDFDFDMSRLDGSTPRQVGTPELVRKILDDSEGDSITDVISLRHVKLAGV
RVQSSSPEGSFEREEKRVDKTAQLLNEERAFTPCKLRNSPMDFSPSKKLVFKKSSSIITA
FPQVTAESTVGDITTETVYSQSFSEDSANAGQ

121 >cyberlindnera_jadinii

MLLIGIFGVLITTLLVPLLMGVTDMDHLVHSLGSQLQRFIGPSDDIVHDKIAVAHTPLNL
THISVLDPMYSHGLGRKDLIHQVESLVQGVEEKLESRLSQCVAQFLPESMACKIHSVADA
WKQQRLYSVYKDYEALFKALALSAMGLLMLKILNLLLTLIVPKREHLEEVETPPVIDKRY
TDFETFPVCTGKGKGDAEEFLMLNSYKDNNIELPSYGTEDLSILGGSTPIKSSSIPVEVE
IVGCGDTGVPVHISESESEGEEEEEDDDDDDGEQRDISQEQGQRSNATQPILSLSSFEKV
EAQNDDLAMELNRARSLTPTRIDSGRKGTTELSPSRKLVYKKSSSILTAFPKVTSETTVG
DVTHETVYSQGFSSSGSGDSSPAPSKPGSHA

122 >cyberlindnera_fabianii
MIHNKQRGKEHVVEIYNQYVPQQVATYVNTSFSYWKTHSSYKVYREHEGVINNVLLVLGL
LITLKVIGSFLKLIFGPAALDDPDEDEMEKTVGAVVDKRGYIKKEYTEFETIPICVGKGD
EEEALVMQSYAKLRTQAGSESVILQELNKKNVDYRINGFTGPFGNGNGNGHKFELSIIDG
STPLRSTLEDPYTSTDTTSTSQSGLSSPMIAKQHDPFTVLMEEKRNKSIVSIQDVSYSGS
PISTKSRSTSPEDSFEREERVVDKLLSRLNEGESDRQVERPYTPSKLSMSSSLEYSPSKK
LVFKKSTSILTAHPITSVAETSVGSITKEDVYSKEFSSDHSSDILSPSLRKSL

123 >cyberlindnera_maclurae
MLMGLVITYRVIHSFYKMFLHKEEPEPVTLDKRYTEFDTIPACTGKGDMEELLMLRSYHD
MQIQLPGLGLDMSVLEGYTPKQMETPELVRKIAEDSENDSMTDASSLRHSQLLGGRLQSP
SPEDSFEREERRVDKSAQLLNEQRAYTPSKSPKSGADFSPSKKLVFKKSFSIITAFPQVT
AESTVGDITTETVYSRGLSEDSADAR

124 >wickerhamomyce_salni
MSDARIPSSRSNITFDPLMDTGRGTPYKVLNRDLVGDFETLLNQSKEAIVHTVSETYYNY
IPLHIREPLSQAYGSWKLHPSYNFYKNHELVFSFIFQTFAVFVVVKLLKSIVMSILSLFH
DPNPQIQAQPELVLNKSYLDFETIPLCVGKGDAEELLYIQTLQEQSILKHDDSYLTQFGI
SSDDTTANEEIVDSSTPLGLSAAVEITTLETNESLQLSEFSMSMNSEASSIYVSNPTSAE
STSKSTSPEDSFERQERVLDKKLARLNDTVTGSLTPKPSKVFNTMNTSPSRVLYYKKSTS
ILTTPRPKPTIGDITQATVYSEPFSSDDSLGSPISFKTRANGNLPRKPKMT

125 >wickerhamomyces_canadensis
MSDSRPQSSKSVDYLPFETIPMCVGKGDAEELLVLQSYEELSALKHDSKILKQFDIATEE
YRDEISSKLTSSTRISLDGPVDVTTLSTDSESLQISESTNNDHLDTDIQYKFPTSSSPSS
NSNDSPSPEDSFERQERKLDKNLSNLNRSNAPESVSMTPTSSSVLEFTMSTSPTRKLVYK
QSFSLLSGHSPTPASASATIGDITQETVYSEPFSSDDSASSPIRLKNSG

126 >wickerhamomyces_ciferrii
MNTNYIDNKDKFLGQSDLNVEELDINDVPLKIQTVAIDDELVVTAVDVASDAATTTPLPL
PLLILNKNPHYLLHDFKQFINESKSSIQLQFWEIHDKYSPQQLKSIEEQFLEFYKSEQVS
NFLDKYEKPLNNLFIGLTIIIILQIFKFFFNQFTKKPSKTIIPKEALTEKATKDALAGFK
DYNLETIPICIGKGDEEELQLLNSYTEILQNSNDEEILTKYGYNINDLINSSELNGLNSS
NDNSNLSDMFESTPLDIDEAPLDQSIDSNLNSNSNSSTGNSSNQNSSSNSPPVKSEDSFE
KHENLLDDEFSLRLINNSINLDSKSSKSSILEKFSTSPSKKLVYKKSTAILTNNAFPKES
VADTTVGDITTETVYSEPFTSDAETSFKNSL

127 >Wickerhamomyces_chambardii
MLLIPVLIGFIATTLFIPFLPQAVEYLEQQFPGLGSQLRHLLDQPNNASAKVSETVSFSE
SIILKTIADEHSTVVQTSTSSVYVTSFVTETPVYSTPEVQQSSVRQDFQLMINGIYDKIV
IPSVDYVLKKYNEYINEETKDTLQQYADKASSQGSILFNKSLEYGQEYAQLAYDQSQVYT
NKAYQLIVTEKTLFFYNLYSYEIKVLGLSFTIAFFLWLFFGRPSKNAARYQKNDTSNSDS
DLQKVCSINDKSEKEEGDILRQTNNLRTTKSNKEQELRPTSADFINNLRQINSKEEQKKA
KEGLDSLNSIALKADIVIAQVSPFEDLNLGKTNEIEIKTQELIEEDDEDEESSKLSLSIS
STTPSQGSSFERQESQLDNELDLLNKSATDTNFNNTQSINTKQRILNTKRSSSSIITSPS
PSKNIILKSSSASILSSSYTFPKLTYMKTLQNKNTISSLSLSNSNMGPLDDEFDNSVGDV
SIGEINQTNVYSQSFIFTHASDHDIQEQLSSLA

128 >wickerhamomyces_hampshirensis

MLALSIITAILATILLPLLTSVDPDDIGFYYNKFISGAPKSSVRMIEEAPVREADPHKEP
LTSFLLPSIDLVYEIESLVQENKEYLEDKLTEVYELLVPTTVKRNINRLYQSWRQHESYK
IYKAQEHKIFNLASLLLAILIIRRAKSIYNTIFISEKHLKKDESPPVVTIDARYTEFEVL
PPCVGRCDAEELLMIQSYDEMGISYADKNEEEQTVWDTEESMDQSNFDISILEGSTPQQV
NSPTENSSSDIQSPMPNGKKSLIVQKFLDDDANNDGKPEVHLTSSPFSSGSSKSGSTSPE
DSFEKLERQVDKSLSKLNENESYTPSRLSGNSSFNTYSPSKLSRNSSFNAYSPSKKLIYK
KSALILTNKKASSAKVTISAESTVGETTVGDVTEETVYSEPFSSDEHIVSPLRRKISIQK

129 >wickerhamomyces_mucosus
MIILPLIGFILSIVFLIPWLNEFVIDQVPLLKSSLIIRFINGSNIATSSDVSNESRNLID
NVISEENIIHPSSSTVLHLSNATINQIYNNAIDHVFIKELLRILNEKILIIKDKVNHIYY
NYIPVEISQYLDNLIRKSDPIIEYYLKNREIINNSISIIAIFSISKIIISKINKVKISKK
QKNANKTGEYFNIEEIERNYLTTFETIPMCKPNFNSEELLVLKSYQEIKNVDDGNTYDFT
ISSEISEEDTPSIFGIHKSNANSNSNSKSPSVGSQFESIDIIDTPSQTISKSPSPDDSFE
KAEKLLDKELSNLNMQEESEYQPSLSLSPSSLFKKNSNMSFNEIHSSSSKKLIYKNSFGI
LTSSVESGSEIFADSTSNVTEETVYSDFFPSSPSANKTII

130 >wickerhamomyces_piperi
MVTNESITTDKLIDWINNLDASTSDTSPLNQVPELITSTIEALKSTTALDAFQHDLKDEV
HTLLEPKLEIFKDFVDSKIEILGSQSTIKESDMYSTLLWFYTQYSDQINVVIMALAIAVA
IKLLRTLTFKLLGDYPAQNEGTYGVDDDIVSTEISMISQIEREYTQNFSVIPMCKPNYNR
EESLLDQLYENLGVDSGSESFLAGVVTDEASLETTAEEESLENTLSQALLTSQGLKHKTS
KQLVFKNSSSVLESPKKSNDKNCSPVINITEETVYSSDFSKSDTLQDSYLR

131 >Wickerhamomyces_anomalus
MPWISVIVNERMPQLSAQFSKFLSQPEPEPIIEKFIEENGGLENITGQHSSKLLNLNPHD
VLKDVNIFIGETKESLYDQWIVLYDKHAPVLLTDKVDSALSFIRSDKVWPYYEKYQKPID
NLLIGLAILILLRVVKVLARLVKPAKTIREIDEVTKVVLTEKDFETIPICVGKGDAEELL
LIQSYADILKSSNDEEVLKKYGYDKDFSDTNSKSDLSILNDFSPSRIDSPNDFTASDVSA
IFQTNQDIGLDAPVEIELRKGTESPSLEDSFERAENILDNKLESLNSSNMVVTPSSGGNS
IDQYSTSPSKLIYKKTSISILTNNTFPTTASVADTTVGDITEETVYSEPFAGDENTSFKD
SL

132 >komagataella_phaffi
MGAPLLALTIYYLILPLISFFAVRHLNLLDAPVNTPIKIAQKQDNTSLVELTYTDTRNLE
IRQPYQEIMEQLTFDSGFINCAGLIIFIAVLGVFFYYVSLLMLGGISDLYKGLNEQQKFR
TADLNPVNGSSIYQEQTFELGDHDVEVDKDPAVQERTKTDTSIRTTATETTPTPKSENGS
LNTSTDSLEKVLQSINTLDDDNDDDEGNDTPSKGLKPEFWGRNSMLQKYAERNPYTPRQS
LSPVRSVSPKKDKAESNEVSLKINQLANITRNSVCHSPGTSFASSVLSTTTAFPQTTVTH
INDDSASILEDIVLQQ

133 >komagataella_populi
MGVSLVALAIYYLVLPLISFFAVQHLKLLDAPADVPLKLTQKQSNTSLIELKDSNTQNFG
LHQPYQDIMEVLTIDSGFINCAGLVIFIAVLCVFFYYLSLLLLGGISDLYKGLNEQQKFR
TAPLSPVKESSIYQEQTFEYVDVDHEVEAEKDAAEQISATKTNNSSSQTTTAETTPTPKY
EEGSRNTSAASLERFMQSIDSSEEDSDADEGHNTPSNRLKPESWGRNSMLQKYAERNPYT
PRQSLSPVRSVSPKKGKAEPTEVSLKINQLANMTRSSVCNSPGTSFASSVLSTTTAFPQT
SVIQTNDDSASILEDIVLQH

134 >komagataella_pastoris
MGVPLLALTVYYMILPLISFFAIRHLNLLDAPVNSPMKIAQKQGNTSLIELTHTNTRNLD
IRQPYQEIMEQLTFDSGFINCAGLVIFIAVLAVFFYYVSLLMLGGISDLYKGLNEQQKFR
TADLNPVNGSSIYQEQTFEIVDHDIEVDKDIVMEETTKTDKTDTSSARTTATETTPTPKS
ENGSLNTSTDSLEKVMQSINTLDDDNDYDDDEGYDTPSKGLKPEFWGRNSMLQRYAERNP
YTPRQSLSPVRSVSPKKDKAESNEVSLKINQLANITRNSVCHSPGTSFASSVLSTTTAFP
QTTVTHTNDDSASILEDIVLQQ

135 >komagataella_kurtzmanii
MGAPLLALTIYYLILPLISFFAVRHLNLLDAPVNTPIKIAQKQDNTSLVELTYTDTRNLE
IRQPYQEIMEQLTFDSGFINCAGLIMFIAVLGVFFYYVSLLMLGGISDLYKGLNEQQKFR
TADLNPVNGSSIYQEQTFELGDHDVEVDKDPAVQERTKTDTSIRTTATETTPTPKSENGS
LNTSTDSLEKVLQSINTLDDDNDDDEGNDIPSKGLKPEFWGRNSMLQKYAERNPYTPRQS
LSPVRSVSPKKDKAEFNEVSLKINQLANITRNSVCHSPGTSFASSVLSTTTAFPQTTATH
INDDSASILEDIVLQQ

136 >komatagaella_mondaviorum
MGLSLVALAIYYLVLPLISFFAVQHLKLLDAPADVPLKLTQKQSNTSLIELKDSNTQNFG
LHQPYQDIMEVLTIDSGFINCAGLVIFIAVLCVFFYYLSLLLLGGISDLYKGLNEQQKFR
TAPLSPVQESSIYQEQTFEYVDVDHDVEVEKDTAEQMSITKTNNSSSQTTTAETTPTPKF
EEGSLNTSAASLERFMQSIDSSGEDSDSDEGYNTPSNRLKPESWGRNSMLQKYAERNPYT
PRQSLSPVRSVSPKKGKAEPTEVSLKINQLANMTRSSVCNSPGTSFASSVLSTTTAFPQT
SVIQTSDDSASILEDIVLQQ

137 >kuraishia_floccosa
MEPPYLLFAYYVMLPILSLFFIYDASIPGVDSFYKLIGFSSPSELVTISDSGLSISDAVG
IASAHTHPFFESPLYELYGSVHFYFTKYVLENELIGSVILQWCENVSPVAEWVSERLSPL
YYRFLAKYCAEASMFYDEHLFRLTAEYWPYIQQIGGLVLGFMLAMLLKDYIFRLVNSVMG
DVFDLSASINSQYSAKPTYGDLRDITDEQFVRALRSVDSEFVFQRSSIDQDDGLDEIIME
TITTTEDLNGNLKDVEVSFAKATEEDIDIGKVNRLVDLINKKMGSLNESEEEEEEEEDDE
ESQDATPESVMLNEDKIIDAALKSISESDSDNGGVFATPRKIDLDNHVTLSTSPKKINHS
NSFSSLSVSSRDNDKHGFKPYSDVYSSSPAYAPGTSRTTSTVSEENVLIAATSSSHSPSS
IRILSDINSGKLPI
